# Comparative Evaluation of Glycoproteomics Software for Rare Glycopeptide Identification

**DOI:** 10.1101/2025.07.10.664079

**Authors:** Hiroaki Sakaue, Kunio Kawanishi, Azusa Tomioka, Chiaki Nagai-Okatani, Hiroyuki Kaji, Atsushi Kuno

**Author notes:** Correspondence: Hiroaki Sakaue, Atsushi Kuno.

## Abstract

Advancements in glycoproteomics software have improved glycopeptide identification; however, algorithm differences cause discrepancies in identified glycopeptide, even when identical datasets. We compared five state-of-the-art glycoproteomics software programs (Byonic, MSFragger-Glyco, pGlyco3, Glyco-Decipher, and GRable), investigating their unique capabilities, and examined their ability to identify rare sialic acid–containing glycopeptides (NeuGc and KDN) derived from BJAB-K20 cells, which lack UDP-*N*-acetylglucosamine 2-epimerase, the rate-limiting enzyme for sialic acid synthesis. Approximately half of the identified glycopeptides were unique to individual tools. Byonic identified the highest number of glycopeptides, whereas Glyco-Decipher and GRable identified complex highly branched glycan structures. NeuGc- and KDN-containing glycopeptides were identified by specific programs, highlighting their capability to handle rare glycan structures. To assess the reliability of these identifications, we reanalyzed the MS/MS spectra for the presence of diagnostic ions corresponding to each identified glycopeptide. Some software programs identified glycopeptides without detecting the corresponding diagnostic ions, raising concerns regarding result reliability. However, leveraging the distinct capabilities of each software enabled us to achieve a comprehensive and reliable analysis of glycopeptides, including those with rare glycan structures. Combining multiple glycoproteomics software programs with complementary strengths and incorporating post-verification steps, such as diagnostic ion analysis, enhances the accuracy and depth of glycopeptide identification.

## INTRODUCTION

Glycosylation is a major post-translational modification that is crucial for cell–cell interactions and signal transduction. Cell surface glycans vary by cell type and condition, making alterations in protein glycosylation valuable biomarkers for several diseases, such as cancer and metabolic disorders^1, 2^. Glycan analysis primarily relies on released glycan assays^3–5^, where glycans are cleaved from proteins and analyzed using liquid chromatography or mass spectrometry, resulting in glycomics. An alternative glycomic approach, lectin microarray, enables simple and highly sensitive glycan analysis by inferring glycan structures via glycan interactions of lectins with different binding specificities. This technique has facilitated biomarker discoveries for various diseases, including cancer, metabolic disorders, and infectious diseases^6^.

Recent advances in mass spectrometry–based glycoproteomics have enabled the identification of glycosylation sites and detailed glycan structures because of marked improvements in mass spectrometer sensitivity, data acquisition speed^7, 8^, and specialized analytical software^9–25^. Glycoproteomics provides detailed insights into glycan structures and their attachment sites; however, it remains challenging. Misassignments remain a notable issue, underscoring the importance of selecting the appropriate software and analysis parameters. Kawahara *et al*. distributed liquid chromatography–tandem mass spectrometry (LC-MS) data to 22 institutions via the HUPO Human Glycoproteomics Initiative and compared the results of glycoproteomic analysis^26^. The number and glycan structures of the glycopeptides identified by each institution differed and included misassigned results. Additionally, the identification results differed with the analysis parameter settings, even when the same analysis software was used.

Since this report, studies have increasingly focused on comparing multiple software tools side by side^27–29^. A common finding across these studies is that each software tool, particularly its identification algorithms, has distinct strengths and weaknesses in peptide identification. The software reported to date is often classified into “peptide-first” or “glycan-first.” In peptide-first algorithms, peptides are identified first, and glycans are then determined based on the mass difference (mass offset) with the precursor ions. This approach allows the identification of glycopeptides, even when glycan-derived fragments are few or undetected. In contrast, glycan-first algorithms use specific glycan fragments (such as B-ions and Y-ions) to classify the spectrum as an *N*-glycopeptide and then identify the peptide by analyzing the consensus sequences. This method is advantageous when a few peptide fragments from the glycopeptide are available. No single approach is inherently superior, and recent studies have clearly demonstrated the benefits of using multiple software tools for a given dataset.

The concept of selecting or combining software with different algorithms according to individual analytical needs is becoming increasingly widespread. Although the use of multiple software tools enhances the comprehensiveness of glycopeptide identification, as pointed out in the HGI study, careful attention must also be paid to misidentifications.^30, 31^ These misidentifications may stem from differences in glycan libraries, scoring strategies, or the absence of key diagnostic fragment ions in the spectra. Such variability can obscure biological conclusions if they are not carefully validated. Therefore, incorporating post-validation steps such as manual inspection of diagnostic ions becomes essential to ensure the accuracy of glycopeptide assignments.

In this study, we conducted a comparative evaluation of five glycoproteomics software tools— Byonic, MSFragger-Glyco, Glyco-Decipher, pGlyco3, and GRable^32^—to assess their performance and unique capabilities. Upon integrating the identification results, we re-examined the MS/MS spectra to develop a workflow that ensures more comprehensive and accurate glycopeptide identification. Furthermore, we applied this workflow to glycopeptides containing NeuGc and KDN to enhance the identification of rare and biologically significant glycans such as NeuGc and KDN. The characteristics of the software employed in this study are summarized in Supplementary Materials. Peptide-first software, which does not require Y-ions for identification, includes Byonic and MSFragger-Glyco, whereas glycan-first software, which requires the presence of Y-ions for identification, includes pGlyco3 and Glyco-Decipher. GRable identifies the peptide portion using isotope-coded glycosylation-site-specific tagging (IGOT)^33^ and subsequently estimates the glycan using MS1 information. In this estimation process, the presence of Y-ions is not essential; however, because the identification in this study is based on the presence of Y0-ions, it is classified as glycan-first.

## RESULTS

### Overview of parallel identification using identical LC-MS/MS data

Glycopeptides prepared from BJAB-K20 cell membrane fraction were analyzed using four open-source software tools (Glyco-Decipher, MSFragger-Glyco, pGlyco3, and GRable) and the widely used commercial software, Byonic, following the experimental scheme in Figure 1.

**Figure 1.**
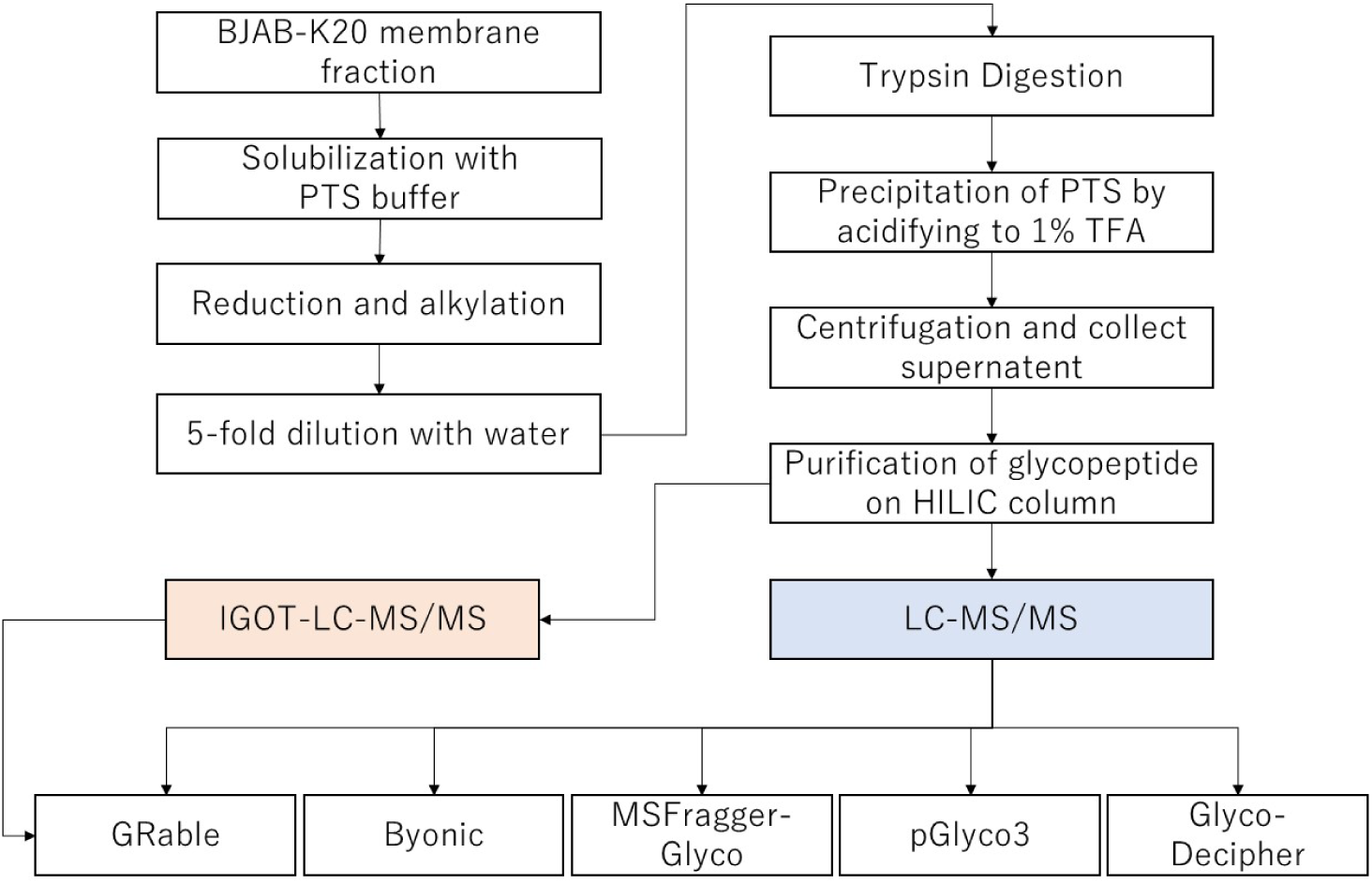
Experimental workflow for glycopeptide analysis using multiple software tools. Enriched glycopeptides extracted from BJAB-K20 cells were analyzed using nanoLC-MS/MS on an Orbitrap Fusion mass spectrometer. The obtained data were analyzed using glycoproteomics software.

The number of glycopeptides identified by each software package was compared. Considering that glycan databases are required for all four software tools except for Glyco-Decipher, a built-in library containing 132 human N-glycan structures provided by Byonic was used to standardize the experimental conditions. Figure 2A shows the number of glycopeptides identified using each software tool. In MS2-based analysis, Byonic yielded the highest number of identifications, followed by Glyco-Decipher, MSFragger-Glyco, pGlyco3, and GRable. GRable assigns glycopeptide signals from MS1 and subsequently reads the MS/MS spectra to validate the diagnostic ions. The MS1 signals included a total of 1,229 potential glycopeptide signals (green bar in Figure 2A), suggesting the need for further analytical sensitivity for MS/MS in glycopeptide analysis.

**Figure 2.**
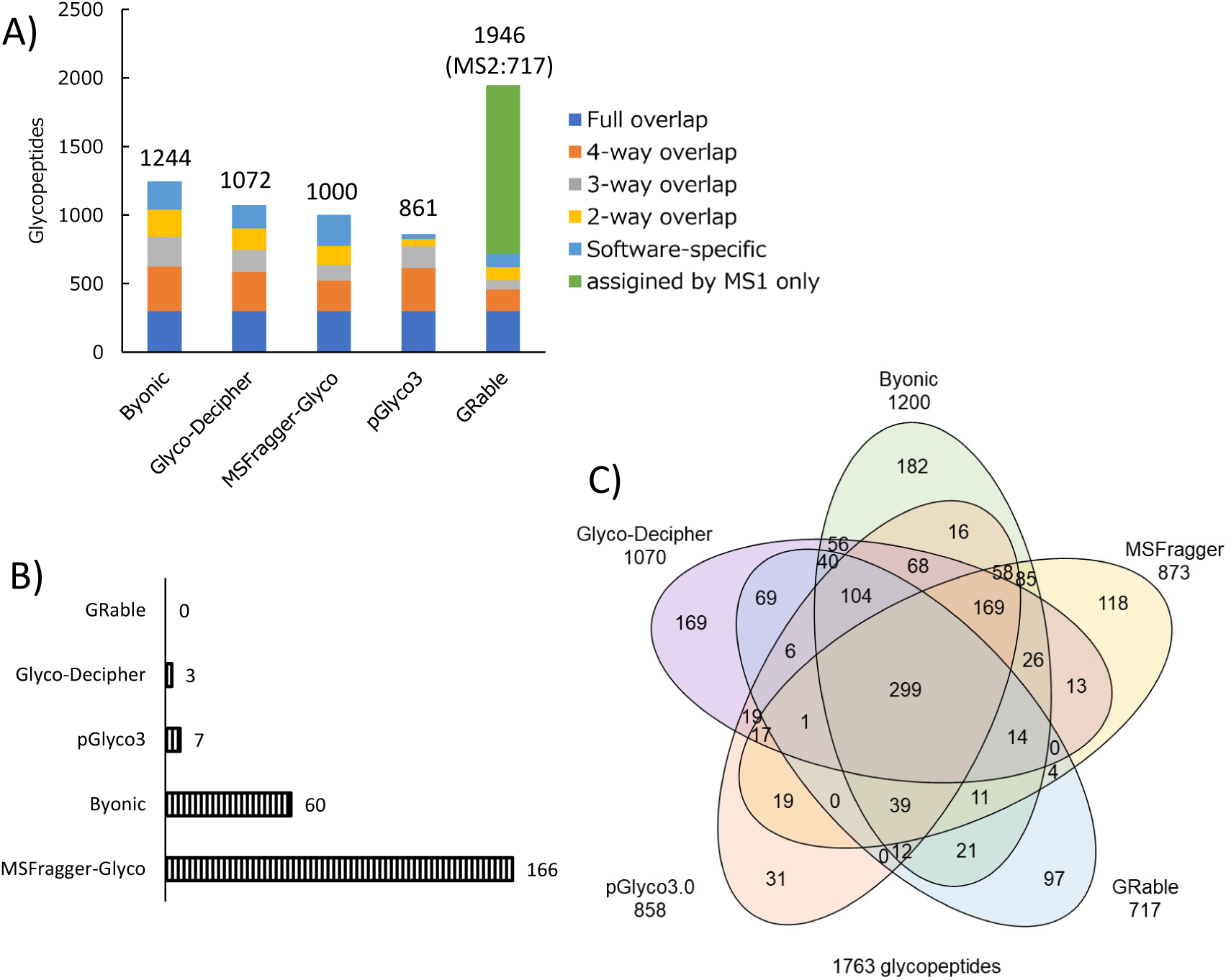
Comparison of the number of glycopeptides identified using five different glycoproteomics software tools. A) Number of glycopeptides identified by each software. The bar graph represents the number of glycopeptides identified by each software. Blue, orange, gray, yellow, and light blue indicate glycopeptides identified using all five, four, three, two, and one software tools, respectively. B) Number of glycopeptides identified without Y0/Y1 signals. C) Refined Venn diagram of glycopeptide identification after removing misidentifications.

Y0/Y1 ions have been reported to be highly useful for the accurate identification of glycopeptides^34^. Figure 2B shows the number of glycopeptide signals lacking Y0/Y1 ions identified by each software. Glycopeptides without Y0/Y1 ions were detected in 166 of 1,000 glycopeptides (16.6%) using MSFragger-Glyco and in 60 of 1,244 glycopeptides (4.8%) using Byonic. Similar identifications were rare (∼1%) when pGlyco3, Glyco-Decipher, or GRable were used. All the 203 MS/MS scans, assigned as lacking Y0/Y1 ions, were verified to confirm the presence of b- and y-ions. Consequently, 148 scans could not be confidently identified due to the absence of both b- and y-ions. The remaining 55 MS/MS scans detected at least one b- or y-ion (details shown in the Supplementary data, Table S1). Validation using the five characteristic HexNAc-derived signals^34^ (m/z 138.0555, 144.0661, 168.0660, 186.0766, and 204.0872) as diagnostic ions for glycopeptides confirmed 23 out of the 1,916 glycopeptides lacked the m/z 144.0661 signal. A method for distinguishing between GlcNAc and GalNAc has been developed based on differences in their fragmentation patterns at m/z 144 and m/z 138^35^. In α-GalNAc, the signals at m/z 138 and m/z 144 were detected with similar intensities, whereas in β-GlcNAc, the intensity of m/z 144 was substantially lower. Consequently, the signal at m/z 144 was more prone to loss than the other HexNAc-derived fragments. Among the remaining four HexNAc-derived fragments, any instance in which even one was missing resulted in insufficient signal information to confidently identify the glycopeptide, and these cases were excluded from identification. Based on these findings, we reviewed the identification results and established the following criteria for valid glycopeptide identification.

1. Glycopeptide spectra containing at least four of the five diagnostic HexNAc-derived ions
2. Glycopeptide spectra containing any Y0/Y1 or b/y-ions

Figure 2C shows a Venn diagram of the identified peptides refined based on this criterion. Of the 1,763 glycopeptides identified, 33.9% (597/1,763) were uniquely identified using the glycan analysis software. Only 17.0% (299/1,763) of the glycopeptides were commonly identified using all software packages. To evaluate the reliability of these results, we first investigated whether the HexNAc fragment signal, 204.0872 ± 0.05 Da^36, 37^, was present in the MS/MS scan corresponding to the identified glycopeptide. Using a VBA function in Microsoft Excel, we examined the scan numbers from the mgf file to verify the presence of the 204.0872 ± 0.05 signal in the corresponding fragment ions. HexNAc signals were absent in only two peptides identified using MSFragger-Glyco.

### Discrepancy in parallel identification

Among all software tools, Byonic identified the highest number of glycopeptides, missing only one glycopeptide, NGSAAMCHFTTK+ HexNAc(3)Hex(5), which was detected by all other tools. This peptide was identified in the 6283rd MS/MS scan. As shown in Figure 3, all the fragment ions of NGSAAMCHFTTK, from y1 to y11, were detected. Additionally, Y-series ions were observed, confirming the correct identification.

**Figure 3.**
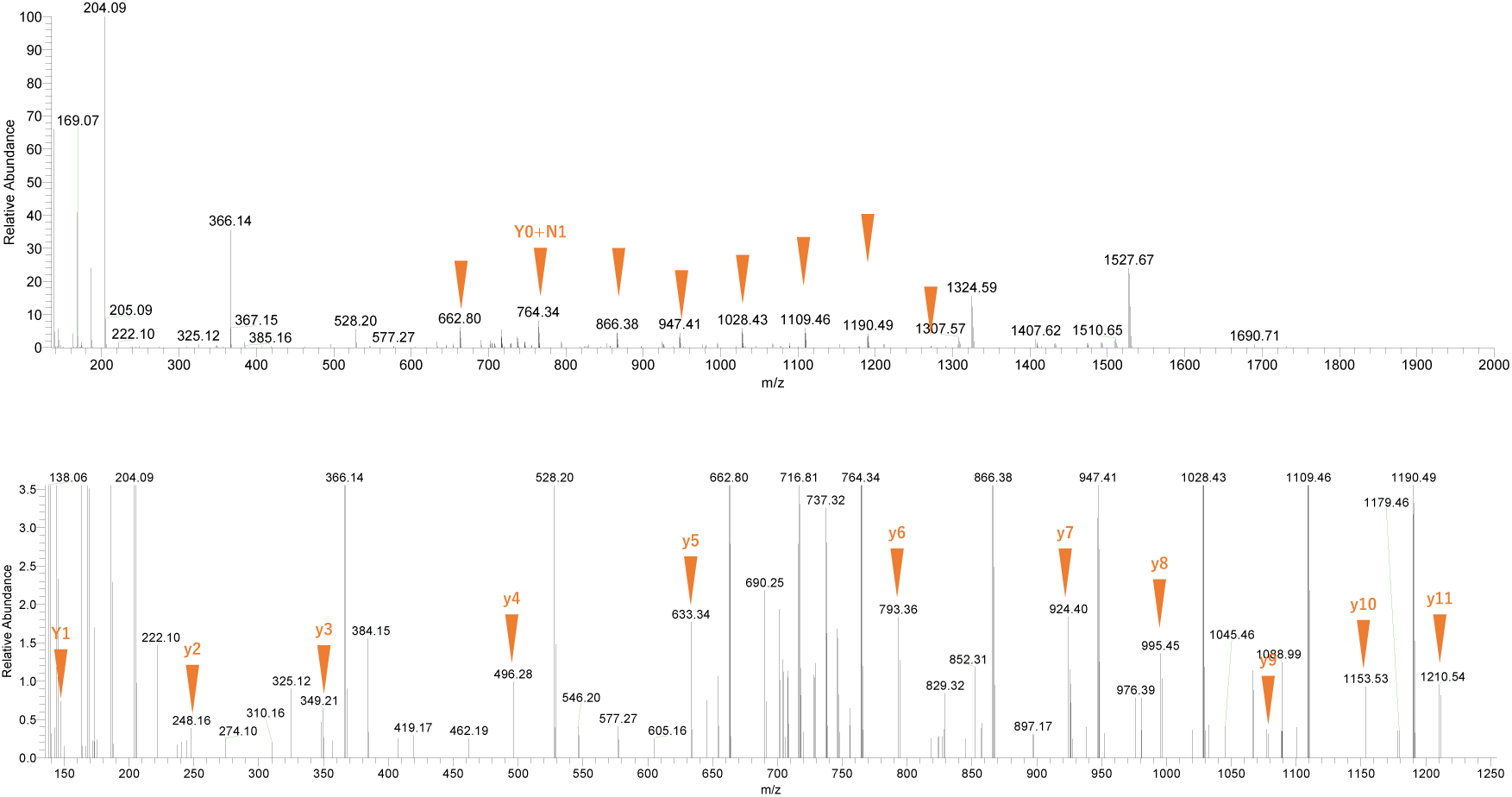
NGSAAMCHFTTK+ HexNAc(3)Hex(5) glycopeptide signal identified by all software tools excluding Byonic (scan no. 6283)

In contrast, scan number 6283 was identified as a glycopeptide with a completely different core sequence, GNWTTNEM(oxi)EVK+ HexNAc(3)Hex(5), by Byonic. However, the b- and y-ions derived from this core peptide were not detected, strongly suggesting misidentification by Byonic. The mass difference between NGSAAMCHFTTK+ HexNAc(3)Hex(5) and GNWTTNEM(oxi)EVK+ HexNAc(3)Hex(5) was 0.0065 Da, which likely contributed to the misidentification, as Byonic identified glycopeptides based on spectrum matching. The similarity in the Y-series ion masses potentially led to the incorrect assignment of the glycopeptides.

Furthermore, two additional glycopeptides were identified by Byonic using GNWTTNEM(oxi)EVK as the core peptide: HexNAc(3)Hex(6)Fuc(1) from scan 6350 and HexNAc(3)Hex(6)NeuAc(1) from scan 6893. These ions were misidentified because the corresponding b- and y-ions were not detected. This suggests that the misidentification issue is not an isolated case but rather a systematic limitation of Byonic’s spectrum-matching approach.

Several cases were observed in which different glycopeptides were identified in the same scan. In this analysis, 136 glycopeptides were identified in 86 scans. However, as mentioned above, this does not necessarily indicate misidentification by overlapping fragmentation patterns. Some of these examples are included in the Supplementary Information (Figures S1, S2, and S3).

Typically, a single peptide spectrum match (PSM) is generated from a single MS/MS scan. However, both pGlyco3 and GRable generated different PSMs from a single MS/MS scan. In pGlyco3, 10 glycopeptides were identified in five scans, whereas in GRable, 28 glycopeptides were identified in 14 scans. In GRable, if the retention time (RT) range and mass difference were within the set parameters for glycopeptide detection, the same scan was duplicated and clustered. Consequently, multiple PSMs can be assigned to a single scan. However, the specific algorithm used by pGlyco3 to assign multiple peptides from a single MS/MS scan remains unclear, as it is not well-documented. However, this approach appears useful for expanding the coverage and depth of glycopeptide analysis by capturing multiple plausible glycopeptide assignments that might be overlooked by other software.

As shown in Supplementary Figures 1–3, misidentification still occurs even when the FDR is set to < 1%, particularly in cases where the diagnostic ion signals are weak or ambiguous. We believe that the presence of diagnostic ions in the signal should be revalidated to achieve more reliable glycopeptide identification. Therefore, to enhance the reliability of glycopeptide identification, we verified the presence of Y0 and Y1 ions (univalent, divalent, and trivalent) in the MS2 scans of all the identified glycopeptides.

### Redundancy of core peptides

Figure 2C presents a Venn diagram showing the number of identified glycopeptides, and Figure 4A shows a Venn diagram illustrating the number of core peptides identified across the different software tools. Byonic and MSFragger-Glyco uniquely detected a significant number of peptide cores. These two software tools can be classified as peptide-first software, not requiring Y0/Y1 ions for identification. This strongly suggests that peptide-first (PF) approaches are advantageous for detecting a broader range of glycopeptide cores. Additionally, this result indicates that many glycopeptides lack glycan-derived fragments, highlighting the value of combining PF and glycan-first approaches to achieve more comprehensive glycopeptide identification. Our comparative framework illustrates this complementary nature in concrete terms.

**Figure 4.**
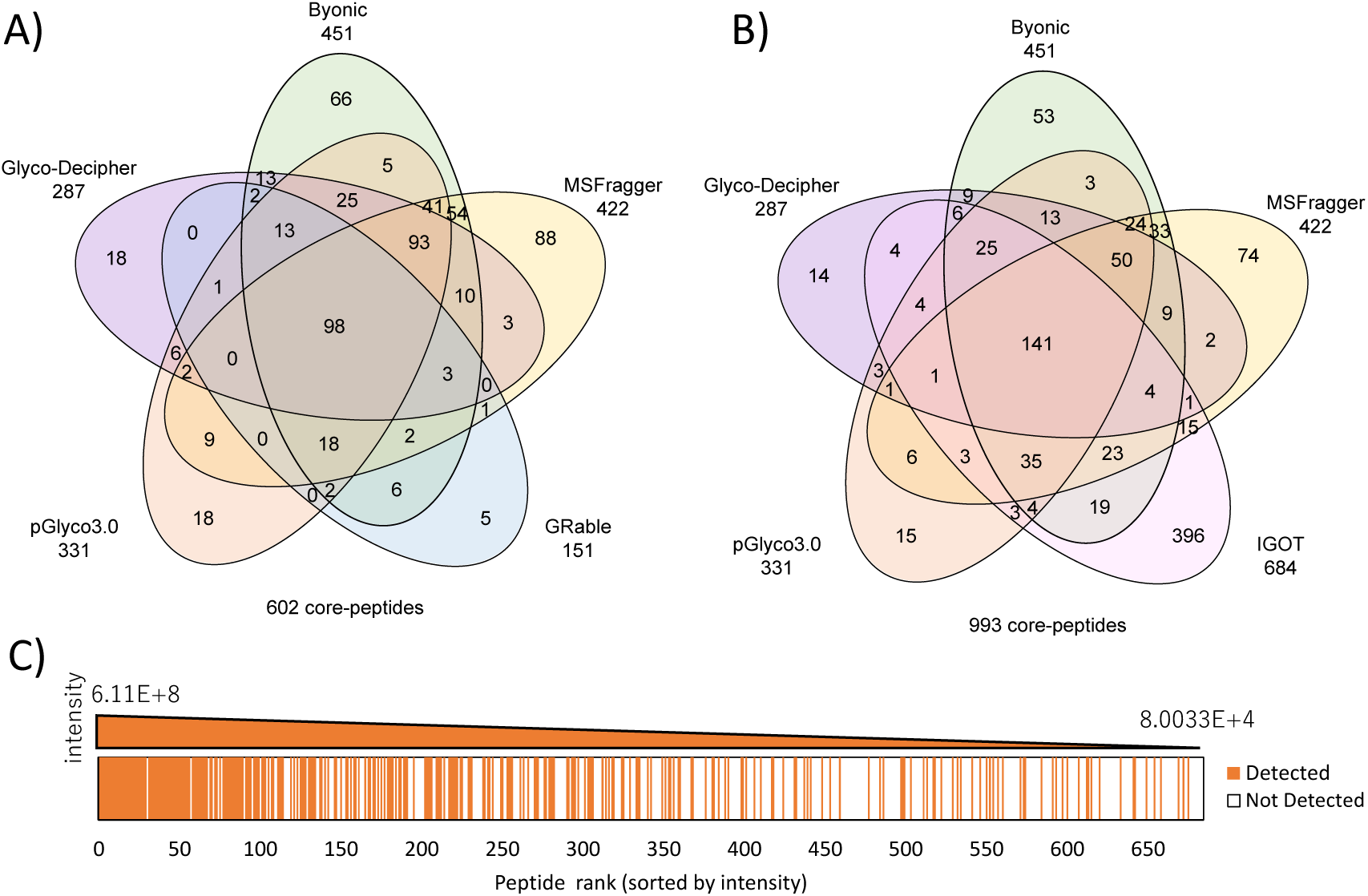
Comparison of the numbers of core glycopeptides. A) Venn diagram of the core peptides identified by each software. B) Venn diagram of core glycopeptides, including the results of IGOT-LC-MS/MS analysis. C) Intensity-based distribution of core peptides identified by IGOT-LC-MS/MS and other glycoproteomics software.

Based on the results shown in Figures 2C and 4A, the average number of glycan compositions (GC) identified per core peptide was calculated. The results show that Byonic identified an average of 2.76 GC, MSFragger identified 2.37 GC, Glyco-Decipher identified 3.73 GC, pGlyco3 identified 2.60 GC, and GRable identified 4.76 GC. These findings suggest that Glyco-Decipher and GRable can analyze a greater number of GC per core peptide than other software tools.

However, the total number of core peptides detected by GRable was notably lower (less than half the number detected by other software tools). In particular, GRable did not detect 93 core peptides, whereas the other four software tools did. The reduced number of core peptides identified in GRable is a key factor contributing to its lower overall glycopeptide identification compared with the other software programs.

As GRable performs its analysis based on core peptides identified by IGOT-LC-MS, it cannot analyze glycans attached to peptides not identified by IGOT-LC-MS. To assess the extent of this limitation, we investigated the overlap between the core peptides identified by IGOT-LC-MS and those identified by all software.

As shown in Figure 4B, IGOT-LC-MS detected 684 core peptides out of a total of 993 peptides. This coverage was up to twice as comprehensive as those detected by other software tools. However, IGOT-LC-MS failed to detect 309 core peptides. Notably, 153 of the 309 core peptides missed by GRable were identified by more than two software tools. This indicates that approximately half of the 309 missing core peptides were due to the limitations of IGOT-LC-MS data.

Table S4 provides detailed information on the 309 core peptides. Although many peptides not identified by IGOT-LC-MS are presumed to have incomplete glycan release by PNGF, no clear trends were observed in terms of peptide length or the positional distribution of glycosylation sites within the peptides.

As mentioned above, IGOT-LC-MS possibly achieved a more comprehensive core peptide identification because the deglycosylation by *N*-glycanase considerably reduced the complexity. However, even for peptides with glycans confirmed by IGOT-LC-MS, GC could not be assigned if the glycopeptide signals were too weak for analysis. This highlights the need for further improvements in the sensitivity of mass spectrometry.

Figure 4C shows the 684 core peptides identified by IGOT-LC-MS, sorted by intensity. The orange plots represent the core peptides for which glycopeptides were identified using at least one of the five software tools. The clustering of IGOT-identified glycopeptides at higher intensities suggests that low-abundance glycopeptides are more challenging to identify.

### Decreased coverage because of clustering conditions in GRable

In GRable, glycopeptides with different GC on the same core peptide are clustered based on differences in elution time and mass. In this analysis, the minimum cluster membership was set to three, implying that glycopeptides with fewer than three structural variations were not detected.

Among the glycopeptides identified in this study, 140 glycopeptides (corresponding to 99 core peptides) had less than two glycan structural varieties and could not be identified by GRable because of the clustering threshold.

For example, in this study, five types of high-mannose-type glycans, ranging from HexNAc(2)Hex(6) to HexNAc(2)Hex(10), were identified on GNCSLVIR peptide. However, the monoisotopic mass corresponding to pep+HexNAc(2)Hex(8) was not picked up by GRable’s internal processing. As a result, the software failed to detect three consecutive glycopeptide variants, preventing their identification.

This issue could be addressed by reducing the minimum cluster membership to 2, which would allow for the detection of such glycopeptide clusters. However, lowering the threshold also increases the risk of clustering signals that only coincidentally match in mass, potentially leading to false positives.

### Characteristics of glycopeptides identified by each software

The characteristics of the glycopeptides identified using each software are shown in Figure 5. The identified glycopeptides were categorized as high-mannose/pauci-, hybrid, or complex glycans. Among these tools, MSFragger-Glyco exhibited a notably high identification ratio for high-mannose/pauci-glycans.

**Figure 5.**
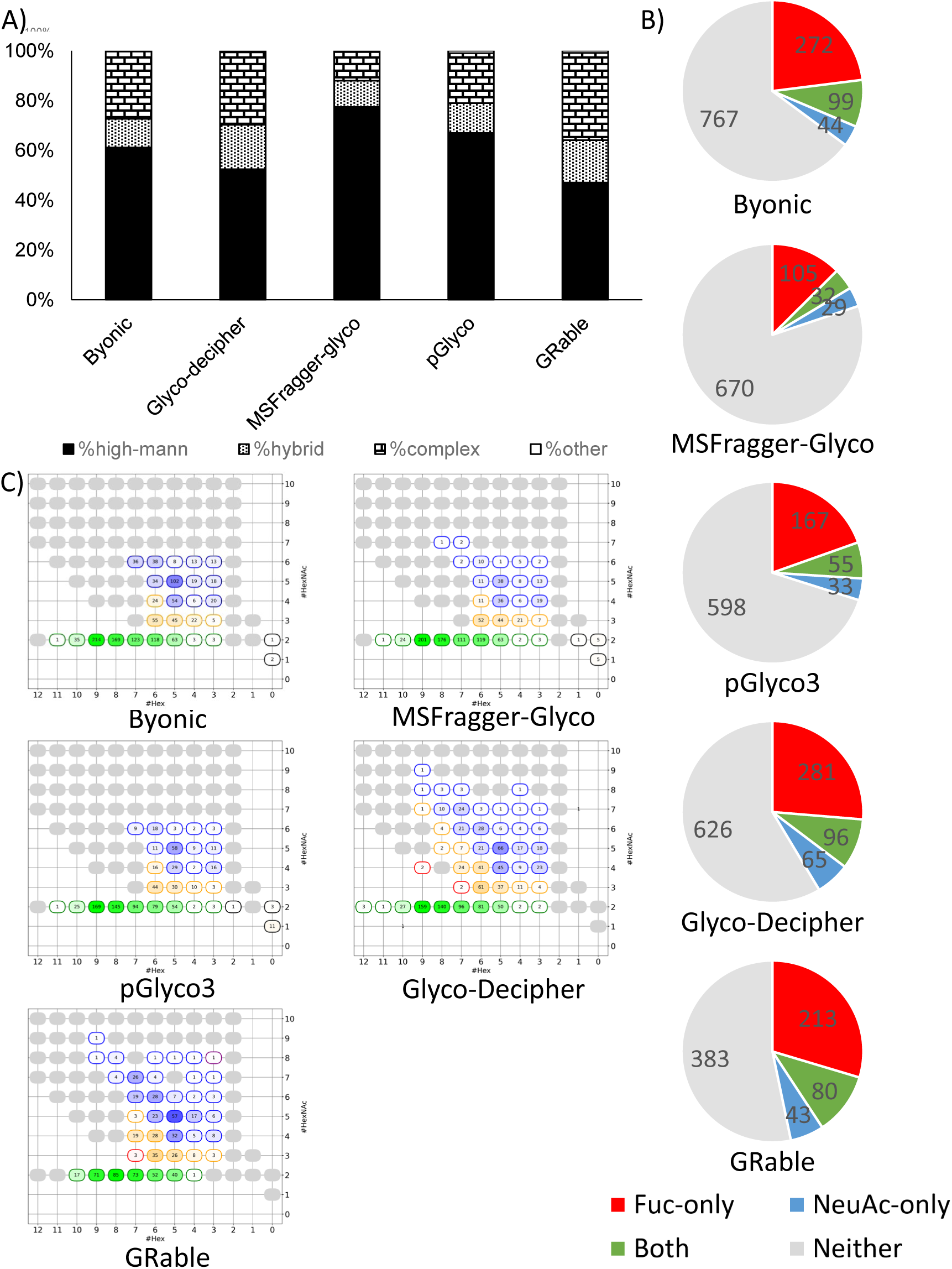
Characterization of glycopeptides identified by each software. A) The proportion of glycan types identified by each software. B) Proportion of glycans containing fucose and sialic acid in their structures. The numbers inside the circles indicate the number of identified glycopeptides. C) Diversity of glycopeptides shown using the Glyco-stem matrix^38^. The numbers inside the circles represent the identification count, with green indicating high mannose-type, orange indicating hybrid-type, and blue indicating complex-type. The intensity of the color reflects the number of identified peptides.

As shown in Figure 2, MSFragger-Glyco could not identify 104 glycopeptides that were detected using other software. Further analysis revealed that 76 of these 104 glycopeptides (73%) consisted only of Hex and HexNAc, suggesting difficulties in identifying glycopeptides containing fucose and sialic acid.

In contrast, Glyco-Decipher and GRable demonstrated high identification rates for hybrid and complex glycan structures. Glyco-Decipher utilizes de novo analysis via monosaccharide stepping, allowing the identification of glycans not listed in the glycan database. This functionality has been shown to be effective for identifying complex glycan structures. Similarly, GRable can analyze glycans beyond those listed in the database.

Among the glycopeptides identified exclusively by Glyco-Decipher or GRable, 82 of the 169 glycopeptides identified by Glyco-Decipher and 45 of the 99 identified by GRable contained glycan structures that were not found in the database. Furthermore, of the 69 glycopeptides identified by both tools, 51 contained non-database glycans. Although Glyco-Decipher occasionally misidentified glycan structures, such as HexNAc(1)Hex(10) and HexNAc(7)Hex(1), incorrect identifications were rare.

A visual representation of glycan identification provided further insights. Previously, we proposed the concept of Glyco-stem, which categorizes glycan types based on the number of Hex and HexNAc. This format enables visual evaluation of elongated glycan structures. The plot indicates that Glyco-Decipher and GRable are more effective in identifying glycopeptides with extended glycan structures than other tools.

In contrast, Byonic, MSFragger, and pGlyco3 are limited to detecting glycan structures listed in the glycan database. Among these, Byonic exhibited a balanced identification of all glycan types, whereas MSFragger and pGlyco3 showed a bias toward high-mannose glycans.

Figure 5B shows a pie chart that categorizes the GC into four groups: i) those containing only fucose, ii) those containing only NeuAc, iii) those containing both fucose and NeuAc, and iv) those containing neither. As shown in Figure 5A, MSFragger-Glyco and pGlyco3, which primarily identify high-mannose-type glycans, exhibited a low identification rate for glycopeptides containing fucose and NeuAc. For instance, fewer than 30% of glycopeptides identified by these tools included fucose or NeuAc. This suggests a potential limitation in detecting more complex or diverse glycan structures.

In contrast, Glyco-Decipher and GRable exhibited significantly higher identification rates for glycopeptides containing fucose and NeuAc. Approximately half of the glycopeptides identified by these tools contained these glycan components. The ability of these tools to identify complex glycan structures may be attributed to their advanced algorithms, which analyze GC beyond the constraints of predefined glycan databases.

### Comparison of the analytical depth of each software

In glycopeptide analysis, two key factors must be considered: (i) the number of glycans that can be analyzed for a single peptide (depth) and (ii) the total number of glycopeptides that can be identified (coverage). To evaluate depth, we selected the top nine highly heterogeneous glycopeptides with a common core peptide, identified by all software tools, and compared their identification counts (Figure 6). Despite some variation, the results showed that the analytical depth increased in the following order: Glyco-Decipher = Grable > Byonic > pGlyco3 > MSFragger-Glyco.

**Figure 6.**
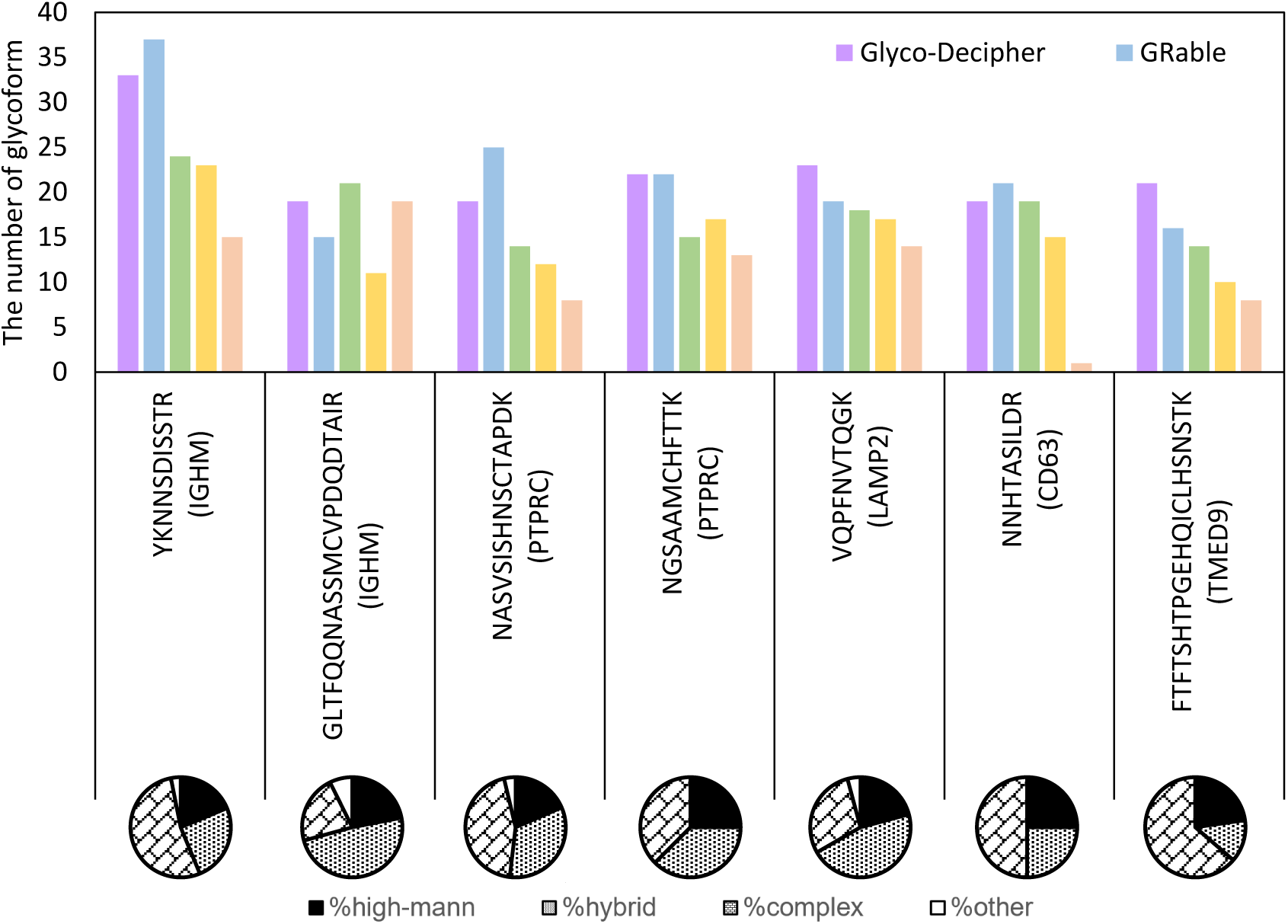
Glycoproteomics software for deep analysis of the heterogeneity of glycans attached to a single peptide. The bar graph shows the number of glycan varieties identified on a single peptide by each software. The pie chart below displays the types of glycans attached to the single peptide.

The pie chart below the bar graph illustrates the distribution of the identified glycan types. Glycopeptides with high structural heterogeneity tend to contain abundant hybrid and complex glycans, emphasizing the importance of analyzing these glycan types to achieve a high-depth glycan analysis.

As Glyco-Decipher and GRable can analyze glycans beyond those listed in the glycan database, their de novo analysis capabilities strongly contribute to their superior identification depth compared to other tools.

### Analysis of glycopeptides from MS1 signals

As mentioned previously, GRable extracts potential glycopeptides based on three types of information: mass difference, elution time difference, and de-glycoproteomic information. Subsequently, reliable peptides are annotated using MS2 data.

Based on the parameters used in this study, signals were clustered if the mass difference between them matched the expected monosaccharide mass and the elution time difference fell within a predefined range (e.g., -1 to +0.1 min for neutral monosaccharides and 0 to +3 min for acidic monosaccharides).

The number of glycopeptides predicted by GRable from the MS1 data reached 1,946, which considerably exceeded those identified by the other four software in this study. However, only 717 of these were identified via MS/MS, highlighting that although a large number of glycopeptides can be predicted from MS1 data, the depth of glycopeptide analysis remains insufficient.

Glycopeptides predicted from MS1 data can be categorized into two types based on their reliability.

1. Clusters where most signals have MS/MS data, but some lack MS/MS data.
2. Clusters where most signals lack MS/MS data, and the predictions are based on limited supporting data.

Figure 7A illustrates a cluster of glycopeptides with the core peptide NASVSISHNSCTAPDK. Toward the center of Figure 7A, a group of signals corresponding to tri-antennary glycan structures can be observed. Purple lines extending to the right of these signals indicate glycopeptides with attached sialic acids. Although MS/MS data were not available, the variation in the glycans comprising the cluster strongly suggests the presence of sialic acid– containing glycopeptides. Given that the addition of sialic acid often reduces ionization efficiency, the signal intensity may have been insufficient to acquire MS2 data. Thus, GRable predicts glycan composition related to other glycan structures even for signals lacking MS2 information. Although the detected glycan structures cannot be verified directly, the ability to identify glycan structures that are potentially present is a significant advantage of using MS1 data for glycopeptide detection.

**Figure 7.**
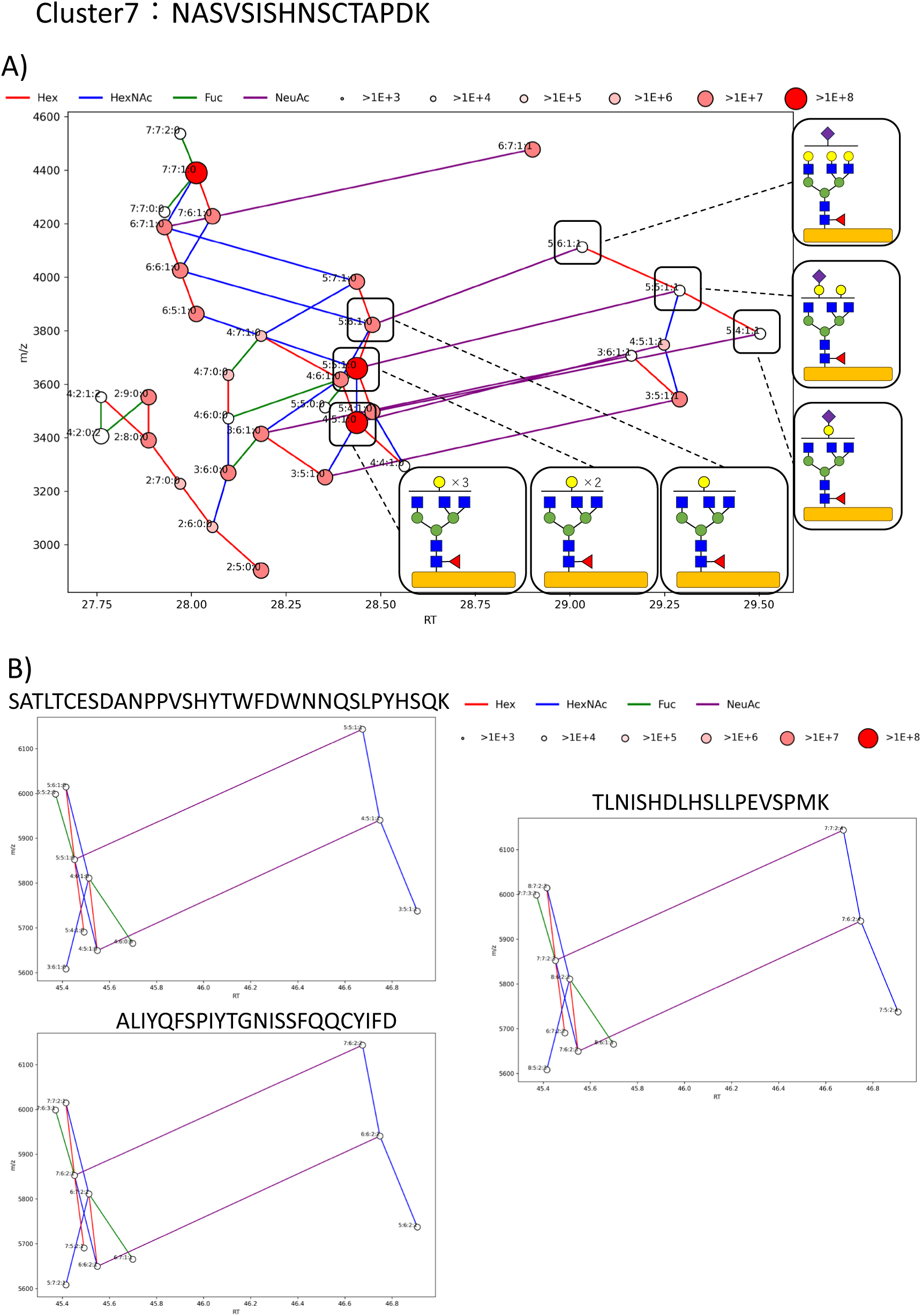
Structural prediction of glycopeptides lacking MS/MS signal acquisition. A) Cluster containing multiple glycopeptide signals. B) Clusters formed based only on the RT and m/z of the signals. Red circles indicate signals with MS2 information where Y0 or Y1 ions have been detected. The presence of Y0/Y1 ions in most signals within the cluster suggests that these signals correspond to glycopeptides with the same core sequence. In contrast, white circles represent signals without MS2 information.

### Clusters with limited MS2 information

Some clusters consisted entirely of signals lacking MS2 information. Figure 7B illustrates the predicted glycopeptide clusters based on mass, RT, and deglycoproteomic information. Three clusters with different GC and core peptides were predicted:

1. TLNISHDLHSLLPEVSPMK+HexNAc(7)Hex(6)Fuc(2)NeuAc(4): This structure suggests the presence of two fucose residues and complete capping by sialic acid; however, it is unlikely to exist.
2. SATLCESDANPPVSHYTWFDWNNQSLPYHSQK+HexNAc(4)Hex(5)Fuc(1)NeuAc(1): This represents a biantennary glycan with one sialic acid and core fucose. However, peptide+HexNAc(4)Hex(5)Fuc(1), which lacks one sialic acid, likely corresponds to a typical biantennary glycan. As Hex cannot attach to this glycan unless α-Gal is considered, this cluster is also unlikely to exist. On the other hand, if this composition is interpreted as a hybrid-type glycan, structures with up to HexNAc(4)Hex(7) are also plausible. Furthermore, considering the possibility of LacDiNAc, a HexNAc(5)Hex(5) structure could also be possible.
3. ALIYQFSPIYTGNISSFQQCYIFD: This cluster includes various glycans forming tetra-antennary structures. However, the composition HexNAc(5)Hex(7)Fuc(2)NeuAc(2) is not feasible unless α-Gal is considered. While these structures are theoretically possible, their high degree of fucosylation and sialylation makes their actual presence unlikely.

In recent years, MS/MS scan speeds have increased significantly, with the latest instruments capable of acquiring data at 300 Hz while maintaining excellent sensitivity. This advancement has enabled the identification of over 5,000 proteins in single cells^39^. Such improvements in mass spectrometry technology will greatly enhance the depth and comprehensiveness of glycopeptide identification using MS2-based software, reducing the number of glycopeptides predicted solely based on MS1 and RT, as predicted by GRable. This enhancement would strengthen the evidence-based validity of the GRable. One of the key advantages of GRable is its ability to predict the presence of glycans even without access to high-end mass spectrometers. Therefore, selecting and combining software based on available resources and the desired level of identification can be highly beneficial.

### Identification of NeuGc- and KDN-containing glycopeptides

As NeuGc and KDN are considered potential substitutes for NeuAc, we created new GC by individually replacing NeuAc with NeuGc or KDN within the glycan structures listed in the human 132 *N*-glycan dataset of Byonic. These new compositions were then added to the original database, resulting in a glycan database consisting of 500 GC containing NeuGc and KDN.

Among the five software tools compared in this study, four tools (Byonic, MSFragger-Glyco, pGlyco3, and GRable) could analyze NeuGc-containing glycopeptides in principle. Three tools (Byonic, pGlyco3, and GRable) could analyze KDN-containing glycopeptides. Glyco-Decipher can identify glycans not listed in the database via monosaccharide stepping. However, NeuGc- and KDN-containing glycopeptides were not detected. This is because Glyco-Decipher identifies unknown glycans by calculating the gaps between the Y-ions. However, sialic acids are typically found at terminal positions, making them unlikely to be detected via Y-ion gap calculations. MSFragger-Glyco does not register KDN as a monosaccharide; therefore, glycopeptides containing KDN could not be analyzed, even when these glycans were added to the database.

For glycopeptides containing sialic acid, MS2 spectra showed signals corresponding to constituent monosaccharides, their dehydrated forms, and non-reducing terminal HexNAc+Hex+sialic acid. These signals could be used as diagnostic ions for glycan identification. Figure 8 shows the MS2 spectra of a glycopeptide in which a hybrid-type glycan was attached to the peptide, YKNNSDISSTR. The terminal glycan was replaced with three different glycan components: NeuAc, NeuGc, and KDN.

**Figure 8.**
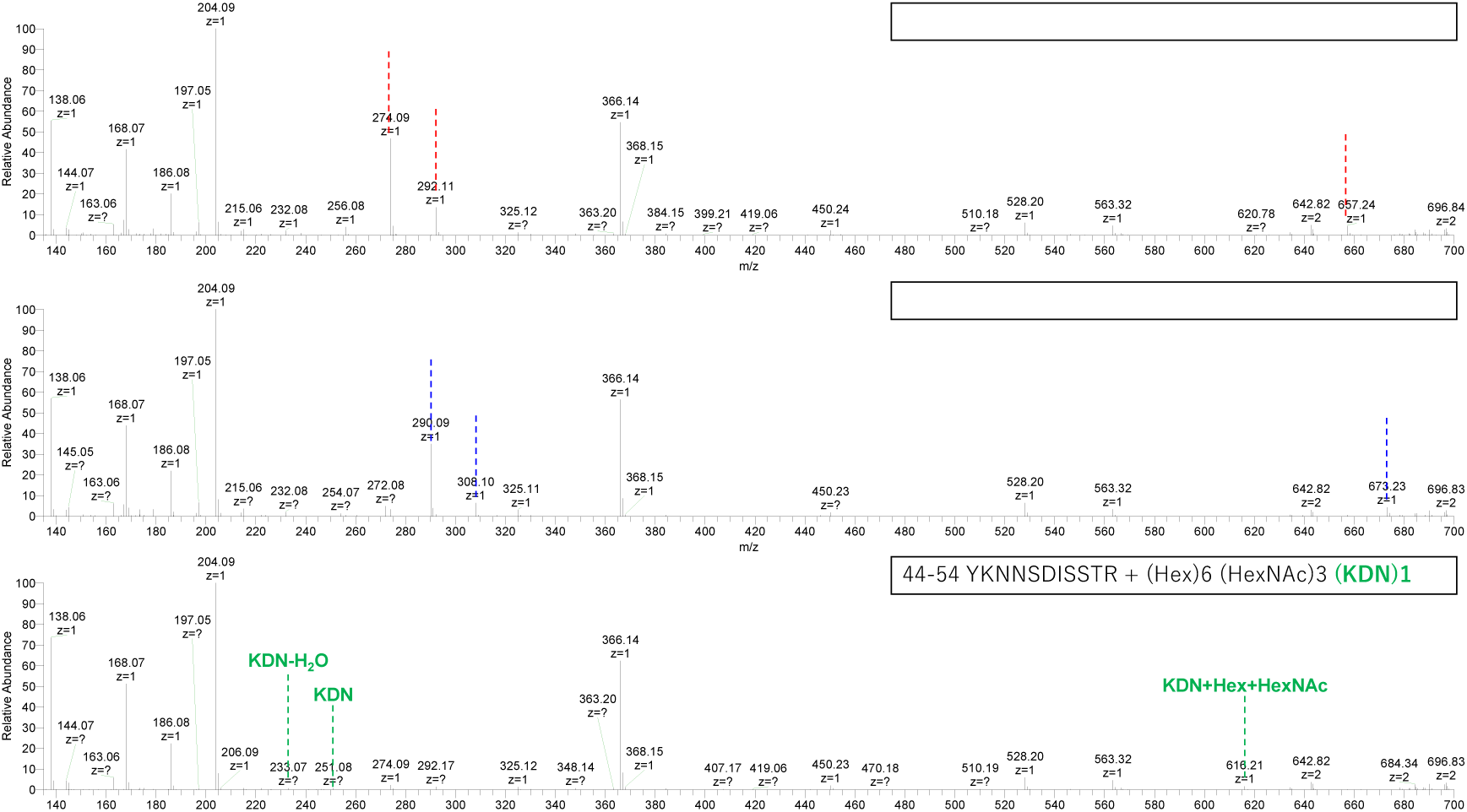
Differences in diagnostic ions by the presence of sialic acid

Figure 9 presents a Venn diagram showing the glycopeptides containing NeuGc and KDN identified by each software tool. The criteria used for peptide identification were the same as those used for NeuAc-containing glycopeptides. Although glycopeptides containing NeuGc and KDN were identified, their numbers were lower than those containing NeuAc. The lists of identified glycopeptides containing NeuGc or KDN are provided in Tables S2 and S3 in the Supplemental Information.

**Figure 9.**
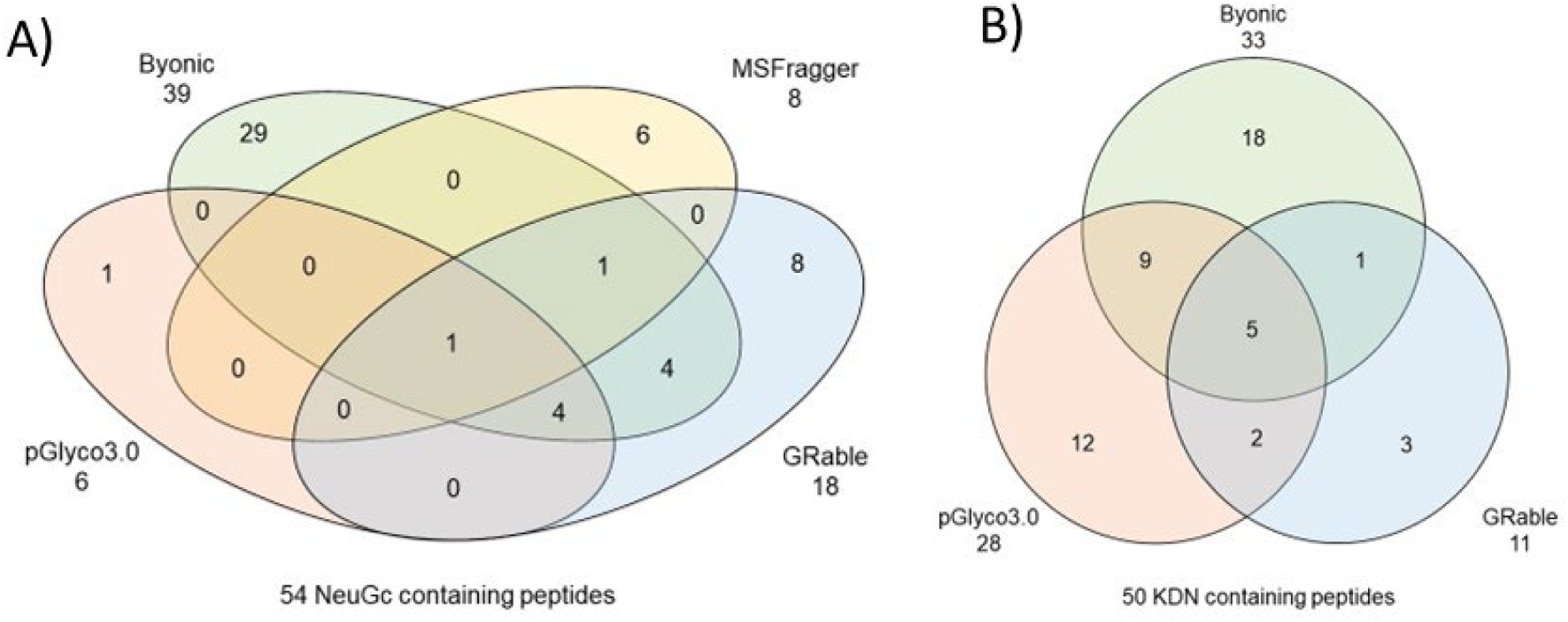
Venn diagram of the number of glycopeptides. A) Number of glycopeptides containing NeuGc. B) Number of glycopeptides containing KDN.

Although many membrane proteins were identified in this study because of the extraction and analysis of membrane protein fractions, the secreted protein, IgM, was also identified as a carrier protein for NeuGc and KDN. The carrier proteins of NeuGc and KDN were almost identical.

The BJAB-K20 cells used in this study lacked UDP-*N*-acetylglucosamine 2-epimerase, preventing the synthesis of sialic acid^40^. Therefore, we expected NeuAc-containing glycopeptides to be scarce. However, most glycopeptides contained NeuAc, and the levels of KDN-containing peptides and NeuGc-containing proteins identified were similar.

This study was conducted with 10% FBS added to the cell culture. It is presumed that free NeuAc and NeuGc present in bovine serum contributed to glycan synthesis. Despite this, our results demonstrated that glycoproteomics software can successfully analyze glycans that are rarely used in human cells, such as NeuGc and KDN.

## DISCUSSION

In this study, we compared five glycoproteomics analysis software tools: Byonic, MSFragger-Glyco, pGlyco3, Glyco-Decipher, and GRable. A custom program was developed to systematically evaluate the validity of all glycopeptide identifications by analyzing the MS/MS spectra obtained from each software. The presence of Y0/Y1 ions strongly supports glycopeptide identification because most of the identified spectra contain these ions. However, some software programs produced spectra lacking Y0/Y1 ions. In many such cases, b/y-ions indicative of the presence of peptides were also not detected, reinforcing the idea that Y0/Y1 ions provide strong evidence for glycopeptide identification.

pGlyco3 treats spectra containing the m/z 204 signal (indicative of HexNAc) as glycopeptides and deduces glycan structures on peptides by subtracting the mass of Y-ions from the precursor mass. In contrast, Glyco-Decipher virtually removes glycan-derived signals to identify the core peptide and subsequently assigns Y-ions to determine the glycan composition. GRable examines MS2 spectra derived from MS1 signals clustered as potential glycopeptides and confirms the presence of Y-ions and oxonium ions. As shown in Figure 1B, very few glycopeptides identified by these three software tools lacked Y0/Y1 ions.

PF algorithms, such as Byonic and MSFragger, yielded numerous spectra exhibiting b/y-ions and HexNAc-derived signals, despite the absence of Y0/Y1 ions. As shown in Figure 3A, both software programs uniquely identified a substantial number of core peptides. These results suggest that PF algorithms may have an advantage in the detection of unique glycopeptide cores. Thus, integrating different analytical approaches is crucial for achieving more comprehensive identification.

Our findings highlight the importance of post-validation in excluding unreliable identifications from large datasets obtained using glycoproteomic software. Even when the FDR is set to below 1%, false identifications cannot be completely eliminated. To address this issue, we proposed criteria for the post-validation of identified peptides by reanalyzing their MS/MS spectra, ultimately leading to more accurate glycopeptide identification.

Furthermore, we explored the potential for the comprehensive identification of rare glycans using glycoproteomics software. KDN is a type of sialic acid, structurally similar to *N*-acetylneuraminic acid (NeuAc) and *N*-glycolylneuraminic acid (NeuGc) and classified as an acidic sugar. NeuAc and NeuGc are the predominant sialic acids in animals, with their relative proportions varying by species and tissue type. NeuGc is biosynthesized from NeuAc by the enzyme CMP-*N*-acetylneuraminic acid hydroxylase (CMAH). However, in humans and birds, the CMAH gene is inactive, preventing endogenous NeuGc production. NeuGc found in humans is primarily diet-derived, originating from red meat (e.g., beef or pork), and is recognized as a foreign substance by the human immune system. This may trigger immune responses, such as antibody production, and has been implicated in vascular diseases and cancer^41^.

Seo *et al.* developed a highly sensitive method for detecting NeuAc and NeuGc, revealing that serum NeuGc levels in healthy humans are approximately 1/10,000 of NeuAc levels^42^. Their study also suggested that sialic acids exist not in free form but as components of proteins and lipids. Similarly, KDN was initially identified in rainbow trout eggs and was long considered to be absent in humans. However, recent studies have demonstrated that KDN is produced in humans as a metabolite of mannose^43^. Mammalian complex lipids containing KDN constitute less than 1% of total sialic acids^44^, indicating that its *in vivo* abundance is exceptionally low, similar to NeuGc.

Identifying carrier proteins for NeuGc and KDN in human tissues and cultured cells has been challenging due to their trace levels. However, recent findings have identified KDN-containing glycans in prostate-specific antigen (PSA)^45^. Wei *et al.* provided the first evidence of KDN-containing glycans in human glycoproteins by detecting them in clinical urine samples via mass spectrometry. Although no significant differences in KDN glycans were observed between patients with prostate cancer and those without, the analysis of trace glycans remains a valuable tool for biomarker discovery.

### Future perspectives

In this study, we demonstrated for the first time that Byonic, pGlyco3, and GRable enable comprehensive searches for proteins containing KDN. Examination of the MS/MS spectra of the identified peptides revealed that although KDN-related signals were significantly weaker than those for NeuAc and NeuGc, diagnostic ions were still detectable. These findings broaden the scope of glycoproteomics, offering promising prospects for elucidating novel biological phenomena and discovering new biomarkers in various diseases. In addition to the development of glycoproteomic software, new techniques for glycopeptide enrichment are being developed^46, 47^, and detailed analyses of attached glycans are underway.

Recent advancements in the Pan-Cancer project^48^ have enabled more in-depth investigations of **cancer-associated glycan alterations** across multiple cancer types^49^. These cross-cancer studies aim to identify common and unique glycan modifications that may serve as biomarkers or therapeutic targets. Along with individual advancements in software development, pretreatment techniques, and glycan databases, the synergistic progress of these technologies is expected to further enhance the depth of glycopeptide analysis.

In such analyses, generating **highly reliable glycopeptide datasets** is crucial. Our study significantly contributes to this field by providing a robust validation framework for glycopeptide identification, ensuring higher accuracy and confidence in glycoproteomic datasets.

As glycoproteomics continues to evolve, integrating **diverse analytical approaches** and applying rigorous validation criteria will be essential for deciphering glycan-related biological mechanisms. Our findings underscore the importance of selecting and combining software tools based on the analytical objectives and available resources. The ability to accurately identify rare glycans, such as KDN, will further enhance our understanding of glycan diversity and its functional roles in health and diseases.

## METHODS

### Cell culture

BJAB-K20 cells (originally established by Stephan Hinderlich and the late Werner Reutter at the Beuth University of Applied Sciences, Berlin, Germany) were kindly provided by Dr. Ajit Varki (University of California, San Diego) with permission from the original developers, and were cultured in RPMI 1640 medium supplemented with 10% fetal bovine serum (FBS) and 5 mM mannose.

### Glycopeptide preparation

Membrane proteins were prepared by ultracentrifugation, as previously reported. For protein enrichment, four times the volume of cold acetone was added to the cell suspension, followed by storing at -80 °C for 1 h. The precipitated proteins were collected by centrifugation at 15,000× g for 3 min, and the supernatant was discarded. The pellet was dissolved in 200 μL of phase transfer surfactant buffer and reduced at room temperature for 2 h with dithiothreitol at a concentration equal to the protein weight. Alkylation was performed by adding 2.5 times the volume of iodoacetamide relative to the protein weight, followed by incubation for 2 h in the dark at room temperature. Proteins were diluted 5-fold with ultrapure water and digested with lysyl endopeptidase at a protein-to-enzyme ratio of 200:1 for 1 h at 37 °C. Trypsin was then added at a protein-to-enzyme ratio of 100:1, and digestion was carried out overnight at 37 °C. Digestion was stopped by adding phenylmethylsulfonyl fluoride, and the surfactants were precipitated with 1% trifluoroacetic acid (TFA). After centrifugation at 1,000× g for 3 min at 4 °C, the supernatant was filtered using a 0.22 μm poly(1,1,2,2-tetrafluoroethylene) filter.

Glycopeptides were purified using an amide column (Amide-80, 2 x 50 mm; Tosoh, Japan). The sample was injected into the high-performance liquid chromatography system operating in isocratic mode with 75% acetonitrile containing 0.1% TFA, removing non-glycopeptides that did not bind to the column. Glycopeptides were then eluted with 50% acetonitrile containing 0.1% TFA at a flow rate of 0.2 mL/min, and detection was performed at 215 nm.

### IGOT-LC-MS

Aliquots of the collected glycopeptide fraction were lyophilized and subjected to PNGase F digestion in ^18^O-labeled water. The digests were analyzed using nanoLC-MS/MS on an Orbitrap Fusion mass spectrometer (Thermo Fisher Scientific, USA) coupled to a nano-flow LC system (Ultimate3000; Thermo Fisher Scientific, USA).

Samples were loaded onto a C18 tip column (0.15 mm id × 100 mm; 3 μm particles; Nikkyo Technos, Japan) and separated with a linear gradient of 5– 60% acetonitrile in 0.1% formic acid at a flow rate of 300 nl/min.

The raw data files were processed using Mascot Distiller software (ver.2.6; Matrix Science, USA) and converted into mgf format. The mgf files were then subjected to database searches using Mascot (ver.2.6.2; Matrix Science) with the UniProtKB database (downloaded on January 10, 2020; 7330 Homosapience entries). The following search conditions were used: enzyme, trypsin + LEP; maximum missed cleavages, 2; fixed modification, carbamidomethyl (C); variable modifications, ammonia loss (N-term C [carbamidomethyl]), Delta:H(1)N(- 1)18O(1) (N), Gln > pyroGlu (N-term Q), oxidation (M), MS1 tolerance of 7 ppm, and MS2 tolerance of 0.02 Da. Mascot exported the results file (. dat) as a CSV file and processed using Microsoft Excel.

### Parameter setting for each software

Byonic, pGlyco3, and MSFragger were operated in *N*-glycan mode. The search parameters for all software tools were standardized as follows: peptide weight range, 350–1000 Da; Peptide length range, 5–50 aa. precursorr mass tolerance: 7 ppm; MS2 fragment tolerance: 0.05 Da.

Fixed modification: Carbamidomethyl (C); variable modifications: ammonia loss (N-term C [carbamidomethyl]), Gln > pyroGlu (N-term Q), and oxidation (M). To search for glycopeptides containing NeuAc, we used the glycan database provided by Byonic, which consists of 132 human N-glycan structures (Byonic_human_132). To search for glycopeptides containing NeuGc and KDN, we constructed a library that included all possible combinations in which NeuAc in Byonic_human_132 was individually replaced with NeuGc and KDN, resulting in a total of 500 structures.

### Data analysis

The glycopeptides identified by each software tool were compiled and standardized into a common format. Glycopeptides with matching amino acid sequences and GC across different software tools were considered to have overlapping identifications. To ensure accuracy, several custom scripts were generated using ChatGPT to systematically verify the presence of Y0/Y1 ions, B/Y-ions, and various oxonium ion signals in all scans. All identified glycopeptides, including raw MS data, are available in the jPOST Repository.

## ACKNOWLEDGEMENTS

We gratefully acknowledge Masako Sukegawa and Mika Fujita for their kind support in the preparation of experimental samples. This work was supported by JSPS KAKENHI Grant Number JP24K02257 (to K.K.)

## COMPETING INTERESTS

The authors declare no conflicts of interest.

## AUTHOR CONTRIBUTIONS

H.S. and A.K. designed the study and wrote the manuscript; H.S., K.K., A.T., and M.M. conducted experiments; C.N. and H.K. contributed to the interpretation of the data. All the authors contributed to the manuscript.

## DATA AVAILABILITY

Mass spectrometry data and search results were deposited in the ProteomeXchange repository via jPOST with the dataset identifier PXD065680.

---

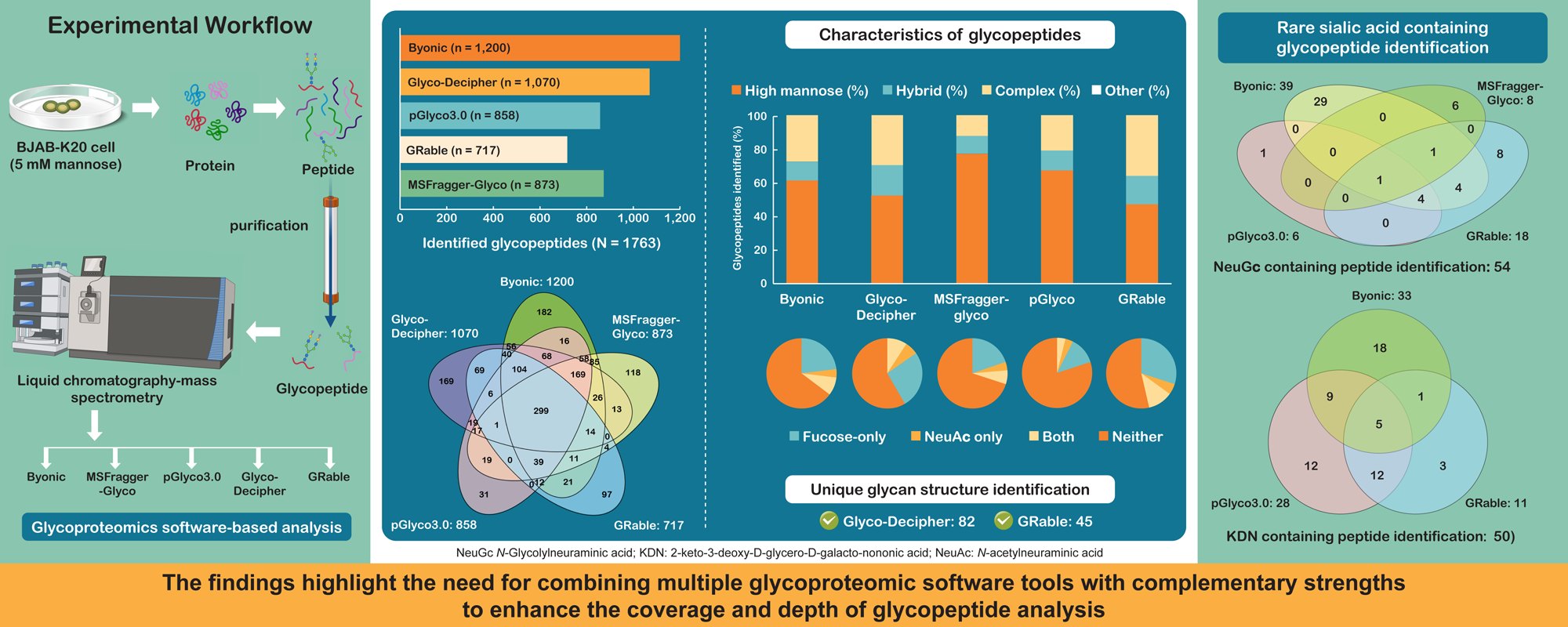

## Notes

### Competing Interest Statement

The authors have declared no competing interest.

## REFERENCES

1. Varki, A. Biological roles of glycans. Glycobiology 27, 3–49 (2017). 10.1093/glycob/cww086, PubMed: 27558841.

2. He, M., Zhou, X. & Wang, X. Glycosylation: Mechanisms, biological functions and clinical implications. Signal Transduct. Target. Ther. 9, 194 (2024). 10.1038/s41392-024-01886-1, PubMed: 39098853.

3. Reiding, K. R., Blank, D., Kuijper, D. M., Deelder, A. M. & Wuhrer, M. High-throughput profiling of protein N-glycosylation by MALDI-TOF-MS employing linkage-specific sialic acid esterification. Anal. Chem. 86, 5784–5793 (2014). 10.1021/ac500335t, PubMed: 24831253.

4. Tomiya, N. et al. Analyses of N-linked oligosaccharides using a two-dimensional mapping technique. Anal. Biochem. 171, 73–90 (1988). 10.1016/0003-2697(88)90126-1, PubMed: 3407923.

5. Nakagawa, H. et al. Identification of neutral and sialyl N-linked oligosaccharide structures from human serum glycoproteins using three kinds of high-performance liquid chromatography. Anal. Biochem. 226, 130–138 (1995). 10.1006/abio.1995.1200, PubMed: 7785764.

6. Toshima, T. et al. A novel serum marker, glycosylated Wisteria floribunda agglutinin-positive Mac-2 binding protein (WFA(+)-M2BP), for assessing liver fibrosis. J. Gastroenterol. 50, 76–84 (2015). 10.1007/s00535-014-0946-y, PubMed: 24603981.

7. Kaspar-Schoenefeld, S. et al. High-throughput proteome profiling with low variation in a multi-center study using dia-PASEF. bioRxiv, 2024.2005.2029.596405 (2024).

8. Stewart, H. I. et al. Parallelized acquisition of Orbitrap and astral analyzers enables high-throughput quantitative analysis. Anal. Chem. 95, 15656–15664 (2023). 10.1021/acs.analchem.3c02856, PubMed: 37815927.

9. Bern, M. & Kil, Y. J. (2012) **Chapter 13**. Becker C. Byonic: Advanced peptide and protein identification software. in Curr Protoc Bioinformatics, 13 11–13 20 14.

10. Fang, Z. et al. Glyco-Decipher enables glycan database-independent peptide matching and in-depth characterization of site-specific N-glycosylation. Nat. Commun. 13, 1900 (2022). 10.1038/s41467-022-29530-y, PubMed: 35393418.

11. Zeng, W. F., Cao, W. Q., Liu, M. Q., He, S. M. & Yang, P. Y. Precise, fast and comprehensive analysis of intact glycopeptides and modified glycans with pGlyco3. Nat. Methods 18, 1515–1523 (2021). 10.1038/s41592-021-01306-0, PubMed: 34824474.

12. Polasky, D. A., Yu, F., Teo, G. C. & Nesvizhskii, A. I. Fast and comprehensive N- and O-glycoproteomics analysis with MSFragger-Glyco. Nat. Methods 17, 1125–1132 (2020). 10.1038/s41592-020-0967-9, PubMed: 33020657.

13. Zeng, W. F. et al. pGlyco: A pipeline for the identification of intact N-glycopeptides by using HCD- and CID-MS/MS and MS3. Sci. Rep. 6, 25102 (2016). 10.1038/srep25102, PubMed: 27139140.

14. Liu, M. Q. et al. pGlyco 2.0 enables precision N-glycoproteomics with comprehensive quality control and one-step mass spectrometry for intact glycopeptide identification. Nat. Commun. 8, 438 (2017). 10.1038/s41467-017-00535-2, PubMed: 28874712.

15. Kong, S. et al. pGlycoQuant with a deep residual network for quantitative glycoproteomics at intact glycopeptide level. Nat. Commun. 13, 7539 (2022). 10.1038/s41467-022-35172-x, PubMed: 36477196.

16. Baker, P. R., Trinidad, J. C. & Chalkley, R. J. Modification site localization scoring integrated into a search engine. Mol. Cell. Proteomics 10, M111.008078 (2011). 10.1074/mcp.M111.008078, PubMed: 21490164.

17. Zeng, W. F., Yan, G., Zhao, H. H., Liu, C. & Cao, W. Uncovering missing glycans and unexpected fragments with pGlycoNovo for site-specific glycosylation analysis across species. Nat. Commun. 15, 8055 (2024). 10.1038/s41467-024-52099-7, PubMed: 39277585.

18. Devabhaktuni, A. et al. TagGraph reveals vast protein modification landscapes from large tandem mass spectrometry datasets. Nat. Biotechnol. 37, 469–479 (2019). 10.1038/s41587-019-0067-5, PubMed: 30936560.

19. An, Z. et al. N-linked glycopeptide identification based on open mass spectral library search. BioMed Res. Int. 2018, 1564136 (2018). 10.1155/2018/1564136, PubMed: 30186849.

20. Pioch, M., Hoffmann, M., Pralow, A., Reichl, U. & Rapp, E. glyXtoolMS: An open-source pipeline for semiautomated analysis of glycopeptide mass spectrometry data. Anal. Chem. 90, 11908–11916 (2018). 10.1021/acs.analchem.8b02087, PubMed: 30252445.

21. Nasir, W. et al. SweetNET: A bioinformatics workflow for glycopeptide MS/MS spectral analysis. J. Proteome Res. 15, 2826–2840 (2016). 10.1021/acs.jproteome.6b00417, PubMed: 27399812.

22. Park, G. W. et al. Integrated GlycoProteome analyzer (I-GPA) for automated identification and quantitation of site-specific N-glycosylation. Sci. Rep. 6, 21175 (2016). 10.1038/srep21175, PubMed: 26883985.

23. Toghi Eshghi, S., Shah, P., Yang, W., Li, X. & Zhang, H. GPQuest: A spectral library matching algorithm for site-specific assignment of tandem mass spectra to intact N-glycopeptides. Anal. Chem. 87, 5181–5188 (2015). 10.1021/acs.analchem.5b00024, PubMed: 25945896.

24. Lynn, K. S. et al. MAGIC: An automated N-linked glycoprotein identification tool using a Y1-ion pattern matching algorithm and in silico MS² approach. Anal. Chem. 87, 2466–2473 (2015). 10.1021/ac5044829, PubMed: 25629585.

25. Shen, J. et al. StrucGP: De novo structural sequencing of site-specific N-glycan on glycoproteins using a modularization strategy. Nat. Methods 18, 921–929 (2021). 10.1038/s41592-021-01209-0, PubMed: 34341581.

26. Kawahara, R. et al. Community evaluation of glycoproteomics informatics solutions reveals high-performance search strategies for serum glycopeptide analysis. Nat. Methods 18, 1304–1316 (2021). 10.1038/s41592-021-01309-x, PubMed: 34725484.

27. Hogan, R. A., Pepi, L. E., Riley, N. M. & Chalkley, R. J. Comparative Analysis of Glycoproteomic Software using a Tailored Glycan Database. bioRxiv, 2024.2007.2024.604997, 2024.07.24.604997 (2024). 10.1101/2024.07.24.604997, PubMed: 39091841.

28. Rangel-Angarita, V., Mahoney, K. E., Ince, D. & Malaker, S. A. A systematic comparison of current bioinformatic tools for glycoproteomics data. bioRxiv, 2022.2003.2015.484528 (2022).

29. Chalkley, R. J. & Baker, P. R. Improving the depth and reliability of glycopeptide identification using protein prospector. Mol. Cell. Proteomics 24, 100903 (2025). 10.1016/j.mcpro.2025.100903, PubMed: 39788319.

30. Lee, L. Y. et al. Toward automated N-glycopeptide identification in glycoproteomics. J. Proteome Res. 15, 3904–3915 (2016). 10.1021/acs.jproteome.6b00438, PubMed: 27519006.

31. Darula, Z. & Medzihradszky, K. F. Carbamidomethylation side reactions may lead to glycan misassignments in glycopeptide analysis. Anal. Chem. 87, 6297–6302 (2015). 10.1021/acs.analchem.5b01121, PubMed: 25978763.

32. Nagai-Okatani, C. et al. *GRable*. version 1.0: A Software Tool for Site-Specific Glycoform Analysis with Improved MS1-Based Glycopeptide Detection with Parallel Clustering and Confidence Evaluation with MS2 Information. Mol. Cell. Proteomics, 23. 10.0833(2024).

33. Kaji, H. et al. Lectin affinity capture, isotope-coded tagging and mass spectrometry to identify N-linked glycoproteins. Nat. Biotechnol. 21, 667–672 (2003). 10.1038/nbt829, PubMed: 12754521.

34. Nilsson, J. Liquid chromatography-tandem mass spectrometry-based fragmentation analysis of glycopeptides. Glycoconj. J. 33, 261–272 (2016). 10.1007/s10719-016-9649-3, PubMed: 26780731.

35. Halim, A. et al. Assignment of saccharide identities through analysis of oxonium ion fragmentation profiles in LC-MS/MS of glycopeptides. J. Proteome Res. 13, 6024–6032 (2014). 10.1021/pr500898r, PubMed: 25358049.

36. Campbell, M. P. A review of software applications and databases for the interpretation of glycopeptide data. Trends Glycoscience and Glycotechnology, 29, E51–E62 (2017). 10.4052/tigg.1601.1E.

37. Yu, J. et al. Distinctive MS/MS fragmentation pathway of glycopeptide-generated oxonium ions provide evidence of the glycan structure. Chemistry 22, 1114–1124 (2016). 10.1002/chem.201503659, PubMed: 26663535.

38. Hiono, T. et al. Combinatorial Approach with mass spectrometry and Lectin microarray Dissected Site-Specific Glycostem and Glycoleaf Features of the Virion-Derived Spike Protein of Ancestral and gamma Variant SARS-CoV-2 Strains. J. Proteome Res. 23, 1408–1419 (2024). 10.1021/acs.jproteome.3c00874, PubMed: 38536229.

39. Bubis, J. A. et al. Challenging the Astral mass analyzer to quantify up to 5,300 proteins per single cell at unseen accuracy to uncover cellular heterogeneity. Nat. Methods 22, 510–519 (2025). 10.1038/s41592-024-02559-1, PubMed: 39820751.

40. Hinderlich, S., Berger, M., Keppler, O. T., Pawlita, M. & Reutter, W. Biosynthesis of N-acetylneuraminic acid in cells lacking UDP-N-acetylglucosamine 2-epimerase/N-acetylmannosamine kinase. Biol. Chem. 382, 291–297 (2001). 10.1515/BC.2001.036, PubMed: 11308027.

41. Kawanishi, K. et al. Human species-specific loss of CMP-N-acetylneuraminic acid hydroxylase enhances atherosclerosis via intrinsic and extrinsic mechanisms. Proc. Natl Acad. Sci. U. S. A. 116, 16036–16045 (2019). 10.1073/pnas.1902902116, PubMed: 31332008.

42. Seo, N. et al. In-depth characterization of non-human sialic acid (Neu5Gc) in human serum using label-free ZIC-HILIC/MRM-MS. Anal. Bioanal. Chem. 413, 5227–5237 (2021). 10.1007/s00216-021-03495-1, PubMed: 34235565.

43. Kawanishi, K. et al. Evolutionary conservation of human ketodeoxynonulosonic acid production is independent of sialoglycan biosynthesis. J. Clin. Invest. 131, e137681 (2021). 10.1172/JCI137681, PubMed: 33373330.

44. Jahan, M., Thomson, P. C., Wynn, P. C. & Wang, B. Red meat derived glycan, N-acetylneuraminic acid (Neu5Ac) is a major sialic acid in different skeletal muscles and organs of nine animal species-A guideline for human consumers. Foods 12, 337 (2023). 10.3390/foods12020337, PubMed: 36673429.

45. Wang, W. et al. Human prostate-specific antigen carries N-glycans with Ketodeoxynononic acid. Engineering 26, 119–131 (2023). 10.1016/j.eng.2023.02.009.

46. Potel, C. M. et al. Uncovering protein glycosylation dynamics and heterogeneity using deep quantitative glycoprofiling (DQGlyco). Nat. Struct. Mol. Biol. (2025). 10.1038/s41594-025-01485-w, PubMed: 39930009.

47. Fang, P. et al. Ultradeep N-glycoproteome atlas of mouse reveals spatiotemporal signatures of brain aging and neurodegenerative diseases. bioRxiv, 2025.2002.2015.638397 (2025).

48. Ma, X. et al. Pan-cancer genome and transcriptome analyses of 1,699 paediatric leukaemias and solid tumours. Nature 555, 371–376 (2018). 10.1038/nature25795, PubMed: 29489755.

49. Zhang, H. & Hu, Y. GPnotebook: A pan-cancer glycoproteomic database and toolkit for analysis of protein glycosylation changes associated with cancer phenotypes. bioRxiv, 2024.2004.2018.589619 (2024).

