## supplemental information for "Comparative Evaluation of Glycoproteomics Software for Rare Glycopeptide Identification"

1. **The glycoproteomics software used in this study**

**Byonic**

The Byonic software was developed and commercialized by Protei Metrix Inc. The detailed algorithms behind Byonic are unpublished; however, the peptide identification algorithm from its predecessor, ByOnic, can provide insights regarding the behavior of Byonic in glycopeptide identification. ByOnic utilizes Lookup Peaks, a hybrid method that combines conventional database searches with de novo sequencing, thereby enabling efficient and sensitive peptide identification.

Look-up Peaks identify continuous b- and y-ion pairs of peptides by determining whether the mass difference between the two peaks in the spectrum corresponds to the mass of an amino acid residue. For example, if the two peaks at 500 and 615 Da differ by 115 Da, they correspond to aspartic acid. Thus, these peaks were considered part of a peptide. The partial amino acid sequence of the peptide is thus identified, and the next database search is performed. Candidate peptides were identified by matching the calculated peptide masses from protein databases with those of the partial amino acid sequences estimated from the Lookup Peaks. In silico spectra for these selected candidate peptides were generated and compared with the measured spectra, and the match was assigned a score based on mass error and peak intensity. For the modified peptides, the allowed mass error for the precursor ions was set wider than usual. The possibility of modification was predicted based on the difference between the measured and theoretical masses of the precursor ions. An in silico spectrum of the modified peptide was generated and compared to the measured spectrum. Although detailed algorithms for glycopeptide analysis remain unpublished, developers have disclosed that their method relies on comparing the intensities of the predicted and measured spectra, considering both bare peptides and peptide+HexNAc. Consequently, the glycopeptide identification algorithm in Byonic can be summarized as follows. First, the peptide is identified. Then, the glycan mass is calculated by subtracting the mass of the bare peptide from that of the precursor ion. The predicted glycan mass was selected to match the glycan masses listed in the glycan database. Finally, an in silico spectrum was generated and matched with the measured spectrum.

**MSFragger-Glyco**

MSFragger-Glyco identified the glycopeptides by initially checking for the presence of oxonium ions in the acquired MS/MS spectra. Peptides lacking oxonium ions are considered non-glycopeptides. During the database searches, proteins listed in the protein database are cleaved in silico, and peptides with N-X-S/T consensus sequences are extracted. Peptide sequences without consensus are used for non-glycopeptide identification.

In contrast, signals confirmed to contain oxonium ions are classified as glycopeptides, and mass-offset searches were performed. This approach is based on the hypothesis that the difference between the observed mass of the precursor ion and the theoretical mass of the peptide (without glycans) corresponds to the mass of the glycan. The calculated glycan mass is then compared with a glycan list, and the attached glycan is identified.

**pGlyco3**

Similar to Byonic and MSFragger-Glyco, pGlyco3 requires a glycan database for analysis. However, unlike the other two software, glycan analysis is performed before peptide identification. In pGlyco3, the spectrum containing m/z 204, indicating the presence of HexNAc, is assigned to a glycopeptide. Subsequently, the Y-ion series, consisting of peptide and glycan fragment combinations, was compared.

The Y-complementary ion was defined as the difference between the precursor mass and the observed Y-ion mass. This ion consists solely of glycans and lacks peptides, enabling the identification of glycans prior to peptides.

**Glyco-Decipher**

GlycoDecipher is a glycan list-independent glycoproteomic software that is characterized by in silico deglycosylation. During in silico deglycosylation, oxonium, B, and Y ions derived from glycans are removed in silico, leaving only peptide-derived b- and y-ions for peptide identification. Glycopeptides with the same peptide backbone exhibited similar fragmentation pattern spectra. Spectrum expansion was applied to peptides that were poorly identified in the first analysis, thereby increasing the possibility of peptide identification by comparing peptide fragmentation patterns. After identifying the peptide backbone, the glycan mass was calculated from the difference in the masses of the precursor and peptide backbone. If a glycan is listed in the database, it is identified based on a match between the theoretical and experimental B/Y ions. Even if glycans such as modified glycans are not present in the database, they can still be identified by calculating the mass difference between Y ions using monosaccharide stepping.

**GlRable**

GRable is a new glycoproteomics software tool based on the Glyco-RIDGE method that can predict glycan structures from glycopeptides without MS/MS spectral data. GRable requires two sets of data: deglycoproteomics and glycoproteomics. Deglycoproteomic data are obtained after N-glycanase treatment of glycopeptides in 18O water, which specifically labels the glycosylation sites. While glycopeptides exhibit heterogeneity, removal of glycans significantly reduces the structural complexity and concentrates the data into a single core peptide. This enables comprehensive identification similar to that achieved using standard proteomics. Subsequently, information regarding the core peptide sequence, glycosylation site, mass, and retention time (RT) is obtained using IGOT-LC-MS. Next, the glycan structures are predicted from the glycoproteomic data. Glycopeptides with identical core peptides exhibited similar elution times during LC. However, when a neutral monosaccharide (Hex, HexNAc, or fucose) is attached to a peptide, the RT shifts up to 1 min earlier, whereas the attachment of an acidic monosaccharide (N-acetylneuraminic acid [NeuAc] or N-glycolylneuraminic acid [NeuGc]) causes a delay of up to 3 min. Using the mass and RT of the core peptide identified by IGOT-LC-MS, glycan structures were predicted by identifying signals that showed an increased mass corresponding to the monosaccharide and a shift in RT depending on the type of attached glycan. One limitation of GRable is that glycan analysis can only be performed if multiple glycans exist in a single-core peptide. However, it can predict glycan structures even from weak signals, even when MS/MS data is unavailable. For signals in the acquired MS/MS spectra, the reliability of the assigned glycopeptides was ensured by verifying diagnostic ions (oxonium and Y ions).

2. **Supplemental Figures**

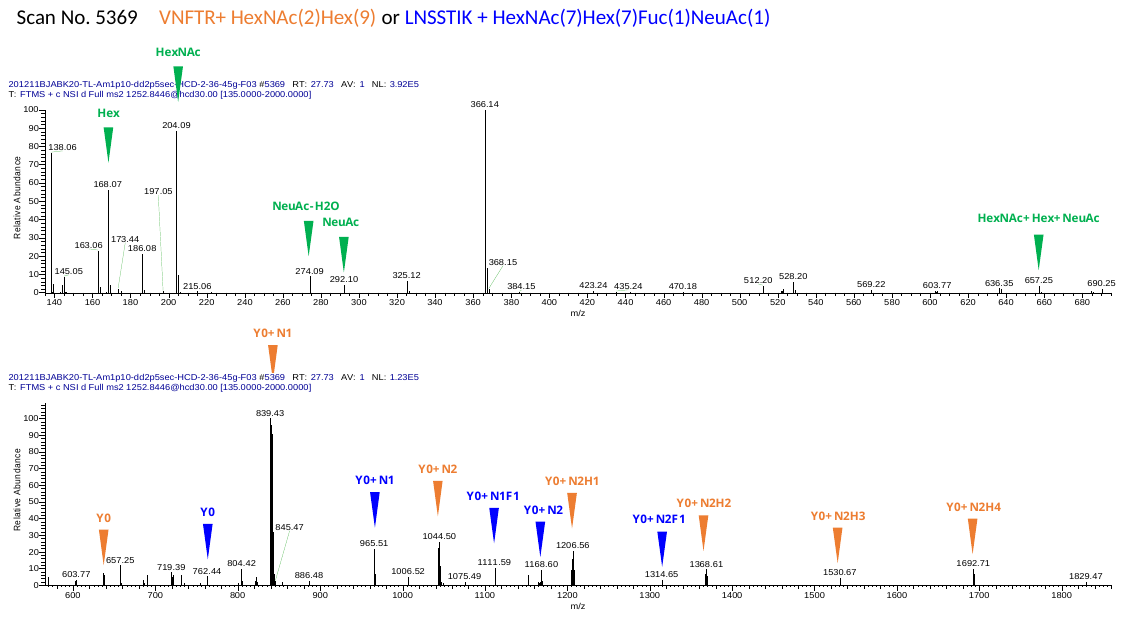

Figure S1 The MS/MS spectra of scan 5369 containing different peptides identified by Glyco-Decipher and GRable.

From this spectrum, Glyco-Decipher identifies VNFTR + HexNAc(2)Hex(9) and GRable identifies LNSSTIK + HexNAc(7)Hex(7)Fuc(1)NeuAc(1). Both b-, y-, and Y-series ions from YNFTR and LNSSTIK are detected in this spectrum. In addition, a Y1F signal (Y0+N1F1) and a HexNAc+Hex+NeuAc signal were detected, suggesting the presence of the HexNAc(7)Hex(7)Fuc(1)NeuAc(1) structure suggested by GRable.

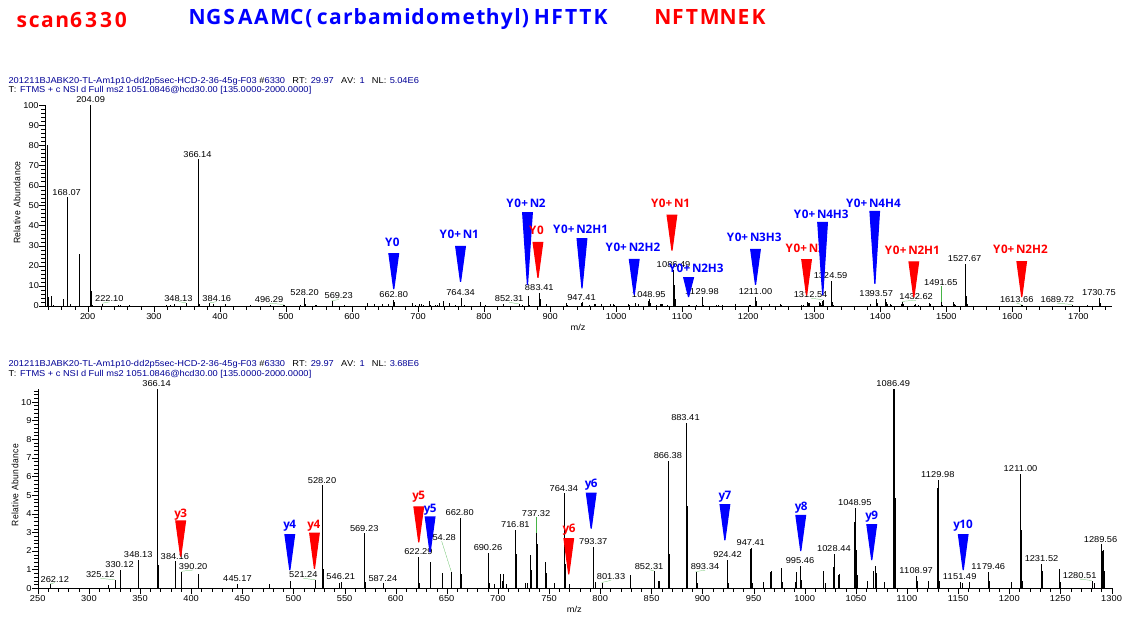

Figure S2 The MS/MS spectra of scan 6330 containing different peptides identified by pGlyco3 and GRable.

In scan 6330, NFTMNEK2:5:0:0 (2099.8207) was identified from pGlyco3 and NGSAAMCHFTTK5:5:0:0 (3093.2168) from GRable. The m/z of the precursor ion in this spectrum is 1050.7, which is considered to be a mixture of two components: a divalent ion of NFTMNEK2:5:0:0 ([M+2H+]2+＝1050.414) and a trivalent ion of NGSAAMCHFTTK5:5:0:0 ([M+3H+]3+1050.7509).

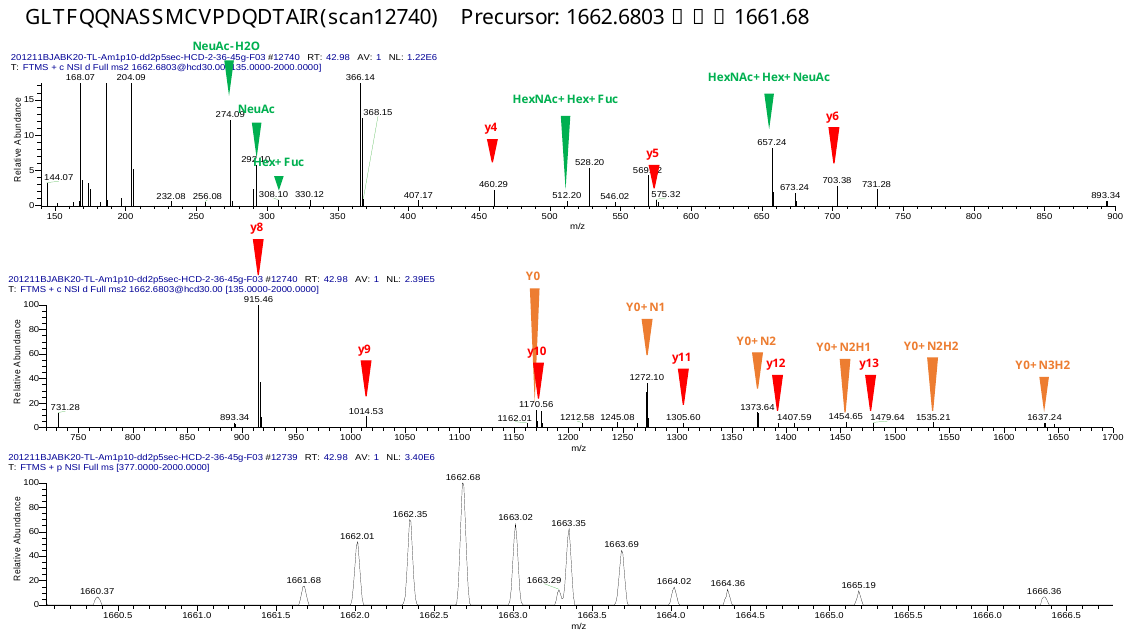

Figure S3 The MS/MS spectra of scan 12740 containing different peptides identified by Byonic, Glyco-Decipher and MSFragger-Glyco.

In scan12740, common core peptides GLTFQQNASSMCVPDQDTAIR were identified from Byonic, Glyco-Decipher, and MSFragger, but the glycan structures were different: HexNAc(6)Hex(7)NeuAc(1) in Byonic, HexNAc(6)Hex(7)Fuc(2) in Glyco-Decipher, HexNAc(6)Hex(5)NeuAc(1) + oxidized methionine in MSFragger. The masses of two Fuc (292.1158) and one NeuAc (291.0954) are close, so this is probably one of the most misidentified combinations.

The glycopeptide identified by MSFragger-Glyco was incorrect because there was no evidence for the presence of oxidized methionine. On the other hand, Hex+HexNAc+Fuc (m/z 512.2) and Hex+Fuc (m/z 308.1) indicating the presence of fucose at the non-reducing end, NeuAc-H2O (m/z 274) , NeuAc (m/z 292) and Hex+HexNAc+NeuAc (m/z 657.2) signals indicating the presence of NeuAc were observed.

The m/z for the precursor ion in this scan was 1662.6803, while the m/z for the trivalent ion of GLTFQQNASSMCVPDQDTAIR+HexNAc(6)Hex(7)NeuAc(1) is 1661.6771 and that of GLTFQQNASSMCVPDQDTAIR+HexNAc(6)Hex(7)Fuc(2) is 1663.0313, which is intermediate between the two.

Confirming the MS1 signal, the monoisotopic mass is 1661.68, but the peak shapes at m/z 1663.02 and m/z 1663.35 are a little different from the normal distribution. This is probably due to the overlap of the GLTFQQNASSMCVPDQDTAIR+HexNAc(6)Hex(7)Fuc(2) signals. This suggests that the 12740 scan is a chimeric spectrum of two glycopeptides and that different peptides were identified by Byonic and Glyco-Decipher, respectively.
