## Supplementary material for "Comparative Evaluation of Glycoproteomics Software for Rare Glycopeptide Identification": Table S1

| Software | No. | Scan | Proteins | Genes | Pep. seq. | pep_var_mod_pos | Glycosylation | No. of HexNAc | No. of Hex | No. of Fuc | No. of NeuAc | Composition (Hex:HexNAc:Fuc:NeuAc) | glycan type | Listed in the Glycan database (Byonic 132) (0: Not listed, 1: listed) | Identified by Byonic (0: Not identified, 1: identified) | Identified by Glyco-Decipher (0: Not identified, 1: identified) | Identified by MSFragger (0: Not identified, 1: identified) | Identified by pGlyco3 (0: Not identified, 1: identified) | Identified by GRable (0: Not identified, 1: identified)GRable | Intensity of HexNAc signal (138.0555) | Intensity of HexNAc signal (144.0661) | Intensity of HexNAc signal (168.0660) | Intensity of HexNAc signal (186.0766) | Intensity of HexNAc signal (204.0872) | Intensity of NeuAc signal (292.1032) | Intensity of NeuAc-H2O signal (274.0927) | Intensity of Hex+HexNAc signal (366.14) | Intensity of Hex+HexNAc+Fuc signal (512.1979) | Intensity of Hex2+HexNAc signal (617.2166) | Intensity of Hex+HexNAc+NeuAc signal (657.2354) | Intensity of Hex+NeuAc signal (454.156) | Intensity of Hex+NeuGc signal (470.1509) | Intensity of Fuc signal (147.0657) | Intensity of Hex+HexNAc+Fuc signal (511.1901) | Intensity of Hex+Fuc signal (308.1107) | Intensity of HesNAc+Fuc signal (349.1373) | Intensity of Y0 signal | Intensity of signal Y1 | Intensity of signal Y1F | Intensity of signal Y0(z=2) | Intensity of Y1(z=2) signal | Intensity of Y1F(z=2) signal | Intensity of Y0(z=3) signal | Intensity of Y1(z=3) signal | Intensity of Y1F(z=3) signal | Y0 or Y1 ions detected or not ? (0: Not detected, 1: Detected) | Detected y ions | Detected b ions |
|---|---|---|---|---|---|---|---|---|---|---|---|---|---|---|---|---|---|---|---|---|---|---|---|---|---|---|---|---|---|---|---|---|---|---|---|---|---|---|---|---|---|---|---|---|---|---|---|---|
| Byonic | 1 | 20497 | Q9GZR5 | ELOVL4 | GLLDSEPGSVLNVVSTALNDTVEFYR | N19(HexNAc(2)Hex(9)) | 20 | 2 | 9 | 0 | 0 | 2:9:0:0 | pauci- or high mannose | 1 | 1 | 0 | 1 | 0 | 0 | 39798.4 | 3387.9 | 988.068 | 12651.1 | 65697.3 | 1112.9 | 2828.64 | 32238.8 | × | × | × | × | × | × | × | × | × | × | × | × | 6395.47 | 22587.6 | × | × | × | × | 1 | × | × |
| Byonic | 5 | 17196 | Q96G23 | CERS2 | LWLPVNLTWADLEDRDGR | N6(HexNAc(2)Hex(9)) | 19 | 2 | 9 | 0 | 0 | 2:9:0:0 | pauci- or high mannose | 1 | 1 | 0 | 1 | 1 | 0 | 283886 | 25921.7 | 6726.62 | 118377 | 598888 | × | 11490.4 | 259166 | × | × | × | × | × | × | × | × | × | × | × | × | 99099.1 | 1240390 | × | × | 30257.6 | × | 1 | × | × |
| Byonic | 6 | 18185 | Q96G23 | CERS2 | LWLPVNLTWADLEDR | N6(HexNAc(2)Hex(9)) | 19 | 2 | 9 | 0 | 0 | 2:9:0:0 | pauci- or high mannose | 1 | 1 | 1 | 1 | 1 | 0 | 137292 | 1202.28 | 81401.3 | 1827.8 | 266370 | 10874.6 | 32079.9 | 94884.7 | × | × | × | × | × | × | × | × | × | 7880.98 | × | × | 8228.23 | 39731 | × | × | × | × | 1 | × | × |
| Byonic | 7 | 17688 | Q8NSB7 | CERS5 | FWLPENVSWADLEGPADGYGYPR | N6(HexNAc(2)Hex(9)) | 26 | 2 | 9 | 0 | 0 | 2:9:0:0 | pauci- or high mannose | 1 | 1 | 0 | 0 | 0 | 0 | 137481 | 10999.4 | 2544.14 | 43195.3 | 163014 | 5038.49 | 7544.75 | 101425 | × | × | 2878.47 | × | × | × | × | × | × | × | × | × | 2118.55 | 10904 | × | 1476.7 | × | × | 1 | × | × |
| Byonic | 10 | 8444 | Q8IWB1 | ITPRIP | ENATVPENEEEIIRK | N2(HexNAc(2)Hex(9)) | 27 | 2 | 9 | 0 | 0 | 2:9:0:0 | pauci- or high mannose | 1 | 1 | 0 | 0 | 1 | 1 | 213425 | 12235.9 | 143939 | 76793.9 | 363986 | 7451.88 | 13141.9 | 174733 | × | × | 2834.13 | × | × | × | × | × | × | 11238.7 | 35051.1 | × | 22279.7 | 286888 | × | × | 3452.82 | × | 1 | y2, y4, y6, y10, y11, y12, y15 | b2, b3, b5, b9 |
| Byonic | 19 | 8432 | Q92542 | NCSTN | KIYIPLNK | N7(HexNAc(2)Hex(9)) | 45 | 2 | 9 | 0 | 0 | 2:9:0:0 | pauci- or high mannose | 1 | 1 | 1 | 0 | 0 | 0 | 197446 | 17070.5 | 133057 | 78908.8 | 335000 | 7107.97 | 18381.4 | 121448 | × | × | × | × | × | × | × | × | × | 54902.8 | 257019 | × | 4116.59 | 125850 | × | × | × | × | 1 | y4, y6, y8 | b2, b4 |
| Byonic | 20 | 7018 | P79483 | HLA-DRB3 | SECHFFNGTER | C3(Carbamidomethyl); N7(HexNAc(2)Hex(9)) | 48 | 2 | 9 | 0 | 0 | 2:9:0:0 | pauci- or high mannose | 1 | 1 | 1 | 0 | 1 | 0 | 241527 | 14611.8 | 139559 | 66593.4 | 250721 | 1230.33 | 5262.89 | 160666 | × | × | × | × | × | 1229.43 | × | × | × | 9976.71 | 76226.8 | × | 11511 | 94156.4 | × | × | × | × | 1 | y1, y4, y11 | b2, b3, b4, b5, b6 |
| Byonic | 23 | 11183 | Q8TEM1 | NUP210 | VNFTLEASEGCYR | N2(HexNAc(2)Hex(9)); C11(Carbamidomethyl) | 44 | 2 | 9 | 0 | 0 | 2:9:0:0 | pauci- or high mannose | 1 | 1 | 1 | 1 | 1 | 0 | 395378 | 37398.3 | 282095 | 9797.86 | 1096410 | × | × | 315792 | × | × | × | × | × | × | × | × | × | 143477 | 358585 | × | 13930.8 | 226084 | × | × | × | × | 1 | y1, y2, y4, y5, y6, y7, y8, y9, y10, y11, y13 | b2, b3, b6 |
| Byonic | 25 | 4145 | P01871 | IGHM | YKNNSDISSTR | N3(HexNAc(2)Hex(9)) | 46 | 2 | 9 | 0 | 0 | 2:9:0:0 | pauci- or high mannose | 1 | 1 | 1 | 1 | 1 | 0 | 207607 | 15966.4 | 143505 | 76176.1 | 425042 | × | × | 130364 | × | × | × | × | × | 17480 | × | × | × | 13528.2 | 49811.7 | × | 13503.8 | 55044.1 | × | × | × | × | 1 | y1, y2, y3, y4, y5, y6, y7, y8, y9, y10, y11 | b1, b1, b2, b9, b11 |
| Byonic | 58 | 3894 | P01871 | IGHM | NNSDISSTR | N1(HexNAc(2)Hex(9)) | 46 | 2 | 9 | 0 | 0 | 2:9:0:0 | pauci- or high mannose | 1 | 1 | 1 | 0 | 0 | 1 | 74024.7 | 4254.75 | 47337.4 | 22882.1 | 140175 | 1284.29 | 4116.68 | 79343.8 | × | × | 1464.99 | × | × | × | × | × | × | 25614.1 | 146988 | × | 1224.07 | 1244.78 | × | × | × | × | 1 | y1, y4, y5, y7, y8, y9 | × |
| Byonic | 76 | 10704 | P05107 | ITGB2 | LNFTGPGDPDSIR | N2(HexNAc(2)Hex(9)) | 50 | 2 | 9 | 0 | 0 | 2:9:0:0 | pauci- or high mannose | 1 | 1 | 1 | 1 | 1 | 1 | 5090.93 | 37929.8 | 355491 | 224417 | 1523850 | × | × | 478685 | × | × | × | × | × | × | × | × | × | 200071 | 858938 | × | 105734 | 660666 | × | × | × | × | 1 | y1, y2, y3, y3, y4, y5, y6, y7, y8, y9, y10, y11, y13 | b2, b2, b3, b3, b4, b5, b7, b8 |
| Byonic | 83 | 14834 | Q9BY67 | CADM1 | DVTVIEGEVATISCQVNK | C14(Carbamidomethyl); N17(HexNAc(2)Hex(9)) | 67 | 2 | 9 | 0 | 0 | 2:9:0:0 | pauci- or high mannose | 1 | 1 | 0 | 0 | 0 | 0 | 286782 | 25165.8 | 193038 | 96705.4 | 560616 | × | 8164.86 | 257109 | × | × | × | × | × | × | × | × | × | 20633.5 | × | × | 22963.3 | 12800.7 | × | × | × | × | 1 | × | × |
| Byonic | 94 | 5034 | Q92478 | CLEC2B | EEGDWNSSK | N6(HexNAc(2)Hex(9)) | 57 | 2 | 9 | 0 | 0 | 2:9:0:0 | pauci- or high mannose | 1 | 1 | 1 | 0 | 1 | 0 | 116145 | 10114 | 74855.4 | 46360.8 | 242153 | × | 1102.36 | 117533 | × | 998.485 | × | × | × | × | × | × | × | 42759.1 | 257990 | × | × | 9305.52 | × | × | × | × | 1 | y2, y3, y4, y5, y7, y9 | b2, b5, b6, b7, b8, b9 |
| Byonic | 96 | 6973 | P20645 | M6PR | LKPLFNK | N6(HexNAc(2)Hex(9)) | 57 | 2 | 9 | 0 | 0 | 2:9:0:0 | pauci- or high mannose | 1 | 1 | 1 | 0 | 1 | 1 | 252978 | 17311.6 | 205427 | 123936 | 739280 | × | 1941.47 | 231222 | × | × | × | × | × | 2914.43 | × | × | × | 186911 | 1107470 | × | × | 96297.4 | × | × | × | × | 1 | y1, y2, y3, y5, y7 | b2, b5, b6, b7 |
| Byonic | 101 | 9623 | P53801 | PTTG1IP | NVSCLWCNTNK | N1(HexNAc(2)Hex(9)); C4(Carbamidomethyl); C7(Carbamidomethyl) | 54 | 2 | 9 | 0 | 0 | 2:9:0:0 | pauci- or high mannose | 1 | 1 | 0 | 0 | 0 | 0 | 117098 | 7577.93 | 66336.7 | 37552.6 | 184734 | × | 3295.49 | 79495.5 | × | × | × | × | × | × | × | × | × | 22906 | 43384.8 | × | × | 6540.88 | × | × | × | × | 1 | y4, y5, y6, y9, y11 | b2 |
| [illegible] | [illegible] | [illegible] | [illegible] | [illegible] | [illegible] | [illegible] | [illegible] | [illegible] | [illegible] | [illegible] | [illegible] | [illegible] | [illegible] | [illegible] | [illegible] | [illegible] | [illegible] | [illegible] | [illegible] | [illegible] | [illegible] | [illegible] | [illegible] | [illegible] | [illegible] | [illegible] | [illegible] | [illegible] | [illegible] | [illegible] | [illegible] | [illegible] | [illegible] | [illegible] | [illegible] | [illegible] | [illegible] | [illegible] | [illegible] | [illegible] | [illegible] | [illegible] | [illegible] | [illegible] | [illegible] | [illegible] | [illegible] | [illegible] |

| | | | | | | | | | | | |
|---|---|---|---|---|---|---|---|---|---|---|---|
| pGlyco207 | 1551 | 6415 | O75951 | LYZL6 | FNISK | | 58 | 2 | 9 | 0 | [illegible] |
| pGlyco490 | 1562 | 9566 | P04233 | CD74 | GHHNCSESLELEDPSSGLGVTK | 5,Carbamidomethyl[C]; | 270 | 2 | 9 | 0 | [illegible] |
| pGlyco615 | 1569 | 11152 | Q9BRN9 | TM2D3 | LPADCIDCTTNFSCTYGK | 5,Carbamidomethyl[C];8,Carbamidomethyl[C];14,Carbamidomethyl[C]; | 87 | 2 | 9 | 0 | [illegible] |
| pGlyco620 | 1570 | 11278 | Q96BZ4 | PLD4 | NISLAVATSSPTLAR | | 171 | 2 | 9 | 0 | [illegible] |
| pGlyco681 | 1575 | 12137 | P11717 | IGF2R | TNITLVCKPGDLESAPVLR | 7,Carbamidomethyl[C]; | 581 | 2 | 9 | 0 | [illegible] |
| pGlyco764 | 1580 | 13815 | Q9NV96 | TMEM30A | YSLNVTYNYPVHYFDGR | | 294 | 2 | 9 | 0 | [illegible] |
| pGlyco837 | 1586 | 16565 | Q92854 | SEMA4D | EVHLVQFHEPDIYNYSALLLSEDKDTLYIGAR | | 49 | 2 | 9 | 0 | [illegible] |
| pGlyco841 | 1587 | 16709 | Q7Z2K6 | ERMP1 | DSGIWINGFDYTGISHITPHIPEINDSIR | | 730 | 2 | 9 | 0 | [illegible] |
| pGlyco844 | 1588 | 17056 | Q9UP95 | SLC12A4 | SIFDPPVFPVCMLGNR | 11,Carbamidomethyl[C]; | 312 | 2 | 9 | 0 | [illegible] |
| pGlyco855 | 1589 | 17629 | Q9BXR5 | TLR10 | LETLSLVSCFANNTPLEHLDLSQNLLQHK | 9,Carbamidomethyl[C]; | 395 | 2 | 9 | 0 | [illegible] |
| pGlyco860 | 1590 | 18412 | Q9NR96 | TLR9 | QALEVAPGALLGLGNLTHLSLK | 1,Gln->pyro-Glu[AnyN-termQ]; | 200 | 2 | 9 | 0 | [illegible] |
| pGlyco861 | 1591 | 18416 | O75629 | CREG1 | LNITNIWVLDYFGGPK | | 193 | 2 | 9 | 0 | [illegible] |
| MSFragger | 1604 | 6346 | O00584 | RNASET2 | QDQQLQNCTEPGEQPSPK | 8C(57.0214), 7N(1864.6342) | 212 | 2 | 9 | 0 | [illegible] |
| MSFragger | 1614 | 7649 | Q16533 | SNAPC1 | EDKPLSLSMPVITEEEENESLSGTEFTASKK | 9M(15.9949), 18N(1864.6342) | 341 | 2 | 9 | 0 | [illegible] |
| MSFragger | 1633 | 10469 | Q99729 | HNRNPAB | GRGWTGAAAGAGGATAAPPSGNQNGAEGDQINASKNEE | 32N(1864.6342) | 30 | 2 | 9 | 0 | [illegible] |
| MSFragger | 1666 | 13348 | Q9Y487 | ATP6V0A2 | SVNLFGSGWNVSAMYSSSHPPAEHKK | 10N(1864.6342) | 474 | 2 | 9 | 0 | [illegible] |
| MSFragger | 1668 | 13462 | Q92542-2 | NCSTN | DLYEYSWVQGPLHSNETDR | 15N(1864.6342) | 582 | 2 | 9 | 0 | [illegible] |
| MSFragger | 1671 | 13672 | P08575-10 | PTPRC | SAPPSQVWNMTVSMTSDNSMHVK | 9N(1864.6342) | 377 | 2 | 9 | 0 | [illegible] |
| MSFragger | 1676 | 14022 | P20273-2 | CD22 | SATLTCESDANPPVSHYTWFDWNNQSLPYHSQK | 6C(57.0214), 24N(1864.6342) | 437 | 2 | 9 | 0 | [illegible] |
| MSFragger | 1683 | 14378 | P68104-2 | EEF1A1 | VETGVLKPGMVVTFAPVNVTTEVK | 18N(1864.6342) | 253 | 2 | 9 | 0 | [illegible] |
| MSFragger | 1701 | 15749 | Q969N2-6 | PIGT | KTYAIYDLLDTAMINNSR | 15N(1864.6342) | 317 | 2 | 9 | 0 | [illegible] |
| MSFragger | 1704 | 15869 | P18850 | ATF6 | NQTNTFFGSPPAATEATHVVSTIPESLQ | 1N(1864.6342) | 643 | 2 | 9 | 0 | [illegible] |
| MSFragger | 1716 | 16343 | P32942 | ICAM3 | VELAPLPPWQPVGQNFTLR | 15N(1864.6342) | 124 | 2 | 9 | 0 | [illegible] |
| MSFragger | 1719 | 16487 | O96005-3 | CLPTM1 | RAEDYGPVEVISHWHPNITINIVDDHTPWVK | 17N(1864.6342) | 129 | 2 | 9 | 0 | [illegible] |
| MSFragger | 1721 | 16588 | Q10589-2 | BST2 | GFQDVEAQAATCNHTVMALMASLDAEK | 12C(57.0214), 13N(1864.6342) | 70 | 2 | 9 | 0 | [illegible] |
| MSFragger | 1729 | 16809 | Q8N2K0-2 | ABHD12 | KPQDQGLNHTCNYYLQPEEDVTIGVWHTVPAVWWK | 11C(57.0214), 8N(1864.6342) | 123 | 2 | 9 | 0 | [illegible] |
| MSFragger | 1732 | 16849 | Q8N766-2 | EMC1 | DSLACFNQTYTINLYLVETGR | 5C(57.0214), 7N(1864.6342) | 369 | 2 | 9 | 0 | [illegible] |
| MSFragger | 1757 | 17509 | P01137 | TGFB1 | LASPPSQGEVPPGPLPEAVLALYNSTR | 24N(1864.6342) | 62 | 2 | 9 | 0 | [illegible] |
| MSFragger | 1767 | 17754 | Q8TEM1 | NUP210 | RFDNFSSLSIQWESTRPVLASIEPELPMQLVSQDDESGQK | 4N(1864.6342) | 801 | 2 | 9 | 0 | [illegible] |
| MSFragger | 1772 | 17892 | P34810-2 | CD68 | DLQAPLGQSFSCSNSSIILSPAVHLDLLSLR | 12C(57.0214), 14N(1864.6342) | 212 | 2 | 9 | 0 | [illegible] |
| MSFragger | 1779 | 18129 | Q32P28-3 | P3H1 | EKPFEDFLPSHYNYLQFAYYNIGNYTQAVECAK | 31C(57.0214), 24N(1864.6342) | 296 | 2 | 9 | 0 | [illegible] |
| MSFragger | 1783 | 18207 | Q6NUS6-2 | TCTN3 | VNATNFQALAAEFGGESFTSTFQTQSPPSFYR | 2N(1864.6342) | 179 | 2 | 9 | 0 | [illegible] |
| MSFragger | 1790 | 18414 | Q9H330-3 | TMEM245 | VHQESVHMIEVTSNLINETLANHPEWANWLPEAQVVQR | 17N(1864.6342) | 42 | 2 | 9 | 0 | [illegible] |
| MSFragger | 1792 | 18837 | Q9UHG3-2 | PCYOX1 | KMSNITFLNFDPPIEEFHQYYQHIVTTLVK | 4N(1864.6342) | 246 | 2 | 9 | 0 | [illegible] |
| MSFragger | 1793 | 18842 | Q68CR1-2 | SEL1L3 | NVSEVSVEYLCSQPCVVNLEAVVSSEFR | 11C(57.0214), 15C(57.0214), 1N(1864.6342) | 60 | 2 | 9 | 0 | [illegible] |
| MSFragger | 1811 | 19635 | Q9Y4L1 | HYOU1 | QTVHFQISSQLQFSPEEVLGMVLNYSR | 24N(1864.6342) | 135 | 2 | 9 | 0 | [illegible] |
| GRable | 1907 | 6216 | Q9H173 | SIL1 | FNSSSSSLEEK | 0.02000000000.0 | 193 | 2 | 9 | 0 | [illegible] |
| Byonic | 2 | 5039 | P49207 | RPL34 | LSYNTASNK | N8(HexNAc(2)Hex(8)) | 18 | 2 | 8 | 0 | [illegible] |
| Byonic | 3 | 15023 | Q6ZMG9 | CERS6 | FWLPHNVTWADLK | N6(HexNAc(2)Hex(8)) | 18 | 2 | 8 | 0 | [illegible] |
| Byonic | 4 | 17231 | Q96G23 | CERS2 | LWLPVNLTWADLEDRDGR | N6(HexNAc(2)Hex(8)) | 19 | 2 | 8 | 0 | [illegible] |
| Byonic | 9 | 8449 | Q8IWB1 | ITPRIP | ENATVPENEEEIIRK | N2(HexNAc(2)Hex(8)) | 27 | 2 | 8 | 0 | [illegible] |
| Byonic | 22 | 11184 | Q8TEM1 | NUP210 | VNFTLEASEGCYR | N2(HexNAc(2)Hex(8)); C11(Carbamidomethyl) | 44 | 2 | 8 | 0 | [illegible] |
| Byonic | 31 | 4135 | P01871 | IGHM | YKNNSDISSTR | N3(HexNAc(2)Hex(8)) | 46 | 2 | 8 | 0 | [illegible] |
| Byonic | 56 | 3955 | P01871 | IGHM | NNSDISSTR | N1(HexNAc(2)Hex(8)) | 46 | 2 | 8 | 0 | [illegible] |
| Byonic | 73 | 11713 | P43121 | MCAM | CGLSQSQGNLSHVDWFSVHK | C1(Carbamidomethyl); N9(HexNAc(2)Hex(8)) | 56 | 2 | 8 | 0 | [illegible] |
| Byonic | 74 | 10721 | P05107 | ITGB2 | LNFTGPGDPDSIR | N2(HexNAc(2)Hex(8)) | 50 | 2 | 8 | 0 | [illegible] |
| Byonic | 88 | 6009 | P20645 | M6PR | RLKPLFNK | N7(HexNAc(2)Hex(8)) | 57 | 2 | 8 | 0 | [illegible] |
| Byonic | 89 | 7070 | P20645 | M6PR | LKPLFNK | N6(HexNAc(2)Hex(8)) | 57 | 2 | 8 | 0 | [illegible] |
| Byonic | 98 | 9599 | P53801 | PTTG1IP | NVSCLWCNTNK | N1(HexNAc(2)Hex(8)); C4(Carbamidomethyl); C7(Carbamidomethyl) | 54 | 2 | 8 | 0 | [illegible] |
| Byonic | 107 | 8809 | Q16563 | SYPL1 | GQTEIQVNCPPAVTENK | C9(Carbamidomethyl); N16(HexNAc(2)Hex(8)) | 71 | 2 | 8 | 0 | [illegible] |
| Byonic | 111 | 16637 | P13284 | IFI30 | SNAPLVNVTLYYEALCGGCR | N7(HexNAc(2)Hex(8)); C16(Carbamidomethyl); C19(Carbamidomethyl) | 63 | 2 | 8 | 0 | [illegible] |
| Byonic | 114 | 17627 | Q92478 | CLEC2B | YNCSTQHADLTIIDNIEEMNFLR | N2(HexNAc(2)Hex(8)); C3(Carbamidomethyl) | 62 | 2 | 8 | 0 | [illegible] |
| Byonic | 122 | 13419 | P40259 | CD79B | MHCYMNSASGNVSWLWK | C3(Carbamidomethyl); N11(HexNAc(2)Hex(8)) | 73 | 2 | 8 | 0 | [illegible] |
| Byonic | 125 | 8138 | Q9NR96 | TLR9 | GNVTSLSLSSNR | N2(HexNAc(2)Hex(8)) | 64 | 2 | 8 | 0 | [illegible] |
| Byonic | 129 | 13724 | O95158 | NXPH4 | AWSWAWPTNHTGALAR | N9(HexNAc(2)Hex(8)) | 72 | 2 | 8 | 0 | [illegible] |
| Byonic | 130 | 11950 | Q08722 | CD47 | DIYTFDGALNK | N10(HexNAc(2)Hex(8)) | 73 | 2 | 8 | 0 | [illegible] |
| Byonic | 134 | 15539 | Q9H0U3 | MAGT1 | APPRNYSVIVMFTALQLHR | N5(HexNAc(2)Hex(8)) | 71 | 2 | 8 | 0 | [illegible] |
| Byonic | 135 | 11452 | Q9H5K3 | POMK | NCSPWLSCEELR | N1(HexNAc(2)Hex(8)); C2(Carbamidomethyl); C8(Carbamidomethyl) | 67 | 2 | 8 | 0 | [illegible] |
| Byonic | 143 | 8688 | P01591 | JCHAIN | ENISDPTSPLR | N2(HexNAc(2)Hex(8)) | 71 | 2 | 8 | 0 | [illegible] |
| Byonic | 144 | 11580 | O15342 | ATP6V0E1 | NETIWYLK | N1(HexNAc(2)Hex(8)) | 70 | 2 | 8 | 0 | [illegible] |
| Byonic | 146 | 16267 | Q6UW56 | ATRAID | GTILGLDLQNCSLEDPGPNFHQAHTTVIIDLQANPLK | N10(HexNAc(2)Hex(8)); C11(Carbamidomethyl) | 79 | 2 | 8 | 0 | [illegible] |
| Byonic | 148 | 10588 | Q9NY35 | CLDND1 | YNGTVGLWR | N2(HexNAc(2)Hex(8)) | 72 | 2 | 8 | 0 | [illegible] |
| Byonic | 156 | 11191 | Q9BRN9 | TM2D3 | LPADCIDCTTNFSCTYGK | C5(Carbamidomethyl); C8(Carbamidomethyl); N11(HexNAc(2)Hex(8)); C14(Carbamidomethyl) | 87 | 2 | 8 | 0 | [illegible] |
| Byonic | 181 | 9038 | P43308 | SSR2 | IAPASNVSHTVVLRPLK | N6(HexNAc(2)Hex(8)) | 88 | 2 | 8 | 0 | [illegible] |
| Byonic | 184 | 7977 | P43308 | SSR2 | IAPASNVSHTVVLR | N6(HexNAc(2)Hex(8)) | 88 | 2 | 8 | 0 | [illegible] |
| Byonic | 196 | 7148 | Q96KA5 | CLPTM1L | TVNVSVPK | N3(HexNAc(2)Hex(8)) | 91 | 2 | 8 | 0 | [illegible] |
| Byonic | 205 | 6470 | Q14738 | PPP2R5D | FNLSK | N2(HexNAc(2)Hex(8)) | 93 | 2 | 8 | 0 | [illegible] |
| Byonic | 206 | 12004 | Q9BX74 | TM2D1 | DSSGNETHFTGNEVGFFKPISCR | N5(HexNAc(2)Hex(8)); C22(Carbamidomethyl) | 96 | 2 | 8 | 0 | [illegible] |
| Byonic | 207 | 17148 | Q86WD7 | SERPINA9 | TQILQGLGFNLTHTPESAIHQGFQHLVHSLTVPSK | N10(HexNAc(2)Hex(8)) | 101 | 2 | 8 | 0 | [illegible] |
| Byonic | 221 | 13342 | P40259 | CD79B | MEESQNESLATLTIQGIR | N6(HexNAc(2)Hex(8)) | 101 | 2 | 8 | 0 | [illegible] |
| Byonic | 238 | 9922 | P11279 | LAMP1 | GHTLTLNFTR | N7(HexNAc(2)Hex(8)) | 103 | 2 | 8 | 0 | [illegible] |
| Byonic | 242 | 15139 | Q6P179 | ERAP2 | IEVLVSNATQFIILHSK | N7(HexNAc(2)Hex(8)) | 103 | 2 | 8 | 0 | [illegible] |
| Byonic | 260 | 15613 | Q96KA5 | CLPTM1L | NNGTLYAYIFLHHAGVLPWHDGK | N2(HexNAc(2)Hex(8)) | 101 | 2 | 8 | 0 | [illegible] |
| Byonic | 262 | 8952 | Q7Z7H5 | TMED4 | FTFTSHTPGDHQICLHSNSTR | C14(Carbamidomethyl); N18(HexNAc(2)Hex(8)) | 117 | 2 | 8 | 0 | [illegible] |
| Byonic | 264 | 18125 | P43308 | SSR2 | AGYFNFTSATITYLAQEDGPVVIGSTSAPGQGGILAQR | N5(HexNAc(2)Hex(8)) | 104 | 2 | 8 | 0 | [illegible] |
| Byonic | 282 | 15934 | P01903 | HLA-DRA | SNYTPITNVPPEVTVLTNSPVELR | N2(HexNAc(2)Hex(8)) | 103 | 2 | 8 | 0 | [illegible] |
| Byonic | 284 | 14364 | P05107 | ITGB2 | VTLYLRPGQAAAFNVTFR | N14(HexNAc(2)Hex(8)) | 116 | 2 | 8 | 0 | [illegible] |
| Byonic | 291 | 5620 | O75477 | ERLIN1 | NYTADYDK | N1(HexNAc(2)Hex(8)) | 106 | 2 | 8 | 0 | [illegible] |
| Byonic | 298 | 6685 | P10321 | HLA-C | GYYNQSEDGSHTLQR | N4(HexNAc(2)Hex(8)) | 110 | 2 | 8 | 0 | [illegible] |
| Byonic | 304 | 8341 | P11912 | CD79A | VQEGNESYQQSCGTYLR | N5(HexNAc(2)Hex(8)); C12(Carbamidomethyl) | 112 | 2 | 8 | 0 | [illegible] |
| Byonic | 308 | 8685 | Q9BVK6 | TMED9 | FTFTSHTPGEHQICLHSNSTK | C14(Carbamidomethyl); N18(HexNAc(2)Hex(8)) | 125 | 2 | 8 | 0 | [illegible] |
| Byonic | 322 | 9357 | P20036 | HLA-DPA1 | SNHTQATNDPPEVTVFPK | N2(HexNAc(2)Hex(8)) | 109 | 2 | 8 | 0 | [illegible] |
| Byonic | 330 | 16924 | P04233 | CD74 | MATPLLMQALPMGALPQGPMQNATK | N22(HexNAc(2)Hex(8)) | 130 | 2 | 8 | 0 | [illegible] |
| Byonic | 351 | 13259 | Q6P4Q7 | CNNM4 | DLVVQQLVNVSR | N9(HexNAc(2)Hex(8)) | 122 | 2 | 8 | 0 | [illegible] |
| Byonic | 355 | 12553 | Q9UI09 | NDUFA12 | FNVTGTPEQYVPYSTTR | N2(HexNAc(2)Hex(8)) | 116 | 2 | 8 | 0 | [illegible] |
| Byonic | 362 | 15154 | Q9H497 | TOR3A | ISNNFTGLEWDLNVR | N4(HexNAc(2)Hex(8)) | 122 | 2 | 8 | 0 | [illegible] |
| Byonic | 382 | 8156 | P40259 | CD79B | CNNTSEVYQGCGTELR | C1(Carbamidomethyl); N2(HexNAc(2)Hex(8)); C11(Carbamidomethyl) | 127 | 2 | 8 | 0 | [illegible] |
| Byonic | 386 | 17415 | P43307 | SSR1 | YPQDYQFYIQNFTALPLNTVVPPQR | N11(HexNAc(2)Hex(8)) | 136 | 2 | 8 | 0 | [illegible] |
| Byonic | 400 | 5911 | P08962 | CD63 | NNHTASILDR | N2(HexNAc(2)Hex(8)) | 130 | 2 | 8 | 0 | [illegible] |
| Byonic | 425 | 14384 | P04233 | CD74 | YGNMTEDHVMHLLQNADPLK | N3(HexNAc(2)Hex(8)) | 136 | 2 | 8 | 0 | [illegible] |
| Byonic | 433 | 15466 | P01903 | HLA-DRA | FTPPVVNVTWLR | N7(HexNAc(2)Hex(8)) | 143 | 2 | 8 | 0 | [illegible] |
| Byonic | 437 | 10626 | Q9UIQ6 | LNPEP | KNQSIGLIQPFATNGK | N2(HexNAc(2)Hex(8)) | 145 | 2 | 8 | 0 | [illegible] |
| Byonic | 443 | 5123 | Q9BRN9 | TM2D3 | YPANCTVR | N4(HexNAc(2)Hex(8)); C5(Carbamidomethyl) | 157 | 2 | 8 | 0 | [illegible] |
| Byonic | 447 | 12265 | Q969N2 | PIGT | PLGLANDTDHYFLR | N6(HexNAc(2)Hex(8)) | 164 | 2 | 8 | 0 | [illegible] |
| Byonic | 454 | 5571 | P11117 | ACP2 | YEQLQNETR | N6(HexNAc(2)Hex(8)) | 167 | 2 | 8 | 0 | [illegible] |
| Byonic | 460 | 7154 | Q92896 | GLG1 | LNLTTDPK | N2(HexNAc(2)Hex(8)) | 165 | 2 | 8 | 0 | [illegible] |
| Byonic | 470 | 10452 | Q13586 | STIM1 | LAVTNTTMTGTVLK | N5(HexNAc(2)Hex(8)) | 171 | 2 | 8 | 0 | [illegible] |
| Byonic | 474 | 6276 | O60637 | TSPAN3 | NQSVPLSCCR | N1(HexNAc(2)Hex(8)); C8(Carbamidomethyl); C9(Carbamidomethyl) | 167 | 2 | 8 | 0 | [illegible] |
| Byonic | 489 | 6319 | P25942 | CD40 | DLVVQQAGTNK | N10(HexNAc(2)Hex(8)) | 180 | 2 | 8 | 0 | [illegible] |
| Byonic | 491 | 11342 | Q96BZ4 | PLD4 | NISLAVATSSPTLAR | N1(HexNAc(2)Hex(8)) | 171 | 2 | 8 | 0 | [illegible] |
| Byonic | 510 | 17331 | P43307 | SSR1 | DLNGNVFQDAVFNQTVTVIER | N13(HexNAc(2)Hex(8)) | 191 | 2 | 8 | 0 | [illegible] |
| Byonic | 512 | 14125 | Q9NUM4 | TMEM106B | LNNITIIGPLDMK | N3(HexNAc(2)Hex(8)) | 183 | 2 | 8 | 0 | [illegible] |
| Byonic | 513 | 15302 | Q9UHG3 | PCYOX1 | LLHALGGDDFLGMLNR | N15(HexNAc(2)Hex(8)) | 196 | 2 | 8 | 0 | [illegible] |
| Byonic | 515 | 18157 | A2RU67 | FAM234B | PAANLVCLSGMNGSTLWSSLLPEEAR | C7(Carbamidomethyl); N12(HexNAc(2)Hex(8)) | 193 | 2 | 8 | 0 | [illegible] |
| Byonic | 519 | 10437 | P48723 | HSPA13 | NSTIEAANLAGLK | N1(HexNAc(2)Hex(8)) | 184 | 2 | 8 | 0 | [illegible] |
| Byonic | 533 | 15275 | Q99467 | CD180 | SLEQAINLSLNFNGNNVK | N7(HexNAc(2)Hex(8)) | 201 | 2 | 8 | 0 | [illegible] |
| Byonic | 535 | 13686 | P28907 | CD38 | FAEAACDVVHVMLNGSR | C6(Carbamidomethyl); N14(HexNAc(2)Hex(8)) | 209 | 2 | 8 | 0 | [illegible] |
| Byonic | 539 | 12635 | Q9HDC9 | APMAP | NMSFVNDLTVTQDGR | N1(HexNAc(2)Hex(8)) | 196 | 2 | 8 | 0 | [illegible] |
| Byonic | 549 | 14988 | Q68CR1 | SEL1L3 | NYTLLQTIPPFERPFK | N1(HexNAc(2)Hex(8)) | 201 | 2 | 8 | 0 | [illegible] |
| Byonic | 569 | 12066 | P01871 | IGHM | GLTFQQNASSMCVPDQDTAIR | N7(HexNAc(2)Hex(8)); C12(Carbamidomethyl) | 209 | 2 | 8 | 0 | [illegible] |
| Byonic | 579 | 5556 | P20138 | CD33 | PQDHGTNLTCQVK | N7(HexNAc(2)Hex(8)); C10(Carbamidomethyl) | 209 | 2 | 8 | 0 | [illegible] |
| Byonic | 593 | 6301 | Q8NBL1 | POGLUT1 | NSTAYFR | N1(HexNAc(2)Hex(8)) | 204 | 2 | 8 | 0 | [illegible] |
| Byonic | 603 | 10224 | P09326 | CD48 | NGTVCLSPPCTLAR | N1(HexNAc(2)Hex(8)); C5(Carbamidomethyl); C10(Carbamidomethyl) | 206 | 2 | 8 | 0 | [illegible] |
| Byonic | 609 | 7472 | P05107 | ITGB2 | LTNNSNQFQTEVGK | N3(HexNAc(2)Hex(8)) | 212 | 2 | 8 | 0 | [illegible] |
| Byonic | 614 | 6445 | Q9HD67 | MYO10 | TVYNNNSSRFGK | N6(HexNAc(2)Hex(8)) | 217 | 2 | 8 | 0 | [illegible] |
| Byonic | 615 | 13697 | P07602 | PSAP | TNSTFVQALVEHVK | N2(HexNAc(2)Hex(8)) | 215 | 2 | 8 | 0 | [illegible] |
| Byonic | 617 | 12523 | P14625 | HSP90B1 | HNNDTQHIWESDSNEFSVIADPR | N3(HexNAc(2)Hex(8)) | 217 | 2 | 8 | 0 | [illegible] |
| Byonic | 618 | 10740 | P28907 | CD38 | IFDKNSTFGSVEVHNLQPEK | N5(HexNAc(2)Hex(8)) | 219 | 2 | 8 | 0 | [illegible] |
| Byonic | 634 | 9385 | P28907 | CD38 | NSTFGSVEVHNLQPEK | N1(HexNAc(2)Hex(8)) | 219 | 2 | 8 | 0 | [illegible] |
| Byonic | 637 | 8161 | Q96KA5 | CLPTM1L | VKDLMVINR | N8(HexNAc(2)Hex(8)) | 229 | 2 | 8 | 0 | [illegible] |
| Byonic | 643 | 9249 | Q96KA5 | CLPTM1L | DLMVINR | N6(HexNAc(2)Hex(8)) | 229 | 2 | 8 | 0 | [illegible] |
| Byonic | 644 | 16406 | O96005 | CLPTM1 | RAEDYGPVEVISHWHPNITINIVDDHTPWVK | N17(HexNAc(2)Hex(8)) | 241 | 2 | 8 | 0 | [illegible] |
| Byonic | 651 | 5372 | Q9Y639 | NPTN | ANATIEVK | N2(HexNAc(2)Hex(8)) | 229 | 2 | 8 | 0 | [illegible] |
| Byonic | 663 | 5000 | Q5JRA6 | MIA3 | LKVPESENNK | N9(HexNAc(2)Hex(8)) | 246 | 2 | 8 | 0 | [illegible] |
| Byonic | 670 | 11053 | Q14108 | SCARB2 | CNMINGTDGDSFHPLITK | C1(Carbamidomethyl); N5(HexNAc(2)Hex(8)) | 249 | 2 | 8 | 0 | [illegible] |
| Byonic | 673 | 7521 | O75326 | SEMA7A | EDNPDKNPEAPLNVSR | N13(HexNAc(2)Hex(8)) | 258 | 2 | 8 | 0 | [illegible] |
| Byonic | 679 | 9844 | Q96JJ7 | TMX3 | LVALAVIDEKNTSVEHTR | N11(HexNAc(2)Hex(8)) | 258 | 2 | 8 | 0 | [illegible] |
| Byonic | 705 | 7438 | Q13510 | ASAH1 | TVLENSTSYEEAK | N5(HexNAc(2)Hex(8)) | 259 | 2 | 8 | 0 | [illegible] |
| Byonic | 713 | 7924 | P11279 | LAMP1 | LLNINPNK | N7(HexNAc(2)Hex(8)) | 261 | 2 | 8 | 0 | [illegible] |
| Byonic | 717 | 15224 | Q9UBV2 | SEL1L | GQTALGFLYASGLGVNSSQAK | N16(HexNAc(2)Hex(8)) | 272 | 2 | 8 | 0 | [illegible] |
| Byonic | 725 | 4998 | P35613 | BSG | ALMNGSESR | N4(HexNAc(2)Hex(8)) | 268 | 2 | 8 | 0 | [illegible] |
| Byonic | 735 | 9904 | P11117 | ACP2 | KNLTLMATTSQLPK | N2(HexNAc(2)Hex(8)) | 267 | 2 | 8 | 0 | [illegible] |
| Byonic | 757 | 16123 | Q8NAT1 | POMGNT2 | LNVSHTGVPLGEEYILVFSR | N2(HexNAc(2)Hex(8)) | 276 | 2 | 8 | 0 | [illegible] |
| Byonic | 759 | 8868 | Q96HE7 | ERO1A | WGHNITEFQQR | N4(HexNAc(2)Hex(8)) | 280 | 2 | 8 | 0 | [illegible] |
| Byonic | 783 | 16675 | P04843 | RPN1 | TILPAAAQDVYYRDEIGNVSTSHLLILDDSVEMEIRPR | N18(HexNAc(2)Hex(8)) | 299 | 2 | 8 | 0 | [illegible] |
| Byonic | 793 | 15354 | O96005 | CLPTM1 | DYYPINESLASLPLR | N6(HexNAc(2)Hex(8)) | 295 | 2 | 8 | 0 | [illegible] |
| Byonic | 799 | 7608 | Q6P179 | ERAP2 | NQTHYALQASLK | N1(HexNAc(2)Hex(8)) | 294 | 2 | 8 | 0 | [illegible] |
| Byonic | 802 | 15430 | P04843 | RPN1 | DEIGNVSTSHLLILDDSVEMEIRPR | N5(HexNAc(2)Hex(8)) | 299 | 2 | 8 | 0 | [illegible] |
| Byonic | 804 | 17083 | Q9UP95 | SLC12A4 | SIFDPPVFPVCMLGNR | C11(Carbamidomethyl); N15(HexNAc(2)Hex(8)) | 312 | 2 | 8 | 0 | [illegible] |
| Byonic | 813 | 7132 | Q9NXH8 | TOR4A | FVLQNASR | N5(HexNAc(2)Hex(8)) | 314 | 2 | 8 | 0 | [illegible] |
| Byonic | 817 | 11814 | O75764 | TCEA3 | KNCTYNQVQTR | N2(HexNAc(2)Hex(8)); C3(Carbamidomethyl) | 317 | 2 | 8 | 0 | [illegible] |
| Byonic | 844 | 15070 | Q8TEM1 | NUP210 | LPNSTIYVVEPGYLGFTVHPGDR | N3(HexNAc(2)Hex(8)) | 337 | 2 | 8 | 0 | [illegible] |
| Byonic | 851 | 4906 | Q96LC7 | SIGLEC10 | VMVSQANR | N7(HexNAc(2)Hex(8)) | 355 | 2 | 8 | 0 | [illegible] |
| Byonic | 853 | 11102 | Q9UHG3 | PCYOX1 | GELNTSIFSSR | N4(HexNAc(2)Hex(8)) | 353 | 2 | 8 | 0 | [illegible] |
| Byonic | 855 | 11089 | Q9UHG3 | PCYOX1 | GELNTSIFSSRPIDK | N4(HexNAc(2)Hex(8)) | 353 | 2 | 8 | 0 | [illegible] |
| Byonic | 869 | 8054 | P13473 | LAMP2 | VQPFNVTQGK | N5(HexNAc(2)Hex(8)) | 356 | 2 | 8 | 0 | [illegible] |
| Byonic | 886 | 4337 | Q96SB8 | SMC6 | AYNEAEVLYNR | N10(HexNAc(2)Hex(8)) | 368 | 2 | 8 | 0 | [illegible] |
| Byonic | 892 | 6377 | Q4KMQ2 | ANO6 | LNITCESSK | N2(HexNAc(2)Hex(8)); C5(Carbamidomethyl) | 361 | 2 | 8 | 0 | [illegible] |
| Byonic | 896 | 16137 | Q9Y6X5 | ENPP4 | HSTINIVDIYPMMCHILGLKPHPNNGTFGHTK | C14(Carbamidomethyl); N25(HexNAc(2)Hex(8)) | 386 | 2 | 8 | 0 | [illegible] |
| Byonic | 898 | 16833 | Q8N766 | EMC1 | DSLACFNQTYTINLYLVETGR | C5(Carbamidomethyl); N7(HexNAc(2)Hex(8)) | 370 | 2 | 8 | 0 | [illegible] |
| Byonic | 905 | 9075 | Q5JPE7 | NOMO2 | LENITTGTYTIHAQK | N3(HexNAc(2)Hex(8)) | 369 | 2 | 8 | 0 | [illegible] |
| Byonic | 909 | 9567 | Q96S52 | PIGS | TYNASVLPVR | N3(HexNAc(2)Hex(8)) | 370 | 2 | 8 | 0 | [illegible] |
| Byonic | 922 | 7072 | Q68CR1 | SEL1L3 | SYHNQTISFR | N4(HexNAc(2)Hex(8)) | 382 | 2 | 8 | 0 | [illegible] |
| Byonic | 927 | 13377 | Q6NSJ0 | MYORG | VTLWVHPFVNYNSSR | N12(HexNAc(2)Hex(8)) | 398 | 2 | 8 | 0 | [illegible] |
| Byonic | 950 | 12462 | Q9UBV2 | SEL1L | MYSEGSDIVPQSNETALHYFK | N13(HexNAc(2)Hex(8)) | 431 | 2 | 8 | 0 | [illegible] |
| Byonic | 958 | 8465 | Q969V3 | NCLN | VIYNLTEK | N4(HexNAc(2)Hex(8)) | 428 | 2 | 8 | 0 | [illegible] |
| Byonic | 963 | 5065 | Q6P179 | ERAP2 | DSLNSSRPISK | N4(HexNAc(2)Hex(8)) | 431 | 2 | 8 | 0 | [illegible] |
| Byonic | 970 | 9189 | Q92542 | NCSTN | NISGVVLADHSGAFHNK | N1(HexNAc(2)Hex(8)) | 435 | 2 | 8 | 0 | [illegible] |
| Byonic | 972 | 8242 | O14672 | ADAM10 | NISQVLEK | N1(HexNAc(2)Hex(8)) | 439 | 2 | 8 | 0 | [illegible] |
| Byonic | 989 | 6179 | P08575 | PTPRC | NGSAAMCHFTTK | N1(HexNAc(2)Hex(8)); C7(Carbamidomethyl) | 470 | 2 | 8 | 0 | [illegible] |
| Byonic | 999 | 14270 | Q9Y487 | ATP6V0A2 | SVNLFGSGWNVSAMYSSSHPPAEHK | N10(HexNAc(2)Hex(8)) | 484 | 2 | 8 | 0 | [illegible] |
| Byonic | 1026 | 10835 | P05107 | ITGB2 | KDNNSIICSGLGDCVCGQCLCHTSDVPGK | N3(HexNAc(2)Hex(8)); C8(Carbamidomethyl); C14(Carbamidomethyl); C16(Carbamidomethyl); C19(Carbamidomethyl); C21(Carbamidomethyl) | 501 | 2 | 8 | 0 | [illegible] |
| Byonic | 1035 | 11928 | P05107 | ITGB2 | DNNSIICSGLGDCVCGQCLCHTSDVPGK | N2(HexNAc(2)Hex(8)); C7(Carbamidomethyl); C13(Carbamidomethyl); C15(Carbamidomethyl); C18(Carbamidomethyl); C20(Carbamidomethyl) | 501 | 2 | 8 | 0 | [illegible] |
| Byonic | 1042 | 9259 | Q9Y4L1 | HYOU1 | VFGSQNLTTVK | N6(HexNAc(2)Hex(8)) | 515 | 2 | 8 | 0 | [illegible] |
| Byonic | 1048 | 7111 | P05556 | ITGB1 | KENSSEICSNNGECVCGQCVCR | N3(HexNAc(2)Hex(8)); C8(Carbamidomethyl); C14(Carbamidomethyl); C16(Carbamidomethyl); C19(Carbamidomethyl); C21(Carbamidomethyl) | 520 | 2 | 8 | 0 | [illegible] |
| Byonic | 1054 | 17375 | P46977 | STT3A | VMSWWDYGYQITAMANR | N16(HexNAc(2)Hex(8)) | 537 | 2 | 8 | 0 | [illegible] |
| Byonic | 1059 | 5697 | Q68CR1 | SEL1L3 | NQNESVSEIGGK | N3(HexNAc(2)Hex(8)) | 527 | 2 | 8 | 0 | [illegible] |
| Byonic | 1063 | 5580 | P26010 | ITGB7 | APNGTGPLCSGK | N3(HexNAc(2)Hex(8)); C9(Carbamidomethyl) | 531 | 2 | 8 | 0 | [illegible] |
| Byonic | 1067 | 5043 | P78536 | ADAM17 | CQEAINATCK | C1(Carbamidomethyl); N6(HexNAc(2)Hex(8)); C9(Carbamidomethyl) | 539 | 2 | 8 | 0 | [illegible] |
| Byonic | 1069 | 10382 | P46977 | STT3A | TILVDNNTWNNTHISR | N10(HexNAc(2)Hex(8)) | 548 | 2 | 8 | 0 | [illegible] |
| Byonic | 1072 | 5226 | Q9H330 | TMEM245 | VNNTAVIEK | N2(HexNAc(2)Hex(8)) | 551 | 2 | 8 | 0 | [illegible] |
| Byonic | 1080 | 12126 | P11717 | IGF2R | TNITLVCKPGDLESAPVLR | N2(HexNAc(2)Hex(8)); C7(Carbamidomethyl) | 581 | 2 | 8 | 0 | [illegible] |
| Byonic | 1085 | 15918 | P26006 | ITGA3 | SLDAYPILNQAQALENHTEVQFQK | N16(HexNAc(2)Hex(8)) | 605 | 2 | 8 | 0 | [illegible] |
| Byonic | 1086 | 16666 | Q9Y2U8 | LEMD3 | CVGFGPEEELTNITDVQFLQSTR | C1(Carbamidomethyl); N12(HexNAc(2)Hex(8)) | 604 | 2 | 8 | 0 | [illegible] |
| Byonic | 1088 | 6537 | Q9Y4L1 | HYOU1 | ENGTDTVQEEEESPAEGSK | N2(HexNAc(2)Hex(8)) | 596 | 2 | 8 | 0 | [illegible] |
| Byonic | 1091 | 8546 | Q9UBV2 | SEL1L | EASIVGENETYPR | N8(HexNAc(2)Hex(8)) | 608 | 2 | 8 | 0 | [illegible] |
| Byonic | 1092 | 17447 | Q8TCJ2 | STT3B | VMSWWDYGYQIAGMANR | N16(HexNAc(2)Hex(8)) | 616 | 2 | 8 | 0 | [illegible] |
| Byonic | 1095 | 9962 | Q5JPE7 | NOMO2 | ENVGIYNLSK | N7(HexNAc(2)Hex(8)) | 618 | 2 | 8 | 0 | [illegible] |
| Byonic | 1098 | 12084 | Q9BU23 | LMF2 | SANSTLAQALHWTR | N3(HexNAc(2)Hex(8)) | 616 | 2 | 8 | 0 | [illegible] |
| Byonic | 1102 | 11792 | Q8TCJ2 | STT3B | TTLVDNNTWNNSHIALVGK | N10(HexNAc(2)Hex(8)) | 627 | 2 | 8 | 0 | [illegible] |
| Byonic | 1107 | 7369 | Q9UH99 | SUN2 | ALSPNSTISSAPK | N5(HexNAc(2)Hex(8)) | 636 | 2 | 8 | 0 | [illegible] |
| Byonic | 1111 | 5441 | Q8TCJ2 | STT3B | AMSSNETAAYK | N5(HexNAc(2)Hex(8)) | 641 | 2 | 8 | 0 | [illegible] |
| Byonic | 1117 | 9541 | P05107 | ITGB2 | NCSAACPGLQLSNNPVK | N1(HexNAc(2)Hex(8)); C2(Carbamidomethyl); C6(Carbamidomethyl) | 642 | 2 | 8 | 0 | [illegible] |
| Byonic | 1125 | 15916 | Q8TEM1 | NUP210 | FFQNVTAEDTDSIGLALFAPHSSR | N4(HexNAc(2)Hex(8)) | 681 | 2 | 8 | 0 | [illegible] |
| Byonic | 1133 | 7082 | Q9NR96 | TLR9 | ALTNGSLPAGTR | N4(HexNAc(2)Hex(8)) | 694 | 2 | 8 | 0 | [illegible] |
| Byonic | 1135 | 16337 | A2VDJ0 | TMEM131L | NNLTVIDMIGVEGFGAR | N2(HexNAc(2)Hex(8)) | 709 | 2 | 8 | 0 | [illegible] |
| Byonic | 1138 | 9940 | P02786 | TFRC | KQNNGAFNETLFR | N8(HexNAc(2)Hex(8)) | 727 | 2 | 8 | 0 | [illegible] |
| Byonic | 1143 | 11406 | P02786 | TFRC | QNNGAFNETLFR | N7(HexNAc(2)Hex(8)) | 727 | 2 | 8 | 0 | [illegible] |
| Byonic | 1163 | 5515 | Q9Y6M7 | SLC4A7 | NLTVSECK | N1(HexNAc(2)Hex(8)); C7(Carbamidomethyl) | 796 | 2 | 8 | 0 | [illegible] |
| Byonic | 1166 | 8590 | Q68CR1 | SEL1L3 | NQTLAGEYFHK | N1(HexNAc(2)Hex(8)) | 818 | 2 | 8 | 0 | [illegible] |
| Byonic | 1170 | 16444 | Q9Y4L1 | HYOU1 | LSALDNLLNHSSMFLK | N9(HexNAc(2)Hex(8)) | 830 | 2 | 8 | 0 | [illegible] |
| Byonic | 1175 | 11643 | Q9Y4L1 | HYOU1 | VINETWAWK | N3(HexNAc(2)Hex(8)) | 862 | 2 | 8 | 0 | [illegible] |
| Byonic | 1178 | 10363 | Q92673 | SORL1 | LTIVNSSVLDRPR | N5(HexNAc(2)Hex(8)) | 871 | 2 | 8 | 0 | [illegible] |
| Byonic | 1182 | 4394 | Q9Y4L1 | HYOU1 | NATLAEQAK | N1(HexNAc(2)Hex(8)) | 869 | 2 | 8 | 0 | [illegible] |
| Byonic | 1185 | 8435 | Q8N766 | EMC1 | FINYNQTVSR | N5(HexNAc(2)Hex(8)) | 913 | 2 | 8 | 0 | [illegible] |
| Byonic | 1192 | 6295 | Q9Y4L1 | HYOU1 | AEPPLNASASDQGEK | N6(HexNAc(2)Hex(8)) | 931 | 2 | 8 | 0 | [illegible] |
| Byonic | 1196 | 14053 | P26006 | ITGA3 | VWNSTFIEDYRDFDR | N3(HexNAc(2)Hex(8)) | 935 | 2 | 8 | 0 | [illegible] |
| Byonic | 1197 | 13035 | P26006 | ITGA3 | VWNSTFIEDYR | N3(HexNAc(2)Hex(8)) | 935 | 2 | 8 | 0 | [illegible] |
| Byonic | 1200 | 13075 | O15031 | PLXNB2 | SINVTGQGFSLIQR | N3(HexNAc(2)Hex(8)) | 1002 | 2 | 8 | 0 | [illegible] |
| Byonic | 1209 | 5909 | P11717 | IGF2R | MNFTGGDTCHK | N2(HexNAc(2)Hex(8)); C9(Carbamidomethyl) | 1312 | 2 | 8 | 0 | [illegible] |
| Byonic | 1216 | 4956 | Q8TEM1 | NUP210 | GPTNNTCVVR | N4(HexNAc(2)Hex(8)); C7(Carbamidomethyl) | 1441 | 2 | 8 | 0 | [illegible] |
| Byonic | 1227 | 7613 | Q92673 | SORL1 | GIGNWSDSK | N4(HexNAc(2)Hex(8)) | 1733 | 2 | 8 | 0 | [illegible] |
| Byonic | 1229 | 4934 | Q92673 | SORL1 | SLTTSLHNK | N8(HexNAc(2)Hex(8)) | 1894 | 2 | 8 | 0 | [illegible] |
| Byonic | 1239 | 11006 | Q92673 | SORL1 | YHIIVQLGNMSK | N9(HexNAc(2)Hex(8)) | 2010 | 2 | 8 | 0 | [illegible] |
| Glyco-Decip | 1270 | 5075 | Q92478 | CLC2B | EEGDWNSSK | | 57 | 2 | 8 | 0 | [illegible] |
| Glyco-Decip | 1271 | 5208 | P05556 | ITB1 | CHEGNGTFECGACR | 1,Carbamidomethyl(+57.0215);10,Carbamidomethyl(+57.0215);13,Carbamidomethyl(+57.0215); | 481 | 2 | 8 | 0 | [illegible] |
| Glyco-Decip | 1272 | 5290 | O60449 | LY75 | ENNNITMR | | 1593 | 2 | 8 | 0 | [illegible] |
| Glyco-Decip | 1279 | 5442 | O75900 | MMP23 | NVTFR | | 316 | 2 | 8 | 0 | [illegible] |
| Glyco-Decip | 1296 | 5717 | P06276 | CHLE | DNNSIITR | | 369 | 2 | 8 | 0 | [illegible] |
| Glyco-Decip | 1305 | 6095 | Q9H6U8 | ALG9 | PVNVCVGK | 5,Carbamidomethyl(+57.0215); | 465 | 2 | 8 | 0 | [illegible] |
| Glyco-Decip | 1307 | 6206 | Q9H173 | SIL1 | FNSSSSSLEEK | | 193 | 2 | 8 | 0 | [illegible] |
| Glyco-Decip | 1323 | 6586 | O60568 | PLOD3 | EQYIHENYSR | | 548 | 2 | 8 | 0 | [illegible] |
| Glyco-Decip | 1335 | 7016 | Q96LC7 | SIG10 | GNCSLVIR | 3,Carbamidomethyl(+57.0215); | 100 | 2 | 8 | 0 | [illegible] |
| Glyco-Decip | 1336 | 7042 | P79483 | DRB3 | SECHFFNGTER | 3,Carbamidomethyl(+57.0215); | 48 | 2 | 8 | 0 | [illegible] |
| Glyco-Decip | 1345 | 7233 | P42892 | ECE1 | HLLENSTASVSEAER | | 166 | 2 | 8 | 0 | [illegible] |
| Glyco-Decip | 1368 | 8188 | Q9BU23 | LMF2 | PGNLSRPPPVVVPHQPR | | 489 | 2 | 8 | 0 | [illegible] |
| Glyco-Decip | 1386 | 8697 | O14672 | ADA10 | INTTADEKDPTNPFR | | 278 | 2 | 8 | 0 | [illegible] |
| Glyco-Decip | 1410 | 9611 | P04233 | HG2A | GHHNCSESLELEDPSSGLGVTK | 5,Carbamidomethyl(+57.0215); | 270 | 2 | 8 | 0 | [illegible] |
| Glyco-Decip | 1415 | 9763 | Q92542 | NICA | IYIPLNK | | 45 | 2 | 8 | 0 | [illegible] |
| Glyco-Decip | 1451 | 12120 | P54709 | AT1B3 | NLTVCPDGALFEQK | 5,Carbamidomethyl(+57.0215); | 124 | 2 | 8 | 0 | [illegible] |
| Glyco-Decip | 1474 | 13455 | O00584 | RNT2 | AYWPDVIHSFPNR | | 106 | 2 | 8 | 0 | [illegible] |
| Glyco-Decip | 1483 | 14238 | Q0P6H9 | TMM62 | SNHTIWFGHFTTSTILSPSPGIR | | 223 | 2 | 8 | 0 | [illegible] |
| Glyco-Decip | 1495 | 15095 | Q9BY67 | CADM1 | FQLLNFSSSELK | | 101 | 2 | 8 | 0 | [illegible] |
| Glyco-Decip | 1499 | 15560 | P11912-2 | CD79A | VLHGNYTWPPEFLGPGEDPNEPPPRPFLDMGEGTK | | 73 | 2 | 8 | 0 | [illegible] |
| Glyco-Decip | 1514 | 17996 | P11215 | ITAM | LNFSLVGTPLSAFGNLRPVLAEDAQR | | 734 | 2 | 8 | 0 | [illegible] |
| Glyco-Decip | 1519 | 18440 | Q9NYU2 | UGGG1 | GTEVNTTVIGENDPIDEVQGFLFGK | | 269 | 2 | 8 | 0 | [illegible] |
| Glyco-Decip | 1526 | 19092 | Q9H6E4 | CC134 | WGISFCNQTGVFNQGPHSPILSLMAQELGISEK | 6,Carbamidomethyl(+57.0215); | 148 | 2 | 8 | 0 | [illegible] |
| Glyco-Decip | 1528 | 19128 | Q9NSC7 | SIA7A | LEHFAPPFGFMELNYSLVQKVVTR | 11,Oxidation(+15.9949); | 331 | 2 | 8 | 0 | [illegible] |
| pGlyco567 | 1565 | 10438 | Q96FM1 | PGAP3 | DCVLQCEEQNCSGGALNHFR | 2,Carbamidomethyl[C];6,Carbamidomethyl[C];11,Carbamidomethyl[C]; | 40 | 2 | 8 | 0 | [illegible] |
| pGlyco576 | 1567 | 10540 | Q08722 | CD47 | SDAVSHTGNYTCEVTELTR | 12,Carbamidomethyl[C]; | 111 | 2 | 8 | 0 | [illegible] |
| pGlyco623 | 1571 | 11352 | Q9HDC9 | APMAP | AGPNGTLFVADAYK | | 160 | 2 | 8 | 0 | [illegible] |
| pGlyco695 | 1576 | 12334 | Q10589 | BST2 | NVTHLLQQELTEAQK | | 65 | 2 | 8 | 0 | [illegible] |
| pGlyco779 | 1581 | 14367 | Q5VTE0 | EEF1A1P5 | VETGVLKPGMVVTFAPVNVTTEVK | | 284 | 2 | 8 | 0 | [illegible] |
| MSFragger | 1600 | 5478 | P08575-10 | PTPRC | NASVSISHNSCTAPDK | 11C(57.0214), 1N(1702.5814) | 165 | 2 | 8 | 0 | [illegible] |
| MSFragger | 1606 | 6784 | P18850 | ATF6 | DHLLLPATTHNK | 11N(1702.5814) | 574 | 2 | 8 | 0 | [illegible] |
| MSFragger | 1617 | 7894 | P05556-2 | ITGB1 | ENSSEICSNNGECVCGQCVCR | 13C(57.0214), 15C(57.0214), 18C(57.0214), 20C(57.0214), 7C(57.0214), 2N(1702.5814) | 520 | 2 | 8 | 0 | [illegible] |
| MSFragger | 1631 | 10276 | Q32P28-3 | P3H1 | VPLQSAHLYYNVTEK | 11N(1702.5814) | 530 | 2 | 8 | 0 | [illegible] |
| MSFragger | 1650 | 11731 | O76024 | WFS1 | CLYGEAYPACSPGNTSTAEEELCR | 10C(57.0214), 1C(57.0214), 23C(57.0214), 14N(1702.5814) | 736 | 2 | 8 | 0 | [illegible] |

| | | | | | | | | | | | | | | | | | | | | | |
|---|---|---|---|---|---|---|---|---|---|---|---|---|---|---|---|---|---|---|---|---|---|
| MSFragger | 1663 | 13231 | P12074 | COX6A1 | TKPFPWGDGNHTLFHNPHVNPLPTGYEDE | 10N(1702.5814) | 80 | 2 | 8 | 0 | 0 | 2:8:0:0 | pauci- or high mannose | 0 | 0 | 0 | 1 | 0 | 0 | 89672.5 | 8433.96 |
| MSFragger | 1673 | 13768 | P42785-2 | PRCP | NISSHTNIVFSNGELDPWSGGGVTK | 1N(1702.5814) | 436 | 2 | 8 | 0 | 0 | 2:8:0:0 | pauci- or high mannose | 3 | 0 | 0 | 1 | 0 | 0 | 83839.5 | 7010.36 |
| MSFragger | 1674 | 13789 | P05556-2 | ITGB1 | SCGECIQAGPNCGWCTNSTFLQEGMPTSAR | 12C(57.0214), 15C(57.0214), 2C(57.0214), 5C(57.0214), 17N(1702.5814) | 40 | 2 | 8 | 0 | 0 | 2:8:0:0 | pauci- or high mannose | 1 | 0 | 0 | 1 | 0 | 0 | 119289 | 8698.41 |
| MSFragger | 1698 | 15421 | A2VDJ0-5 | TMEM131L | VQSIQLSQMQAETTNTSLLQVQLECSLHNK | 25C(57.0214), 15N(1702.5814) | 215 | 2 | 8 | 0 | 0 | 2:8:0:0 | pauci- or high mannose | 0 | 0 | 0 | 1 | 0 | 0 | 137811 | 6397.85 |
| MSFragger | 1709 | 16004 | Q8NC56 | LEMD2 | CIPVMEAQEYIANVTSSSSAK | 1C(57.0214), 13N(1702.5814) | 301 | 2 | 8 | 0 | 0 | 2:8:0:0 | pauci- or high mannose | 0 | 0 | 0 | 1 | 0 | 0 | 112086 | 4609.83 |
| MSFragger | 1720 | 16579 | Q10589-2 | BST2 | GFQDVEAQAATCNHTVMALMASLDAEK | 12C(57.0214), 13N(1702.5814) | 70 | 2 | 8 | 0 | 0 | 2:8:0:0 | pauci- or high mannose | 1 | 0 | 0 | 1 | 0 | 0 | 110606 | 8535.98 |
| MSFragger | 1741 | 17041 | P04844 | RPN2 | SNLDPSNVDSLFYAAQASQALSGCEISISNETK | 24C(57.0214), 30N(1702.5814) | 76 | 2 | 8 | 0 | 0 | 2:8:0:0 | pauci- or high mannose | 0 | 0 | 0 | 1 | 0 | 0 | 112073 | 9828.96 |
| MSFragger | 1744 | 17084 | Q9BV10 | ALG12 | VLASVVGTTGVSLNLTQLPPFNVHLQTK | 14N(1702.5814) | 453 | 2 | 8 | 0 | 0 | 2:8:0:0 | pauci- or high mannose | 0 | 0 | 0 | 1 | 0 | 0 | 135004 | 9881.79 |
| MSFragger | 1749 | 17140 | Q86WC4 | OSTM1 | ELLLDFANSSAELTGCLVR | 16C(57.0214), 8N(1702.5814) | 93 | 2 | 8 | 0 | 0 | 2:8:0:0 | pauci- or high mannose | 0 | 0 | 0 | 1 | 0 | 0 | 87548 | 7526.9 |
| MSFragger | 1761 | 17585 | P01137 | TGFB1 | LASPPSQGEVPPGPLPEAVLALYNSTR | 24N(1702.5814) | 62 | 2 | 8 | 0 | 0 | 2:8:0:0 | pauci- or high mannose | 0 | 0 | 0 | 1 | 0 | 0 | 163055 | 14819 |
| MSFragger | 1770 | 17835 | Q8TEM1 | NUP210 | RFDNFSSLSIQWESTRPVLASIEPELPMQLVSQDDESGQK | 4N(1702.5814) | 801 | 2 | 8 | 0 | 0 | 2:8:0:0 | pauci- or high mannose | 0 | 0 | 0 | 1 | 0 | 0 | 156251 | 3833.83 |
| MSFragger | 1775 | 17970 | O14656-2 | TOR1A | IIAENIYEGGLNSDYVHLFVATLHFPHASNITLYK | 30N(1702.5814) | 113 | 2 | 8 | 0 | 0 | 2:8:0:0 | pauci- or high mannose | 0 | 0 | 0 | 1 | 0 | 0 | 85441.4 | 10397.6 |
| MSFragger | 1791 | 18678 | P27797 | CALR | SGTIFDNFLITNDEAYAEEFGNETWGVTK | 22N(1702.5814) | 324 | 2 | 8 | 0 | 0 | 2:8:0:0 | pauci- or high mannose | 1 | 0 | 0 | 1 | 0 | 0 | 120243 | 11217.1 |
| MSFragger | 1794 | 18854 | P61009 | SPCS3 | NVTLTLSWNVVPNAGILPLVTGSGHVSVPFPDTYEITK | 1N(1702.5814) | 141 | 2 | 8 | 0 | 0 | 2:8:0:0 | pauci- or high mannose | 0 | 0 | 0 | 1 | 0 | 0 | 122773 | 8620.22 |
| MSFragger | 1795 | 18855 | Q68CR1-2 | SEL1L3 | NVSEVSVEYLCSQPCVVNLEAVVSSEFR | 11C(57.0214), 15C(57.0214), 1N(1702.5814) | 60 | 2 | 8 | 0 | 0 | 2:8:0:0 | pauci- or high mannose | 0 | 0 | 0 | 1 | 0 | 0 | 99230.6 | 7231.89 |
| MSFragger | 1801 | 19358 | Q9Y2U8 | LEMD3 | DLGPEYEGIFNTSLQWILENGK | 11N(1702.5814) | 566 | 2 | 8 | 0 | 0 | 2:8:0:0 | pauci- or high mannose | 10 | 0 | 0 | 1 | 0 | 0 | 169233 | 13368.4 |
| GRable | 1853 | 7185 | Q13740-2 | GN | TVNSLNVSANENR | 0.0000020000000.0 | 499 | 2 | 8 | 0 | 0 | 2:8:0:0 | pauci- or high mannose | 0 | 0 | 0 | 0 | 0 | 1 | 325403 | 25192.6 |
| GRable | 1875 | 11270 | P35613 | 2 | ILLTCSLNDSATEVTGHR | 0.000000020000000000.0 | 160 | 2 | 8 | 0 | 0 | 2:8:0:0 | pauci- or high mannose | 0 | 0 | 0 | 0 | 0 | 1 | 260554 | 14422.5 |
| GRable | 1876 | 7369 | O94901 | SUN1 | TLSPTGNISSAPK | 0.0000002000000.0 | 732 | 2 | 8 | 0 | 0 | 2:8:0:0 | pauci- or high mannose | 0 | 0 | 0 | 0 | 0 | 1 | 316600 | 24391.7 |
| GRable | 1894 | 7813 | P78536 | 9606 | VVVVDGKNESEYTVK | 0.000000020000000.0 | 103 | 2 | 8 | 0 | 0 | 2:8:0:0 | pauci- or high mannose | 0 | 0 | 0 | 0 | 0 | 1 | 113664 | 4020.3 |
| GRable | 1906 | 15916 | Q6NUS6 | 2 | NLASSCTLDSALNAASYYNFTVLK | 0.00000000000000000000200000.0 | 279 | 2 | 8 | 0 | 0 | 2:8:0:0 | pauci- or high mannose | 0 | 0 | 0 | 0 | 0 | 1 | 266043 | 17606.6 |
| GRable | 1918 | 9546 | P05556 | SV | KDTCTQECSYFNITK | 0.000000000002000.0 | 669 | 2 | 8 | 0 | 0 | 2:8:0:0 | pauci- or high mannose | 0 | 0 | 0 | 0 | 0 | 1 | 204135 | 14799.8 |
| Byonic | 12 | 7275 | P52798 | EFNA4 | HVVYWNSSNPR | N6(HexNAc(2)Hex(7)) | 33 | 2 | 7 | 0 | 0 | 2:7:0:0 | pauci- or high mannose | 1 | 1 | 0 | 0 | 0 | 0 | 82244.5 | 7509.11 |
| Byonic | 24 | 11262 | Q8TEM1 | NUP210 | VNFTLEASEGCYR | N2(HexNAc(2)Hex(7)); C11(Carbamidomethyl) | 44 | 2 | 7 | 0 | 0 | 2:7:0:0 | pauci- or high mannose | 1 | 1 | 1 | 1 | 1 | 0 | 2819.61 | 3910.36 |
| Byonic | 30 | 4156 | P01871 | IGHM | YKNNSDISSTR | N3(HexNAc(2)Hex(7)) | 46 | 2 | 7 | 0 | 0 | 2:7:0:0 | pauci- or high mannose | 1 | 1 | 1 | 1 | 1 | 1 | 1339890 | 74595.9 |
| Byonic | 64 | 3983 | P01871 | IGHM | NNSDISSTR | N1(HexNAc(2)Hex(7)) | 46 | 2 | 7 | 0 | 0 | 2:7:0:0 | pauci- or high mannose | 1 | 1 | 1 | 1 | 1 | 0 | 134505 | 8698.97 |
| Byonic | 81 | 6074 | P20645 | M6PR | RLKPLFNK | N7(HexNAc(2)Hex(7)) | 57 | 2 | 7 | 0 | 0 | 2:7:0:0 | pauci- or high mannose | 1 | 1 | 0 | 0 | 0 | 1 | 181737 | 15663 |
| Byonic | 86 | 6875 | P53634 | CTSC | DVNCSVMGPQEK | N3(HexNAc(2)Hex(7)); C4(Carbamidomethyl) | 53 | 2 | 7 | 0 | 0 | 2:7:0:0 | pauci- or high mannose | 1 | 1 | 1 | 1 | 1 | 1 | 140205 | 9153.42 |
| Byonic | 91 | 7036 | P20645 | M6PR | LKPLFNK | N6(HexNAc(2)Hex(7)) | 57 | 2 | 7 | 0 | 0 | 2:7:0:0 | pauci- or high mannose | 1 | 1 | 1 | 1 | 1 | 1 | 324974 | 19634.1 |
| Byonic | 95 | 5108 | Q92478 | CLEC2B | EEGDWNSSK | N6(HexNAc(2)Hex(7)) | 57 | 2 | 7 | 0 | 0 | 2:7:0:0 | pauci- or high mannose | 1 | 1 | 0 | 0 | 0 | 0 | 129901 | 8739.28 |
| Byonic | 97 | 9639 | P53801 | PTTG1IP | NVSCLWCNTNK | N1(HexNAc(2)Hex(7)); C4(Carbamidomethyl); C7(Carbamidomethyl) | 54 | 2 | 7 | 0 | 0 | 2:7:0:0 | pauci- or high mannose | 1 | 1 | 1 | 1 | 1 | 1 | 1023730 | 77464.6 |
| Byonic | 104 | 8759 | Q16563 | SYPL1 | GQTEIQVNCPPAVTENK | C9(Carbamidomethyl); N16(HexNAc(2)Hex(7)) | 71 | 2 | 7 | 0 | 0 | 2:7:0:0 | pauci- or high mannose | 1 | 1 | 1 | 1 | 1 | 1 | 1590290 | 138600 |
| Byonic | 113 | 17578 | P01137 | TGFB1 | LASPPSQGEVPPGPLPEAVLALYNSTR | N24(HexNAc(2)Hex(7)) | 82 | 2 | 7 | 0 | 0 | 2:7:0:0 | pauci- or high mannose | 1 | 1 | 0 | 1 | 0 | 0 | 158363 | 10681.5 |
| Byonic | 117 | 17673 | Q92478 | CLEC2B | YNCSTQHADLTIIDNIEEMNFLR | N2(HexNAc(2)Hex(7)); C3(Carbamidomethyl) | 62 | 2 | 7 | 0 | 0 | 2:7:0:0 | pauci- or high mannose | 1 | 1 | 0 | 1 | 1 | 0 | 162117 | 15167 |
| Byonic | 119 | 8649 | Q8NBN3 | TMEM87A | NTTIFLK | N1(HexNAc(2)Hex(7)) | 62 | 2 | 7 | 0 | 0 | 2:7:0:0 | pauci- or high mannose | 1 | 1 | 1 | 0 | 0 | 0 | 291676 | 23992.4 |

[illegible]

| | | | | | | | | | | | | | | | | | | | | | | | | | | | | | | |
|---|---|---|---|---|---|---|---|---|---|---|---|---|---|---|---|---|---|---|---|---|---|---|---|---|---|---|---|---|---|---|
| Byonic | 849 | 17057 | Q13510 | ASAH1 | TSQENISFETMYDVLSTKPVLNK | N5(HexNAc(2)Hex(6)Phospho(1)) | 348 | 2 | 6 | 0 | 0 | 2:6:0:0 | pauci- or high mannose | 6 | 1 | 0 | 0 | 0 | 0 | 113699 | 10698.3 | 72407.7 | 36411.5 | 149563 | 2911.17 | 17093.4 | 99134.8 | × | × | [illegible] |
| Byonic | 873 | 8111 | P13473 | LAMP2 | VQPFNVTQGK | N5(HexNAc(2)Hex(6)) | 356 | 2 | 6 | 0 | 0 | 2:6:0:0 | pauci- or high mannose | 1 | 1 | 1 | 1 | 1 | 1 | 360657 | 27462.6 | 240688 | 153405 | 839022 | × | 3248.23 | 250436 | × | × | [illegible] |
| Byonic | 884 | 4371 | Q96SB8 | SMC6 | AYNEAEVLYNR | N10(HexNAc(2)Hex(6)) | 368 | 2 | 6 | 0 | 0 | 2:6:0:0 | pauci- or high mannose | 1 | 1 | 0 | 0 | 0 | 0 | 288728 | 20124.2 | 196088 | 93081.2 | 536234 | × | × | 158326 | × | × | [illegible] |
| Byonic | 897 | 12011 | P15848 | ARSB | GHTNGTKPLDGFDVWK | N4(HexNAc(2)Hex(6)Phospho(1)) | 366 | 2 | 6 | 0 | 0 | 2:6:0:0 | pauci- or high mannose | 1 | 1 | 0 | 1 | 0 | 0 | 212314 | 20403.4 | 125474 | 72534.6 | 275990 | × | 7499.61 | 115942 | × | × | [illegible] |
| Byonic | 899 | 10915 | O00754 | MAN2B1 | ANLTWSVK | N2(HexNAc(2)Hex(6)Phospho(1)) | 367 | 2 | 6 | 0 | 0 | 2:6:0:0 | pauci- or high mannose | 1 | 1 | 1 | 1 | 1 | 0 | 584343 | 66565.1 | 397630 | 257977 | 1351460 | × | 7856.56 | 369151 | × | × | [illegible] |
| Byonic | 904 | 9140 | Q5JPE7 | NOMO2 | LENITTGTYTIHAQK | N3(HexNAc(2)Hex(6)) | 369 | 2 | 6 | 0 | 0 | 2:6:0:0 | pauci- or high mannose | 1 | 1 | 1 | 1 | 1 | 1 | 1159750 | 88795 | 804957 | 465265 | 2148550 | × | 7571.14 | 686027 | × | × | [illegible] |
| Byonic | 908 | 9643 | Q96S52 | PIGS | TYNASVLPVR | N3(HexNAc(2)Hex(6)) | 370 | 2 | 6 | 0 | 0 | 2:6:0:0 | pauci- or high mannose | 1 | 1 | 1 | 1 | 1 | 0 | 348395 | 23809.3 | 223824 | 125603 | 769965 | × | 7230.06 | 214531 | × | × | [illegible] |
| Byonic | 954 | 12486 | Q9UBV2 | SEL1L | MYSEGSDIVPQSNETALHYFK | N13(HexNAc(2)Hex(6)) | 431 | 2 | 6 | 0 | 0 | 2:6:0:0 | pauci- or high mannose | 1 | 1 | 1 | 1 | 1 | 1 | 235744 | 18619.5 | 140691 | 82869.2 | 353353 | 1432.76 | 13056.8 | 134321 | 1549.25 | × | [illegible] |
| Byonic | 959 | 8593 | Q969V3 | NCLN | VIYNLTEK | N4(HexNAc(2)Hex(6)) | 428 | 2 | 6 | 0 | 0 | 2:6:0:0 | pauci- or high mannose | 1 | 1 | 1 | 0 | 0 | 0 | 340585 | 19177.8 | 195534 | 109852 | 473523 | 3933.25 | 17304.7 | 186374 | × | × | [illegible] |
| Byonic | 961 | 12090 | P07602 | PSAP | NSTKQEILAALEK | N1(HexNAc(2)Hex(6)Phospho(1)) | 426 | 2 | 6 | 0 | 0 | 2:6:0:0 | pauci- or high mannose | 1 | 1 | 0 | 1 | 0 | 0 | 289122 | 16618.9 | 192591 | 109012 | 538198 | × | × | 207218 | × | × | [illegible] |
| Byonic | 965 | 14693 | P01871 | IGHM | STGKPTLYNVSLVMSDTAGTCY | N9(HexNAc(2)Hex(6)); C21(Carbamidomethyl) | 440 | 2 | 6 | 0 | 0 | 2:6:0:0 | pauci- or high mannose | 1 | 1 | 1 | 1 | 1 | 1 | 366784 | 30616 | 211370 | 125884 | 669977 | × | 7795.42 | 202475 | × | × | [illegible] |
| Byonic | 975 | 8391 | O14672 | ADAM10 | NISQVLEK | N1(HexNAc(2)Hex(6)) | 439 | 2 | 6 | 0 | 0 | 2:6:0:0 | pauci- or high mannose | 10 | 1 | 1 | 1 | 1 | 0 | 467939 | 37154.7 | 340204 | 241363 | 1488940 | × | 5312.49 | 372589 | × | × | [illegible] |
| Byonic | 979 | 11665 | Q8NHP8 | PLBD2 | RNQSLVQDMDSMVR | N2(HexNAc(2)Hex(6)) | 465 | 2 | 6 | 0 | 0 | 2:6:0:0 | pauci- or high mannose | 10 | 1 | 0 | 0 | 0 | 0 | 292195 | 21973.3 | 156069 | 84585.5 | 289664 | × | 5935.8 | 156642 | × | × | [illegible] |
| Byonic | 986 | 6252 | P08575 | PTPRC | NGSAAMCHFTTK | N1(HexNAc(2)Hex(6)); C7(Carbamidomethyl) | 470 | 2 | 6 | 0 | 0 | 2:6:0:0 | pauci- or high mannose | 10 | 1 | 1 | 1 | 1 | 1 | 308411 | 24058.7 | 198449 | 120604 | 556829 | 20158.5 | 52664.7 | 243985 | × | × | [illegible] |
| Byonic | 1006 | 11444 | Q8TEM1 | NUP210 | AHGGSGNFSWSSSSHLVATVTVK | N7(HexNAc(2)Hex(6)) | 484 | 2 | 6 | 0 | 0 | 2:6:0:0 | pauci- or high mannose | 0 | 1 | 0 | 1 | 1 | 0 | 171154 | 16524.4 | 107187 | 67614.1 | 320244 | × | × | 87789 | × | × | [illegible] |
| Byonic | 1022 | 6243 | P01880 | IGHD | TLLNASR | N4(HexNAc(2)Hex(6)) | 496 | 2 | 6 | 0 | 0 | 2:6:0:0 | pauci- or high mannose | 6 | 1 | 1 | 0 | 0 | 0 | 262335 | 20468.5 | 197278 | 132060 | 668774 | 2830.57 | 4836.01 | 181062 | × | × | [illegible] |
| Byonic | 1024 | 12466 | Q9H3S1 | SEMA4A | ANCSVYESCVDCVLAR | N2(HexNAc(2)Hex(6)); C3(Carbamidomethyl); C9(Carbamidomethyl); C12(Carbamidomethyl) | 496 | 2 | 6 | 0 | 0 | 2:6:0:0 | pauci- or high mannose | 6 | 1 | 1 | 1 | 1 | 0 | 258742 | 20170.5 | 149690 | 83933.8 | 399550 | 3689.59 | 5456.93 | 166438 | × | × | [illegible] |
| Byonic | 1028 | 10893 | P05107 | ITGB2 | KDNNSIICSGLGDCVCGQCLCHTSDVPGK | N3(HexNAc(2)Hex(6)); C8(Carbamidomethyl); C14(Carbamidomethyl); C16(Carbamidomethyl); C19(Carbamidomethyl); C21(Carbamidomethyl) | 501 | 2 | 6 | 0 | 0 | 2:6:0:0 | pauci- or high mannose | 6 | 1 | 1 | 1 | 1 | 1 | 1152810 | 82166 | 695126 | 15512.9 | 2099800 | × | × | 610215 | × | × | [illegible] |
| Byonic | 1033 | 11967 | P05107 | ITGB2 | DNNSIICSGLGDCVCGQCLCHTSDVPGK | N2(HexNAc(2)Hex(6)); C7(Carbamidomethyl); C13(Carbamidomethyl); C15(Carbamidomethyl); C18(Carbamidomethyl); C20(Carbamidomethyl) | 501 | 2 | 6 | 0 | 0 | 2:6:0:0 | pauci- or high mannose | 6 | 1 | 1 | 1 | 0 | 1 | 1777130 | 139275 | 1051690 | 621340 | 3438590 | × | × | 1083080 | × | × | [illegible] |
| Byonic | 1040 | 9399 | Q9Y4L1 | HYOU1 | VFGSQNLTTVK | N6(HexNAc(2)Hex(6)) | 515 | 2 | 6 | 0 | 0 | 2:6:0:0 | pauci- or high mannose | 6 | 1 | 1 | 0 | 0 | 1 | 728629 | 23315.9 | 464915 | 285833 | 1247670 | × | 11681.5 | 474412 | × | × | [illegible] |
| Byonic | 1050 | 7145 | P05556 | ITGB1 | KENSSEICSNNGECVCGQCVCR | N3(HexNAc(2)Hex(6)); C8(Carbamidomethyl); C14(Carbamidomethyl); C16(Carbamidomethyl); C19(Carbamidomethyl); C21(Carbamidomethyl) | 520 | 2 | 6 | 0 | 0 | 2:6:0:0 | pauci- or high mannose | 3 | 1 | 0 | 1 | 1 | 0 | 354446 | 24174.8 | 229004 | 122631 | 546288 | × | 6606.68 | 340371 | × | × | [illegible] |
| Byonic | 1064 | 10306 | Q32P28 | P3H1 | VPLQSAHLYYNVTEK | N11(HexNAc(2)Hex(6)) | 540 | 2 | 6 | 0 | 0 | 2:6:0:0 | pauci- or high mannose | 3 | 1 | 0 | 1 | 1 | 1 | 307136 | 23568.8 | 201058 | 118597 | 504156 | × | 2899.21 | 162483 | × | × | [illegible] |
| Byonic | 1078 | 6793 | P18850 | ATF6 | DHLLLPATTHNK | N11(HexNAc(2)Hex(6)) | 584 | 2 | 6 | 0 | 0 | 2:6:0:0 | pauci- or high mannose | 3 | 1 | 1 | 0 | 0 | 0 | 116739 | 7795.25 | 68492.2 | 34109.4 | 107034 | 715.777 | 5039.92 | 58472.2 | × | × | [illegible] |
| Byonic | 1087 | 6621 | Q9Y4L1 | HYOU1 | ENGTDTVQEEEESPAEGSK | N2(HexNAc(2)Hex(6)) | 596 | 2 | 6 | 0 | 0 | 2:6:0:0 | pauci- or high mannose | 3 | 1 | 1 | 1 | 1 | 1 | 211016 | 28550.6 | 146996 | 113448 | 504787 | 6767.95 | 20151.3 | 179101 | × | × | [illegible] |
| [illegible] | | | | | | | | | | | | | | | | | | | | | | | | | | | | | | |

| | | | | | | | |
|---|---|---|---|---|---|---|---|
| Byonic | 371 | 11872 | P54709 | ATP1B3 | NLTVCPDGALFEQK | N1(HexNAc(6)Hex(6)Fuc(1)); C5(Carbamidomethyl) | [illegible] |
| Byonic | 387 | 8105 | P40259 | CD79B | CNNTSEVYQGCGTELR | C1(Carbamidomethyl); N3(HexNAc(6)Hex(6)Fuc(1)); C11(Carbamidomethyl) | [illegible] |
| Byonic | 413 | 8412 | P25445 | FAS | CKPNFFCNSTVCEHCDPCTK | C1(Carbamidomethyl); C7(Carbamidomethyl); N8(HexNAc(6)Hex(6)Fuc(1)); C12(Carbamidomethyl); C15(Carbamidomethyl); [illegible] | [illegible] |
| Byonic | 587 | 12028 | P01871 | IGHM | GLTFQQNASSMCVPDQDTAIR | N7(HexNAc(6)Hex(6)Fuc(1)); C12(Carbamidomethyl) | [illegible] |
| Byonic | 589 | 5576 | P20138 | CD33 | PQDHGTNLTCQVK | N7(HexNAc(6)Hex(6)Fuc(1)); C10(Carbamidomethyl) | [illegible] |
| Byonic | 612 | 7480 | P05107 | ITGB2 | LTNNSNQFQTEVGK | N3(HexNAc(6)Hex(6)Fuc(1)) | [illegible] |
| Byonic | 659 | 11977 | P08575 | PTPRC | YANITVDYLYNK | N3(HexNAc(6)Hex(6)Fuc(1)) | [illegible] |
| Byonic | 664 | 13812 | P02786 | TFRC | KDFEDLYTPVNGSIVIVR | N11(HexNAc(6)Hex(6)Fuc(1)) | [illegible] |
| Byonic | 665 | 15431 | P02786 | TFRC | DFEDLYTPVNGSIVIVR | N10(HexNAc(6)Hex(6)Fuc(1)) | [illegible] |
| Byonic | 697 | 8433 | P13473 | LAMP2 | VASVININPNTTHSTGSCR | N10(HexNAc(6)Hex(6)Fuc(1)); C18(Carbamidomethyl) | [illegible] |
| Byonic | 708 | 7828 | P11279 | LAMP1 | LLNINPNK | N7(HexNAc(6)Hex(6)Fuc(1)) | [illegible] |
| Byonic | 750 | 4746 | P13473 | LAMP2 | LNSSTIK | N2(HexNAc(6)Hex(6)Fuc(1)) | [illegible] |
| Byonic | 771 | 5477 | P08575 | PTPRC | NASVSISHNSCTAPDK | N1(HexNAc(6)Hex(6)Fuc(1)); C11(Carbamidomethyl) | [illegible] |
| Byonic | 784 | 14165 | Q9BY67 | CADM1 | VDDEMPQHAVLSGPNLFINNLNK | N22(HexNAc(6)Hex(6)Fuc(1)) | [illegible] |
| Byonic | 810 | 3811 | Q9BY67 | CADM1 | TDNGTYR | N3(HexNAc(6)Hex(6)Fuc(1)) | [illegible] |
| Byonic | 811 | 4329 | Q13740-2 | ALCAM | NATGDYK | N1(HexNAc(6)Hex(6)Fuc(1)) | [illegible] |
| Byonic | 829 | 11441 | P08575 | PTPRC | NIETFTCDTQNITYR | C7(Carbamidomethyl); N11(HexNAc(6)Hex(6)Fuc(1)) | [illegible] |
| Byonic | 1049 | 8441 | P08575 | PTPRC | YHLEVEAGNTLVRNESHK | N14(HexNAc(6)Hex(6)Fuc(1)) | [illegible] |
| Glyco-Decip | 1352 | 7748 | Q99523 | SORT | LANNTHQHVFDDLR | | [illegible] |
| Glyco-Decip | 1384 | 8626 | Q9BVK6 | TMED9 | FTFTSHTPGEHQICLHSNSTK | 14,Carbamidomethyl(+57.0215); | [illegible] |
| Glyco-Decip | 1436 | 10769 | P05107 | ITB2 | KDNNSIICSGLGDCVCGQCLCHTSDVPGK | 8,Carbamidomethyl(+57.0215);14,Carbamidomethyl(+57.0215);16,Carbamidomethyl(+57.0215);19,Carbamidomethyl(+57.02[illegible] | [illegible] |
| Glyco-Decip | 1492 | 14735 | Q9BY67 | CADM1 | FQLLNFSSSELK | | [illegible] |
| Glyco-Decip | 1518 | 18368 | Q9UIQ6 | LCAP | LFDDWMASNGTQSLPTDVMTTVFK | | [illegible] |
| pGlyco374 | 1559 | 8246 | P11912 | CD79A | VQEGNESYQQSCGTYLR | 12,Carbamidomethyl[C]; | [illegible] |
| [illegible] | [illegible] | [illegible] | [illegible] | [illegible] | [illegible] | [illegible] | [illegible] |

| | | | | | | |
|---|---|---|---|---|---|---|
| MSFragger- | 1669 | 13474 | O43852-10 | CALU | NATYGYVLDDPDPDDGFNYK | 1N(1647.6132) |
| MSFragger- | 1746 | 17116 | P04844 | RPN2 | SNLDPSNVDSLFYAAQASQALSGCEISISNETK | 24C(57.0214), 30N(1444.5339) |
| MSFragger- | 1747 | 17118 | P04844 | RPN2 | SNLDPSNVDSLFYAAQASQALSGCEISISNETK | 24C(57.0214), 30N(1647.6132) |
| Byonic | 34 | 4683 | P01871 | IGHM | YKNNSDISSTR | N3(HexNAc(3)Hex(5)Fuc(1)NeuAc(1)) |
| Byonic | 51 | 4490 | P01871 | IGHM | NNSDISSTR | N1(HexNAc(3)Hex(5)Fuc(1)NeuAc(1)) |
| Byonic | 108 | 9464 | Q16563 | SYPL1 | GQTEIQVNCPPAVTENK | C9(Carbamidomethyl); N16(HexNAc(3)Hex(5)Fuc(1)NeuAc(1)) |
| Byonic | 169 | 7685 | P20645 | M6PR | EAGNHTSGAGLVQINK | N4(HexNAc(3)Hex(5)Fuc(1)NeuAc(1)) |
| Byonic | 220 | 14332 | P40259 | CD79B | MEESQNESLATLTIQGIR | N6(HexNAc(3)Hex(5)Fuc(1)NeuAc(1)) |
| Byonic | 243 | 10326 | P28907 | CD38 | HPCNITEEDYQPLMK | C3(Carbamidomethyl); N4(HexNAc(3)Hex(5)Fuc(1)NeuAc(1)) |
| Byonic | 373 | 13099 | P54709 | ATP1B3 | NLTVCPDGALFEQK | N1(HexNAc(3)Hex(5)Fuc(1)NeuAc(1)); C5(Carbamidomethyl) |
| Byonic | 377 | 15719 | P15391 | CD19 | WNVSDLGGLGCGLK | N2(HexNAc(3)Hex(5)Fuc(1)NeuAc(1)); C11(Carbamidomethyl) |
| Byonic | 402 | 6742 | P08962 | CD63 | NNHTASILDR | N2(HexNAc(3)Hex(5)Fuc(1)NeuAc(1)) |
| Byonic | 485 | 9854 | Q9HD45 | TM9SF3 | IVDVNLTSEGK | N5(HexNAc(3)Hex(5)Fuc(1)NeuAc(1)) |
| Byonic | 498 | 13384 | P09326 | CD48 | ELQNSVLETTLMPHNYSR | N15(HexNAc(3)Hex(5)Fuc(1)NeuAc(1)) |
| Byonic | 694 | 9309 | P13473 | LAMP2 | VASVININPNTTHSTGSCR | N10(HexNAc(3)Hex(5)Fuc(1)NeuAc(1)); C18(Carbamidomethyl) |
| Byonic | 726 | 5828 | P35613 | BSG | ALMNGSESR | N4(HexNAc(3)Hex(5)Fuc(1)NeuAc(1)) |
| Byonic | 863 | 8995 | P13473 | LAMP2 | VQPFNVTQGK | N5(HexNAc(3)Hex(5)Fuc(1)NeuAc(1)) |
| Byonic | 878 | 17497 | P08195 | SLC3A2 | DASSFLAEWQNITK | N11(HexNAc(3)Hex(5)Fuc(1)NeuAc(1)) |
| Byonic | 890 | 4798 | Q96SB8 | SMC6 | AYNEAEVLYNR | N10(HexNAc(3)Hex(5)Fuc(1)NeuAc(1)) |
| Byonic | 995 | 6899 | P08575 | PTPRC | NGSAAMCHFTTK | N1(HexNAc(3)Hex(5)Fuc(1)NeuAc(1)); C7(Carbamidomethyl) |
| Byonic | 1013 | 14675 | P08575 | PTPRC | SAPPSQVWNMTVSMTSDNSMHVK | N9(HexNAc(3)Hex(5)Fuc(1)NeuAc(1)) |
| Glyco-Decip | 1273 | 5319 | Q9Y6M7 | S4A7 | NLTVSECKK | 7,Carbamidomethyl(+57.0215); |
| Glyco-Decip | 1340 | 7124 | O60637 | TSN3 | NQSVPLSCCR | 8,Carbamidomethyl(+57.0215);9,Carbamidomethyl(+57.0215); |
| MSFragger- | 1621 | 9169 | Q96RQ1 | ERGIC2 | GHAHLAALVNHESYNFSHR | 15N(1856.6557) |
| MSFragger- | 1773 | 17951 | P19397 | CD53 | AAWDSIQSFLQCCGINGTSDWTSGPPASCPSDR | 12C(57.0214), 13C(57.0214), 29C(57.0214), 16N(1856.6557) |
| GRable | 1831 | 6096 | P08575 | PTPRC | NASVSISHNSCTAPDK | 0.2000000000000000.0 |
| Byonic | 159 | 11329 | P11279 | LAMP1 | SSCGKENTSDPSLVIAFGR | C3(Carbamidomethyl); N7(HexNAc(5)Hex(4)Fuc(1)) |
| Byonic | 253 | 6577 | P09326 | CD48 | VQKEDNSTYIMR | N6(HexNAc(5)Hex(4)Fuc(1)) |
| Byonic | 275 | 7357 | P09326 | CD48 | EDNSTYIMR | N3(HexNAc(5)Hex(4)Fuc(1)) |
| Byonic | 290 | 9941 | P25445 | FAS | LCDEGHGLEVEINCTR | C2(Carbamidomethyl); N13(HexNAc(5)Hex(4)Fuc(1)); C14(Carbamidomethyl) |
| Byonic | 317 | 8807 | Q9BVK6 | TMED9 | FTFTSHTPGEHQICLHSNSTK | C14(Carbamidomethyl); N18(HexNAc(5)Hex(4)Fuc(1)) |
| Byonic | 348 | 13240 | P54709 | ATP1B3 | FLKPYTLEEQKNLTVCPDGALFEQK | N12(HexNAc(5)Hex(4)Fuc(1)); C16(Carbamidomethyl) |
| Byonic | 401 | 6190 | P08962 | CD63 | NNHTASILDR | N2(HexNAc(5)Hex(4)Fuc(1)) |
| Byonic | 432 | 12948 | Q9C0H2 | TTYH3 | VWDTAVGLNHTAEPSLQTLER | N9(HexNAc(5)Hex(4)Fuc(1)) |
| Byonic | 481 | 5710 | Q8WTV0 | SCARB1 | AFMNR | N4(HexNAc(5)Hex(4)Fuc(1)) |
| Byonic | 766 | 5732 | P08575 | PTPRC | NASVSISHNSCTAPDK | N1(HexNAc(5)Hex(4)Fuc(1)); C11(Carbamidomethyl) |
| Byonic | 782 | 4984 | P08473 | MME | EIANATAKPEDR | N4(HexNAc(5)Hex(4)Fuc(1)) |
| Byonic | 828 | 11539 | P08575 | PTPRC | NIETFTCDTQNITYR | C7(Carbamidomethyl); N11(HexNAc(5)Hex(4)Fuc(1)) |
| Byonic | 860 | 8084 | P13473 | LAMP2 | VQPFNVTQGK | N5(HexNAc(5)Hex(4)Fuc(1)) |
| Byonic | 918 | 3863 | P08575 | PTPRC | FTNASK | N3(HexNAc(5)Hex(4)Fuc(1)) |
| Byonic | 988 | 6374 | P08575 | PTPRC | NGSAAMCHFTTK | N1(HexNAc(5)Hex(4)Fuc(1)); C7(Carbamidomethyl) |
| Byonic | 1052 | 8531 | P08575 | PTPRC | YHLEVEAGNTLVRNESHK | N14(HexNAc(5)Hex(4)Fuc(1)) |
| Byonic | 1154 | 10174 | O00754 | MAN2B1 | LNQTEPVAGNYYPVNTR | N2(HexNAc(5)Hex(4)Fuc(1)) |
| Byonic | 1224 | 7352 | Q92673 | SORL1 | AASNFTEIK | N4(HexNAc(5)Hex(4)Fuc(1)) |
| Byonic | 1237 | 6486 | Q92673 | SORL1 | SRNSTVEYTLNK | N3(HexNAc(5)Hex(4)Fuc(1)) |
| Glyco-Decip | 1338 | 7100 | P20645 | MPRD | EAGNHTSGAGLVQINK | |
| MSFragger- | 1610 | 7257 | O15031 | PLXNB2 | SCVAVTSAQPQNMSR | 13M(15.9949), 2C(57.0214), 12N(1809.6661) |
| GRable | 1916 | 11654 | O14773 | SV | FLSSSPHLPPSSYFNASGR | 0.0000000000000020000.0 |
| Byonic | 26 | 4706 | P01871 | IGHM | YKNNSDISSTR | N3(HexNAc(4)Hex(5)Fuc(1)NeuAc(1)) |
| Byonic | 50 | 4525 | P01871 | IGHM | NNSDISSTR | N1(HexNAc(4)Hex(5)Fuc(1)NeuAc(1)) |
| Byonic | 136 | 18253 | P07602 | PSAP | DVVTAAGDMLKDNATEEEILVYLEK | N13(HexNAc(4)Hex(5)Fuc(1)NeuAc(1)) |
| Byonic | 166 | 7644 | P20645 | M6PR | EAGNHTSGAGLVQINK | N4(HexNAc(4)Hex(5)Fuc(1)NeuAc(1)) |
| Byonic | 347 | 4523 | O00481 | BTN3A1 | IHNVTASDSGK | N3(HexNAc(4)Hex(5)Fuc(1)NeuAc(1)) |
| Byonic | 423 | 17672 | P25942 | CD40 | QIATGVSDTICEPCPVGFFSNVSSAFEK | C11(Carbamidomethyl); C14(Carbamidomethyl); N21(HexNAc(4)Hex(5)Fuc(1)N[illegible] |
| Byonic | 553 | 12818 | P01871 | IGHM | GLTFQQNASSMCVPDQDTAIR | N7(HexNAc(4)Hex(5)Fuc(1)NeuAc(1)); C12(Carbamidomethyl) |
| Byonic | 678 | 3668 | P11279 | LAMP1 | KDNTTVTR | N3(HexNAc(4)Hex(5)Fuc(1)NeuAc(1)) |
| Byonic | 733 | 5802 | P35613 | BSG | ALMNGSESR | N4(HexNAc(4)Hex(5)Fuc(1)NeuAc(1)) |
| Byonic | 778 | 6027 | P08575 | PTPRC | NASVSISHNSCTAPDK | N1(HexNAc(4)Hex(5)Fuc(1)NeuAc(1)); C11(Carbamidomethyl) |
| Byonic | 881 | 13864 | O00592 | PODXL | QLVLNLTGNTLCAGGASDEK | N5(HexNAc(4)Hex(5)Fuc(1)NeuAc(1)); C12(Carbamidomethyl) |
| Byonic | 889 | 4796 | Q96SB8 | SMC6 | AYNEAEVLYNR | N10(HexNAc(4)Hex(5)Fuc(1)NeuAc(1)) |
| Byonic | 947 | 12487 | P08575 | PTPRC | SFHNFTLCYIK | N4(HexNAc(4)Hex(5)Fuc(1)NeuAc(1)); C8(Carbamidomethyl) |
| Byonic | 1145 | 11770 | O15031 | PLXNB2 | LSHDANETLPLHLYVK | N6(HexNAc(4)Hex(5)Fuc(1)NeuAc(1)) |
| Glyco-Decip | 1331 | 6890 | P08575 | PTPRC | NGSAAMCHFTTK | 7,Carbamidomethyl(+57.0215); |
| Glyco-Decip | 1475 | 13570 | Q96BZ4 | PLD4 | VFIVPVGNHSNIPFSR | |
| Glyco-Decip | 1490 | 14545 | P07602 | SAP | TNSTFVQALVEHVK | |
| GRable | 1914 | 12312 | O14773 | SV | FLSSSPHLPPSSYFNASGR | 0.0000000000000020000.0 |
| Byonic | 33 | 4616 | P01871 | IGHM | YKNNSDISSTR | N3(HexNAc(3)Hex(6)Fuc(1)NeuAc(1)) |
| Byonic | 61 | 4428 | P01871 | IGHM | NNSDISSTR | N1(HexNAc(3)Hex(6)Fuc(1)NeuAc(1)) |
| Byonic | 84 | 14522 | Q9BY67 | CADM1 | DVTVIEGEVATISCQVNK | C14(Carbamidomethyl); N17(HexNAc(6)Hex(7)Fuc(1)) |
| Byonic | 105 | 9422 | Q16563 | SYPL1 | GQTEIQVNCPPAVTENK | C9(Carbamidomethyl); N16(HexNAc(3)Hex(6)Fuc(1)NeuAc(1)) |
| Byonic | 131 | 12873 | Q08722 | CD47 | DIYTFDGALNK | N10(HexNAc(3)Hex(6)Fuc(1)NeuAc(1)) |
| Byonic | 152 | 3771 | Q15043 | SLC39A14 | GNVTQHVQGHR | N2(HexNAc(6)Hex(7)Fuc(1)) |
| Byonic | 155 | 8831 | Q9BRN9 | TM2D3 | LPADCIDCTTNFSCTYGK | C5(Carbamidomethyl); C8(Carbamidomethyl); N11(HexNAc(6)Hex(7)Fuc(1)); C1[illegible] |
| Byonic | 176 | 7597 | P20645 | M6PR | EAGNHTSGAGLVQINK | N4(HexNAc(3)Hex(6)Fuc(1)NeuAc(1)) |
| Byonic | 178 | 6824 | P20645 | M6PR | EAGNHTSGAGLVQINK | N4(HexNAc(6)Hex(7)Fuc(1)) |
| Byonic | 219 | 14235 | P40259 | CD79B | MEESQNESLATLTIQGIR | N6(HexNAc(3)Hex(6)Fuc(1)NeuAc(1)) |
| Byonic | 235 | 5475 | P28907 | CD38 | HPCNITEEDYQPLMK | C3(Carbamidomethyl); N4(HexNAc(6)Hex(7)Fuc(1)); M14(Oxidation) |
| Byonic | 240 | 10261 | P28907 | CD38 | HPCNITEEDYQPLMK | C3(Carbamidomethyl); N4(HexNAc(3)Hex(6)Fuc(1)NeuAc(1)) |
| Byonic | 336 | 9762 | Q9BY67 | CADM1 | VSLTNVSISDEGR | N5(HexNAc(6)Hex(7)Fuc(1)) |
| Byonic | 392 | 8072 | P40259 | CD79B | CNNTSEVYQGCGTELR | C1(Carbamidomethyl); N3(HexNAc(6)Hex(7)Fuc(1)); C11(Carbamidomethyl) |
| Byonic | 393 | 8950 | P40259 | CD79B | CNNTSEVYQGCGTELR | C1(Carbamidomethyl); N3(HexNAc(3)Hex(6)Fuc(1)NeuAc(1)); C11(Carbamidom[illegible] |
| Byonic | 404 | 6668 | P08962 | CD63 | NNHTASILDR | N2(HexNAc(3)Hex(6)Fuc(1)NeuAc(1)) |
| Byonic | 493 | 7089 | P25942 | CD40 | DLVVQQAGTNK | N10(HexNAc(3)Hex(6)Fuc(1)NeuAc(1)) |
| Byonic | 563 | 12784 | P01871 | IGHM | GLTFQQNASSMCVPDQDTAIR | N7(HexNAc(3)Hex(6)Fuc(1)NeuAc(1)); C12(Carbamidomethyl) |
| Byonic | 590 | 6170 | P20138 | CD33 | PQDHGTNLTCQVK | N7(HexNAc(3)Hex(6)Fuc(1)NeuAc(1)); C10(Carbamidomethyl) |
| Byonic | 655 | 11966 | P08575 | PTPRC | YANITVDYLYNK | N3(HexNAc(6)Hex(7)Fuc(1)) |
| Byonic | 687 | 8388 | P13473 | LAMP2 | VASVININPNTTHSTGSCR | N10(HexNAc(6)Hex(7)Fuc(1)); C18(Carbamidomethyl) |
| Byonic | 707 | 8913 | P11279 | LAMP1 | LLNINPNK | N7(HexNAc(3)Hex(6)Fuc(1)NeuAc(1)) |
| Byonic | 715 | 7800 | P11279 | LAMP1 | LLNINPNK | N7(HexNAc(6)Hex(7)Fuc(1)) |
| Byonic | 768 | 5455 | P08575 | PTPRC | NASVSISHNSCTAPDK | N1(HexNAc(6)Hex(7)Fuc(1)); C11(Carbamidomethyl) |
| Byonic | 823 | 11271 | P08575 | PTPRC | NIETFTCDTQNITYR | C7(Carbamidomethyl); N11(HexNAc(6)Hex(7)Fuc(1)) |
| Byonic | 871 | 7832 | P13473 | LAMP2 | VQPFNVTQGK | N5(HexNAc(6)Hex(7)Fuc(1)) |
| Byonic | 876 | 17483 | P08195 | SLC3A2 | DASSFLAEWQNITK | N11(HexNAc(3)Hex(6)Fuc(1)NeuAc(1)) |
| Byonic | 900 | 4355 | O00754 | MAN2B1 | ANLTWSVK | N2(HexNAc(6)Hex(7)Fuc(1)) |
| Byonic | 1014 | 14620 | P08575 | PTPRC | SAPPSQVWNMTVSMTSDNSMHVK | N9(HexNAc(3)Hex(6)Fuc(1)NeuAc(1)) |
| Glyco-Decip | 1297 | 5758 | P35613 | BASI | ALMNGSESR | |
| Glyco-Decip | 1391 | 8927 | P13473 | LAMP2 | VQPFNVTQGK | |
| Glyco-Decip | 1403 | 9253 | P13473 | LAMP2 | VASVININPNTTHSTGSCR | 18,Carbamidomethyl(+57.0215); |
| Glyco-Decip | 1468 | 13005 | P54709 | AT1B3 | FLKPYTLEEQKNLTVCPDGALFEQK | 16,Carbamidomethyl(+57.0215); |
| Glyco-Decip | 1484 | 14271 | P04233 | HG2A | YGNMTEDHVMHLLQNADPLK | |
| Glyco-Decip | 1517 | 18276 | Q9UIQ6 | LCAP | LFDDWMASNGTQSLPTDVMTTVFK | |
| GRable | 1847 | 5954 | P08962 | SV | NNHTASILDR | 0.0200000000.0 |
| GRable | 1861 | 14748 | Q9BY67 | CADM1 | FQLLNFSSSELK | 0.000020000000.0 |
| Byonic | 16 | 6283 | Q96IR3 | | GNWTTNEMEVK | N2(HexNAc(3)Hex(5)); M8(Oxidation) |
| Byonic | 42 | 4294 | P01871 | IGHM | YKNNSDISSTR | N3(HexNAc(3)Hex(5)) |
| Byonic | 48 | 4331 | P01871 | IGHM | YKNNSDISSTR | N3(HexNAc(3)Hex(5)Fuc(1)) |
| Byonic | 179 | 7045 | P20645 | M6PR | EAGNHTSGAGLVQINK | N4(HexNAc(3)Hex(5)Fuc(1)) |
| Byonic | 321 | 8764 | Q9BVK6 | TMED9 | FTFTSHTPGEHQICLHSNSTK | C14(Carbamidomethyl); N18(HexNAc(3)Hex(5)Fuc(1)) |
| Byonic | 329 | 14883 | P04233 | CD74 | MATPLLMQALPMGALPQGPMQNATK | M12(Oxidation); N22(HexNAc(3)Hex(5)) |
| Byonic | 376 | 12190 | P54709 | ATP1B3 | NLTVCPDGALFEQK | N1(HexNAc(3)Hex(5)Fuc(1)); C5(Carbamidomethyl) |
| Byonic | 457 | 7373 | P42892 | ECE1 | HLLENSTASVSEAER | N5(HexNAc(3)Hex(5)Fuc(1)) |
| Byonic | 464 | 6522 | O60637 | TSPAN3 | NQSVPLSCCR | N1(HexNAc(3)Hex(5)Fuc(1)); C8(Carbamidomethyl); C9(Carbamidomethyl) |
| Byonic | 562 | 12206 | P01871 | IGHM | GLTFQQNASSMCVPDQDTAIR | N7(HexNAc(3)Hex(5)Fuc(1)); C12(Carbamidomethyl) |
| Byonic | 683 | 8617 | P13473 | LAMP2 | VASVININPNTTHSTGSCR | N10(HexNAc(3)Hex(5)Fuc(1)); C18(Carbamidomethyl) |
| Byonic | 712 | 8108 | P11279 | LAMP1 | LLNINPNK | N7(HexNAc(3)Hex(5)Fuc(1)) |
| Byonic | 776 | 5619 | P08575 | PTPRC | NASVSISHNSCTAPDK | N1(HexNAc(3)Hex(5)Fuc(1)); C11(Carbamidomethyl) |
| Byonic | 824 | 10952 | P10619 | CTSA | MDPPCTNTTAASTYLNNPYVR | M1(Oxidation); C5(Carbamidomethyl); N7(HexNAc(3)Hex(5)) |
| Byonic | 832 | 11543 | P08575 | PTPRC | NIETFTCDTQNITYR | C7(Carbamidomethyl); N11(HexNAc(3)Hex(5)Fuc(1)) |
| Byonic | 865 | 8069 | P13473 | LAMP2 | VQPFNVTQGK | N5(HexNAc(3)Hex(5)Fuc(1)) |
| Byonic | 872 | 8200 | P13473 | LAMP2 | VQPFNVTQGK | N5(HexNAc(3)Hex(5)) |
| Byonic | 1011 | 10550 | P08575 | PTPRC | SAPPSQVWNMTVSMTSDNSMHVK | N9(HexNAc(3)Hex(5)); M10(Oxidation) |
| Byonic | 1121 | 9532 | P05107 | ITGB2 | NCSAACPGLQLSNNPVK | N1(HexNAc(3)Hex(5)); C2(Carbamidomethyl); C6(Carbamidomethyl) |
| Glyco-Decip | 1312 | 6283 | P08575 | PTPRC | NGSAAMCHFTTK | 7,Carbamidomethyl(+57.0215); |
| Glyco-Decip | 1438 | 10883 | P05107 | ITB2 | KDNNSIICSGLGDCVCGQCLCHTSDVPGK | 8,Carbamidomethyl(+57.0215);14,Carbamidomethyl(+57.0215);16,Carbamidom[illegible] |
| MSFragger- | 1594 | 4327 | O75629 | CREG1 | VNETEMDIAK | 6M(15.9949), 2N(1419.5023) |
| MSFragger- | 1597 | 4977 | P53634 | CTSC | DVNCSVMGPQEK | 4C(57.0214), 7M(15.9949), 3N(1419.5023) |
| MSFragger- | 1598 | 5032 | P53634 | CTSC | DVNCSVMGPQEK | 4C(57.0214), 7M(15.9949), 3N(1565.5602) |
| MSFragger- | 1632 | 10377 | P08575-10 | PTPRC | SAPPSQVWNMTVSMTSDNSMHVK | 10M(15.9949), 14M(15.9949), 9N(1565.5602) |
| MSFragger- | 1694 | 15086 | P04233 | CD74 | MATPLLMQALPMGALPQGPMQNATK | 1M(15.9949), 7M(15.9949), 22N(1565.5602) |
| MSFragger- | 1781 | 18192 | P43308 | SSR2 | AGYFNFTSATITYLAQEDGPVVIGSTSAPGQGGILAQR | 5N(1419.5023) |
| GRable | 1851 | 6364 | O60637 | 1 | NQSVPLSCCR | 0.2000000000.0 |
| Glyco-Decip | 1249 | 4069 | P01871 | IGHM | NNSDISSTR | |
| Glyco-Decip | 1256 | 4305 | P01871 | IGHM | YKNNSDISSTR | |
| Glyco-Decip | 1265 | 4863 | P13473 | LAMP2 | LNSSTIK | |
| Glyco-Decip | 1309 | 6230 | P08575 | PTPRC | NGSAAMCHFTTK | 7,Carbamidomethyl(+57.0215); |
| Glyco-Decip | 1314 | 6381 | O60637 | TSN3 | NQSVPLSCCR | 8,Carbamidomethyl(+57.0215);9,Carbamidomethyl(+57.0215); |
| Glyco-Decip | 1324 | 6637 | Q7LGA3 | HS2ST | YHVLHINTTK | |
| Glyco-Decip | 1363 | 8058 | P11279 | LAMP1 | LLNINPNK | |
| Glyco-Decip | 1366 | 8127 | P13473 | LAMP2 | VQPFNVTQGK | |
| Glyco-Decip | 1369 | 8233 | Q96KP6-2 | TNIP3 | NLCSLKTPGK | 3,Carbamidomethyl(+57.0215); |
| Glyco-Decip | 1373 | 8353 | P11912 | CD79A | VQEGNESYQQSCGTYLR | 12,Carbamidomethyl(+57.0215); |
| Glyco-Decip | 1397 | 9066 | Q9HD45 | TM9S3 | IVDVNLTSEGK | |
| Glyco-Decip | 1409 | 9570 | P05107 | ITB2 | NCSAACPGLQLSNNPVK | 2,Carbamidomethyl(+57.0215);6,Carbamidomethyl(+57.0215); |
| Glyco-Decip | 1417 | 9960 | P11279 | LAMP1 | GHTLTLNFTR | |
| Glyco-Decip | 1457 | 12439 | Q9H3S1 | SEM4A | ANCSVYESCVDCVLAR | 3,Carbamidomethyl(+57.0215);9,Carbamidomethyl(+57.0215);12,Carbamidom[illegible] |
| GRable | 1858 | 13848 | O00754 | SV | TPLVQEVHQNFSAWCSQVVR | 0.00000000020000000000.0 |
| Byonic | 17 | 6350 | Q96IR3 | | GNWTTNEMEVK | N2(HexNAc(5)Hex(6)Fuc(1)); M8(Oxidation) |
| Byonic | 231 | 4766 | Q8TCT7 | SPPL2B | GNCTFYEK | N2(HexNAc(5)Hex(6)Fuc(1)); C3(Carbamidomethyl) |
| Byonic | 406 | 6180 | P08962 | CD63 | NNHTASILDR | N2(HexNAc(5)Hex(6)Fuc(1)) |
| Byonic | 657 | 12063 | P08575 | PTPRC | YANITVDYLYNK | N3(HexNAc(5)Hex(6)Fuc(1)) |
| Byonic | 688 | 8526 | P13473 | LAMP2 | VASVININPNTTHSTGSCR | N10(HexNAc(5)Hex(6)Fuc(1)); C18(Carbamidomethyl) |
| Byonic | 826 | 11404 | P08575 | PTPRC | NIETFTCDTQNITYR | C7(Carbamidomethyl); N11(HexNAc(5)Hex(6)Fuc(1)) |
| Glyco-Decip | 1257 | 4322 | P01871 | IGHM | YKNNSDISSTR | |
| Glyco-Decip | 1281 | 5481 | P08575 | PTPRC | NASVSISHNSCTAPDK | 11,Carbamidomethyl(+57.0215); |
| Glyco-Decip | 1287 | 5569 | P08575 | PTPRC | NASVSISHNSCTAPDK | 11,Carbamidomethyl(+57.0215); |
| Glyco-Decip | 1311 | 6267 | P08575 | PTPRC | NGSAAMCHFTTK | 7,Carbamidomethyl(+57.0215); |
| Glyco-Decip | 1315 | 6396 | O60637 | TSN3 | NQSVPLSCCR | 8,Carbamidomethyl(+57.0215);9,Carbamidomethyl(+57.0215); |
| Glyco-Decip | 1321 | 6496 | P25942 | TNR5 | DLVVQQAGTNK | |
| Glyco-Decip | 1339 | 7106 | P20645 | MPRD | LKPLFNK | |
| Glyco-Decip | 1348 | 7309 | P42892 | ECE1 | HLLENSTASVSEAER | |
| Glyco-Decip | 1357 | 8002 | P13473 | LAMP2 | VQPFNVTQGK | |
| Glyco-Decip | 1358 | 8017 | P11279 | LAMP1 | LLNINPNK | |
| Glyco-Decip | 1375 | 8423 | P25445 | TNR6 | CKPNFFCNSTVCEHCDPCTK | 1,Carbamidomethyl(+57.0215);7,Carbamidomethyl(+57.0215);12,Carbamidom[illegible] |
| Glyco-Decip | 1381 | 8547 | P13473 | LAMP2 | VASVININPNTTHSTGSCR | 18,Carbamidomethyl(+57.0215); |
| Glyco-Decip | 1419 | 9998 | Q9HD45 | TM9S3 | IVDVNLTSEGK | |
| Glyco-Decip | 1449 | 11975 | Q08722 | CD47 | DIYTFDGALNK | |
| Glyco-Decip | 1472 | 13257 | P54709 | AT1B3 | FLKPYTLEEQKNLTVCPDGALFEQK | 16,Carbamidomethyl(+57.0215); |
| pGlyco709 | 1578 | 12526 | P08575 | PTPRC | SFHNFTLCYIK | 8,Carbamidomethyl[C]; |
| GRable | 1850 | 6350 | P08575 | PTPRC | NGSAAMCHFTTK | 0.200000000000.0 |
| GRable | 1852 | 6509 | O60637 | 1 | NQSVPLSCCR | 0.2000000000.0 |
| GRable | 1857 | 7979 | P13473 | LAMP2 | VQPFNVTQGK | 0.0000200000.0 |
| GRable | 1889 | 8505 | P08575 | PTPRC | YHLEVEAGNTLVRNESHK | 0.0000000000000020000.0 |
| GRable | 1917 | 13403 | P40259 | OX | MHCYMNSASGNVSWLWK | 0.00000000002000000.0 |
| Byonic | 271 | 15340 | P01903 | HLA-DRA | RSNYTPITNVPPEVTVLTNSPVELR | N3(HexNAc(6)Hex(7)Fuc(1)NeuAc(1)) |
| Byonic | 277 | 16425 | P01903 | HLA-DRA | SNYTPITNVPPEVTVLTNSPVELR | N2(HexNAc(6)Hex(7)Fuc(1)NeuAc(1)) |
| Byonic | 344 | 13545 | P54709 | ATP1B3 | FLKPYTLEEQKNLTVCPDGALFEQK | N12(HexNAc(6)Hex(7)Fuc(1)NeuAc(1)); C16(Carbamidomethyl) |
| Byonic | 365 | 16815 | P32942 | ICAM3 | VELAPLPPWQPVGQNFTLR | N15(HexNAc(6)Hex(7)Fuc(1)NeuAc(1)) |
| Byonic | 571 | 12672 | P01871 | IGHM | GLTFQQNASSMCVPDQDTAIR | N7(HexNAc(6)Hex(7)Fuc(1)NeuAc(1)); C12(Carbamidomethyl) |
| Byonic | 693 | 9121 | P13473 | LAMP2 | VASVININPNTTHSTGSCR | N10(HexNAc(6)Hex(7)Fuc(1)NeuAc(1)); C18(Carbamidomethyl) |
| Byonic | 779 | 5880 | P08575 | PTPRC | NASVSISHNSCTAPDK | N1(HexNAc(6)Hex(7)Fuc(1)NeuAc(1)); C11(Carbamidomethyl) |
| Byonic | 827 | 12145 | P08575 | PTPRC | NIETFTCDTQNITYR | C7(Carbamidomethyl); N11(HexNAc(6)Hex(7)Fuc(1)NeuAc(1)) |
| Byonic | 1105 | 11739 | Q8TCJ2 | STT3B | TTLVDNNTWNNSHIALVGK | N10(HexNAc(6)Hex(7)Fuc(1)NeuAc(1)) |
| Glyco-Decip | 1466 | 12836 | P08575 | PTPRC | YANITVDYLYNK | |
| GRable | 1867 | 10468 | P05107 | SV | LTNNSNQFQTEVGK | 0.00200000000000.0 |
| GRable | 1885 | 14816 | Q9BY67 | CADM1 | VDDEMPQHAVLSGPNLFINNLNK | 0.0000000000000000000020.0 |
| GRable | 1911 | 6240 | O15031 | 3 | NCSFQPER | 0.20000000.0 |
| Byonic | 72 | 12372 | P43121 | MCAM | CGLSQSQGNLSHVDWFSVHK | C1(Carbamidomethyl); N9(HexNAc(5)Hex(5)Fuc(3)) |
| Byonic | 160 | 12172 | P11279 | LAMP1 | SSCGKENTSDPSLVIAFGR | C3(Carbamidomethyl); N7(HexNAc(5)Hex(5)Fuc(3)) |
| Byonic | 187 | 8379 | P11117 | ACP2 | YHGFLNTSYHR | N6(HexNAc(5)Hex(5)Fuc(3)) |
| Byonic | 227 | 8479 | Q99523 | SORT1 | LANNTHQHVFDDLR | N3(HexNAc(5)Hex(5)Fuc(3)) |
| Byonic | 257 | 9726 | P20138 | CD33 | NNCSLSIVDAR | N2(HexNAc(5)Hex(5)Fuc(3)); C3(Carbamidomethyl) |
| Byonic | 269 | 15485 | P01903 | HLA-DRA | RSNYTPITNVPPEVTVLTNSPVELR | N3(HexNAc(5)Hex(5)Fuc(3)) |
| Byonic | 414 | 8980 | P25445 | FAS | CKPNFFCNSTVCEHCDPCTK | C1(Carbamidomethyl); C7(Carbamidomethyl); N8(HexNAc(5)Hex(5)Fuc(3)); C12[illegible] |
| Byonic | 639 | 16092 | P54709 | ATP1B3 | LHVGYLQPLVAVQVSFAPNNTGK | N19(HexNAc(5)Hex(5)Fuc(3)) |
| Byonic | 820 | 17166 | Q969P0 | IGSF8 | IGPGEPLELLCNVSGALPPAGR | C11(Carbamidomethyl); N12(HexNAc(5)Hex(5)Fuc(3)) |
| pGlyco63 | 1543 | 4809 | P01871 | IGHM | YKNNSDISSTR | |
| Byonic | 38 | 4409 | P01871 | IGHM | YKNNSDISSTR | N3(HexNAc(3)Hex(4)Fuc(1)) |
| Byonic | 62 | 4188 | P01871 | IGHM | NNSDISSTR | N1(HexNAc(3)Hex(4)Fuc(1)) |
| Byonic | 174 | 7511 | P20645 | M6PR | EAGNHTSGAGLVQINK | N4(HexNAc(5)Hex(6)Fuc(1)NeuAc(1)) |
| Byonic | 198 | 7363 | Q96KA5 | CLPTM1L | TVNVSVPK | N3(HexNAc(3)Hex(4)Fuc(1)) |
| Byonic | 224 | 14065 | P40259 | CD79B | MEESQNESLATLTIQGIR | N6(HexNAc(5)Hex(6)Fuc(1)NeuAc(1)) |
| Byonic | 245 | 10011 | P11279 | LAMP1 | GHTLTLNFTR | N7(HexNAc(4)Hex(5)) |
| Byonic | 316 | 8811 | Q9BVK6 | TMED9 | FTFTSHTPGEHQICLHSNSTK | C14(Carbamidomethyl); N18(HexNAc(3)Hex(4)Fuc(1)) |
| Byonic | 372 | 15523 | P15391 | CD19 | WNVSDLGGLGCGLK | N2(HexNAc(5)Hex(6)Fuc(1)NeuAc(1)); C11(Carbamidomethyl) |
| Byonic | 399 | 6189 | P08962 | CD63 | NNHTASILDR | N2(HexNAc(3)Hex(4)Fuc(1)) |
| Byonic | 473 | 6548 | O60637 | TSPAN3 | NQSVPLSCCR | N1(HexNAc(3)Hex(4)Fuc(1)); C8(Carbamidomethyl); C9(Carbamidomethyl) |
| Byonic | 477 | 6422 | O60637 | TSPAN3 | NQSVPLSCCR | N1(HexNAc(4)Hex(5)); C8(Carbamidomethyl); C9(Carbamidomethyl) |
| Byonic | 555 | 12225 | P01871 | IGHM | GLTFQQNASSMCVPDQDTAIR | N7(HexNAc(3)Hex(4)Fuc(1)); C12(Carbamidomethyl) |
| Byonic | 558 | 12766 | P01871 | IGHM | GLTFQQNASSMCVPDQDTAIR | N7(HexNAc(5)Hex(6)Fuc(1)NeuAc(1)); C12(Carbamidomethyl) |
| Byonic | 754 | 4917 | P13473 | LAMP2 | LNSSTIK | N2(HexNAc(4)Hex(5)) |
| Byonic | 755 | 5375 | P13473 | LAMP2 | LNSSTIK | N2(HexNAc(5)Hex(6)Fuc(1)NeuAc(1)) |
| Byonic | 874 | 8271 | P13473 | LAMP2 | VQPFNVTQGK | N5(HexNAc(4)Hex(5)) |
| Byonic | 919 | 3812 | P08575 | PTPRC | FTNASK | N3(HexNAc(5)Hex(6)) |
| Byonic | 945 | 11672 | P08575 | PTPRC | SFHNFTLCYIK | N4(HexNAc(5)Hex(6)); C8(Carbamidomethyl) |
| Byonic | 976 | 4896 | P78536 | ADAM17 | MFSNCSK | N4(HexNAc(4)Hex(5)); C5(Carbamidomethyl) |
| Byonic | 991 | 6263 | P08575 | PTPRC | NGSAAMCHFTTK | N1(HexNAc(4)Hex(5)); C7(Carbamidomethyl) |
| Byonic | 992 | 7510 | P08575 | PTPRC | NGSAAMCHFTTK | N1(HexNAc(5)Hex(6)); C7(Carbamidomethyl) |
| Byonic | 1129 | 13890 | O00754 | MAN2B1 | TPLVQEVHQNFSAWCSQVVR | N10(HexNAc(4)Hex(5)); C15(Carbamidomethyl) |
| Byonic | 1158 | 12109 | Q9Y5Y6 | ST14 | VINQTTCENLLPQQITPR | N3(HexNAc(4)Hex(5)); C7(Carbamidomethyl) |
| Glyco-Decip | 1264 | 4845 | P01871 | IGHM | YKNNSDISSTR | |
| Glyco-Decip | 1325 | 6642 | P08962 | CD63 | NNHTASILDR | |
| Glyco-Decip | 1395 | 8976 | P25445 | TNR6 | CKPNFFCNSTVCEHCDPCTK | 1,Carbamidomethyl(+57.0215);7,Carbamidomethyl(+57.0215);12,Carbamidom[illegible] |
| Glyco-Decip | 1399 | 9147 | Q9HD45 | TM9S3 | IVDVNLTSEGK | |
| Glyco-Decip | 1413 | 9746 | O00754 | MA2B1 | WWHQQTNATQEVVR | |
| Glyco-Decip | 1444 | 11471 | P08575 | PTPRC | NIETFTCDTQNITYR | 7,Carbamidomethyl(+57.0215); |
| Glyco-Decip | 1510 | 17577 | P04233 | HG2A | MATPLLMQALPMGALPQGPMQNATK | |
| MSFragger- | 1635 | 10550 | P08575-10 | PTPRC | SAPPSQVWNMTVSMTSDNSMHVK | 10M(15.9949), 14M(15.9949), 9N(1403.5074) |
| GRable | 1886 | 14899 | Q9BY67 | CADM1 | VDDEMPQHAVLSGPNLFINNLNK | 0.0000000000000000000020.0 |
| GRable | 1897 | 8667 | P26006 | SV | NITIVTGAPR | 0.2000000000.0 |
| Byonic | 82 | 7052 | P53634 | CTSC | DVNCSVMGPQEK | N3(HexNAc(2)Hex(5)Fuc(1)); C4(Carbamidomethyl) |
| Byonic | 158 | 17117 | P04844 | RPN2 | SNLDPSNVDSLFYAAQASQALSGCEISISNETK | C24(Carbamidomethyl); N30(HexNAc(2)Hex(5)Fuc(1)) |
| Byonic | 255 | 6510 | P09326 | CD48 | VQKEDNSTYIMR | N6(HexNAc(5)Hex(5)) |
| Byonic | 410 | 6117 | P08962 | CD63 | NNHTASILDR | N2(HexNAc(2)Hex(5)Fuc(1)) |
| Byonic | 450 | 7662 | O75629 | CREG1 | VNETEMDIAK | N2(HexNAc(2)Hex(5)Fuc(1)) |
| Byonic | 476 | 6471 | O60637 | TSPAN3 | NQSVPLSCCR | N1(HexNAc(2)Hex(5)Fuc(1)); C8(Carbamidomethyl); C9(Carbamidomethyl) |
| Byonic | 556 | 9640 | P01871 | IGHM | GLTFQQNASSMCVPDQDTAIR | N7(HexNAc(3)Hex(4)); M11(Oxidation); C12(Carbamidomethyl) |
| Byonic | 597 | 11188 | P09326 | CD48 | NGTVCLSPPCTLAR | N1(HexNAc(5)Hex(5)); C5(Carbamidomethyl); C10(Carbamidomethyl) |
| Byonic | 601 | 11281 | P09326 | CD48 | NGTVCLSPPCTLAR | N1(HexNAc(3)Hex(4)); C5(Carbamidomethyl); C10(Carbamidomethyl) |

| | | | | | | | |
|---|---|---|---|---|---|---|---|
| Byonic | 724 | 8643 | P26006 | ITGA3 | NITIVTGAPR | N1(HexNAc(5)Hex(5)) | [illegible] |
| Byonic | 806 | 13010 | Q8NC56 | LEMD2 | CIPVMEAQEYIANVTSSSSAK | C1(Carbamidomethyl); M5(Oxidation); N13(HexNAc(3)Hex(4)) | [illegible] |
| Byonic | 833 | 10987 | P10619 | CTSA | MDPPCTNTTAASTYLNNPYVR | M1(Oxidation); C5(Carbamidomethyl); N7(HexNAc(3)Hex(4)) | [illegible] |
| Byonic | 906 | 9204 | Q5JPE7 | NOMO2 | LENITTGTYTIHAQK | N3(HexNAc(2)Hex(5)Fuc(1)) | [illegible] |
| Byonic | 915 | 3768 | P08575 | PTPRC | FTNASK | N3(HexNAc(5)Hex(5)) | [illegible] |
| Byonic | 939 | 11669 | P08575 | PTPRC | SFHNFTLCYIK | N4(HexNAc(5)Hex(5)); C8(Carbamidomethyl) | [illegible] |
| Byonic | 983 | 6330 | P08575 | PTPRC | NGSAAMCHFTTK | N1(HexNAc(5)Hex(5)); C7(Carbamidomethyl) | [illegible] |
| Byonic | 1010 | 10586 | P08575 | PTPRC | SAPPSQVWNMTVSMTSDNSMHVK | N9(HexNAc(3)Hex(4)); M14(Oxidation) | [illegible] |
| Byonic | 1044 | 7257 | O15031 | PLXNB2 | SCVAVTSAQPQNMSR | C2(Carbamidomethyl); N12(HexNAc(5)Hex(5)) | [illegible] |
| Glyco-Decip | 1250 | 4072 | P01871 | IGHM | NNSDISSTR | | [illegible] |
| Glyco-Decip | 1255 | 4256 | P01871 | IGHM | YKNNSDISSTR | | [illegible] |
| Glyco-Decip | 1267 | 4959 | A8K979 | ERI2 | NLSISTK | | [illegible] |
| Glyco-Decip | 1285 | 5541 | P08575 | PTPRC | NASVSISHNSCTAPDK | 11,Carbamidomethyl(+57.0215); | [illegible] |
| Glyco-Decip | 1308 | 6228 | P08575 | PTPRC | NGSAAMCHFTTK | 7,Carbamidomethyl(+57.0215); | [illegible] |
| Glyco-Decip | 1316 | 6409 | O60637 | TSN3 | NQSVPLSCCR | 8,Carbamidomethyl(+57.0215);9,Carbamidomethyl(+57.0215); | [illegible] |
| Glyco-Decip | 1318 | 6428 | P25942 | TNR5 | DLVVQQAGTNK | | [illegible] |
| Glyco-Decip | 1362 | 8057 | P13473 | LAMP2 | VQPFNVTQGK | | [illegible] |
| Glyco-Decip | 1376 | 8438 | Q9NR96 | TLR9 | YNNLTVVPR | | [illegible] |
| Glyco-Decip | 1418 | 9961 | P11279 | LAMP1 | GHTLTLNFTR | | [illegible] |
| Glyco-Decip | 1424 | 10241 | P09326 | CD48 | NGTVCLSPPCTLAR | 5,Carbamidomethyl(+57.0215);10,Carbamidomethyl(+57.0215); | [illegible] |
| Glyco-Decip | 1427 | 10411 | Q96BZ4 | PLD4 | TWPQNFSSHFNR | | [illegible] |
| MSFragger- | 1601 | 5764 | Q6PJF5-2 | RHBDF2 | SNHTGFLHMDCEIK | 11C(57.0214), 9M(15.9949), 2N(1257.4495) | [illegible] |
| MSFragger- | 1641 | 10886 | P10619-2 | CTSA | MDPPCTNTTAASTYLNNPYVR | 1M(15.9949), 5C(57.0214), 7N(1825.6610) | [illegible] |
| MSFragger- | 1689 | 14652 | P07602-2 | PSAP | TNSTFVQALVEHVKEECDR | 17C(57.0214), 2N(1362.4808) | [illegible] |
| MSFragger- | 1708 | 15941 | P04233 | CD74 | MATPLLMQALPMGALPQGPMQNATK | [illegible] | [illegible] |
| GRable | 1835 | 4795 | P13473 | LAMP2 | LNSSTIK | [illegible] | [illegible] |
| Byonic | 191 | 18681 | P13473 | LAMP2 | IAVQFGPGFSWIANFTK | [illegible] | [illegible] |
| Byonic | 258 | 9743 | P20138 | CD33 | NNCSLSIVDAR | [illegible] | [illegible] |
| Byonic | 287 | 5351 | Q9UHD9 | UBQLN2 | SQNRPQGQSTQPSNAAGTNTTSASTPR | N19(HexNAc(1)) | [illegible] |
| Byonic | 354 | 17970 | O14656 | TOR1A | IIAENIYEGGLNSDYVHLFVATLHFPHASNITLYK | N30(HexNAc(6)Hex(3)) | [illegible] |
| Byonic | 394 | 14324 | P01137 | TGFB1 | QSTHSIYMFFNTSELR | N11(HexNAc(6)Hex(3)) | [illegible] |
| Byonic | 585 | 10170 | P01871 | IGHM | GLTFQQNASSMCVPDQDTAIR | [illegible] | [illegible] |
| Byonic | 756 | 5529 | P13473 | LAMP2 | LNSSTIK | [illegible] | [illegible] |
| Byonic | 911 | 6292 | Q7Z417 | NUFIP2 | TIQNSSVSPTSSSSSSSSTGETQTQSSSR | N4(HexNAc(1)) | [illegible] |
| Byonic | 917 | 3975 | P08575 | PTPRC | FTNASK | [illegible] | [illegible] |
| Byonic | 940 | 12561 | P08575 | PTPRC | SFHNFTLCYIK | [illegible] | [illegible] |
| Byonic | 985 | 6923 | P08575 | PTPRC | NGSAAMCHFTTK | [illegible] | [illegible] |
| Byonic | 1015 | 12821 | P08575 | PTPRC | SAPPSQVWNMTVSMTSDNSMHVK | [illegible] | [illegible] |
| Byonic | 1045 | 7845 | O15031 | PLXNB2 | SCVAVTSAQPQNMSR | [illegible] | [illegible] |
| Glyco-Decip | 1252 | 4140 | P01871 | IGHM | NNSDISSTR | | [illegible] |
| Glyco-Decip | 1258 | 4342 | P01871 | IGHM | YKNNSDISSTR | | [illegible] |
| Glyco-Decip | 1288 | 5574 | P08575 | PTPRC | NASVSISHNSCTAPDK | [illegible] | [illegible] |
| Glyco-Decip | 1303 | 6015 | P08962 | CD63 | NNHTASILDR | | [illegible] |
| Glyco-Decip | 1319 | 6435 | P25942 | TNR5 | DLVVQQAGTNK | | [illegible] |
| Glyco-Decip | 1332 | 6998 | P20645 | MPRD | EAGNHTSGAGLVQINK | | [illegible] |
| Glyco-Decip | 1355 | 7971 | P11279 | LAMP1 | LLNINPNK | | [illegible] |
| Glyco-Decip | 1471 | 13179 | P54709 | AT1B3 | FLKPYTLEEQKNLTVCPDGALFEQK | [illegible] | [illegible] |
| Glyco-Decip | 1506 | 16476 | Q969P0 | IGSF8 | IGPGEPLELLCNVSGALPPAGR | [illegible] | [illegible] |
| pGlyco54 | 1542 | 4596 | Q9NRH1 | YAE1 | LCENNAEFNKNCSKSHSGIDCSYVECCR | [illegible] | [illegible] |
| pGlyco206 | 1550 | 6391 | Q9BU61 | NDUFAF3 | GIAVEVQDTPNACATFNFLCHEGR | [illegible] | [illegible] |
| pGlyco232 | 1552 | 6657 | Q15648 | MED1 | TPPSSNSCTASSSSFSSSGSSMSSSQNQHGSSK | [illegible] | [illegible] |
| pGlyco509 | 1564 | 9760 | Q5VXU9 | SHOC1 | MMETLEHLNTYLCHDNLSSNDTK | [illegible] | [illegible] |
| MSFragger- | 1605 | 6418 | Q12874 | SF3A3 | NLPLGWDGKPIPYWLYKLHGLNINYNCEICGNYTYR | [illegible] | [illegible] |
| MSFragger- | 1628 | 9945 | P05556-2 | ITGB1 | VCECNPNYTGSACDCSLDTSTCEASNGQICNGR | [illegible] | [illegible] |
| Byonic | 39 | 4773 | P01871 | IGHM | YKNNSDISSTR | [illegible] | [illegible] |
| Byonic | 60 | 4528 | P01871 | IGHM | NNSDISSTR | [illegible] | [illegible] |
| Byonic | 337 | 16874 | P04233 | CD74 | MATPLLMQALPMGALPQGPMQNATK | [illegible] | [illegible] |
| Byonic | 358 | 16809 | Q8N2K0 | ABHD12 | KPQDQGLNHTCNYYLQPEEDVTIGVWHTVPAVWWK | [illegible] | [illegible] |
| Byonic | 396 | 9753 | O00754 | MAN2B1 | WWHQQTNATQEVVR | [illegible] | [illegible] |
| Byonic | 888 | 4870 | Q96SB8 | SMC6 | AYNEAEVLYNR | [illegible] | [illegible] |
| Byonic | 925 | 4819 | Q8IV08 | PLD3 | DNHTHSDIQVK | [illegible] | [illegible] |
| Byonic | 932 | 12587 | Q96BZ4 | PLD4 | SLQALSNPAANVSVDVK | [illegible] | [illegible] |
| Byonic | 1094 | 14022 | P20273 | CD22 | SATLTCESDANPPVSHYTWFDWNNQSLPYHSQK | [illegible] | [illegible] |
| Byonic | 1241 | 11133 | Q02224 | CENPE | QFEAQNLSMQSVR | [illegible] | [illegible] |
| pGlyco611 | 1568 | 11133 | Q96BZ4 | PLD4 | TWPQNFSSHFNR | | [illegible] |
| MSFragger- | 1616 | 7888 | P05556-2 | ITGB1 | ENSSEICSNNGECVCGQCVCR | [illegible] | [illegible] |
| Byonic | 18 | 14280 | P05556 | ITGB1 | SCGECIQAGPNCGWCTNSTFLQEGMPTSAR | [illegible] | [illegible] |
| Byonic | 67 | 4464 | P01871 | IGHM | NNSDISSTR | [illegible] | [illegible] |
| Byonic | 80 | 6966 | P53634 | CTSC | DVNCSVMGPQEK | [illegible] | [illegible] |
| Byonic | 190 | 18622 | P13473 | LAMP2 | IAVQFGPGFSWIANFTK | [illegible] | [illegible] |
| Byonic | 210 | 10415 | P07602 | PSAP | TCDWLPKPNMSASCK | [illegible] | [illegible] |
| Byonic | 265 | 8954 | Q7Z7H5 | TMED4 | FTFTSHTPGDHQICLHSNSTR | [illegible] | [illegible] |
| Byonic | 313 | 8714 | Q9BVK6 | TMED9 | FTFTSHTPGEHQICLHSNSTK | [illegible] | [illegible] |
| Byonic | 364 | 16798 | P32942 | ICAM3 | VELAPLPPWQPVGQNFTLR | [illegible] | [illegible] |
| Byonic | 389 | 8745 | P40259 | CD79B | CNNTSEVYQGCGTELR | [illegible] | [illegible] |
| Byonic | 391 | 9994 | P40259 | CD79B | CNNTSEVYQGCGTELR | [illegible] | [illegible] |
| Byonic | 398 | 6210 | P08962 | CD63 | NNHTASILDR | [illegible] | [illegible] |
| Byonic | 409 | 6282 | P08962 | CD63 | NNHTASILDR | [illegible] | [illegible] |
| Byonic | 480 | 9983 | Q9HD45 | TM9SF3 | IVDVNLTSEGK | [illegible] | [illegible] |
| Byonic | 495 | 10916 | A2RU67 | FAM234B | NGSAVGVSRPAANLVCLSGMNGSTLWSSLLPEEAR | [illegible] | [illegible] |
| Byonic | 502 | 14898 | Q9UIQ6 | LNPEP | YELSLHPNLTSMTFR | [illegible] | [illegible] |
| Byonic | 564 | 12708 | P01871 | IGHM | GLTFQQNASSMCVPDQDTAIR | [illegible] | [illegible] |
| Byonic | 572 | 12755 | P01871 | IGHM | GLTFQQNASSMCVPDQDTAIR | [illegible] | [illegible] |
| Byonic | 584 | 12768 | P01871 | IGHM | GLTFQQNASSMCVPDQDTAIR | [illegible] | [illegible] |
| Byonic | 667 | 16152 | P02786 | TFRC | DFEDLYTPVNGSIVIVR | [illegible] | [illegible] |
| Byonic | 696 | 9068 | P13473 | LAMP2 | VASVININPNTTHSTGSCR | [illegible] | [illegible] |
| Byonic | 742 | 9542 | P04233 | CD74 | GHHNCSESLELEDPSSGLGVTK | [illegible] | [illegible] |
| Byonic | 753 | 5289 | P13473 | LAMP2 | LNSSTIK | [illegible] | [illegible] |
| Byonic | 777 | 5479 | P08575 | PTPRC | NASVSISHNSCTAPDK | [illegible] | [illegible] |
| Byonic | 780 | 4729 | P08473 | MME | EIANATAKPEDR | [illegible] | [illegible] |
| Byonic | 785 | 14880 | Q9BY67 | CADM1 | VDDEMPQHAVLSGPNLFINNLNK | [illegible] | [illegible] |
| Byonic | 831 | 11403 | P08575 | PTPRC | NIETFTCDTQNITYR | [illegible] | [illegible] |
| Byonic | 866 | 9159 | P13473 | LAMP2 | VQPFNVTQGK | [illegible] | [illegible] |
| Byonic | 870 | 8055 | P13473 | LAMP2 | VQPFNVTQGK | [illegible] | [illegible] |
| Byonic | 981 | 6952 | P08575 | PTPRC | NGSAAMCHFTTK | [illegible] | [illegible] |
| Byonic | 1025 | 13306 | Q9H3S1 | SEMA4A | ANCSVYESCVDCVLAR | [illegible] | [illegible] |
| Byonic | 1173 | 18893 | Q9UIQ6 | LNPEP | LFDDWMASNGTQSLPTDVMTTVFK | [illegible] | [illegible] |
| Glyco-Decip | 1259 | 4618 | P01871 | IGHM | YKNNSDISSTR | | [illegible] |
| Glyco-Decip | 1322 | 6509 | P08581 | MET | NFTVACQHR | [illegible] | [illegible] |
| Glyco-Decip | 1334 | 7011 | P79483 | DRB3 | SECHFFNGTER | [illegible] | [illegible] |
| Glyco-Decip | 1359 | 8036 | P09326 | CD48 | EDNSTYIMR | | [illegible] |
| Glyco-Decip | 1401 | 9175 | Q9BVK6 | TMED9 | FTFTSHTPGEHQICLHSNSTK | [illegible] | [illegible] |
| Glyco-Decip | 1456 | 12402 | P08575 | PTPRC | SFHNFTLCYIK | [illegible] | [illegible] |
| Glyco-Decip | 1463 | 12785 | P01871 | IGHM | GLTFQQNASSMCVPDQDTAIR | [illegible] | [illegible] |
| Glyco-Decip | 1476 | 13643 | P54709 | AT1B3 | FLKPYTLEEQKNLTVCPDGALFEQK | [illegible] | [illegible] |
| Glyco-Decip | 1505 | 15731 | Q16563 | SYPL1 | LNEASFQPPPGVNICDVNWK | [illegible] | [illegible] |
| Glyco-Decip | 1508 | 17155 | Q969P0 | IGSF8 | IGPGEPLELLCNVSGALPPAGR | [illegible] | [illegible] |
| MSFragger- | 1611 | 7285 | O15031 | PLXNB2 | SCVAVTSAQPQNMSRR | [illegible] | [illegible] |
| MSFragger- | 1626 | 9779 | P16422 | EPCAM | QCNGTSMCWCVNTAGVR | [illegible] | [illegible] |
| MSFragger- | 1629 | 10130 | Q03001-14 | DST | VNNSGISLCNLISAVTTPAK | [illegible] | [illegible] |
| MSFragger- | 1643 | 11058 | Q92673 | SORL1 | YHIIVQLGNMSK | [illegible] | [illegible] |
| GRable | 1821 | 12464 | P08575 | PTPRC | SFHNFTLCYIK | [illegible] | [illegible] |
| GRable | 1841 | 4769 | P01871 | IGHM | YKNNSDISSTR | [illegible] | [illegible] |
| GRable | 1843 | 4713 | P01871 | IGHM | YKNNSDISSTR | [illegible] | [illegible] |
| GRable | 1873 | 12313 | P43121 | MCAM | CGLSQSQGNLSHVDWFSVHK | [illegible] | [illegible] |
| GRable | 1904 | 6249 | Q92673 | SV | SRNSTVEYTLNK | [illegible] | [illegible] |
| GRable | 1909 | 5359 | O60637 | 1 | TYNGTNPDAASR | [illegible] | [illegible] |
| GRable | 1912 | 6245 | O15031 | 3 | NCSFQPER | [illegible] | [illegible] |
| Byonic | 41 | 4425 | P01871 | IGHM | YKNNSDISSTR | [illegible] | [illegible] |
| Byonic | 45 | 4482 | P01871 | IGHM | YKNNSDISSTR | [illegible] | [illegible] |
| Byonic | 279 | 7292 | Q7LGA3 | HS2ST1 | YHVLHINTTK | [illegible] | [illegible] |
| Byonic | 281 | 16525 | P01903 | HLA-DRA | SNYTPITNVPPEVTVLTNSPVELR | [illegible] | [illegible] |
| Byonic | 320 | 8768 | Q9BVK6 | TMED9 | FTFTSHTPGEHQICLHSNSTK | [illegible] | [illegible] |
| Byonic | 323 | 8890 | Q9BVK6 | TMED9 | FTFTSHTPGEHQICLHSNSTK | [illegible] | [illegible] |
| Byonic | 328 | 9637 | P20036 | HLA-DPA1 | SNHTQATNDPPEVTVFPK | [illegible] | [illegible] |
| Byonic | 405 | 6181 | P08962 | CD63 | NNHTASILDR | [illegible] | [illegible] |
| Byonic | 411 | 6642 | P08962 | CD63 | NNHTASILDR | [illegible] | [illegible] |
| Byonic | 412 | 8976 | P25445 | FAS | CKPNFFCNSTVCEHCDPCTK | [illegible] | [illegible] |
| Byonic | 475 | 6565 | O60637 | TSPAN3 | NQSVPLSCCR | [illegible] | [illegible] |
| Byonic | 566 | 12740 | P01871 | IGHM | GLTFQQNASSMCVPDQDTAIR | [illegible] | [illegible] |
| Byonic | 574 | 9647 | P01871 | IGHM | GLTFQQNASSMCVPDQDTAIR | [illegible] | [illegible] |
| Byonic | 578 | 9646 | P01871 | IGHM | GLTFQQNASSMCVPDQDTAIR | [illegible] | [illegible] |
| Byonic | 648 | 10566 | P20273 | CD22 | FLSNDTVQLNVK | [illegible] | [illegible] |
| Byonic | 752 | 4750 | P13473 | LAMP2 | LNSSTIK | [illegible] | [illegible] |
| Byonic | 796 | 10026 | P32942 | ICAM3 | EIVCNVTLGGER | [illegible] | [illegible] |
| Byonic | 797 | 9423 | P32942 | ICAM3 | EIVCNVTLGGER | [illegible] | [illegible] |
| Byonic | 912 | 19412 | P08195 | SLC3A2 | LLIAGTNSSDLQQILSLLESNK | [illegible] | [illegible] |
| Byonic | 920 | 3824 | P08575 | PTPRC | FTNASK | [illegible] | [illegible] |
| Byonic | 944 | 11479 | P08575 | PTPRC | SFHNFTLCYIK | [illegible] | [illegible] |
| Byonic | 946 | 12527 | P08575 | PTPRC | SFHNFTLCYIK | [illegible] | [illegible] |
| Byonic | 948 | 12583 | P08575 | PTPRC | SFHNFTLCYIK | [illegible] | [illegible] |
| Byonic | 994 | 6889 | P08575 | PTPRC | NGSAAMCHFTTK | [illegible] | [illegible] |
| Byonic | 1009 | 12912 | P08575 | PTPRC | SAPPSQVWNMTVSMTSDNSMHVK | [illegible] | [illegible] |
| Byonic | 1012 | 11062 | P08575 | PTPRC | SAPPSQVWNMTVSMTSDNSMHVK | [illegible] | [illegible] |
| Byonic | 1193 | 6529 | Q9Y4L1 | HYOU1 | AEPPLNASASDQGEK | [illegible] | [illegible] |
| Byonic | 1236 | 3977 | P42858 | HTT | CENLSTPTMLK | [illegible] | [illegible] |
| Glyco-Decip | 1261 | 4635 | P01871 | IGHM | YKNNSDISSTR | | [illegible] |
| Glyco-Decip | 1284 | 5521 | Q7L1W4 | LRC8D | AHTPPGNAEVTTNIPK | | [illegible] |
| Glyco-Decip | 1291 | 5635 | P08575 | PTPRC | NASVSISHNSCTAPDK | [illegible] | [illegible] |
| Glyco-Decip | 1292 | 5651 | P05556 | ITB1 | CHEGNGTFECGACR | [illegible] | [illegible] |
| Glyco-Decip | 1306 | 6172 | P08962 | CD63 | NNHTASILDR | | [illegible] |
| Glyco-Decip | 1344 | 7221 | P25445 | TNR6 | KTVTTVETQNLEGLHHDGQFCHKPCPPGER | [illegible] | [illegible] |
| Glyco-Decip | 1346 | 7292 | Q7LGA3 | HS2ST | YHVLHINTTK | | [illegible] |
| Glyco-Decip | 1365 | 8115 | P11279 | LAMP1 | LLNINPNK | | [illegible] |
| Glyco-Decip | 1367 | 8145 | Q92896 | GSLG1 | LNLTTDPK | | [illegible] |
| Glyco-Decip | 1379 | 8487 | P13473 | LAMP2 | VASVININPNTTHSTGSCR | [illegible] | [illegible] |
| Glyco-Decip | 1443 | 11467 | P08575 | PTPRC | NIETFTCDTQNITYR | [illegible] | [illegible] |
| Glyco-Decip | 1465 | 12796 | P08575 | PTPRC | YANITVDYLYNK | | [illegible] |
| Glyco-Decip | 1516 | 18111 | P07602 | SAP | DVVTAAGDMLKDNATEEEILVYLEK | | [illegible] |
| Glyco-Decip | 1521 | 18523 | P13473 | LAMP2 | IAVQFGPGFSWIANFTK | | [illegible] |
| pGlyco500 | 1563 | 9669 | P20138 | CD33 | NNCSLSIVDAR | [illegible] | [illegible] |
| pGlyco723 | 1579 | 12800 | P0DOX6 | | GLTFQQNASSMCVPDQDTAIR | [illegible] | [illegible] |
| MSFragger- | 1634 | 10500 | P07602-2 | PSAP | TCDWLPKPNMSASCK | [illegible] | [illegible] |
| MSFragger- | 1638 | 10586 | P08575-10 | PTPRC | SAPPSQVWNMTVSMTSDNSMHVK | [illegible] | [illegible] |
| MSFragger- | 1681 | 14141 | P26006-1 | ITGA3 | AHCVWLECPIPDAPVVTNVTVK | [illegible] | [illegible] |
| MSFragger- | 1718 | 16465 | P01903 | HLA-DRA | SNYTPITNVPPEVTVLTNSPVELR | [illegible] | [illegible] |
| GRable | 1820 | 12485 | P08575 | PTPRC | SFHNFTLCYIK | [illegible] | [illegible] |
| GRable | 1834 | 5369 | P13473 | LAMP2 | LNSSTIK | [illegible] | [illegible] |
| GRable | 1860 | 15648 | Q9BY67 | CADM1 | FQLLNFSSSELK | [illegible] | [illegible] |
| GRable | 1865 | 15386 | P15391 | 1 | WNVSDLGGLGCGLK | [illegible] | [illegible] |
| GRable | 1866 | 10468 | Q9BY67 | CADM1 | VSLTNVSISDEGR | [illegible] | [illegible] |
| GRable | 1868 | 9762 | P05107 | SV | LTNNSNQFQTEVGK | [illegible] | [illegible] |
| GRable | 1872 | 12800 | P01871 | IGHM | GLTFQQNASSMCVPDQDTAIR | [illegible] | [illegible] |
| GRable | 1898 | 8431 | P26006 | SV | NITIVTGAPR | [illegible] | [illegible] |
| Byonic | 102 | 3995 | P13473 | LAMP2 | YETTNK | [illegible] | [illegible] |
| Byonic | 162 | 11314 | P11279 | LAMP1 | SSCGKENTSDPSLVIAFGR | [illegible] | [illegible] |
| Byonic | 209 | 7542 | P07602 | PSAP | TCDWLPKPNMSASCK | [illegible] | [illegible] |
| Byonic | 212 | 10383 | P07602 | PSAP | TCDWLPKPNMSASCK | [illegible] | [illegible] |
| Byonic | 288 | 5310 | Q9UHD9 | UBQLN2 | SQNRPQGQSTQPSNAAGTNTTSASTPR | [illegible] | [illegible] |
| Byonic | 326 | 8690 | Q9BVK6 | TMED9 | FTFTSHTPGEHQICLHSNSTK | [illegible] | [illegible] |
| Byonic | 561 | 9641 | P01871 | IGHM | GLTFQQNASSMCVPDQDTAIR | [illegible] | [illegible] |
| Byonic | 751 | 5329 | P13473 | LAMP2 | LNSSTIK | [illegible] | [illegible] |
| Glyco-Decip | 1245 | 3673 | P08575 | PTPRC | FTNASK | | [illegible] |
| Glyco-Decip | 1247 | 3973 | Q15043 | S39AE | GNVTQHVQGHR | | [illegible] |
| Glyco-Decip | 1389 | 8770 | Q9BVK6 | TMED9 | FTFTSHTPGEHQICLHSNSTK | [illegible] | [illegible] |
| Glyco-Decip | 1400 | 9162 | P32942 | ICAM3 | EIVCNVTLGGER | [illegible] | [illegible] |
| Glyco-Decip | 1421 | 10149 | P32942 | ICAM3 | EIVCNVTLGGER | [illegible] | [illegible] |
| Glyco-Decip | 1431 | 10453 | P20273 | CD22 | FLSNDTVQLNVK | | [illegible] |
| Glyco-Decip | 1447 | 11831 | P08575 | PTPRC | YANITVDYLYNK | | [illegible] |
| Glyco-Decip | 1458 | 12549 | P08575 | PTPRC | SFHNFTLCYIK | [illegible] | [illegible] |
| Glyco-Decip | 1464 | 12791 | P08575 | PTPRC | YANITVDYLYNK | | [illegible] |
| Glyco-Decip | 1502 | 15630 | Q9BY67 | CADM1 | FQLLNFSSSELK | | [illegible] |
| Glyco-Decip | 1522 | 18608 | P13473 | LAMP2 | IAVQFGPGFSWIANFTK | | [illegible] |
| Glyco-Decip | 1533 | 19298 | P08195 | 4F2 | LLIAGTNSSDLQQILSLLESNK | | [illegible] |
| pGlyco142 | 1544 | 5810 | Q68E01 | INTS3 | KNATQPPNAEEESGSSSASEEEDTKPK | | [illegible] |
| pGlyco158 | 1547 | 6090 | Q9Y2G8 | DNAJC16 | TEPSFTKENSSK | | [illegible] |
| pGlyco199 | 1549 | 6348 | Q9UK61 | TASOR | LQQFSPLSDYEGQEEEMNGTKMK | | [illegible] |
| pGlyco451 | 1560 | 9084 | Q8IZU8 | DSEL | KRWTQFVNVTFQMEPTITR | | [illegible] |
| MSFragger- | 1603 | 6233 | Q7Z417 | NUFIP2 | TIQNSSVSPTSSSSSSSSTGETQTQSSSR | [illegible] | [illegible] |
| GRable | 1830 | 5443 | P08575 | PTPRC | NASVSISHNSCTAPDK | [illegible] | [illegible] |
| GRable | 1832 | 4723 | P13473 | LAMP2 | LNSSTIK | [illegible] | [illegible] |
| GRable | 1838 | 5291 | P13473 | LAMP2 | LNSSTIK | [illegible] | [illegible] |
| GRable | 1839 | 4701 | P13473 | LAMP2 | LNSSTIK | [illegible] | [illegible] |
| GRable | 1883 | 7915 | P40259 | OX | CNNTSEVYQGCGTELR | [illegible] | [illegible] |
| GRable | 1887 | 14789 | Q9BY67 | CADM1 | VDDEMPQHAVLSGPNLFINNLNK | [illegible] | [illegible] |
| GRable | 1908 | 17849 | P13473 | LAMP2 | IAVQFGPGFSWIANFTK | [illegible] | [illegible] |
| Byonic | 380 | 5270 | O60637 | TSPAN3 | TYNGTNPDAASR | [illegible] | [illegible] |
| Byonic | 501 | 5847 | Q9NYQ3 | HAO2 | NLTLTDLQSPK | [illegible] | [illegible] |
| Byonic | 692 | 8836 | P13473 | LAMP2 | VASVININPNTTHSTGSCR | [illegible] | [illegible] |
| Byonic | 923 | 4473 | Q8IV08 | PLD3 | DNHTHSDIQVK | [illegible] | [illegible] |
| Glyco-Decip | 1289 | 5587 | Q7L1W4 | LRC8D | AHTPPGNAEVTTNIPK | | [illegible] |
| Glyco-Decip | 1294 | 5683 | Q68CR1 | SE1L3 | NQNESVSEIGGK | | [illegible] |
| Glyco-Decip | 1343 | 7198 | Q7LGA3 | HS2ST | YHVLHINTTK | | [illegible] |
| Glyco-Decip | 1360 | 8042 | Q92896 | GSLG1 | LNLTTDPK | | [illegible] |
| Glyco-Decip | 1377 | 8445 | P13473 | LAMP2 | VASVININPNTTHSTGSCR | [illegible] | [illegible] |
| Glyco-Decip | 1405 | 9319 | Q9BVK6 | TMED9 | FTFTSHTPGEHQICLHSNSTK | [illegible] | [illegible] |
| Glyco-Decip | 1407 | 9515 | P05107 | ITB2 | NCSAACPGLQLSNNPVK | [illegible] | [illegible] |
| Glyco-Decip | 1412 | 9738 | Q9BY67 | CADM1 | VSLTNVSISDEGR | | [illegible] |
| Glyco-Decip | 1435 | 10768 | Q92673 | SORL | AINQTAVECTWTGPR | [illegible] | [illegible] |
| Glyco-Decip | 1440 | 11288 | P08575 | PTPRC | NIETFTCDTQNITYR | [illegible] | [illegible] |
| Glyco-Decip | 1450 | 12039 | P08575 | PTPRC | YANITVDYLYNK | | [illegible] |
| Glyco-Decip | 1453 | 12249 | P11279 | LAMP1 | SSCGKENTSDPSLVIAFGR | [illegible] | [illegible] |
| Glyco-Decip | 1477 | 13669 | Q96BZ4 | PLD4 | VFIVPVGNHSNIPFSR | | [illegible] |
| Glyco-Decip | 1511 | 17677 | P07602 | SAP | DVVTAAGDMLKDNATEEEILVYLEK | | [illegible] |
| Glyco-Decip | 1512 | 17977 | P13473 | LAMP2 | IAVQFGPGFSWIANFTK | | [illegible] |
| Glyco-Decip | 1525 | 18845 | P08195 | 4F2 | LLIAGTNSSDLQQILSLLESNK | | [illegible] |
| MSFragger- | 1595 | 4463 | O75629 | CREG1 | VNETEMDIAK | [illegible] | [illegible] |
| MSFragger- | 1599 | 5194 | P53634 | CTSC | DVNCSVMGPQEK | [illegible] | [illegible] |
| GRable | 1824 | 12524 | P08575 | PTPRC | SFHNFTLCYIK | [illegible] | [illegible] |
| GRable | 1829 | 5525 | P08575 | PTPRC | NASVSISHNSCTAPDK | [illegible] | [illegible] |
| GRable | 1846 | 4575 | P01871 | IGHM | YKNNSDISSTR | [illegible] | [illegible] |
| GRable | 1915 | 12388 | O14773 | SV | FLSSSPHLPPSSYFNASGR | [illegible] | [illegible] |
| Byonic | 44 | 4451 | P01871 | IGHM | YKNNSDISSTR | [illegible] | [illegible] |
| Byonic | 192 | 18636 | P13473 | LAMP2 | IAVQFGPGFSWIANFTK | [illegible] | [illegible] |
| Byonic | 340 | 16960 | P04233 | CD74 | MATPLLMQALPMGALPQGPMQNATK | [illegible] | [illegible] |
| Byonic | 467 | 6578 | O60637 | TSPAN3 | NQSVPLSCCR | [illegible] | [illegible] |
| Byonic | 580 | 10754 | P01871 | IGHM | GLTFQQNASSMCVPDQDTAIR | [illegible] | [illegible] |
| Byonic | 653 | 10575 | P20273 | CD22 | FLSNDTVQLNVK | [illegible] | [illegible] |

| | | | | | | | | | | | | | | |
|---|---|---|---|---|---|---|---|---|---|---|---|---|---|---|
| Byonic | 668 | 16153 | P02786 | TFRC | DFEDLYTPVNGSIVIVR | N10(HexNAc(6)Hex(7)Fuc(2)) | 251 | 6 | 7 | 2 | 0 | 6:7:2:0 | complex | 2 |
| Byonic | 821 | 17189 | Q969P0 | IGSF8 | IGPGEPLELLCNVSGALPPAGR | C11(Carbamidomethyl); N12(HexNAc(5)Hex(6)Fuc(2)) | 327 | 5 | 6 | 2 | 0 | 5:6:2:0 | complex | 6 |
| Byonic | 894 | 6001 | Q13740-2 | ALCAM | NATVVWMK | N1(HexNAc(6)Hex(5)) | 361 | 6 | 5 | 0 | 0 | 6:5:0:0 | complex | 1 |
| Byonic | 943 | 13630 | P08575 | PTPRC | SFHNFTLCYIK | N4(HexNAc(5)Hex(5)NeuAc(2)); C8(Carbamidomethyl) | 421 | 5 | 5 | 0 | 2 | 5:5:0:2 | complex | 1 |
| Glyco-Decip | 1276 | 5387 | Q96GC9 | VMP1 | EHHNGNFTDPSSVNEK | | 23 | 2 | 12 | 0 | 0 | 2:12:0:0 | pauci- or high mannose | 1 |
| Glyco-Decip | 1293 | 5658 | P05556 | ITB1 | CHEGNGTFECGACR | 1,Carbamidomethyl(+57.0215);10,Carbamidomethyl(+57.0215);13,Carbamidomethyl(+57.0215); | 481 | 7 | 7 | 0 | 1 | 7:7:0:1 | complex | 3 |
| Glyco-Decip | 1300 | 5899 | Q9P0L0 | VAPA | PHSVSLNDTETR | | 168 | 2 | 12 | 0 | 0 | 2:12:0:0 | pauci- or high mannose | 3 |
| Glyco-Decip | 1330 | 6803 | P20645 | MPRD | EAGNHTSGAGLVQINK | | 94 | 8 | 7 | 1 | 0 | 8:7:1:0 | complex | 15 |
| Glyco-Decip | 1333 | 7003 | Q9BV40 | VAMP8 | NKTEDLEATSEHFK | | 46 | 2 | 12 | 0 | 0 | 2:12:0:0 | pauci- or high mannose | 15 |
| Glyco-Decip | 1351 | 7659 | P25445 | TNR6 | KTVTTVETQNLEGLHHDGQFCHKPCPPGER | 21,Carbamidomethyl(+57.0215);25,Carbamidomethyl(+57.0215); | 48 | 4 | 4 | 0 | 1 | 4:4:0:1 | complex | 6 |
| Glyco-Decip | 1374 | 8416 | Q8N766 | EMC1 | FINYNQTVSR | | 913 | 8 | 7 | 1 | 0 | 8:7:1:0 | complex | 6 |
| Glyco-Decip | 1460 | 12740 | P01871 | IGHM | GLTFQQNASSMCVPDQDTAIR | 12,Carbamidomethyl(+57.0215); | 209 | 6 | 7 | 2 | 0 | 6:7:2:0 | complex | 3 |
| Glyco-Decip | 1462 | 12760 | P01871 | IGHM | GLTFQQNASSMCVPDQDTAIR | 12,Carbamidomethyl(+57.0215); | 209 | 5 | 6 | 2 | 0 | 5:6:2:0 | complex | 6 |
| Glyco-Decip | 1467 | 12972 | P54709 | AT1B3 | FLKPYTLEEQKNLTVCPDGALFEQK | 16,Carbamidomethyl(+57.0215); | 124 | 8 | 7 | 1 | 0 | 8:7:1:0 | complex | 6 |
| Glyco-Decip | 1520 | 18502 | P13473 | LAMP2 | IAVQFGPGFSWIANFTK | | 101 | 7 | 7 | 0 | 1 | 7:7:0:1 | complex | 0 |
| pGlyco344 | 1555 | 8002 | Q9ULW0 | TPX2 | PFNLSQGK | | 308 | 5 | 4 | 2 | 0 | 5:4:2:0 | complex | 0 |
| MSFragger- | 1738 | 16960 | P04233 | CD74 | MATPLLMQALPMGALPQGPMQNATK | 22N(2407.8519) | 110 | 5 | 5 | 0 | 2 | 5:5:0:2 | complex | 1 |
| MSFragger- | 1765 | 17672 | P25942 | CD40 | QIATGVSDTICEPCPVGFFSNVSSAFEK | 11C(57.0214), 14C(57.0214), 21N(2058.7146) | 133 | 4 | 4 | 0 | 1 | 4:4:0:1 | complex | 0 |
| GRable | 1833 | 5287 | P13473 | LAMP2 | LNSSTIK | 0.0200000.0 | 275 | 7 | 7 | 0 | 1 | 7:7:0:1 | complex | 1 |
| GRable | 1869 | 9706 | P05107 | SV | LTNNSNQFQTEVGK | 0.00200000000000.0 | 212 | 6 | 7 | 2 | 0 | 6:7:2:0 | complex | 0 |
| Byonic | 230 | 13301 | P40259 | CD79B | MEESQNESLATLTIQGIR | M1(Oxidation); N6(HexNAc(6)Hex(7)Fuc(2)NeuAc(1)) | 101 | 6 | 7 | 2 | 1 | 6:7:2:1 | complex | 1 |
| Byonic | 559 | 9676 | P01871 | IGHM | GLTFQQNASSMCVPDQDTAIR | N7(HexNAc(4)Hex(3)); C12(Carbamidomethyl) | 209 | 4 | 3 | 0 | 0 | 4:3:0:0 | complex | 1 |
| Byonic | 605 | 9379 | Q92896 | GLG1 | GNITEYQCHQYITK | N2(HexNAc(5)Hex(6)Fuc(1)NeuAc(2)); C8(Carbamidomethyl) | 210 | 5 | 6 | 1 | 2 | 5:6:1:2 | complex | 1 |
| Byonic | 625 | 12571 | P14625 | HSP90B1 | HNNDTQHIWESDSNEFSVIADPR | N3(HexNAc(6)Hex(4)Fuc(2)) | 217 | 6 | 4 | 2 | 0 | 6:4:2:0 | complex | 1 |
| Byonic | 914 | 19967 | P08195 | SLC3A2 | LLIAGTNSSDLQQILSLLESNK | N7(HexNAc(6)Hex(7)Fuc(2)NeuAc(1)) | 381 | 6 | 7 | 2 | 1 | 6:7:2:1 | complex | 1 |
| Byonic | 1004 | 5413 | P05556 | ITGB1 | CHEGNGTFECGACR | C1(Carbamidomethyl); N5(HexNAc(4)Hex(3)); C10(Carbamidomethyl); C13(Carbamidomethyl) | 481 | 4 | 3 | 0 | 0 | 4:3:0:0 | complex | 10 |
| Byonic | 1160 | 5592 | Q92896 | GLG1 | NDTLQEAK | N1(HexNAc(5)Hex(6)Fuc(1)NeuAc(2)) | 786 | 5 | 6 | 1 | 2 | 5:6:1:2 | complex | 1 |
| Glyco-Decip | 1329 | 6759 | P20645 | MPRD | EAGNHTSGAGLVQINK | | 94 | 7 | 8 | 1 | 0 | 7:8:1:0 | complex | 15 |
| Glyco-Decip | 1402 | 9226 | P32942 | ICAM3 | EIVCNVTLGGER | 4,Carbamidomethyl(+57.0215); | 295 | 6 | 7 | 0 | 0 | 6:7:0:0 | complex | 2 |
| Glyco-Decip | 1406 | 9398 | Q16563 | SYPL1 | GQTEIQVNCPPAVTENK | 9,Carbamidomethyl(+57.0215); | 71 | 3 | 7 | 0 | 1 | 3:7:0:1 | Unclassifiable | 2 |
| Glyco-Decip | 1437 | 10784 | Q9HD45 | TM9S3 | IVDVNLTSEGK | | 174 | 4 | 5 | 0 | 2 | 4:5:0:2 | complex | 1 |
| Glyco-Decip | 1439 | 11092 | P07602 | SAP | TCDWLPKPNMSASCK | 2,Carbamidomethyl(+57.0215);14,Carbamidomethyl(+57.0215); | 101 | 4 | 5 | 3 | 0 | 4:5:3:0 | complex | 1 |
| Glyco-Decip | 1461 | 12742 | P01871 | IGHM | GLTFQQNASSMCVPDQDTAIR | 12,Carbamidomethyl(+57.0215); | 209 | 4 | 7 | 2 | 0 | 4:7:2:0 | hybrid | 6 |
| Glyco-Decip | 1501 | 15615 | P15391 | CD19 | WNVSDLGGLGCGLK | 11,Carbamidomethyl(+57.0215); | 125 | 4 | 5 | 3 | 0 | 4:5:3:0 | complex | 6 |
| Glyco-Decip | 1504 | 15726 | Q16563 | SYPL1 | LNEASFQPPPGVNICDVNWK | 15,Carbamidomethyl(+57.0215); | 96 | 4 | 7 | 2 | 0 | 4:7:2:0 | hybrid | 3 |
| Glyco-Decip | 1523 | 18694 | P08195 | 4F2 | LLIAGTNSSDLQQILSLLESNK | | 381 | 8 | 8 | 0 | 0 | 8:8:0:0 | complex | 0 |
| Glyco-Decip | 1535 | 19396 | P08195 | 4F2 | LLIAGTNSSDLQQILSLLESNK | | 381 | 7 | 8 | 1 | 0 | 7:8:1:0 | complex | 1 |
| pGlyco343 | 1554 | 7996 | Q9ULW0 | TPX2 | PFNLSQGK | | 308 | 6 | 4 | 2 | 0 | 6:4:2:0 | complex | 0 |
| pGlyco354 | 1557 | 8069 | Q9ULW0 | TPX2 | PFNLSQGK | | 308 | 5 | 3 | 2 | 0 | 5:3:2:0 | complex | 0 |
| MSFragger- | 1655 | 12672 | P01871 | IGHM | GLTFQQNASSMCVPDQDTAIR | 12C(57.0214), 7N(2786.9634) | 209 | 4 | 5 | 0 | 4 | 4:5:0:4 | complex | 10 |
| MSFragger- | 1656 | 12737 | P01871 | IGHM | GLTFQQNASSMCVPDQDTAIR | 12C(57.0214), 7N(2277.7888) | 209 | 5 | 4 | 0 | 0 | 5:4:0:0 | complex | 10 |
| MSFragger- | 1697 | 15340 | P01903 | HLA-DRA | RSNYTPITNVPPEVTVLTNSPVELR | 3N(2786.9634) | 103 | 4 | 5 | 0 | 4 | 4:5:0:4 | complex | 0 |
| MSFragger- | 1739 | 16971 | O96005-3 | CLPTM1 | AEDYGPVEVISHWHPNITINIVDDHTPWVK | 16N(2569.9047) | 129 | 5 | 6 | 0 | 2 | 5:6:0:2 | complex | 0 |
| GRable | 1823 | 13458 | P08575 | PTPRC | SFHNFTLCYIK | 0.00020000000.0 | 421 | 5 | 6 | 0 | 2 | 5:6:0:2 | complex | 14 |
| GRable | 1836 | 4687 | P13473 | LAMP2 | LNSSTIK | 0.0200000.0 | 275 | 8 | 8 | 0 | 0 | 8:8:0:0 | complex | 0 |
| GRable | 1837 | 4759 | P13473 | LAMP2 | LNSSTIK | 0.0200000.0 | 275 | 6 | 7 | 0 | 0 | 6:7:0:0 | complex | 1 |
| GRable | 1840 | 4493 | P01871 | IGHM | YKNNSDISSTR | 0.00200000000.0 | 46 | 3 | 7 | 0 | 1 | 3:7:0:1 | Unclassifiable | 0 |
| GRable | 1844 | 4496 | P01871 | IGHM | YKNNSDISSTR | 0.00200000000.0 | 46 | 2 | 7 | 0 | 1 | 2:7:0:1 | pauci- or high mannose | 0 |
| GRable | 1849 | 6777 | P08575 | PTPRC | NGSAAMCHFTTK | 0.200000000000.0 | 470 | 2 | 7 | 0 | 1 | 2:7:0:1 | pauci- or high mannose | 1 |
| GRable | 1892 | 6563 | P09326 | SV | VQKEDNSTYIMR | 0.000002000000.0 | 104 | 5 | 4 | 0 | 0 | 5:4:0:0 | complex | 1 |
| Byonic | 46 | 4200 | P01871 | IGHM | YKNNSDISSTR | N3(HexNAc(2)Hex(11)) | 46 | 2 | 11 | 0 | 0 | 2:11:0:0 | pauci- or high mannose | 1 |
| Byonic | 85 | 6059 | P20645 | M6PR | RLKPLFNK | N7(HexNAc(2)Hex(4)) | 57 | 2 | 4 | 0 | 0 | 2:4:0:0 | pauci- or high mannose | 1 |
| Byonic | 795 | 4878 | P10909 | CLU | HNSTGCLR | N2(HexNAc(5)Hex(6)NeuAc(3)); C6(Carbamidomethyl) | 291 | 5 | 6 | 0 | 3 | 5:6:0:3 | complex | 6 |
| Byonic | 913 | 19409 | P08195 | SLC3A2 | LLIAGTNSSDLQQILSLLESNK | N7(HexNAc(5)Hex(6)Fuc(4)NeuAc(1)) | 381 | 5 | 6 | 4 | 1 | 5:6:4:1 | complex | 1 |
| Byonic | 1076 | 15933 | Q92896 | GLG1 | LCHTHGWNETSEFMPQGAVFSCLYR | C2(Carbamidomethyl); N8(HexNAc(5)Hex(6)Fuc(3)NeuAc(1)); C22(Carbamidomethyl) | 581 | 5 | 6 | 3 | 1 | 5:6:3:1 | complex | 3 |
| Glyco-Decip | 1253 | 4188 | P01871 | IGHM | YKNNSDISSTR | | 46 | 5 | 8 | 0 | 0 | 5:8:0:0 | hybrid | 3 |
| Glyco-Decip | 1260 | 4620 | P01871 | IGHM | YKNNSDISSTR | | 46 | 4 | 7 | 1 | 1 | 4:7:1:1 | hybrid | 3 |
| Glyco-Decip | 1277 | 5392 | P13473 | LAMP2 | LNSSTIK | | 275 | 5 | 7 | 0 | 1 | 5:7:0:1 | hybrid | 1 |
| Glyco-Decip | 1283 | 5506 | P08575 | PTPRC | NASVSISHNSCTAPDK | 11,Carbamidomethyl(+57.0215); | 286 | 4 | 9 | 1 | 0 | 4:9:1:0 | Unclassifiable | 1 |
| Glyco-Decip | 1326 | 6650 | P20645 | MPRD | EAGNHTSGAGLVQINK | | 83 | 9 | 9 | 1 | 0 | 9:9:1:0 | complex | 1 |
| Glyco-Decip | 1364 | 8098 | P13473 | LAMP2 | VQPFNVTQGK | | 356 | 3 | 7 | 0 | 0 | 3:7:0:0 | Unclassifiable | 6 |
| Glyco-Decip | 1380 | 8498 | O94923 | GLCE | VDGFEKRAAASESNNYMNHVAK | | 63 | 4 | 9 | 0 | 0 | 4:9:0:0 | Unclassifiable | 0 |
| Glyco-Decip | 1385 | 8678 | Q9BVK6 | TMED9 | FTFTSHTPGEHQICLHSNSTK | 14,Carbamidomethyl(+57.0215); | 125 | 8 | 4 | 1 | 0 | 8:4:1:0 | complex | 0 |
| Glyco-Decip | 1387 | 8706 | Q9BVK6 | TMED9 | FTFTSHTPGEHQICLHSNSTK | 14,Carbamidomethyl(+57.0215); | 125 | 7 | 4 | 0 | 0 | 7:4:0:0 | complex | 0 |
| Glyco-Decip | 1388 | 8711 | Q9BVK6 | TMED9 | FTFTSHTPGEHQICLHSNSTK | 14,Carbamidomethyl(+57.0215); | 125 | 7 | 3 | 1 | 0 | 7:3:1:0 | complex | 0 |
| Glyco-Decip | 1393 | 8960 | P25445 | TNR6 | CKPNFFCNSTVCEHCDPCTK | 1,Carbamidomethyl(+57.0215);7,Carbamidomethyl(+57.0215);12,Carbamidomethyl(+57.0215);15,Carbamidomethyl(+57.0215);18,Carbamidomethyl(+57.0215); | 136 | 4 | 6 | 3 | 0 | 4:6:3:0 | hybrid | 0 |
| Glyco-Decip | 1398 | 9086 | P13473 | LAMP2 | VASVININPNTTHSTGSCR | 18,Carbamidomethyl(+57.0215); | 257 | 6 | 8 | 0 | 1 | 6:8:0:1 | hybrid | 10 |
| Glyco-Decip | 1408 | 9535 | P05107 | ITB2 | NCSAACPGLQLSNNPVK | 2,Carbamidomethyl(+57.0215);6,Carbamidomethyl(+57.0215); | 642 | 4 | 4 | 0 | 2 | 4:4:0:2 | complex | 2 |
| Glyco-Decip | 1411 | 9706 | Q9BY67 | CADM1 | VSLTNVSISDEGR | | 113 | 7 | 7 | 2 | 0 | 7:7:2:0 | complex | 2 |
| Glyco-Decip | 1416 | 9784 | O00754 | MA2B1 | WWHQQTNATQEVVR | | 133 | 4 | 4 | 1 | 1 | 4:4:1:1 | complex | 2 |
| Glyco-Decip | 1420 | 10103 | P20036 | DPA1 | SNHTQATNDPPEVTVFPK | | 109 | 4 | 5 | 4 | 2 | 4:5:4:2 | complex | 10 |
| Glyco-Decip | 1425 | 10273 | P07602 | SAP | TCDWLPKPNMSASCK | 2,Carbamidomethyl(+57.0215);14,Carbamidomethyl(+57.0215); | 101 | 7 | 5 | 1 | 0 | 7:5:1:0 | complex | 10 |
| Glyco-Decip | 1428 | 10417 | Q9BY67 | CADM1 | VSLTNVSISDEGR | | 113 | 7 | 8 | 2 | 0 | 7:8:2:0 | complex | 10 |
| Glyco-Decip | 1433 | 10535 | Q9BY67 | CADM1 | VSLTNVSISDEGR | | 113 | 6 | 8 | 2 | 0 | 6:8:2:0 | hybrid | 10 |
| Glyco-Decip | 1434 | 10743 | P35613 | BASI | ILLTCSLNDSATEVTGHR | 5,Carbamidomethyl(+57.0215); | 160 | 1 | 10 | 1 | 0 | 1:10:1:0 | Unclassifiable | 10 |
| Glyco-Decip | 1480 | 14089 | P40259 | CD79B | MEESQNESLATLTIQGIR | | 101 | 5 | 7 | 2 | 0 | 5:7:2:0 | hybrid | 3 |
| Glyco-Decip | 1481 | 14104 | P04004 | VTNC | NGSLFAFR | | 169 | 3 | 6 | 0 | 2 | 3:6:0:2 | hybrid | 0 |
| Glyco-Decip | 1482 | 14218 | P04004 | VTNC | NGSLFAFR | | 169 | 3 | 5 | 0 | 2 | 3:5:0:2 | hybrid | 0 |
| Glyco-Decip | 1524 | 18721 | P08195 | 4F2 | LLIAGTNSSDLQQILSLLESNK | | 381 | 8 | 9 | 0 | 0 | 8:9:0:0 | complex | 1 |
| Glyco-Decip | 1527 | 19124 | P08195 | 4F2 | LLIAGTNSSDLQQILSLLESNK | | 381 | 6 | 8 | 3 | 2 | 6:8:3:2 | hybrid | 0 |
| Glyco-Decip | 1529 | 19137 | P08195 | 4F2 | LLIAGTNSSDLQQILSLLESNK | | 381 | 7 | 8 | 4 | 1 | 7:8:4:1 | complex | 1 |
| Glyco-Decip | 1530 | 19152 | P08195 | 4F2 | LLIAGTNSSDLQQILSLLESNK | | 381 | 7 | 9 | 4 | 1 | 7:9:4:1 | hybrid | 0 |
| Glyco-Decip | 1531 | 19204 | P08195 | 4F2 | LLIAGTNSSDLQQILSLLESNK | | 381 | 6 | 7 | 5 | 1 | 6:7:5:1 | complex | 6 |
| Glyco-Decip | 1532 | 19216 | P08195 | 4F2 | LLIAGTNSSDLQQILSLLESNK | | 381 | 7 | 1 | 1 | 0 | 7:1:1:0 | Unclassifiable | 3 |
| Glyco-Decip | 1534 | 19371 | P08195 | 4F2 | LLIAGTNSSDLQQILSLLESNK | | 381 | 5 | 8 | 0 | 1 | 5:8:0:1 | hybrid | 0 |
| Glyco-Decip | 1536 | 19404 | P08195 | 4F2 | LLIAGTNSSDLQQILSLLESNK | | 381 | 6 | 8 | 1 | 0 | 6:8:1:0 | hybrid | 0 |
| Glyco-Decip | 1537 | 19409 | P08195 | 4F2 | LLIAGTNSSDLQQILSLLESNK | | 381 | 5 | 6 | 2 | 2 | 5:6:2:2 | complex | 1 |
| Glyco-Decip | 1538 | 19967 | P08195 | 4F2 | LLIAGTNSSDLQQILSLLESNK | | 381 | 6 | 7 | 0 | 2 | 6:7:0:2 | complex | 0 |
| pGlyco52 | 1541 | 4588 | O43493 | TGOLN2 | SGAEAQTPEDSPNR | | 152 | 2 | 2 | 0 | 0 | 2:2:0:0 | Unclassifiable | 3 |
| pGlyco346 | 1556 | 8004 | Q9ULW0 | TPX2 | PFNLSQGK | | 308 | 6 | 3 | 2 | 0 | 6:3:2:0 | complex | 0 |
| MSFragger- | 1613 | 7632 | A0A1B0GV | PRSS51 | PAFPNSSWLPFHERLQVQNGECPWQVSIQMSR | 22C(57.0214), 30M(15.9949), 5N(552.2167) | 33 | 2 | 0 | 1 | 0 | 2:0:1:0 | Unclassifiable | 1 |
| MSFragger- | 1645 | 11238 | Q99729 | HNRNPAB | GWTGAAAGAGGATAAPPSGNQNGAEGDQINASK | 30N(2157.7831) | 30 | 6 | 4 | 0 | 1 | 6:4:0:1 | complex | 1 |
| GRable | 1825 | 12203 | Q96KA5 | sapiens | DLMVINR | 0.0000020.0 | 229 | 8 | 3 | 4 | 0 | 8:3:4:0 | LacdiNAc | 2 |
| GRable | 1826 | 12184 | Q96KA5 | sapiens | DLMVINR | 0.0000020.0 | 229 | 8 | 4 | 4 | 0 | 8:4:4:0 | complex | 5 |
| GRable | 1827 | 12108 | Q96KA5 | sapiens | DLMVINR | 0.0000020.0 | 229 | 8 | 5 | 4 | 0 | 8:5:4:0 | complex | 0 |
| GRable | 1828 | 12066 | Q96KA5 | sapiens | DLMVINR | 0.0000020.0 | 229 | 8 | 6 | 4 | 0 | 8:6:4:0 | complex | 5 |
| GRable | 1842 | 4511 | P01871 | IGHM | YKNNSDISSTR | 0.00200000000.0 | 46 | 3 | 7 | 1 | 1 | 3:7:1:1 | Unclassifiable | 0 |
| GRable | 1856 | 8774 | Q9BVK6 | GN | FTFTSHTPGEHQICLHSNSTK | 0.00000000000000000002000.0 | 125 | 7 | 4 | 1 | 0 | 7:4:1:0 | complex | 6 |
| GRable | 1863 | 14450 | P08575 | PTPRC | YANITVDYLYNK | 0.002000000000.0 | 234 | 5 | 5 | 4 | 1 | 5:5:4:1 | complex | 0 |
| GRable | 1864 | 15523 | P08575 | PTPRC | YANITVDYLYNK | 0.002000000000.0 | 234 | 3 | 4 | 4 | 2 | 3:4:4:2 | hybrid | 0 |
| GRable | 1871 | 6690 | P20645 | 1 | EAGNHTSGAGLVQINK | 0.0002000000000000.0 | 83 | 8 | 9 | 1 | 0 | 8:9:1:0 | complex | 0 |
