## Supplementary material for "Comparative Evaluation of Glycoproteomics Software for Rare Glycopeptide Identification": Table S2

| Software | No. | Scan | Proteins | Genes | Pep. seq. | pep_var_mod_pos | Glycosylation site | No. of HexNAc | No. of Hex | No. of Fuc | No. of NeuAc | No. of NeuGc | Composition (Hex:HexNAc:Fuc:NeuAc:NeuGc) | glycan type |
|---|---|---|---|---|---|---|---|---|---|---|---|---|---|---|
| Byonic | 21 | 14465 | P11912 | CD79A | VPASLMVSLGEDAHFQCPHNSSNNANVTWWR | C17(Carbamidomethyl); N20(HexNAc(6)Hex(7)Fuc(3)NeuGc(2)) | 57 | 6 | 7 | 3 | 0 | 2 | 6:7:3:0:2 | complex |
| Byonic | 46 | 4635 | P01871 | IGHM | YKNNSDISSTR | N3(HexNAc(4)Hex(5)Fuc(1)NeuGc(1)) | 46 | 4 | 5 | 1 | 0 | 1 | 4:5:1:0:1 | complex |
| Byonic | 47 | 4618 | P01871 | IGHM | YKNNSDISSTR | N3(HexNAc(4)Hex(5)Fuc(2)NeuGc(1)) | 46 | 4 | 5 | 2 | 0 | 1 | 4:5:2:0:1 | complex |
| Byonic | 48 | 4493 | P01871 | IGHM | YKNNSDISSTR | N3(HexNAc(3)Hex(6)Fuc(1)NeuGc(1)) | 46 | 3 | 6 | 1 | 0 | 1 | 3:6:1:0:1 | hybrid |
| Byonic | 51 | 4545 | P01871 | IGHM | YKNNSDISSTR | N3(HexNAc(3)Hex(6)NeuGc(1)) | 46 | 3 | 6 | 0 | 0 | 1 | 3:6:0:0:1 | hybrid |
| Byonic | 83 | 12325 | P43121 | MCAM | CGLSQSQGNLSHVDWFSVHK | C1(Carbamidomethyl); N9(HexNAc(4)Hex(5)Fuc(2)NeuGc(1)) | 56 | 4 | 5 | 2 | 0 | 1 | 4:5:2:0:1 | complex |
| Byonic | 107 | 7118 | P20645 | M6PR | LKPLFNK | N6(HexNAc(3)Hex(5)Fuc(1)NeuGc(2)) | 57 | 3 | 5 | 1 | 0 | 2 | 3:5:1:0:2 | hybrid |
| Byonic | 122 | 9398 | Q16563 | SYPL1 | GQTEIQVNCPPAVTENK | C9(Carbamidomethyl); N16(HexNAc(3)Hex(6)Fuc(1)NeuGc(1)) | 71 | 3 | 6 | 1 | 0 | 1 | 3:6:1:0:1 | hybrid |
| Byonic | 129 | 9508 | Q16563 | SYPL1 | GQTEIQVNCPPAVTENK | C9(Carbamidomethyl); N16(HexNAc(3)Hex(6)NeuGc(1)) | 71 | 3 | 6 | 0 | 0 | 1 | 3:6:0:0:1 | hybrid |
| Byonic | 260 | 13301 | P40259 | CD79B | MEESQNESLATLTIQGIR | N6(HexNAc(6)Hex(7)Fuc(2)NeuGc(1)) | 101 | 6 | 7 | 2 | 0 | 1 | 6:7:2:0:1 | complex |
| Byonic | 286 | 9705 | P20138 | CD33 | NNCSLSIVDAR | N2(HexNAc(4)Hex(5)Fuc(1)NeuGc(1)); C3(Carbamidomethyl) | 100 | 4 | 5 | 1 | 0 | 1 | 4:5:1:0:1 | complex |
| Byonic | 288 | 9765 | P20138 | CD33 | NNCSLSIVDAR | N2(HexNAc(4)Hex(5)Fuc(2)NeuGc(1)); C3(Carbamidomethyl) | 100 | 4 | 5 | 2 | 0 | 1 | 4:5:2:0:1 | complex |
| Byonic | 308 | 15414 | P01903 | HLA-DRA | RSNYTPITNVPPEVTVLTNSPVELR | N3(HexNAc(4)Hex(5)Fuc(2)NeuGc(1)) | 103 | 4 | 5 | 2 | 0 | 1 | 4:5:2:0:1 | complex |
| Byonic | 338 | 7114 | P10321 | HLA-C | GYYNQSEDGSHTLQR | N4(HexNAc(4)Hex(5)Fuc(2)NeuGc(1)) | 110 | 4 | 5 | 2 | 0 | 1 | 4:5:2:0:1 | complex |
| Byonic | 341 | 3845 | P11279 | LAMP1 | NATR | N1(HexNAc(6)Hex(6)NeuAc(1)NeuGc(1)) | 107 | 6 | 6 | 0 | 1 | 1 | 6:6:0:1:1 | complex |
| Byonic | 344 | 3701 | P11279 | LAMP1 | NATR | N1(HexNAc(5)Hex(6)NeuAc(1)NeuGc(1)) | 107 | 5 | 6 | 0 | 1 | 1 | 5:6:0:1:1 | complex |
| Byonic | 381 | 16939 | P04233 | CD74 | MATPLLMQALPMGALPQGPMQNATK | N22(HexNAc(5)Hex(6)Fuc(3)NeuGc(1)) | 130 | 5 | 6 | 3 | 0 | 1 | 5:6:3:0:1 | complex |
| Byonic | 415 | 16856 | P32942 | ICAM3 | VELAPLPPWQPVGQNFTLR | N15(HexNAc(6)Hex(6)NeuAc(1)NeuGc(1)) | 134 | 6 | 6 | 0 | 1 | 1 | 6:6:0:1:1 | complex |
| Byonic | 474 | 5036 | P08962 | CD63 | NNHTASILDR | N2(HexNAc(6)Hex(7)Fuc(3)NeuAc(2)NeuGc(2)) | 130 | 6 | 7 | 3 | 2 | 2 | 6:7:3:2:2 | complex |
| Byonic | 484 | 15907 | P04233 | CD74 | YGNMTEDHVMHLLQNADPLK | N3(HexNAc(3)Hex(6)NeuGc(1)); M4(Oxidation) | 136 | 3 | 6 | 0 | 0 | 1 | 3:6:0:0:1 | hybrid |
| Byonic | 491 | 15990 | P04233 | CD74 | YGNMTEDHVMHLLQNADPLK | N3(HexNAc(3)Hex(4)NeuGc(1)); M4(Oxidation) | 136 | 3 | 4 | 0 | 0 | 1 | 3:4:0:0:1 | hybrid |
| Byonic | 494 | 15915 | P04233 | CD74 | YGNMTEDHVMHLLQNADPLK | N3(HexNAc(3)Hex(5)NeuGc(1)); M4(Oxidation) | 136 | 3 | 5 | 0 | 0 | 1 | 3:5:0:0:1 | hybrid |
| Byonic | 503 | 12257 | O60341 | KDM1A | KLPPPPPQAPPEEENESEPEEPSGVEGAAFQSR | N15(HexNAc(3)Hex(6)NeuGc(1)) | 164 | 3 | 6 | 0 | 0 | 1 | 3:6:0:0:1 | hybrid |
| Byonic | 550 | 14104 | P04004 | VTN | NGSLFAFR | N1(HexNAc(3)Hex(5)Fuc(1)NeuAc(1)NeuGc(1)) | 169 | 3 | 5 | 1 | 1 | 1 | 3:5:1:1:1 | hybrid |
| Byonic | 573 | 6450 | P78536 | ADAM17 | NVSR | N1(HexNAc(6)Hex(7)Fuc(3)NeuGc(1)) | 174 | 6 | 7 | 3 | 0 | 1 | 6:7:3:0:1 | complex |
| Byonic | 631 | 11954 | P01871 | IGHM | VDHRGLTFQQNASSMCVPDQDTAIR | N11(HexNAc(3)Hex(5)Fuc(1)NeuGc(2)); M15(Oxidation); C16(Carbamidomethyl) | 209 | 3 | 5 | 1 | 0 | 2 | 3:5:1:0:2 | hybrid |
| Byonic | 652 | 12800 | P01871 | IGHM | GLTFQQNASSMCVPDQDTAIR | N7(HexNAc(4)Hex(5)Fuc(1)NeuGc(1)); C12(Carbamidomethyl) | 209 | 4 | 5 | 1 | 0 | 1 | 4:5:1:0:1 | complex |
| Byonic | 666 | 10089 | P01871 | IGHM | GLTFQQNASSMCVPDQDTAIR | N7(HexNAc(3)Hex(5)NeuGc(1)); C12(Carbamidomethyl) | 209 | 3 | 5 | 0 | 0 | 1 | 3:5:0:0:1 | hybrid |
| Byonic | 680 | 12743 | P01871 | IGHM | GLTFQQNASSMCVPDQDTAIR | N7(HexNAc(4)Hex(5)Fuc(2)NeuGc(1)); C12(Carbamidomethyl) | 209 | 4 | 5 | 2 | 0 | 1 | 4:5:2:0:1 | complex |
| Byonic | 697 | 9272 | Q92896 | GLG1 | GNITEYQCHQYITK | N2(HexNAc(4)Hex(5)Fuc(3)NeuGc(2)); C8(Carbamidomethyl) | 210 | 4 | 5 | 3 | 0 | 2 | 4:5:3:0:2 | complex |
| Byonic | 717 | 13030 | P14625 | HSP90B1 | HNNDTQHIWESDSNEFSVIADPR | N3(HexNAc(5)Hex(6)NeuGc(2)) | 217 | 5 | 6 | 0 | 0 | 2 | 5:6:0:0:2 | complex |
| Byonic | 753 | 12874 | P08575 | PTPRC | YANITVDYLYNK | N3(HexNAc(6)Hex(7)Fuc(1)NeuGc(1)) | 234 | 6 | 7 | 1 | 0 | 1 | 6:7:1:0:1 | complex |
| Byonic | 847 | 16588 | Q7Z401 | DENND4A | VYGAAIQFYEPYSEENLTEK | N16(HexNAc(5)Hex(5)NeuAc(1)NeuGc(1)) | 283 | 5 | 5 | 0 | 1 | 1 | 5:5:0:1:1 | complex |
| Byonic | 856 | 17290 | P53634 | CTSC | ILTNNSQTPILSPQEVVSCSQYAQGCEGGFPYLIAGK | N4(HexNAc(4)Hex(5)Fuc(2)NeuGc(1)); C19(Carbamidomethyl); C26(Carbamidomethyl) | 276 | 4 | 5 | 2 | 0 | 1 | 4:5:2:0:1 | complex |
| Byonic | 908 | 4080 | P10909 | CLU | HNSTGCLR | N2(HexNAc(4)Hex(5)Fuc(2)NeuGc(1)); C6(Carbamidomethyl) | 291 | 4 | 5 | 2 | 0 | 1 | 4:5:2:0:1 | complex |
| Byonic | 965 | 15077 | Q8TEM1 | NUP210 | LPNSTIYVVEPGYLGFTVHPGDR | N3(HexNAc(5)Hex(6)NeuGc(3)) | 337 | 5 | 6 | 0 | 0 | 3 | 5:6:0:0:3 | complex |
| Byonic | 987 | 9040 | P13473 | LAMP2 | VQPFNVTQGK | N5(HexNAc(3)Hex(6)NeuGc(1)) | 356 | 3 | 6 | 0 | 0 | 1 | 3:6:0:0:1 | hybrid |
| Byonic | 1062 | 6907 | Q6AHZ1 | ZNF518A | AEEGPNASSGFMK | N6(HexNAc(4)Hex(5)Fuc(1)NeuGc(1)) | 409 | 4 | 5 | 1 | 0 | 1 | 4:5:1:0:1 | complex |
| Byonic | 1066 | 8392 | A2RU67 | FAM234B | GQNLTPYWALR | N3(HexNAc(6)Hex(6)NeuAc(1)NeuGc(1)) | 415 | 6 | 6 | 0 | 1 | 1 | 6:6:0:1:1 | complex |
| Byonic | 1075 | 12485 | P08575 | PTPRC | SFHNFTLCYIK | N4(HexNAc(4)Hex(5)Fuc(1)NeuGc(1)); C8(Carbamidomethyl) | 421 | 4 | 5 | 1 | 0 | 1 | 4:5:1:0:1 | complex |
| Byonic | 1077 | 12526 | P08575 | PTPRC | SFHNFTLCYIK | N4(HexNAc(5)Hex(5)NeuGc(1)); C8(Carbamidomethyl) | 421 | 5 | 5 | 0 | 0 | 1 | 5:5:0:0:1 | complex |
| Byonic | 1081 | 12556 | P08575 | PTPRC | SFHNFTLCYIK | N4(HexNAc(5)Hex(6)NeuGc(1)); C8(Carbamidomethyl) | 421 | 5 | 6 | 0 | 0 | 1 | 5:6:0:0:1 | complex |
| Byonic | 1130 | 6816 | P08575 | PTPRC | NGSAAMCHFTTK | N1(HexNAc(3)Hex(6)NeuGc(1)); C7(Carbamidomethyl) | 470 | 3 | 6 | 0 | 0 | 1 | 3:6:0:0:1 | hybrid |
| Byonic | 1211 | 7794 | P11717 | IGF2R | NGSK | N1(HexNAc(6)Hex(7)Fuc(3)NeuAc(3)NeuGc(1)) | 543 | 6 | 7 | 3 | 3 | 1 | 6:7:3:3:1 | complex |
| Byonic | 1212 | 6405 | P11717 | IGF2R | NGSK | N1(HexNAc(6)Hex(7)NeuAc(3)NeuGc(1)) | 543 | 6 | 7 | 0 | 3 | 1 | 6:7:0:3:1 | complex |
| Byonic | 1213 | 9375 | P11717 | IGF2R | NGSK | N1(HexNAc(6)Hex(7)Fuc(3)NeuAc(1)NeuGc(3)) | 543 | 6 | 7 | 3 | 1 | 3 | 6:7:3:1:3 | complex |
| Byonic | 1214 | 10485 | P11717 | IGF2R | NGSK | N1(HexNAc(5)Hex(6)Fuc(2)NeuAc(1)NeuGc(2)) | 543 | 5 | 6 | 2 | 1 | 2 | 5:6:2:1:2 | complex |
| Byonic | 1216 | 4431 | P11717 | IGF2R | NGSK | N1(HexNAc(6)Hex(6)NeuGc(1)) | 543 | 6 | 6 | 0 | 0 | 1 | 6:6:0:0:1 | complex |
| Byonic | 1217 | 5384 | P11717 | IGF2R | NGSK | N1(HexNAc(6)Hex(7)Fuc(2)NeuAc(1)NeuGc(1)) | 543 | 6 | 7 | 2 | 1 | 1 | 6:7:2:1:1 | complex |
| Byonic | 1272 | 9494 | P05107 | ITGB2 | NCSAACPGLQLSNNPVK | N1(HexNAc(5)Hex(6)Fuc(4)NeuGc(1)); C2(Carbamidomethyl); C6(Carbamidomethyl) | 642 | 5 | 6 | 4 | 0 | 1 | 5:6:4:0:1 | complex |
| Byonic | 1307 | 18793 | O00754 | MAN2B1 | PTWKLNQTEPVAGNYYPVNTR | N6(HexNAc(5)Hex(6)Fuc(4)NeuAc(1)NeuGc(2)) | 766 | 5 | 6 | 4 | 1 | 2 | 5:6:4:1:2 | complex |
| Byonic | 1316 | 4976 | Q92896 | GLG1 | NDTLQEAK | N1(HexNAc(4)Hex(5)Fuc(2)NeuGc(1)) | 786 | 4 | 5 | 2 | 0 | 1 | 4:5:2:0:1 | complex |
| Byonic | 1323 | 17847 | Q8TEM1 | NUP210 | FDNFSSLSIQWESTRPVLASIEPELPMQLVSQDDESGQK | N3(HexNAc(5)Hex(6)Fuc(1)NeuGc(1)); M27(Oxidation) | 801 | 5 | 6 | 1 | 0 | 1 | 5:6:1:0:1 | complex |
| Byonic | 1360 | 15832 | Q13428 | TCOF1 | PPLIFVDPNRSPAGPAATPAQAQAASTPR | N9(HexNAc(5)Hex(6)Fuc(1)NeuAc(1)NeuGc(2)) | 965 | 5 | 6 | 1 | 1 | 2 | 5:6:1:1:2 | complex |
| Byonic | 1433 | 14080 | Q96L91 | EP400 | NGTKDLIR | N1(HexNAc(4)Hex(5)Fuc(1)NeuGc(2)) | 2034 | 4 | 5 | 1 | 0 | 2 | 4:5:1:0:2 | complex |
| pGlyco3 | 664 | 9746 | O00754 | MAN2B1 | WWHQQTNATQEVVR | | 133 | 3 | 4 | 1 | 0 | 1 | 3:4:1:0:1 | hybrid |
| MSFragger- | 466 | 9398 | Q16563-2 | SYPL1 | GQTEIQVNCPPAVTENK | 9C(57.0214), 16N(2033.6830) | 43 | 3 | 5 | 0 | 0 | 2 | 3:5:0:0:2 | hybrid |
| MSFragger- | 728 | 11700 | P01871 | IGHM | THTNISESHPNATFSAVGEASICEDDWNSGER | 23C(57.0214), 4N(3727.2653) | 272 | 6 | 7 | 1 | 0 | 4 | 6:7:1:0:4 | complex |
| MSFragger- | 856 | 12737 | P01871 | IGHM | GLTFQQNASSMCVPDQDTAIR | 12C(57.0214), 7N(2277.7888) | 209 | 5 | 4 | 0 | 0 | 2 | 5:4:0:0:2 | complex |
| MSFragger- | 857 | 12740 | P01871 | IGHM | GLTFQQNASSMCVPDQDTAIR | 11M(15.9949), 12C(57.0214), 7N(2626.9262) | 209 | 6 | 5 | 0 | 1 | 1 | 6:5:0:1:1 | complex |
| MSFragger- | 905 | 13281 | Q9H3S1-2 | SEMA4A | ANCSVYESCVDCVLAR | 12C(57.0214), 3C(57.0214), 9C(57.0214), 2N(1726.5927) | 364 | 3 | 5 | 0 | 0 | 1 | 3:5:0:0:1 | hybrid |
| MSFragger- | 995 | 14448 | P11912 | CD79A | VPASLMVSLGEDAHFQCPHNSSNNANVTWWR | 17C(57.0214), 20N(3403.1597) | 37 | 6 | 6 | 0 | 1 | 3 | 6:6:0:1:3 | complex |
| MSFragger- | 1032 | 15041 | P04233 | CD74 | MATPLLMQALPMGALPQGPMQNATK | 12M(15.9949), 1M(15.9949), 7M(15.9949), 22N(2033.6830) | 110 | 3 | 5 | 0 | 0 | 2 | 3:5:0:0:2 | hybrid |
| MSFragger- | 1070 | 15378 | P01903 | HLA-DRA | RSNYTPITNVPPEVTVLTNSPVELR | 3N(2804.9739) | 103 | 6 | 6 | 0 | 0 | 2 | 6:6:0:0:2 | complex |
| MSFragger- | 1321 | 17672 | P25942 | CD40 | QIATGVSDTICEPCPVGFFSNVSSAFEK | 11C(57.0214), 14C(57.0214), 21N(2058.7146) | 133 | 4 | 4 | 0 | 1 | 1 | 4:4:0:1:1 | complex |
| MSFragger- | 1407 | 19209 | Q9GZR5 | ELOVL4 | MGLLDSEPGSVLNVVSTALNDTVEFYR | 1M(15.9949), 20N(1888.6454) | 0 | 3 | 6 | 0 | 0 | 1 | 3:6:0:0:1 | hybrid |
| GRable | 54 | 13458 | P08575 | PTPRC | SFHNFTLCYIK | 0.00020000000.0 | 421 | 5 | 5 | 1 | 1 | 1 | 5:5:1:1:1 | complex |
| GRable | 314 | 5312 | P13473 | LAMP2 | LNSSTIK | 0.0200000.0 | 275 | 6 | 6 | 0 | 0 | 1 | 6:6:0:0:1 | complex |
| GRable | 334 | 4769 | P01871 | IGHM | YKNNSDISSTR | 0.00200000000.0 | 46 | 5 | 5 | 1 | 0 | 1 | 5:5:1:0:1 | complex |
| GRable | 340 | 4713 | P01871 | IGHM | YKNNSDISSTR | 0.00200000000.0 | 46 | 3 | 4 | 1 | 0 | 1 | 3:4:1:0:1 | hybrid |
| GRable | 734 | 15630 | Q9BY67 | CADM1 | FQLLNFSSSELK | 0.000020000000.0 | 101 | 7 | 7 | 1 | 0 | 1 | 7:7:1:0:1 | complex |
| GRable | 870 | 10417 | Q9BY67 | CADM1 | VSLTNVSISDEGR | 0.0000200000000.0 | 113 | 7 | 7 | 1 | 0 | 1 | 7:7:1:0:1 | complex |
| GRable | 952 | 12740 | P01871 | IGHM | GLTFQQNASSMCVPDQDTAIR | 0.000000200000000000000.0 | 209 | 6 | 6 | 1 | 0 | 1 | 6:6:1:0:1 | complex |
| GRable | 1066 | 14789 | Q9BY67 | CADM1 | VDDEMPQHAVLSGPNLFINNLNK | 0.000000000000000000000020.0 | 304 | 7 | 7 | 1 | 0 | 1 | 7:7:1:0:1 | complex |

| Software | No. | Identified by Byonic (0: Not identified, 1: identified) | Identified by MSFragger (0: Not identified, 1: identified) | Identified by pGlyco3 (0: Not identified, 1: identified) | Identified by GRable (0: Not identified, 1: identified)GRable | Intensity of HexNAc signal (138.0555) | Intensity of HexNAc signal (144.0661) | Intensity of HexNAc signal (168.0660) | Intensity of HexNAc signal (186.0766) | Intensity of HexNAc signal (204.0872) | Intensity of NeuAc signal (292.1032) | Intensity of NeuAc-H2O signal (274.0927) | Intensity of NeuGc signal (308.0981) | Intensity of NeuGc-$H_2O$ signal (290.0876) | Intensity of Hex+HexNAc signal (366.14) | Intensity of Hex+HexNAc+Fuc signal (512.1979) | Intensity of Hex2+HexNAc signal (617.2166) | Intensity of Hex+HexNAc+NeuAc signal (657.2354) | Intensity of Hex+HexNAc+NeuGc signal (673.2303) | Intensity of Hex+NeuAc signal (454.156) | Intensity of Hex+NeuGc signal (470.1509) |
|---|---|---|---|---|---|---|---|---|---|---|---|---|---|---|---|---|---|---|---|---|---|
| Byonic | 21 | 1 | 0 | 0 | 0 | 353483 | 20384.8 | 167551 | 5384.9 | 543588 | × | × | × | × | 232467 | × | × | × | × | × | × |
| Byonic | 46 | 1 | 0 | 1 | 1 | 945704 | 2040.51 | 683036 | 283938 | 1250830 | 89569 | 257435 | 38317.9 | 194011 | 905748 | 6859.28 | × | 33337.5 | 27914.9 | 2012.62 | 1369.82 |
| Byonic | 47 | 1 | 0 | 0 | 0 | 456593 | 31091.9 | 333172 | 192464 | 844750 | 82709.9 | 253515 | × | × | 469518 | × | × | 44324.1 | × | × | × |
| Byonic | 48 | 1 | 0 | 0 | 0 | 158083 | 9979.94 | 105114 | 2293.7 | 237928 | 2463.97 | 7826.47 | × | × | 146068 | × | × | × | × | × | × |
| Byonic | 51 | 1 | 0 | 1 | 1 | 473772 | 24653 | 364974 | 182706 | 832759 | 4613.41 | 26882.4 | 53087.6 | 291070 | 469679 | × | × | 3891.44 | 35568.8 | × | × |
| Byonic | 83 | 1 | 0 | 0 | 0 | 1157360 | 72321.8 | 737562 | 351022 | 1263760 | × | 13461.2 | × | × | 923059 | × | × | × | × | × | × |
| Byonic | 107 | 1 | 0 | 0 | 0 | 116692 | 8779.72 | 83534.8 | 51010.1 | 264265 | 1581.01 | 8657.53 | × | × | 102821 | × | × | × | × | × | × |
| Byonic | 122 | 1 | 0 | 0 | 0 | 1926100 | 116774 | 1459890 | 673179 | 3913390 | 463663 | 1353150 | × | 54546.7 | 2395820 | × | × | 248683 | × | 14702.7 | × |
| Byonic | 129 | 1 | 1 | 0 | 1 | 453456 | 26107.4 | 353914 | 179996 | 922237 | 81356.7 | 215834 | 4133.39 | 26407.1 | 556802 | × | × | 51661.7 | 5431.51 | × | × |
| Byonic | 260 | 1 | 0 | 0 | 0 | 282927 | 18579.3 | 188714 | 87456.9 | 331221 | 17600.6 | 43358.9 | × | 3164.09 | 273947 | 2751.82 | × | 8331.34 | × | × | × |
| Byonic | 286 | 1 | 0 | 0 | 0 | 147031 | 10582.6 | 100666 | 62829.4 | 264168 | 19141.1 | 59555.7 | × | × | 141102 | × | × | 8824.92 | × | × | × |
| Byonic | 288 | 1 | 0 | 0 | 0 | 280474 | 18879.5 | 192943 | 85101.8 | 389987 | 32197.1 | 97340.9 | × | × | 325804 | × | × | 19419.9 | × | × | × |
| Byonic | 308 | 1 | 0 | 0 | 0 | 222602 | 12575.5 | 148080 | 71968 | 268268 | 5845.08 | 14758.9 | × | × | 253487 | 1116.69 | × | 1787.16 | × | × | × |
| Byonic | 338 | 1 | 0 | 0 | 0 | 617065 | 36505.2 | 423719 | 200818 | 733535 | × | 3916.31 | × | × | 669751 | × | × | × | × | × | × |
| Byonic | 341 | 1 | 0 | 0 | 0 | 184422 | 10586.3 | 114414 | 52549.2 | 165753 | 2925.26 | 12158.2 | × | × | 174949 | × | × | × | × | × | × |
| Byonic | 344 | 1 | 0 | 0 | 0 | 99700.8 | 5927.63 | 60564.2 | 28669.7 | 75981.3 | 1899.46 | 4442.52 | × | × | 78352.6 | × | × | × | × | × | × |
| Byonic | 381 | 1 | 0 | 0 | 0 | 5011.94 | 6332.5 | 59714.8 | 41521.5 | 242852 | 3487.84 | × | × | × | 118746 | × | × | × | × | × | × |
| Byonic | 415 | 1 | 0 | 0 | 0 | 448128 | 17918.6 | 270481 | 100547 | 306674 | 21568.2 | 38535.5 | × | × | 502714 | 4570.36 | × | 37389.5 | × | × | × |
| Byonic | 474 | 1 | 0 | 0 | 0 | 182944 | 6507.67 | 125420 | 41891.8 | 130070 | × | × | × | × | 331061 | 3705.27 | × | × | × | × | × |
| Byonic | 484 | 1 | 0 | 0 | 0 | 481197 | 28401 | 325859 | 178844 | 934809 | × | 5553.68 | × | × | 317465 | × | × | × | × | × | × |
| Byonic | 491 | 1 | 0 | 0 | 0 | 328571 | 18800.7 | 209661 | 91264.4 | 487420 | 5532.07 | × | × | × | 202295 | × | × | × | × | × | × |
| Byonic | 494 | 1 | 0 | 0 | 0 | 306916 | 19985.9 | 174743 | 97737.2 | 486280 | × | 9627.71 | × | × | 197369 | × | × | 2623.64 | × | × | × |
| Byonic | 503 | 1 | 0 | 0 | 0 | 138015 | 8899.52 | 89909.4 | 44470.2 | 169022 | 5375.04 | 16798.9 | × | 982.639 | 139287 | 971.572 | × | 5731.25 | × | × | × |
| Byonic | 550 | 1 | 0 | 0 | 0 | 252092 | 23169.9 | 8412.62 | 161087 | 1048170 | 290093 | 953931 | × | × | 318680 | × | × | 66756.7 | × | 7917.38 | × |
| Byonic | 573 | 1 | 0 | 0 | 0 | 34941.6 | 12722.9 | 20613.8 | 30873.4 | 43844.8 | × | × | × | × | 15123.6 | × | × | × | × | × | 6223.28 |
| Byonic | 631 | 1 | 0 | 0 | 0 | 476449 | 27548.6 | 313485 | 123017 | 447836 | × | 9582.35 | × | × | 574841 | 4748.51 | × | × | × | × | × |
| Byonic | 652 | 1 | 0 | 0 | 1 | 806410 | 51168.1 | 529264 | 221710 | 1060440 | 81255.9 | 171826 | 18285.9 | 50114.2 | 1017490 | 14351.4 | × | 73449.1 | 24337.7 | × | × |
| Byonic | 666 | 1 | 0 | 0 | 0 | 150158 | 8488.28 | 90790.3 | 47858.8 | 154601 | 7899.56 | 1216.49 | × | × | 96378.5 | × | × | 1360.3 | × | × | × |
| Byonic | 680 | 1 | 0 | 0 | 0 | 790282 | 61810.2 | 522829 | 253831 | 1111610 | × | 15642.4 | × | × | 795786 | 8283.04 | × | × | × | × | × |
| Byonic | 697 | 1 | 0 | 0 | 0 | 419209 | 20400 | 295610 | 117820 | 510072 | 33171.6 | 80013.8 | × | × | 525479 | 1926.09 | 1740.47 | 32778 | × | 1283.5 | × |
| Byonic | 717 | 1 | 0 | 0 | 0 | 240627 | 1434.97 | 149505 | 59864.3 | 208480 | 3547.2 | 8270.6 | × | × | 230735 | 1221.94 | × | 1289.83 | × | × | × |
| Byonic | 753 | 1 | 0 | 0 | 1 | 749481 | 37944.9 | 514024 | 164307 | 598069 | 32246.4 | 66942.8 | 10243.8 | 24811.4 | 991486 | 11641 | × | 52054.2 | 20139.2 | × | × |
| Byonic | 847 | 1 | 0 | 0 | 0 | 186199 | 12401.8 | 4244.85 | 63063.2 | 312876 | 2587.79 | 13426.9 | × | × | 201106 | 2346.64 | × | 2729.87 | × | × | × |
| Byonic | 856 | 1 | 0 | 0 | 0 | 362457 | 22449.1 | 223193 | 102774 | 408486 | 7316.76 | 19582.5 | × | × | 422798 | 7054.04 | 3325.85 | 7521.73 | × | × | × |
| Byonic | 908 | 1 | 0 | 0 | 0 | 121541 | 7184.86 | 80978.8 | 40752.8 | 128590 | 10930.5 | 40195.1 | × | × | 106654 | × | × | 5109.99 | × | × | × |
| Byonic | 965 | 1 | 0 | 0 | 0 | 144445 | 8134.3 | 93340.7 | 39857.6 | 155530 | 4314.03 | 11558.5 | × | 1002.41 | 152588 | 1311.93 | × | 3787.9 | × | × | × |
| Byonic | 987 | 1 | 0 | 1 | 1 | 498208 | 36248.3 | 390196 | 198538 | 920408 | 4489.86 | 38744.6 | 40457.3 | 283356 | 489275 | × | × | × | 25582.1 | × | × |
| Byonic | 1062 | 1 | 0 | 0 | 0 | 558379 | 32939.6 | 402368 | 199601 | 741871 | 67250.9 | 218453 | × | × | 620892 | × | × | 23405.7 | × | 4184.16 | × |
| Byonic | 1066 | 1 | 0 | 0 | 0 | 440234 | 24337.9 | 269564 | 103948 | 366264 | 18069.9 | 44249.1 | × | 2566.45 | 439593 | 3089.28 | × | 12544.1 | × | × | × |
| Byonic | 1075 | 1 | 0 | 0 | 1 | 2657720 | 160159 | 1741270 | 816208 | 2989670 | × | 7892.2 | × | 6554.51 | 2457080 | × | 14103.3 | × | × | × | × |
| Byonic | 1077 | 1 | 0 | 1 | 1 | 807128 | 64143.8 | 597808 | 276943 | 1098360 | × | 12943 | 47617.1 | 246767 | 860628 | × | × | × | 56562.8 | × | × |
| Byonic | 1081 | 1 | 0 | 0 | 1 | 278562 | 16074.7 | 187334 | 83132.4 | 308852 | 3756.12 | 15254 | 12504.2 | 44358.4 | 280839 | × | × | 2770.41 | 12509.6 | × | × |
| Byonic | 1130 | 1 | 1 | 1 | 1 | 557488 | 37979.3 | 430816 | 204225 | 1100120 | × | 6785.68 | 75453.5 | 358541 | 614686 | × | × | × | 46091.7 | × | × |
| Byonic | 1211 | 1 | 0 | 0 | 0 | 123043 | 7411.84 | 3952.87 | 40612.2 | 156696 | 5780.98 | 24003.7 | × | × | 163840 | × | × | 8588.22 | × | × | × |
| Byonic | 1212 | 1 | 0 | 0 | 0 | 223176 | 13628.4 | 142098 | 3405.78 | 265500 | 3346.9 | 14795 | × | × | 218918 | × | × | 3342.77 | × | × | × |
| Byonic | 1213 | 1 | 0 | 0 | 0 | 59002.3 | 4483.16 | 47067.5 | 1443.28 | 94435.5 | 1409.21 | 6565.23 | × | × | 56756.4 | × | × | 1377.2 | × | × | × |
| Byonic | 1214 | 1 | 0 | 0 | 0 | 72876.4 | 5585.85 | 37179.2 | 1067.86 | 58207.4 | 2300.45 | 7565.7 | × | × | 51342.1 | × | × | 3197.55 | × | × | × |
| Byonic | 1216 | 1 | 0 | 0 | 0 | 84477.9 | 9969.13 | 1727.33 | 27060.5 | 102247 | × | 2547.78 | × | × | 47830.6 | × | × | × | × | × | × |
| Byonic | 1217 | 1 | 0 | 0 | 0 | 395564 | 16426.2 | 269911 | 99934.9 | 306055 | 16495.2 | 35099.1 | × | × | 522134 | 3920.86 | × | 24487.9 | × | × | × |
| Byonic | 1272 | 1 | 0 | 0 | 0 | 198453 | 17125.6 | 133021 | 67644.6 | 262282 | 11426.2 | 34268 | × | × | 193097 | × | × | × | × | × | × |
| Byonic | 1307 | 1 | 0 | 0 | 0 | 112809 | 8865.87 | 1972.86 | 45998.3 | 218206 | 52190.1 | 159906 | 1504.13 | 8083.41 | 137878 | × | × | 30428.3 | × | 2936.67 | × |
| Byonic | 1316 | 1 | 0 | 0 | 0 | 392490 | 16197.5 | 276472 | 7165.42 | 516534 | 46392.3 | 147494 | × | × | 419335 | × | × | 31752.8 | × | × | × |
| Byonic | 1323 | 1 | 0 | 0 | 0 | 73694.3 | 6174.08 | 42872 | 991.283 | 91246.3 | 3293.13 | 6026.23 | × | × | 78469.1 | × | × | 4283.54 | × | × | × |
| Byonic | 1360 | 1 | 0 | 0 | 0 | 112398 | 7837.35 | 68009.6 | 33599 | 117993 | 4836.08 | 10589.1 | × | × | 100720 | × | × | 1181.33 | × | × | × |
| Byonic | 1433 | 1 | 0 | 0 | 0 | 76255 | 8687.85 | 45782.9 | 28334 | 99683.2 | 1673.91 | 7039.2 | × | × | 46028.3 | × | × | × | × | × | × |
| pGlyco3 | 664 | 0 | 0 | 1 | 0 | 204033 | 13091.6 | 135158 | 70866.9 | 297673 | 6093.43 | 15462 | 13594 | 2145.72 | 150066 | × | × | 2021.96 | 7683.12 | × | × |
| MSFragger- | 466 | 0 | 1 | 0 | 0 | 1926100 | 116774 | 1459890 | 673179 | 3913390 | 463663 | 1353150 | × | 54546.7 | 2395820 | × | × | 248683 | × | 14702.7 | × |
| MSFragger- | 728 | 0 | 1 | 0 | 0 | 75080 | 4816.7 | 45321.3 | 21018.8 | 84808.6 | 4032.58 | 9415.13 | × | × | 93801.4 | 1018.97 | × | 3374.34 | × | × | × |
| MSFragger- | 856 | 0 | 1 | 0 | 0 | 396116 | 856.514 | 246899 | 117112 | 509554 | 4879.59 | 12741.1 | 11871.8 | 34271.7 | 454477 | 5765.12 | 2315.67 | 5712.67 | 17533 | × | × |
| MSFragger- | 857 | 0 | 1 | 0 | 0 | 892813 | 38711.6 | 555468 | 222027 | 976503 | 70511.5 | 148176 | 9094.74 | 29113.8 | 1223480 | 8136.39 | × | 99347 | 21205.1 | × | × |
| MSFragger- | 905 | 0 | 1 | 0 | 0 | 210958 | 14850.4 | 133796 | 66713.7 | 266814 | 16537.3 | 49827.5 | 1422.43 | 6931.88 | 177888 | × | 1471.51 | 17899.3 | × | × | × |
| MSFragger- | 995 | 0 | 1 | 0 | 0 | 189272 | 17845.2 | 106565 | 3507.05 | 343233 | × | 2426.24 | × | × | 150615 | × | × | × | × | × | × |
| MSFragger- | 1032 | 0 | 1 | 0 | 0 | 241900 | 20401.5 | 173223 | 92579.4 | 558803 | × | 8945.37 | × | × | 278226 | × | × | × | × | × | × |
| MSFragger- | 1070 | 0 | 1 | 0 | 0 | 261280 | 11877.2 | 174863 | 66192.3 | 231766 | 15199.7 | 27759.1 | 4398.95 | 8817.88 | 369758 | 3576.8 | × | 16952.2 | 6071.09 | × | × |
| MSFragger- | 1321 | 0 | 1 | 0 | 0 | 285643 | 12832 | 194212 | 91549.7 | 369844 | 33986.7 | 91668.9 | × | 3160.49 | 343392 | 6144.11 | × | 23418.5 | × | × | × |
| MSFragger- | 1407 | 0 | 1 | 0 | 0 | 87374.6 | 5297.85 | 52557.9 | 1114.42 | 105149 | 2918.46 | 9402.86 | × | 731.016 | 98090.1 | × | × | 7806.44 | × | × | × |
| GRable | 54 | 0 | 0 | 0 | 1 | 214288 | 15521.8 | 134660 | 73983.2 | 267332 | 13665.4 | 43516.7 | 2066.8 | 17386.6 | 203328 | × | × | 13427.2 | 6289.7 | × | × |
| GRable | 314 | 0 | 0 | 0 | 1 | 665395 | 30599.5 | 513030 | 179496 | 635161 | 41567.7 | 97416.8 | 19230.8 | 80205.4 | 909732 | × | × | 31897.4 | 18031 | × | × |
| GRable | 334 | 0 | 0 | 0 | 1 | 570059 | 34080.2 | 371156 | 10705.7 | 644030 | × | 29694.4 | 15968.1 | 113918 | 604098 | × | × | × | 20674.7 | × | × |
| GRable | 340 | 0 | 0 | 0 | 1 | 152690 | 11846.1 | 3348.31 | 41559.6 | 176817 | 8015.06 | 22939.2 | 3149.6 | 19316.2 | 104420 | × | × | × | × | × | × |
| GRable | 734 | 0 | 0 | 0 | 1 | 1067170 | 49031.2 | 670596 | 240908 | 817490 | 66266.3 | 147551 | 6065.14 | 21417.4 | 1466800 | 14760.8 | × | 88057.8 | 13504 | 4392.03 | × |
| GRable | 870 | 0 | 0 | 0 | 1 | 1272700 | 51282.8 | 852900 | 294469 | 984267 | 67303.8 | 156123 | 15691 | 39687.1 | 1794440 | 25805.6 | × | 117734 | 28974.4 | × | × |
| GRable | 952 | 0 | 0 | 0 | 1 | 892813 | 38711.6 | 555468 | 222027 | 976503 | 70511.5 | 148176 | 9094.74 | 29113.8 | 1223480 | 8136.39 | × | 99347 | 21205.1 | × | × |
| GRable | 1066 | 0 | 0 | 0 | 1 | 1288070 | 52241.4 | 790305 | 294686 | 1004060 | 49107.5 | 130518 | 9604.33 | 52713 | 1594600 | 24693.7 | × | 78469.7 | 30664.1 | × | × |

| Software | No. | Intensity of Fuc signal (147.0657) | Intensity of Hex+HexNAc+Fuc signal (511.1901) | Intensity of Hex+Fuc signal (308.1107) | Intensity of HesNAc+Fuc signal (349.1373) | Intensity of Y0 signal | Intensity of signal Y1 | Intensity of signal Y1F | Intensity of signal Y0(z=2) | Intensity of Y1(z=2) signal | Intensity of Y1F(z=2) signal | Intensity of Y0(z=3) signal | Intensity of Y1(z=3) signal | Intensity of Y1F(z=3) signal | Y0 or Y1 ions detected or not ? (0: Not detected, 1: Detected) | Detected y ions | Detected b ions |
|---|---|---|---|---|---|---|---|---|---|---|---|---|---|---|---|---|---|
| Byonic | 21 | × | × | × | × | × | × | × | × | × | × | × | × | × | 0 | × | × |
| Byonic | 46 | × | × | 38317.9 | 1433.75 | 22301 | 110045 | 65320.9 | 15871.8 | 50408.5 | 9767.16 | × | × | × | 1 | y1, y2, y3, y4, y5, y6, y7, y8, y9, y11 | b2, b9, b10 |
| Byonic | 47 | × | × | × | × | 25606 | 97630.3 | 66249.9 | 14302.8 | 57681.3 | × | × | × | × | 1 | y4, y5, y6, y7, y8, y9, y11 | b2, b10 |
| Byonic | 48 | × | × | × | × | 3600.93 | 45503.2 | × | 2772.7 | 21290 | × | × | × | × | 1 | y1, y4, y5, y7, y11 | b1, b9 |
| Byonic | 51 | × | × | 53087.6 | × | 42650.7 | 294238 | × | 24636.1 | 106468 | × | × | × | × | 1 | y1, y4, y5, y6, y7, y8, y9, y11 | b2, b6, b9 |
| Byonic | 83 | × | × | × | × | × | × | × | 47582.7 | 249119 | 69884.8 | × | 7119.73 | × | 1 | × | × |
| Byonic | 107 | 1315.25 | × | × | × | 3537.83 | 17221.8 | × | × | 1213.08 | × | × | × | × | 1 | y1, y5, y7 | b2 |
| Byonic | 122 | 11464.2 | × | × | × | 104925 | × | × | 170632 | 699188 | × | × | × | × | 1 | y1, y4, y7, y8, y9, y10, y11, y17 | b2, b3, b4, b5, b6, b7, b8, b9, b10, b12, b13 |
| Byonic | 129 | × | × | 4133.39 | × | 21722.5 | × | × | 29182.5 | 126966 | × | × | × | × | 1 | y4, y5, y8, y9, y10, y11, y17 | b2, b3, b4, b5, b7, b8, b9, b13 |
| Byonic | 260 | 1470.78 | × | × | × | × | × | × | × | × | × | × | × | × | 0 | × | × |
| Byonic | 286 | × | × | × | × | 14752.6 | 96664.4 | × | × | 11395.7 | × | × | × | × | 1 | y1, y2, y3, y6, y11 | b9 |
| Byonic | 288 | 1936.2 | × | × | × | 17033 | 75285 | 48417.2 | × | 17747.8 | × | × | × | × | 1 | y1, y2, y3, y4, y6, y8, y11 | × |
| Byonic | 308 | × | × | × | × | × | × | × | × | 8023.93 | 13669.1 | × | × | × | 1 | × | × |
| Byonic | 338 | × | × | × | × | 13654 | 31077.8 | × | 25470.4 | 73586.4 | 12059.6 | × | × | × | 1 | y1, y3, y4, y7, y8, y10, y15 | b3, b14 |
| Byonic | 341 | × | × | × | × | × | × | × | × | × | × | × | × | × | 0 | y1 | b2 |
| Byonic | 344 | × | × | × | × | × | × | × | × | × | × | × | × | × | 0 | y1 | b2 |
| Byonic | 381 | × | × | × | × | × | × | × | 16353.2 | 57360.2 | × | × | × | × | 1 | × | × |
| Byonic | 415 | × | × | × | × | × | × | × | × | 21567.3 | × | × | × | × | 1 | × | × |
| Byonic | 474 | × | × | × | × | × | × | × | × | × | × | × | × | × | 0 | y1 | b3 |
| Byonic | 484 | 47402 | × | × | × | × | × | × | × | × | × | × | × | × | 0 | × | × |
| Byonic | 491 | 19563 | × | × | × | × | × | × | × | × | × | × | × | × | 0 | × | × |
| Byonic | 494 | 20109 | × | × | × | × | × | × | × | × | × | × | × | × | 0 | × | × |
| Byonic | 503 | 892.026 | × | × | × | × | × | × | × | × | × | × | × | × | 0 | × | × |
| Byonic | 550 | × | × | × | × | 99863.3 | 723272 | × | 3667.89 | 8503.36 | × | × | × | × | 1 | × | × |
| Byonic | 573 | × | × | × | × | × | × | × | × | × | × | × | × | × | 0 | y1, y2 | × |
| Byonic | 631 | × | × | × | × | × | × | × | × | × | × | × | × | × | 0 | × | × |
| Byonic | 652 | × | × | 18285.9 | × | × | × | × | 43951.7 | 108956 | × | × | × | × | 1 | × | × |
| Byonic | 666 | × | × | × | × | × | × | × | × | × | × | × | × | × | 0 | y1, y4, y6, y7, y8, y9 | b2, b6, b8 |
| Byonic | 680 | × | × | × | × | × | × | × | 32458.4 | 81645 | × | × | × | × | 1 | × | × |
| Byonic | 697 | 1981.95 | × | × | × | 6297.4 | 20652.8 | × | 9045.92 | 53417.6 | 12238 | × | × | × | 1 | y1, y2, y3, y4, y5, y6, y7, y8, y9, y11, y14 | b3, b12, b13 |
| Byonic | 717 | 862.996 | × | × | × | × | × | × | × | × | × | × | × | × | 0 | × | × |
| Byonic | 753 | × | × | 10243.8 | × | 17271.4 | 54152.7 | 23931 | × | 10479.9 | × | × | × | × | 1 | × | × |
| Byonic | 847 | × | × | × | × | × | × | × | × | × | × | × | × | × | 0 | × | × |
| Byonic | 856 | × | × | × | × | × | × | × | × | × | × | × | × | × | 0 | × | × |
| Byonic | 908 | 631.063 | × | × | × | × | 7935.95 | 7332.43 | × | 2759.29 | × | × | × | × | 1 | y1 | b1, b6 |
| Byonic | 965 | × | × | × | × | × | × | × | × | × | × | × | × | × | 0 | × | × |
| Byonic | 987 | × | × | 40457.3 | × | 101408 | 450735 | × | 4845.56 | 27709.3 | × | × | × | × | 1 | y2, y3, y4, y8, y10 | b2, b2, b6, b8 |
| Byonic | 1062 | × | × | × | × | 90099.7 | 148994 | × | 23642.6 | 40002 | × | × | × | × | 1 | y10, y13 | b3 |
| Byonic | 1066 | × | × | × | × | × | × | × | × | × | × | × | × | × | 0 | y1, y2, y9 | b2 |
| Byonic | 1075 | 12953.5 | × | × | × | 137815 | 515819 | × | 45177.8 | 219882 | × | × | × | × | 1 | × | × |
| Byonic | 1077 | × | × | 47617.1 | × | 68852.3 | 256380 | × | 13882.3 | 95291.2 | × | × | × | × | 1 | × | × |
| Byonic | 1081 | 1699.24 | × | 12504.2 | × | 10172.3 | 38723.9 | × | 1484.74 | 11336.5 | × | × | × | × | 1 | × | × |
| Byonic | 1130 | × | × | 75453.5 | × | 190549 | 351054 | × | 70058.8 | 77635 | × | × | × | × | 1 | y2, y3, y4, y5, y6, y7, y8, y9, y10, y11, y12 | b4, b8, b9, b12 |
| Byonic | 1211 | 2426.38 | × | × | × | × | × | × | × | × | × | × | × | 3408.99 | 0 | y1 | b3 |
| Byonic | 1212 | × | × | × | × | × | × | × | × | × | × | × | × | × | 0 | × | × |
| Byonic | 1213 | 2804.19 | × | × | × | × | × | × | 4854.96 | × | × | × | × | × | 1 | y1 | b3, b4 |
| Byonic | 1214 | 1393.51 | × | × | × | × | × | × | × | × | × | × | × | × | 0 | y1 | b3, b4 |
| Byonic | 1216 | 5809.04 | × | × | × | × | × | × | × | × | × | × | × | × | 0 | y1 | × |
| Byonic | 1217 | × | × | × | × | × | × | × | × | × | × | × | × | × | 0 | × | × |
| Byonic | 1272 | 6274.61 | × | × | × | × | × | × | 22291 | 42457.5 | × | × | × | × | 1 | y1, y3, y7, y8, y11, y12 | b6 |
| Byonic | 1307 | × | × | 1504.13 | × | × | × | × | × | × | × | × | × | × | 0 | × | × |
| Byonic | 1316 | × | × | × | × | 58531.4 | 102487 | 32716.7 | × | × | × | × | × | × | 1 | y4, y6, y8 | × |
| Byonic | 1323 | 1272.35 | × | × | × | × | × | × | × | × | × | × | × | × | 0 | × | × |
| Byonic | 1360 | × | × | × | × | × | × | × | × | × | × | × | × | × | 0 | × | × |
| Byonic | 1433 | × | × | × | × | × | 6312.94 | × | × | × | × | × | × | × | 1 | × | × |
| pGlyco3 | 664 | 1526.87 | 1489.38 | 13594 | × | 5277.69 | 15982.5 | × | 15668.6 | 62403.3 | 7985.08 | × | × | × | 1 | y1, y2, y3, y4, y5, y6, y7, y8, y9, y14 | b1, b3, b4, b5 |
| MSFragger- | 466 | 11464.2 | × | × | × | 99056.4 | × | × | 177103 | 669251 | 18774.5 | × | × | × | 1 | y1, y4, y7, y8, y9, y10, y11, y17 | b2, b3, b4, b5, b6, b7, b8, b9, b10, b12, b13 |
| MSFragger- | 728 | × | × | × | × | × | × | × | × | × | × | × | × | × | 0 | × | × |
| MSFragger- | 856 | 663.427 | × | 11871.8 | × | × | × | × | 9311.05 | 19742 | 1294.59 | × | × | × | 1 | × | × |
| MSFragger- | 857 | × | × | 9094.74 | × | × | × | × | × | × | × | × | × | × | 0 | × | × |
| MSFragger- | 905 | 1125.52 | × | 1422.43 | × | × | × | × | × | 5255.1 | × | × | × | × | 1 | × | × |
| MSFragger- | 995 | × | × | × | × | × | × | × | × | 7269.88 | × | × | 3049.91 | × | 1 | × | × |
| MSFragger- | 1032 | × | × | × | × | × | × | × | × | × | × | × | × | × | 0 | × | × |
| MSFragger- | 1070 | × | × | 4398.95 | × | × | × | × | × | 2634.54 | 8967.34 | × | × | × | 1 | × | × |
| MSFragger- | 1321 | × | × | × | × | × | × | × | 9729.47 | 44322 | 3517.08 | × | × | × | 1 | × | × |
| MSFragger- | 1407 | × | × | × | × | × | × | × | × | × | × | × | × | × | 0 | × | × |
| GRable | 54 | 1373.07 | × | 2066.8 | × | × | 9601.04 | 6619.66 | 2606.41 | 6252.57 | × | × | × | × | 1 | × | × |
| GRable | 314 | × | × | 19230.8 | × | 43224.1 | 240561 | × | × | × | × | × | × | × | 1 | y7 | × |
| GRable | 334 | × | × | 15968.1 | × | × | 61272.1 | 33261.2 | × | 35057.7 | × | × | × | × | 1 | y5 | × |
| GRable | 340 | 2890.74 | × | 3149.6 | × | 3159.48 | 29589.2 | 8779.44 | × | 13835.6 | × | × | × | × | 1 | y3, y5, y11 | × |
| GRable | 734 | 2715.22 | × | 6065.14 | × | 28488 | 83133.3 | 38616.7 | × | 4218.73 | × | × | × | × | 1 | × | × |
| GRable | 870 | × | × | 15691 | × | 22714.8 | 89416.4 | 87944.7 | × | 13348.5 | × | × | × | × | 1 | y2, y3, y5, y6, y7, y8, y13 | b2 |
| GRable | 952 | × | × | 9094.74 | × | × | × | × | 25167.9 | 69332.8 | × | × | × | × | 1 | × | × |
| GRable | 1066 | × | × | 9604.33 | × | × | × | × | 29852.7 | 95686.9 | 27995.8 | × | × | × | 1 | × | × |
