## Supplementary material for "Comparative Evaluation of Glycoproteomics Software for Rare Glycopeptide Identification": Table S3

| Software | No. | Scan | Proteins | Genes | Pep. seq. | pep_var_mod_pos | Glycosylation | No. of | No. of Hex | No. of Fuc | No. of | No. of | No. of KDN | Composition | glycan type | Identified by Byonic | Identified by pGlyco3 | Identified by GRable | Intensity of | Intensity of | Intensity of | Intensity of | Intensity of | Intensity of | Intensity of | Intensity of | Intensity of | Intensity of | Intensity of | Intensity of | Intensity of | Intensity of | Intensity of | Intensity | Intensity | Intensity of | Intensity of | Hex+KDN | Intensity of | Intensity of | Intensity of | Intensity of | Intensity of | Intensity of | Intensity of | Intensity of | Intensity of | Intensity of | Intensity of | Intensity of | Intensity of | Y0 or Y1 ions | Detected y ions | Detected b ions |
|---|---|---|---|---|---|---|---|---|---|---|---|---|---|---|---|---|---|---|---|---|---|---|---|---|---|---|---|---|---|---|---|---|---|---|---|---|---|---|---|---|---|---|---|---|---|---|---|---|---|---|---|---|---|---|
| Byonic | 36 | 4615 | P01871 | IGHM | YKNNSDISSTR | N3(HexNAc(3)Hex(5)Fuc(1)250.068868) | 46 | 3 | 5 | 1 | 0 | 0 | 1 | 3:5:1:0:0:1 | hybrid | 1 | 1 | 1 | 1147460 | 59667.6 | 778107 | 353719 | 1535560 | × | 6484.95 | × | × | 817170 | 4104.37 | 3937.6 | × | × | 24013.2 | × | × | × | × | × | × | × | × | × | 81830.2 | 388868 | 208828 | 54928.6 | 242934 | 35528.3 | × | × | × | 1 | y1, y2, y3, y4, y5, y6, y7, y8, y9, y11 | b2, b9, b10 |
| Byonic | 44 | 4543 | P01871 | IGHM | YKNNSDISSTR | N3(HexNAc(3)Hex(6)Fuc(1)250.068868) | 46 | 3 | 6 | 1 | 0 | 0 | 1 | 3:6:1:0:0:1 | hybrid | 1 | 1 | 1 | 1237340 | 61434.1 | 896979 | 401099 | 1774170 | × | × | × | × | 1065150 | × | 5667.73 | × | × | 36489.6 | × | × | × | × | × | × | × | × | × | 81336 | 356477 | 187777 | 58866 | 245169 | 37530.3 | × | × | × | 1 | y1, y3, y4, y5, y6, y7, y8, y9, y11 | b1, b2, b9, b10 |
| Byonic | 47 | 4618 | P01871 | IGHM | YKNNSDISSTR | N3(HexNAc(5)Hex(5)Fuc(1)250.068868) | 46 | 5 | 5 | 1 | 0 | 0 | 1 | 5:5:1:0:0:1 | complex | 1 | 1 | 0 | 456593 | 31091.9 | 333172 | 192464 | 844750 | 82709.9 | 253515 | × | × | 469518 | × | × | 44324.1 | × | × | × | 9620.44 | × | × | × | × | × | × | × | 25606 | 97630.3 | 66249.9 | 14302.8 | 57681.3 | × | × | × | × | 1 | y4, y5, y6, y7, y8, y9, y11 | b2, b10 |
| Byonic | 50 | 4620 | P01871 | IGHM | YKNNSDISSTR | N3(HexNAc(5)Hex(6)Fuc(1)250.068868) | 46 | 5 | 6 | 1 | 0 | 0 | 1 | 5:6:1:0:0:1 | complex | 1 | 1 | 0 | 629433 | 33606.2 | 459435 | 207351 | 894315 | 106192 | 337267 | × | × | 801556 | × | × | 62766.6 | × | × | × | 17046.1 | 5805.34 | × | × | × | × | × | × | 29590 | 123579 | 81931.4 | 18250.3 | 59872.6 | 5161.57 | × | × | × | 1 | y2, y4, y5, y7, y8, y11 | b2, b10 |
| Byonic | 52 | 4496 | P01871 | IGHM | YKNNSDISSTR | N3(HexNAc(3)Hex(6)250.068868) | 46 | 3 | 6 | 0 | 0 | 0 | 1 | 3:6:0:0:0:1 | hybrid | 1 | 1 | 1 | 428497 | 24801 | 298455 | 129722 | 583075 | 4760.8 | 12039.3 | × | × | 363095 | × | 949.995 | 970.66 | × | 9006.94 | × | × | × | 996.52 | × | 900.599 | × | × | × | 31817.9 | 217729 | × | 20898.5 | 95766.2 | × | × | × | × | 1 | y1, y3, y4, y5, y6, y7, y8, y9, y11 | b1, b2, b9, b10 |
| Byonic | 85 | 12325 | P43121 | MCAM | CGLSQSQGNLSHVDWFSVHK | C1(Carbamidomethyl); N9(HexNAc(5)Hex(5)Fuc(1)250.068868) | 56 | 5 | 5 | 1 | 0 | 0 | 1 | 5:5:1:0:0:1 | complex | 1 | 1 | 0 | 1157360 | 72321.8 | 737562 | 351022 | 1263760 | × | 13461.2 | × | × | 923059 | × | × | × | × | 18334.2 | × | × | × | × | × | × | × | × | × | × | × | × | 47582.7 | 249119 | 69884.8 | × | 7119.73 | × | 1 | × | × |
| Byonic | 130 | 9467 | Q16563 | SYPL1 | GQTEIQVNCPPAVTENK | C9(Carbamidomethyl); N16(HexNAc(5)Hex(5)250.068868) | 71 | 5 | 5 | 0 | 0 | 0 | 1 | 5:5:0:0:0:1 | complex | 1 | 0 | 0 | 165977 | 11164.5 | 111298 | 53335.1 | 245928 | 22378.3 | 60888 | × | × | 174469 | 1897.04 | × | 12916.8 | × | × | × | 1576.64 | × | × | × | 2345.95 | × | × | × | × | × | × | × | 4089.61 | × | × | × | × | 1 | y1, y4, y8, y9, y10 | b2, b2 |
| Byonic | 275 | 15651 | Q9BY67 | CADM1 | FQLLNFSSSELK | N5(HexNAc(6)Hex(7)Fuc(1)250.068868) | 101 | 6 | 7 | 1 | 0 | 0 | 1 | 6:7:1:0:0:1 | complex | 1 | 0 | 0 | 404266 | 19515 | 254098 | 89729.5 | 288556 | 4976.45 | 13297.6 | × | × | 433149 | 4980.71 | 1829.39 | 1676.84 | × | 8944.54 | × | × | × | × | × | 977.156 | × | × | × | 8761.38 | 20987.3 | 8286.33 | × | 3263.69 | × | × | × | × | 1 | × | × |
| Byonic | 288 | 9677 | P20138 | CD33 | NNCSLSIVDAR | N2(HexNAc(5)Hex(6)250.068868); C3(Carbamidomethyl) | 100 | 5 | 6 | 0 | 0 | 0 | 1 | 5:6:0:0:0:1 | complex | 1 | 1 | 0 | 385381 | 24683.2 | 283143 | 133623 | 614659 | 49550.7 | 145898 | × | 3856.26 | 502374 | × | × | 37071.2 | × | × | × | 2824.09 | × | × | × | × | × | × | × | 25833.1 | 211594 | × | × | 22192.7 | × | × | × | × | 1 | y1, y2, y3, y4, y6, y7, y8, y9, y11 | × |
| Byonic | 290 | 9705 | P20138 | CD33 | NNCSLSIVDAR | N2(HexNAc(5)Hex(5)250.068868); C3(Carbamidomethyl) | 100 | 5 | 5 | 0 | 0 | 0 | 1 | 5:5:0:0:0:1 | complex | 1 | 1 | 0 | 147031 | 10582.6 | 100666 | 62829.4 | 264168 | 19141.1 | 59555.7 | × | × | 141102 | × | × | 8824.92 | × | × | × | 2325.9 | × | × | × | × | × | × | × | 14752.6 | 96664.4 | × | × | 11395.7 | × | × | × | × | 1 | y1, y2, y3, y6, y11 | b9 |
| Byonic | 292 | 9765 | P20138 | CD33 | NNCSLSIVDAR | N2(HexNAc(5)Hex(5)Fuc(1)250.068868); C3(Carbamidomethyl) | 100 | 5 | 5 | 1 | 0 | 0 | 1 | 5:5:1:0:0:1 | complex | 1 | 0 | 0 | 280474 | 18879.5 | 192943 | 85101.8 | 389987 | 32197.1 | 97340.9 | × | × | 325804 | × | × | 19419.9 | × | × | × | 2433.81 | × | × | × | 1936.2 | × | × | × | 17033 | 75285 | 48417.2 | × | 17747.8 | × | × | × | × | 1 | y1, y2, y3, y4, y6, y8, y11 | × |
| Byonic | 312 | 15414 | P01903 | HLA-DRA | RSNYTPITNVPPEVTVLTNSPVELR | N3(HexNAc(5)Hex(5)Fuc(1)250.068868) | 103 | 5 | 5 | 1 | 0 | 0 | 1 | 5:5:1:0:0:1 | complex | 1 | 0 | 0 | 222602 | 12575.5 | 148080 | 71968 | 268268 | 5845.08 | 14758.9 | × | × | 253487 | 1116.69 | × | 1787.16 | × | 4744.02 | × | × | × | × | × | × | × | × | × | × | × | × | × | 8023.93 | 13669.1 | × | × | × | 1 | × | × |
| Byonic | 329 | 7226 | O75326 | SEMA7A | NASVR | N1(HexNAc(5)Hex(6)Fuc(3)NeuAc(2)250.068868) | 105 | 5 | 6 | 3 | 2 | 0 | 1 | 5:6:3:2:0:1 | complex | 1 | 0 | 0 | 355640 | 19483.9 | 242260 | 5188.8 | 472457 | 35796.8 | 75424.2 | × | × | 425521 | × | × | 22099.8 | × | × | × | × | × | × | × | × | × | × | × | × | × | × | × | × | × | × | × | × | 0 | × | b2, b2 |
| Byonic | 342 | 7114 | P10321 | HLA-C | GYYNQSEDGSHTLQR | N4(HexNAc(5)Hex(5)Fuc(1)250.068868) | 110 | 5 | 5 | 1 | 0 | 0 | 1 | 5:5:1:0:0:1 | complex | 1 | 1 | 0 | 617065 | 36505.2 | 423719 | 200818 | 733535 | × | 3916.31 | × | × | 669751 | × | × | × | × | 13541.3 | × | × | × | × | × | × | × | × | × | 13654 | 31077.8 | × | 25470.4 | 73586.4 | 12059.6 | × | × | × | 1 | y1, y3, y4, y7, y8, y10, y15 | b3, b14 |
| Byonic | 384 | 16939 | P04233 | CD74 | MATPLLMQALPMGALPQGPMQNATK | N22(HexNAc(6)Hex(6)Fuc(2)250.068868) | 130 | 6 | 6 | 2 | 0 | 0 | 1 | 6:6:2:0:0:1 | complex | 1 | 0 | 0 | 5011.94 | 6332.5 | 59714.8 | 41521.5 | 242852 | 3487.84 | × | × | × | 118746 | × | × | × | × | × | × | × | × | × | × | × | × | × | × | × | × | × | 16353.2 | 57360.2 | × | × | × | × | 1 | × | × |
| Byonic | 395 | 7791 | Q9NRG0 | CHRAC1 | EEDEENDNDNESDHDEADS | N10(HexNAc(3)Hex(5)250.068868) | 122 | 3 | 5 | 0 | 0 | 0 | 1 | 3:5:0:0:0:1 | hybrid | 1 | 0 | 0 | 1280.86 | × | 12222.7 | 5318.86 | 15355.4 | × | × | × | × | 19134.9 | × | × | × | × | × | × | × | × | × | × | × | × | × | × | × | × | × | × | × | × | × | × | × | 0 | × | b2, b3, b4 |
| Byonic | 485 | 14102 | P04233 | CD74 | YGNMTEDHVMHLLQNADPLK | N3(HexNAc(3)Hex(4)Fuc(2)250.068868) | 136 | 3 | 4 | 2 | 0 | 0 | 1 | 3:4:2:0:0:1 | hybrid | 1 | 0 | 0 | 373072 | 29702.3 | 212169 | 145673 | 782004 | 36250.3 | 92439 | × | × | 238479 | × | × | 9123.77 | × | × | × | × | × | × | × | 31532.3 | × | × | × | × | × | × | × | × | × | × | × | × | 0 | × | × |
| Byonic | 512 | 13987 | Q99523 | SORT1 | DITDLINNTFIRTEFGMAIGPENSGK | N7(HexNAc(6)Hex(7)Fuc(1)NeuAc(1)500.137736); M17(Oxidation) | 162 | 6 | 7 | 1 | 1 | 0 | 2 | 6:7:1:1:0:2 | complex | 1 | 0 | 0 | 153453 | 13026.2 | 90147.5 | 42385.5 | 214057 | 3135.29 | 11793 | × | × | 151911 | × | × | 2754.78 | × | × | × | × | × | × | × | × | × | × | × | × | × | × | × | × | × | × | × | × | 0 | × | × |
| Byonic | 531 | 8042 | Q92896 | GLG1 | LNLTTDPK | N2(HexNAc(5)Hex(6)250.068868) | 165 | 5 | 6 | 0 | 0 | 0 | 1 | 5:6:0:0:0:1 | complex | 1 | 1 | 0 | 1355820 | 94485.8 | 1070990 | 495735 | 2322260 | 209757 | 648942 | × | × | 1687780 | × | × | 97044.2 | × | × | × | 37008.5 | 9387.86 | × | × | × | × | × | × | 137271 | 787612 | × | × | × | × | × | × | × | 1 | y2, y5, y8 | b3, b6 |
| Byonic | 553 | 14104 | P04004 | VTN | NGSLFAFR | N1(HexNAc(4)Hex(5)NeuAc(1)250.068868) | 169 | 4 | 5 | 0 | 1 | 0 | 1 | 4:5:0:1:0:1 | complex | 1 | 1 | 0 | 252092 | 23169.9 | 8412.62 | 161087 | 1048170 | 290093 | 953931 | × | × | 318680 | × | × | 66756.7 | × | × | 1265.01 | 35679 | 7917.38 | × | × | × | × | × | × | 99863.3 | 723272 | × | 3667.89 | 8503.36 | × | × | × | × | 1 | × | × |
| Byonic | 633 | 11954 | P01871 | IGHM | VDHRGLTFQQNASSMCVPDQDTAIR | N11(HexNAc(4)Hex(5)NeuGc(1)250.068868); M15(Oxidation); C16(Carbamidomethyl) | 209 | 4 | 5 | 0 | 0 | 1 | 1 | 4:5:0:0:1:1 | complex | 1 | 0 | 0 | 476449 | 27548.6 | 313485 | 123017 | 447836 | × | 9582.35 | × | × | 574841 | 4748.51 | × | × | × | × | × | × | × | × | × | × | × | × | × | × | × | × | × | × | × | × | × | × | 0 | × | × |
| Byonic | 662 | 12095 | P01871 | IGHM | GLTFQQNASSMCVPDQDTAIR | N7(HexNAc(5)Hex(6)NeuAc(1)500.137736); C12(Carbamidomethyl) | 209 | 5 | 6 | 0 | 1 | 0 | 2 | 5:6:0:1:0:2 | complex | 1 | 0 | 0 | 154816 | 6549.92 | 97992 | 37695.1 | 151518 | × | 5940.62 | × | × | 148980 | × | × | 3792.85 | × | × | × | × | × | × | × | × | × | × | × | × | × | × | × | 4752.32 | × | × | × | × | 1 | × | × |
| Byonic | 681 | 12743 | P01871 | IGHM | GLTFQQNASSMCVPDQDTAIR | N7(HexNAc(5)Hex(5)Fuc(1)250.068868); C12(Carbamidomethyl) | 209 | 5 | 5 | 1 | 0 | 0 | 1 | 5:5:1:0:0:1 | complex | 1 | 0 | 1 | 790282 | 61810.2 | 522829 | 253831 | 1111610 | × | 15642.4 | × | × | 795786 | 8283.04 | × | × | × | 9937.02 | × | × | × | × | × | × | × | × | × | × | × | × | 32458.4 | 81645 | × | × | × | × | 1 | × | × |
| Byonic | 696 | 9272 | Q92896 | GLG1 | GNITEYQCHQYITK | N2(HexNAc(5)Hex(6)Fuc(1)NeuAc(1)250.068868); C8(Carbamidomethyl) | 210 | 5 | 6 | 1 | 1 | 0 | 1 | 5:6:1:1:0:1 | complex | 1 | 1 | 0 | 419209 | 20400 | 295610 | 117820 | 510072 | 33171.6 | 80013.8 | × | × | 525479 | 1926.09 | 1740.47 | 32778 | × | 11661.9 | × | 4561.32 | 1283.5 | × | × | 1981.95 | × | × | × | 6297.4 | 20652.8 | × | 9045.92 | 53417.6 | 12238 | × | × | × | 1 | y1, y2, y3, y4, y5, y6, y7, y8, y9, y11, y14 | b3, b12, b13 |
| Byonic | 726 | 8365 | Q9GZU1 | MCOLN1 | NLTLK | N1(HexNAc(5)Hex(6)NeuAc(1)NeuGc(1)250.068868) | 220 | 5 | 6 | 0 | 1 | 1 | 1 | 5:6:0:1:1:1 | complex | 1 | 0 | 0 | 184940 | 11832.8 | 109512 | 52395.7 | 169938 | 3966.16 | 16002.4 | × | × | 139764 | × | × | × | × | × | × | × | × | × | × | 2413.34 | × | × | × | × | × | × | × | 42342.9 | × | × | × | × | 1 | y1 | × |
| Byonic | 797 | 3958 | P05107 | ITGB2 | NVTR | N1(HexNAc(5)Hex(6)Fuc(4)NeuAc(2)250.068868) | 254 | 5 | 6 | 4 | 2 | 0 | 1 | 5:6:4:2:0:1 | complex | 1 | 0 | 0 | 505753 | 18869.6 | 344422 | 116310 | 361922 | × | × | × | 1466.36 | 668230 | 8158.07 | × | × | × | × | × | × | × | × | × | × | × | × | × | × | × | × | × | × | × | × | × | × | 0 | y1 | × |
| Byonic | 854 | 17290 | P53634 | CTSC | ILTNNSQTPILSPQEVVSCSQYAQGCEGGFPYLIAGK | N4(HexNAc(5)Hex(5)Fuc(1)250.068868); C19(Carbamidomethyl); C26(Carbamidomethyl) | 276 | 5 | 5 | 1 | 0 | 0 | 1 | 5:5:1:0:0:1 | complex | 1 | 0 | 0 | 362457 | 22449.1 | 223193 | 102774 | 408486 | 7316.76 | 19582.5 | × | × | 422798 | 7054.04 | 3325.85 | 7521.73 | × | 16080.6 | × | × | × | × | × | × | × | × | × | × | × | × | × | × | × | × | × | × | 0 | × | × |
| Byonic | 904 | 4080 | P10909 | CLU | HNSTGCLR | N2(HexNAc(5)Hex(5)Fuc(1)250.068868); C6(Carbamidomethyl) | 291 | 5 | 5 | 1 | 0 | 0 | 1 | 5:5:1:0:0:1 | complex | 1 | 0 | 0 | 121541 | 7184.86 | 80978.8 | 40752.8 | 128590 | 10930.5 | 40195.1 | × | × | 106654 | × | × | 5109.99 | × | × | × | 950.273 | × | × | × | 631.063 | × | × | × | × | 7935.95 | 7332.43 | × | 2759.29 | × | × | × | × | 1 | y1 | b1, b6 |
| Byonic | 1049 | 6275 | O15265 | ATXN7 | NKTR | N1(HexNAc(6)Hex(6)Fuc(3)NeuAc(1)500.137736) | 391 | 6 | 6 | 3 | 1 | 0 | 2 | 6:6:3:1:0:2 | complex | 1 | 0 | 0 | 533156 | 64653 | 14060.1 | 232832 | 513084 | × | × | × | × | 582808 | × | × | × | × | × | × | × | × | 42799.5 | × | × | × | × | × | × | × | × | × | × | × | 48549.7 | × | × | 1 | y1 | × |
| Byonic | 1057 | 6907 | Q6AHZ1 | ZNF518A | AEEGPNASSGFMK | N6(HexNAc(5)Hex(5)250.068868) | 409 | 5 | 5 | 0 | 0 | 0 | 1 | 5:5:0:0:0:1 | complex | 1 | 0 | 0 | 558379 | 32939.6 | 402368 | 199601 | 741871 | 67250.9 | 218453 | × | × | 620892 | × | × | 23405.7 | × | × | × | 5045.59 | 4184.16 | × | × | × | × | × | × | 90099.7 | 148994 | × | 23642.6 | 40002 | × | × | × | × | 1 | y10, y13 | b3 |
| Byonic | 1068 | 12485 | P08575 | PTPRC | SFHNFTLCYIK | N4(HexNAc(5)Hex(5)250.068868); C8(Carbamidomethyl) | 421 | 5 | 5 | 0 | 0 | 0 | 1 | 5:5:0:0:0:1 | complex | 1 | 1 | 1 | 2657720 | 160159 | 1741270 | 816208 | 2989670 | × | 7892.2 | × | 6554.51 | 2457080 | × | 14103.3 | × | × | 54884.8 | × | × | × | × | × | 12953.5 | × | × | × | 137815 | 515819 | × | 45177.8 | 219882 | × | × | × | × | 1 | × | × |
| Byonic | 1118 | 6777 | P08575 | PTPRC | NGSAAMCHFTTK | N1(HexNAc(3)Hex(6)250.068868); C7(Carbamidomethyl) | 470 | 3 | 6 | 0 | 0 | 0 | 1 | 3:6:0:0:0:1 | hybrid | 1 | 1 | 1 | 696148 | 2394.08 | 527490 | 247554 | 1082510 | × | 6124.3 | × | × | 618281 | × | × | × | × | 19074.8 | × | × | × | × | × | 4056.12 | × | × | × | 163712 | 307456 | × | 57760 | 97741.3 | × | × | × | × | 1 | y1, y2, y5, y6, y7, y8, y9, y11, y12 | b8 |
| Byonic | 1204 | 10485 | P11717 | IGF2R | NGSK | N1(HexNAc(6)Hex(7)NeuAc(2)250.068868) | 543 | 6 | 7 | 0 | 2 | 0 | 1 | 6:7:0:2:0:1 | complex | 1 | 0 | 0 | 72876.4 | 5585.85 | 37179.2 | 1067.86 | 58207.4 | 2300.45 | 7565.7 | × | × | 51342.1 | × | × | 3197.55 | × | × | × | 636.869 | × | × | × | 1393.51 | × | × | × | × | × | × | × | × | × | × | × | × | 0 | y1 | b3, b4 |
| Byonic | 1207 | 16754 | P11717 | IGF2R | NGSK | N1(HexNAc(6)Hex(7)NeuAc(1)NeuGc(1)500.137736) | 543 | 6 | 7 | 0 | 1 | 1 | 2 | 6:7:0:1:1:2 | complex | 1 | 0 | 0 | 70790 | 4684.74 | 33109.6 | 15079.6 | 47492.7 | 1424.71 | 5194.82 | × | × | 50117.1 | × | × | 1296.78 | × | × | × | × | × | × | × | × | × | × | × | × | × | × | × | × | × | × | × | × | 0 | × | × |
| Byonic | 1208 | 3527 | P11717 | IGF2R | NGSK | N1(HexNAc(5)Hex(6)Fuc(3)NeuAc(1)500.137736) | 543 | 5 | 6 | 3 | 1 | 0 | 2 | 5:6:3:1:0:2 | complex | 1 | 0 | 0 | 90114.2 | 4544.01 | 57792.2 | 25520.8 | 63066.6 | × | 1220.72 | × | × | 88995.6 | × | × | × | × | × | × | × | × | × | × | × | × | × | × | × | × | × | × | × | × | × | × | × | 0 | × | × |
| Byonic | 1209 | 3593 | P11717 | IGF2R | NGSK | N1(HexNAc(6)Hex(7)Fuc(1)NeuAc(2)500.137736) | 543 | 6 | 7 | 1 | 2 | 0 | 2 | 6:7:1:2:0:2 | complex | 1 | 0 | 0 | 95705.3 | 502.833 | 59721 | 27100 | 70165.8 | × | 3245.58 | × | × | 96464.6 | × | × | × | × | × | × | × | × | × | × | × | × | × | × | × | × | × | × | × | × | × | × | × | 0 | × | × |
| Byonic | 1294 | 18793 | O00754 | MAN2B1 | PTWKLNQTEPVAGNYYPVNTR | N6(HexNAc(6)Hex(6)Fuc(3)NeuAc(1)NeuGc(1)250.068868) | 766 | 6 | 6 | 3 | 1 | 1 | 1 | 6:6:3:1:1:1 | complex | 1 | 0 | 0 | 112809 | 8865.87 | 1972.86 | 45998.3 | 218206 | 52190.1 | 159906 | 1504.13 | 8083.41 | 137878 | × | × | 30428.3 | × | × | 824.324 | 5308.39 | 2936.67 | × | × | × | × | 1504.13 | × | × | × | × | × | × | × | × | × | × | 0 | × | × |
| Byonic | 1303 | 4976 | Q92896 | GLG1 | NDTLQEAK | N1(HexNAc(5)Hex(5)Fuc(1)250.068868) | 786 | 5 | 5 | 1 | 0 | 0 | 1 | 5:5:1:0:0:1 | complex | 1 | 0 | 0 | 392490 | 16197.5 | 276472 | 7165.42 | 516534 | 46392.3 | 147494 | × | × | 419335 | × | × | 31752.8 | × | × | × | × | × | × | × | × | × | × | × | 58531.4 | 102487 | 32716.7 | × | × | × | × | × | × | 1 | y4, y6, y8 | × |
| Byonic | 1310 | 17847 | Q8TEM1 | NUP210 | FDNFSSLSIQWESTRPVLASIEPELPMQLVSQDDESGQK | N3(HexNAc(6)Hex(6)250.068868); M27(Oxidation) | 801 | 6 | 6 | 0 | 0 | 0 | 1 | 6:6:0:0:0:1 | complex | 1 | 0 | 0 | 73694.3 | 6174.08 | 42872 | 991.283 | 91246.3 | 3293.13 | 6026.23 | × | × | 78469.1 | × | × | 4283.54 | × | × | × | × | × | × | × | 1272.35 | × | × | × | × | × | × | × | × | × | × | × | × | 0 | × | × |
| Byonic | 1348 | 6131 | A2VDJ0 | TMEM131L | NLTLPK | N1(HexNAc(5)Hex(6)Fuc(2)NeuAc(1)250.068868) | 1045 | 5 | 6 | 2 | 1 | 0 | 1 | 5:6:2:1:0:1 | complex | 1 | 0 | 0 | 142851 | 13576.3 | 90711.6 | 2598.51 | 177885 | 2132.87 | 10382.5 | × | × | 134290 | × | × | × | × | × | × | × | × | × | × | × | × | × | × | × | × | × | × | × | × | × | × | × | 0 | y2 | b4 |
| Byonic | 1349 | 5166 | A2VDJ0 | TMEM131L | NLTLPK | N1(HexNAc(6)Hex(7)Fuc(3)NeuAc(1)NeuGc(1)250.068868) | 1045 | 6 | 7 | 3 | 1 | 1 | 1 | 6:7:3:1:1:1 | complex | 1 | 0 | 0 | 76361.8 | 4152.1 | 52726.6 | 18540.7 | 66343.3 | 1540.39 | 4521.19 | × | × | 108676 | 1041.66 | × | 3109.27 | × | × | × | × | × | × | × | 854.459 | × | × | × | × | × | × | × | × | × | × | × | × | 0 | y1 | b4 |
| Byonic | 1354 | 7623 | Q9BQG0 | MYBBP1A | VPAQANGTPTTK | N6(HexNAc(3)Hex(4)250.068868) | 1225 | 3 | 4 | 0 | 0 | 0 | 1 | 3:4:0:0:0:1 | hybrid | 1 | 0 | 0 | 164982 | 13754.4 | 4223.68 | 61025.1 | 300870 | × | 10128.8 | × | × | 103674 | × | × | × | × | × | × | × | × | × | × | 2592.06 | × | × | × | × | × | × | × | 14638.2 | × | × | × | × | 1 | y1, y2, y3, y7, y9, y10 | b2, b5, b7, b8 |
| Byonic | 1362 | 12116 | Q8IWZ3 | ANKHD1 | NASILLK | N1(HexNAc(6)Hex(7)Fuc(3)NeuAc(1)500.137736) | 1427 | 6 | 7 | 3 | 1 | 0 | 2 | 6:7:3:1:0:2 | complex | 1 | 0 | 0 | 223373 | 18179.1 | 4808.55 | 78981.5 | 290710 | 10094.3 | 20525.2 | × | × | 147103 | × | × | 3774.86 | × | × | × | × | × | × | × | × | × | × | × | × | × | × | × | × | × | × | × | × | 0 | × | × |
| pGlyco3 | 53 | 4428 | P01871 | IGHM | NNSDISSTR |  | 46 | 4 | 5 | 1 | 0 | 0 | 1 | 4:5:1:0:0:1 | complex | 0 | 1 | 0 | 160598 | 9250.09 | 130121 | 58227.5 | 319603 | 50614 | 91913.4 | × | × | 300568 | 3843.46 | × | 49004.2 | × | × | × | 1223.64 | 5640.18 | × | × | × | × | × | × | 19069 | 77229.4 | 69704.8 | × | × | × | × | × | × | 1 | y1, y3, y4, y5, y6, y7, y8, y9 | b8 |
| pGlyco3 | 326 | 6893 | P08575 | PTPRC | NGSAAMCHFTTK | 7,Carbamidomethyl[C]; | 470 | 4 | 5 | 0 | 0 | 0 | 1 | 4:5:0:0:0:1 | complex | 0 | 1 | 0 | 168190 | 12580.5 | 115419 | 66823 | 300579 | 33823.9 | 123952 | × | 825.811 | 130942 | × | × | 8130.91 | × | × | × | 6144.97 | × | × | × | 3253.95 | × | × | × | 64321.8 | 47391.9 | × | 56288.9 | 96909.3 | × | × | × | × | 1 | y1, y2, y3, y4, y5, y6, y7, y8, y9, y10, y11, y1 | b2, b3, b5, b12 |
| pGlyco3 | 361 | 7089 | P25942 | CD40 | DLVVQQAGTNK |  | 180 | 4 | 5 | 1 | 0 | 0 | 1 | 4:5:1:0:0:1 | complex | 0 | 1 | 0 | 342614 | 24491.3 | 248959 | 149645 | 671910 | 47830.6 | 178263 | × | × | 364887 | 2267.51 | × | 15223.8 | × | × | × | 8311.43 | × | × | × | 1880.09 | × | × | × | 53157.4 | 252763 | 38531.4 | × | 10680.5 | × | × | × | × | 1 | y1, y4, y5, y11 | b2, b3, b4, b6, b7, b8, b9 |
| pGlyco3 | 375 | 7198 | Q7LGA3 | HS2ST1 | YHVLHINTTK |  | 108 | 5 | 6 | 0 | 0 | 0 | 1 | 5:6:0:0:0:1 | complex | 0 | 1 | 0 | 525065 | 39584.6 | 319476 | 182582 | 666482 | 44767.8 | 226290 | × | × | 294805 | × | × | 3693.63 | × | × | × | 11940.6 | × | × | × | 2649.39 | × | × | × | 34403.9 | 130641 | × | 3949.34 | 67273.7 | × | × | × | × | 1 | y1, y10 | b1, b2, b3, b4, b5, b6, b8 |
| pGlyco3 | 627 | 9410 | O94923 | GLCE | NLTEKPPHIEVYETAEDRDK |  | 225 | 5 | 6 | 0 | 0 | 0 | 1 | 5:6:0:0:0:1 | complex | 0 | 1 | 0 | 245609 | 17076.3 | 160396 | 86560.8 | 347787 | 31113.2 | 97521.4 | × | × | 210140 | × | × | 10566.5 | × | × | × | 4123.52 | × | × | × | 994.254 | × | × | × | × | × | × | 51573.2 | 60558.9 | × | 14392 | 41945.7 | × | 1 | y1, y3, y4, y5, y6, y7, y8, y9, y10, y11, y12, y1 | b2, b3, b4, b5 |
| pGlyco3 | 672 | 9746 | O00754 | MAN2B1 | WWHQQTNATQEVVR |  | 133 | 4 | 4 | 0 | 0 | 0 | 1 | 4:4:0:0:0:1 | complex | 0 | 1 | 0 | 204033 | 13091.6 | 135158 | 70866.9 | 297673 | 6093.43 | 15462 | 13594 | 2145.72 | 150066 | × | × | 2021.96 | 7683.12 | × | × | 945.43 | × | × | × | 1526.87 | 1489.38 | 13594 | × | 5277.69 | 15982.5 | × | 15668.6 | 62403.3 | 7985.08 | × | × | × | 1 | y1, y2, y3, y4, y5, y6, y7, y8, y9, y1 | b1, b3, b4, b5 |
| pGlyco3 | 946 | 12784 | P0DOX6 |  | GLTFQQNASSMCVPDQDTAIR | 12,Carbamidomethyl[C]; | 332 | 4 | 5 | 1 | 0 | 0 | 1 | 4:5:1:0:0:1 | complex | 0 | 1 | 1 | 313090 | 16703.5 | 196891 | 82465.7 | 331160 | 4248.93 | 11554.9 | × | × | 340755 | 5566.66 | 1289.25 | 4214.71 | × | 10221.3 | × | × | × | × | × | 938.252 | × | × | × | × | × | × | 12808.2 | 33295.8 | 2201.14 | × | × | × | 1 | × | × |
| pGlyco3 | 950 | 12800 | P0DOX6 |  | GLTFQQNASSMCVPDQDTAIR | 12,Carbamidomethyl[C]; | 332 | 5 | 5 | 0 | 0 | 0 | 1 | 5:5:0:0:0:1 | complex | 0 | 1 | 0 | 806410 | 51168.1 | 529264 | 221710 | 1060440 | 81255.9 | 171826 | 18285.9 | 50114.2 | 1017490 | 14351.4 | × | 73449.1 | 24337.7 | × | × | 7579.65 | × | × | × | × | × | 18285.9 | × | × | × | × | 43951.7 | 108956 | × | × | × | × | 1 | × | × |
| pGlyco3 | 956 | 12873 | Q08722 | CD47 | DIYTFDGALNK |  | 73 | 4 | 5 | 1 | 0 | 0 | 1 | 4:5:1:0:0:1 | complex | 0 | 1 | 0 | 369568 | 34547.5 | 249898 | 141503 | 680449 | 54317 | 198101 | × | × | 326999 | 2132.63 | × | 17027 | × | × | × | 7745.34 | 2486.24 | × | × | 3315.87 | × | × | × | 62400.7 | 241698 | 36786.4 | × | 3226.2 | × | × | × | × | 1 | × | × |
| pGlyco3 | 1015 | 14222 | P40259 | CD79B | MEESQNESLATLTIQGIR |  | 101 | 4 | 5 | 1 | 0 | 0 | 1 | 4:5:1:0:0:1 | complex | 0 | 1 | 0 | 205771 | 18718.6 | 133695 | 76500.1 | 336123 | 1151.45 | 126167 | × | × | 155351 | × | × | 10738.6 | × | × | × | 6109.34 | × | × | × | 2541.95 | × | × | × | × | × | × | 17086.2 | 43080 | × | × | × | × | 1 | × | × |
| pGlyco3 | 1019 | 14282 | Q9BU23 | LMF2 | PGNLSRPPPVVVPHQPR |  | 489 | 3 | 5 | 0 | 0 | 0 | 1 | 3:5:0:0:0:1 | hybrid | 0 | 1 | 0 | 58783.9 | 3788.4 | 25740.3 | 18065.3 | 58163.5 | × | 777.391 | × | 10554.2 | 11737.8 | × | × | × | × | × | × | × | × | × | 1551.93 | 1208.78 | × | × | × | × | × | × | 11281.9 | 32011.4 | 6231.74 | × | × | × | 1 | × | × |
| pGlyco3 | 1115 | 16971 | O96005 | CLPTM1 | AEDYGPVEVISHWHPNITINIVDDHTPWVK |  | 241 | 6 | 5 | 2 | 0 | 0 | 1 | 6:5:2:0:0:1 | complex | 0 | 1 | 0 | 33638.2 | 1014.32 | 24181.8 | 748.627 | 59373.6 | × | 1772.18 | × | × | 23745.3 | × | × | × | × | × | × | × | × | × | × | 4181.77 | × | × | × | × | × | × | × | 2965.89 | × | 1917 | 16096.3 | × | 1 | × | × |
| pGlyco3 | 1127 | 17483 | P08195 | SLC3A2 | DASSFLAEWQNITK |  | 365 | 4 | 5 | 1 | 0 | 0 | 1 | 4:5:1:0:0:1 | complex | 0 | 1 | 0 | 253562 | 19270.6 | 170445 | 83651.2 | 410306 | 32884.7 | 103335 | × | × | 242017 | 2031.67 | × | 15253.6 | × | × | × | 3028.03 | × | × | × | 2555.06 | × | × | × | 18807.9 | 75173.1 | 11178.1 | 3210.77 | 14260.8 | × | × | × | × | 1 | × | × |
| pGlyco3 | 1143 | 19385 | P08195 | SLC3A2 | LLIAGTNSSDLQQILSLLESNK |  | 381 | 6 | 7 | 0 | 0 | 0 | 1 | 6:7:0:0:0:1 | complex | 0 | 1 | 1 | 246087 | 11059 | 170654 | 63458.2 | 236963 | 18398 | 50483.6 | × | × | 318989 | × | 1170.74 | 9782.22 | × | 6146.72 | × | 1516.72 | × | × | × | × | × | × | × | × | × | × | 4113.51 | 12728.2 | × | × | × | × | 1 | × | × |
| GRable | 42 | 12402 | P08575 | PTPRC | SFHNFTLCYIK | 0.00020000000.0 | 421 | 5 | 5 | 1 | 0 | 0 | 1 | 5:5:1:0:0:1 | complex | 0 | 0 | 1 | 1734950 | 78959.7 | 1232810 | 487778 | 1779590 | 52113.7 | 131099 | × | × | 1844760 | 18760.4 | × | 36332.8 | × | 21914.6 | × | × | × | × | × | × | × | × | × | 52785.9 | 150797 | 69654 | 20953 | 107103 | 8097.52 | × | × | × | 1 | × | × |
| GRable | 375 | 4511 | P01871 | IGHM | YKNNSDISSTR | 0.00200000000.0 | 46 | 4 | 6 | 1 | 0 | 0 | 1 | 4:6:1:0:0:1 | hybrid | 0 | 0 | 1 | 351767 | 27271.6 | 240472 | 133256 | 568601 | × | 3154.87 | × | × | 347836 | × | 2210.56 | × | × | 9498.69 | × | × | × | × | × | × | × | × | × | 19569 | 81678.2 | 44749.9 | 4760.75 | 44585.3 | 4289.08 | × | × | × | 1 | y1, y4, y5, y7, y9, y11 | b2, b10 |
| GRable | 379 | 4526 | P01871 | IGHM | YKNNSDISSTR | 0.00200000000.0 | 46 | 4 | 7 | 1 | 0 | 0 | 1 | 4:7:1:0:0:1 | hybrid | 0 | 0 | 1 | 392918 | 20792.6 | 292407 | 128926 | 492911 | 2010.71 | 9108.8 | × | × | 486096 | × | 3106.82 | × | × | 13199.4 | × | × | × | × | × | × | × | × | × | 14208.8 | 62554.4 | 39631.7 | 7008.38 | 30591.4 | 3325.14 | × | × | × | 1 | y1, y3, y5, y7, y8, y11 | b2, b10 |
