## Supplementary material for "Comparative Evaluation of Glycoproteomics Software for Rare Glycopeptide Identification": Table S4

| Peptide sequence | Peptide length | Position of N-glycosylation site from the N-terminus | Position of N-glycosylation site from the C-terminus | At which position from the termini is the glycan attached? |
|---|---|---|---|---|
| FNLSK | 5 | 2 | 4 | 2 |
| AFMNR | 5 | 4 | 2 | 2 |
| YHINK | 5 | 4 | 2 | 2 |
| VNFTR | 5 | 2 | 4 | 2 |
| NVTFR | 5 | 1 | 5 | 1 |
| FNFSK | 5 | 2 | 4 | 2 |
| FNISK | 5 | 2 | 4 | 2 |
| YETTNK | 6 | 5 | 2 | 2 |
| NYTLWR | 6 | 1 | 6 | 1 |
| ANFSIK | 6 | 2 | 5 | 2 |
| FNHTDR | 6 | 2 | 5 | 2 |
| FTNASK | 6 | 3 | 4 | 3 |
| NFTMNEK | 7 | 1 | 7 | 1 |
| NSTAYFR | 7 | 1 | 7 | 1 |
| TDNGTYR | 7 | 3 | 5 | 3 |
| NATGDYK | 7 | 1 | 7 | 1 |
| MFSNCSK | 7 | 4 | 4 | 4 |
| FTPVHNR | 7 | 6 | 2 | 2 |
| NLSISTK | 7 | 1 | 7 | 1 |
| IADNHTPK | 8 | 4 | 5 | 4 |
| VKPQWFNR | 8 | 7 | 2 | 2 |
| LVPPGGNK | 8 | 7 | 2 | 2 |
| IHLNVSER | 8 | 4 | 5 | 4 |
| NGSLFAFR | 8 | 1 | 8 | 1 |
| NASNMEYR | 8 | 1 | 8 | 1 |
| TPEFHPNK | 8 | 7 | 2 | 2 |
| KDNTTVTR | 8 | 3 | 6 | 3 |
| HNSTGCLR | 8 | 2 | 7 | 2 |
| FVLQNASR | 8 | 5 | 4 | 4 |
| NISQVLEK | 8 | 1 | 8 | 1 |
| NDTLQEAK | 8 | 1 | 8 | 1 |
| ENNNITMR | 8 | 4 | 5 | 4 |
| DNNSIITR | 8 | 2 | 7 | 2 |
| PVNVCVGK | 8 | 7 | 2 | 2 |
| YNVSSLEK | 8 | 2 | 7 | 2 |
| PFNLSQGK | 8 | 3 | 6 | 3 |
| NNSDISSTR | 9 | 1 | 9 | 1 |
| NFIINMTCR | 9 | 5 | 5 | 5 |
| DHVHCLGNR | 9 | 8 | 2 | 2 |
| IIFANVSVR | 9 | 5 | 5 | 5 |
| KNASNMEYR | 9 | 2 | 8 | 2 |
| LVNVTIHFR | 9 | 3 | 7 | 3 |
| LIDNNKTEK | 9 | 5 | 5 | 5 |
| NATLAEQAK | 9 | 1 | 9 | 1 |
| NLTVSECKK | 9 | 1 | 9 | 1 |
| ADSASCENR | 9 | 8 | 2 | 2 |
| VNSSLLPPK | 9 | 2 | 8 | 2 |
| NYSVLYFQQK | 10 | 1 | 10 | 1 |
| IINYTPDMAR | 10 | 3 | 8 | 3 |
| MAAALNATGR | 10 | 6 | 5 | 5 |
| GSLSYLNVTR | 10 | 7 | 4 | 4 |
| EADNHTAFIR | 10 | 4 | 7 | 4 |
| SYHNQTISFR | 10 | 4 | 7 | 4 |
| FINYNQTVSR | 10 | 5 | 6 | 5 |
| GPTNNTCVVR | 10 | 4 | 7 | 4 |
| NLCSLKTPGK | 10 | 9 | 2 | 2 |
| VLEVTNSSLR | 10 | 6 | 5 | 5 |
| SINGSLYIFR | 10 | 3 | 8 | 3 |
| HVVYWNSSNPR | 11 | 6 | 6 | 6 |
| GNWTTNEMEVK | 11 | 2 | 10 | 2 |
| SECHFFNGTER | 11 | 7 | 5 | 5 |
| QECYAFNGTQR | 11 | 7 | 5 | 5 |
| GNVTQHVQGHR | 11 | 2 | 10 | 2 |
| IHNVTASDSGK | 11 | 3 | 9 | 3 |
| NLTLTDLQSPK | 11 | 1 | 11 | 1 |
| KNCTYNQVQTR | 11 | 2 | 10 | 2 |
| GELNTSIFSSR | 11 | 4 | 8 | 4 |
| AYNEAEVLYNR | 11 | 10 | 2 | 2 |
| LLLSINVTNTR | 11 | 6 | 6 | 6 |
| NLSAPVTLNLR | 11 | 1 | 11 | 1 |
| IQRMNSSFSVK | 11 | 5 | 7 | 5 |
| AHNDHPNNFSR | 11 | 8 | 4 | 4 |
| CENLSTPTMLK | 11 | 3 | 9 | 3 |
| NCSPWLSCEELR | 12 | 1 | 12 | 1 |
| ATLSTLAVAVNR | 12 | 11 | 2 | 2 |
| DLVVQQLVNVSR | 12 | 9 | 4 | 4 |
| NINCSIEESMGK | 12 | 3 | 10 | 3 |
| TVYNNNSSRFGK | 12 | 5 | 8 | 5 |
| EIANATAKPEDR | 12 | 4 | 9 | 4 |
| IFIFNQTGIEAK | 12 | 5 | 8 | 5 |
| EIVCNVTLGGER | 12 | 5 | 8 | 5 |
| ELSLLSGEVCNR | 12 | 11 | 2 | 2 |
| KLLLSINVTNTR | 12 | 7 | 6 | 6 |
| ALTNGSLPAGTR | 12 | 4 | 9 | 4 |
| LHVTLYNCSFGR | 12 | 7 | 6 | 6 |
| PAHNSTDLDPFK | 12 | 4 | 9 | 4 |
| PHSVSLNDTETR | 12 | 7 | 6 | 6 |
| KWGHNITEFQQR | 12 | 5 | 8 | 5 |

| | | | | |
|---|---|---|---|---|
| TEPSFTKENSSK | 12 | 9 | 4 | 4 |
| VNFTLEASEGCYR | 13 | 2 | 12 | 2 |
| GRDIYTFDGALNK | 13 | 12 | 2 | 2 |
| AYWPDVIHSFPNR | 13 | 12 | 2 | 2 |
| NCTITANAECACR | 13 | 1 | 13 | 1 |
| LNNITIIGPLDMK | 13 | 3 | 11 | 3 |
| NHSIFLADINQER | 13 | 1 | 13 | 1 |
| PQDHGTNLTCQVK | 13 | 7 | 7 | 7 |
| GETASLLCNISVR | 13 | 9 | 5 | 5 |
| KQNNGAFNETLFR | 13 | 8 | 6 | 6 |
| RDENESPFPDIPK | 13 | 4 | 10 | 4 |
| QFEAQNLSMQSVR | 13 | 6 | 8 | 6 |
| IAPASNVSHTVVLR | 14 | 6 | 9 | 6 |
| LNDTYVNVGLYSTK | 14 | 2 | 13 | 2 |
| PLGLANDTDHYFLR | 14 | 6 | 9 | 6 |
| KNLTLMATTSQLPK | 14 | 2 | 13 | 2 |
| VCERENSTVPWFVK | 14 | 6 | 9 | 6 |
| RNQSLVQDMDSMVR | 14 | 2 | 13 | 2 |
| CHEGNGTFECGACR | 14 | 5 | 10 | 5 |
| LNFTLVGKPLLAFR | 14 | 2 | 13 | 2 |
| NGSSIVDLSPLIHR | 14 | 1 | 14 | 1 |
| SGAEAQTPEDSPNR | 14 | 13 | 2 | 2 |
| QPGENGSIVTCGHR | 14 | 5 | 10 | 5 |
| LWLPVNLTWADLEDR | 15 | 6 | 10 | 6 |
| HPDAVAWANLTNAIR | 15 | 9 | 7 | 7 |
| YELSLHPNLTSMTFR | 15 | 8 | 8 | 8 |
| DLGPTLANSTHHNVR | 15 | 8 | 8 | 8 |
| VTLWVHPFVNYNSSR | 15 | 12 | 4 | 4 |
| VFHIHNESWVLLTPK | 15 | 6 | 10 | 6 |
| AINQTAVECTWTGPR | 15 | 3 | 13 | 3 |
| NTTQFAAVCPQHLDER | 16 | 1 | 16 | 1 |
| EKLNDTYVNVGLYSTK | 16 | 4 | 13 | 4 |
| QSTHSIYMFFNTSELR | 16 | 11 | 6 | 6 |
| QFCVHVNNSNLNYFQK | 16 | 7 | 10 | 7 |
| NYTLLQTIPPFERPFK | 16 | 1 | 16 | 1 |
| EDNPDKNPEAPLNVSR | 16 | 13 | 4 | 4 |
| GHTNGTKPLDGFDVWK | 16 | 4 | 13 | 4 |
| TILVDNNTWNNTHISR | 16 | 6 | 11 | 6 |
| RFDNFSSLSIQWESTR | 16 | 4 | 13 | 4 |
| TSSDDPNDSGELTQEK | 16 | 7 | 10 | 7 |
| AHTPPGNAEVTTNIPK | 16 | 15 | 2 | 2 |
| LNITNIWVLDYFGGPK | 16 | 2 | 15 | 2 |
| SCVAVTSAQPQNMSRR | 16 | 12 | 5 | 5 |
| IEVLVSNATQFIILHSK | 17 | 7 | 11 | 7 |
| FNVTGTPEQYVPYSTTR | 17 | 2 | 16 | 2 |
| FAEAACDVVHVMLNGSR | 17 | 14 | 4 | 4 |
| TYAIYDLLDTAMINNSR | 17 | 14 | 4 | 4 |
| LIAEPGLMLNFSATALR | 17 | 10 | 8 | 8 |
| VMSWWDYGYQIAGMANR | 17 | 16 | 2 | 2 |
| VVSGLFRSLQNDSVKVR | 17 | 11 | 7 | 7 |
| YSLNVTYNYPVHYFDGR | 17 | 4 | 14 | 4 |
| QCNGTSMCWCVNTAGVR | 17 | 3 | 15 | 3 |
| LWLPVNLTWADLEDRDGR | 18 | 6 | 13 | 6 |
| VTLYLRPGQAAAFNVTFR | 18 | 14 | 5 | 5 |
| PTCDEKYANITVDYLYNK | 18 | 9 | 10 | 9 |
| NESDKAPWPLSLPGCPHR | 18 | 1 | 18 | 1 |
| QDQQLQNCTEPGEQPSPK | 18 | 7 | 12 | 7 |
| KTYAIYDLLDTAMINNSR | 18 | 15 | 4 | 4 |
| APPRNYSVIVMFTALQLHR | 19 | 5 | 15 | 5 |
| SSCGKENTSDPSLVIAFGR | 19 | 7 | 13 | 7 |
| TALWVATDHNMDNTSTVLR | 19 | 13 | 7 | 7 |
| SDAVSHTGNYTCEVTELTR | 19 | 9 | 11 | 9 |
| VTTYCNETMTGWVHDVLGR | 19 | 6 | 14 | 6 |
| GHAHLAALVNHESYNFSHR | 19 | 15 | 5 | 5 |
| AEPGTHLCIDVEDAMNITR | 19 | 16 | 4 | 4 |
| TNITLVCKPGDLESAPVLR | 19 | 2 | 18 | 2 |
| KRWTQFVNVTFQMEPTITR | 19 | 8 | 12 | 8 |
| NLSHLPTFSSPAIESHIHR | 19 | 1 | 19 | 1 |
| PAAVVSEACDADDADNASK | 19 | 16 | 4 | 4 |
| AAQTGAAGVRLKEGCNINR | 19 | 18 | 2 | 2 |
| LGDCISEDSYPDGNITWYR | 19 | 14 | 6 | 6 |
| ELLLDFANSSAELTGCLVR | 19 | 8 | 12 | 8 |
| SNAPLVNVTLYYEALCGGCR | 20 | 7 | 14 | 7 |
| CKPNFFCNSTVCEHCDPCTK | 20 | 8 | 13 | 8 |
| TELDMQPQGLGLFVNTSAPR | 20 | 15 | 6 | 6 |
| LNVSHTGVPLGEEYILVFSR | 20 | 2 | 19 | 2 |
| EGDTVTLSCNYNSSNPSVTR | 20 | 12 | 9 | 9 |
| NLTEKPPHIEVYETAEDRDK | 20 | 1 | 20 | 1 |
| VNNSGISLCNLISAVTTPAK | 20 | 2 | 19 | 2 |
| NCSAAPQPEPAAGLASYPELR | 21 | 1 | 21 | 1 |
| FTFTSHTPGDHQICLHSNSTR | 21 | 18 | 4 | 4 |
| VWDTAVGLNHTAEPSLQTLER | 21 | 9 | 13 | 9 |
| DSLACFNQTYTINLYLVETGR | 21 | 7 | 15 | 7 |
| LQFYIGEHLLPYNMTVYQAVR | 21 | 13 | 9 | 9 |
| ENSSEICSNNGECVCGQCVCR | 21 | 2 | 20 | 2 |
| HHYAGMVSLMDEAVGNVTAALK | 22 | 16 | 7 | 7 |
| TNHTVMTMGSDFQYENANMWFK | 22 | 2 | 21 | 2 |
| DHFTFCQQLNISICPLSQTAAR | 22 | 10 | 13 | 10 |
| KENSSEICSNNGECVCGQCVCR | 22 | 3 | 20 | 3 |

| | | | | |
|---|---|---|---|---|
| DLGPEYEGIFNTSLQWILENGK | 22 | 11 | 12 | 11 |
| VDGFEKRAAASESNNYMNHVAK | 22 | 21 | 2 | 2 |
| PYTLEEQKNLTVCPDGALFEQK | 22 | 9 | 14 | 9 |
| FWLPENVSWADLEGPADGYGYPR | 23 | 6 | 18 | 6 |
| YNCSTQHADLTIIDNIEEMNFLR | 23 | 2 | 22 | 2 |
| DSSGNETHFTGNEVGFFKPISCR | 23 | 5 | 19 | 5 |
| NETQHEPYPLMLPGCSPSCPLER | 23 | 1 | 23 | 1 |
| TSQENISFETMYDVLSTKPVLNK | 23 | 5 | 19 | 5 |
| AHGGSGNFSWSSSSHLVATVTVK | 23 | 7 | 17 | 7 |
| CVGFGPEEELTNITDVQFLQSTR | 23 | 12 | 12 | 12 |
| SNHTIWFGHFTTSTILSPSPGIR | 23 | 2 | 22 | 2 |
| LQQFSPLSDYEGQEEEMNGTKMK | 23 | 18 | 6 | 6 |
| MMETLEHLNTYLCHDNLSSNDTK | 23 | 16 | 8 | 8 |
| KRHFIGNTSMDAGLSFIMANHGK | 23 | 7 | 17 | 7 |
| ENCSPTIPASNTADEEYEDGIER | 23 | 2 | 22 | 2 |
| TLMLNVSLFNAGDDAYETTLHVK | 23 | 5 | 19 | 5 |
| GPIGPGEPLELLCNVSGALPPAGR | 24 | 14 | 11 | 11 |
| VTNSNANAASPLIVAGYNVSGSVR | 24 | 18 | 7 | 7 |
| FVMVKFLNDSIVDPVDSEWFGFYR | 24 | 8 | 17 | 8 |
| LVSNASMLVSMHGAQLVTTLFLPR | 24 | 4 | 21 | 4 |
| SLDAYPILNQAQALENHTEVQFQK | 24 | 16 | 9 | 9 |
| FFQNVTAEDTDSIGLALFAPHSSR | 24 | 4 | 21 | 4 |
| LFDDWMASNGTQSLPTDVMTTVFK | 24 | 9 | 16 | 9 |
| LEHFAPPFGFMELNYSLVQKVVTR | 24 | 14 | 11 | 11 |
| GIAVEVQDTPNACATFNFLCHEGR | 24 | 23 | 2 | 2 |
| DGGNQSEGSEFLLLGMSESPEQQR | 24 | 4 | 21 | 4 |
| CLYGEAYPACSPGNTSTAEEELCR | 24 | 14 | 11 | 11 |
| YPQDYQFYIQNFTALPLNTVVPPQR | 25 | 11 | 15 | 11 |
| GTEVNTTVIGENDPIDEVQGFLFGK | 25 | 5 | 21 | 5 |
| LCHTHGWNETSEFMPQGAVFSCLYR | 25 | 8 | 18 | 8 |
| GTVFFDEFTFVKLTGVAGNYTVCQK | 25 | 19 | 7 | 7 |
| NISSHTNIVFSNGELDPWSGGGVTK | 25 | 1 | 25 | 1 |
| GPLKKSNAPLVNVTLYYEALCGGCR | 25 | 12 | 14 | 12 |
| PAANLVCLSGMNGSTLWSSLLPEEAR | 26 | 12 | 15 | 12 |
| CFVCPVEYNNDTNSFTVDCEPSDLFR | 26 | 10 | 17 | 10 |
| LNFSLVGTPLSAFGNLRPVLAEDAQR | 26 | 2 | 25 | 2 |
| SVNLFGSGWNVSAMYSSSHPPAEHKK | 26 | 10 | 17 | 10 |
| GNYDFVQAMIVNNHTSLDVEEAAAVR | 26 | 13 | 14 | 13 |
| CDTPANCTYLDLLGTWVFQVGSSGSQR | 27 | 6 | 22 | 6 |
| SQNRPQGQSTQPSNAAGTNTTSASTPR | 27 | 19 | 9 | 9 |
| ITIAINNTLTPTTLPPGTIQYLTDTSK | 27 | 6 | 22 | 6 |
| CDGWADCTDHSDELNCSCDAGHQFTCK | 27 | 15 | 13 | 13 |
| VVFLSPAVPEEPEAYNLTVLIEMDGHR | 27 | 16 | 12 | 12 |
| KNATQPPNAEEESGSSSASEEEDTKPK | 27 | 2 | 26 | 2 |
| QTVHFQISSQLQFSPEEVLGMVLNYSR | 27 | 24 | 4 | 4 |
| MGLLDSEPGSVLNVVSTALNDTVEFYR | 27 | 20 | 8 | 8 |
| LPSVVIPLHYDLFVHPNLTSLDFVASEK | 28 | 17 | 12 | 12 |
| LCENNAEFNKNCSKSHSGIDCSYVECCR | 28 | 11 | 18 | 11 |
| GHHVSQEPWNSSITLTSQAGAVFQSFAK | 28 | 10 | 19 | 10 |
| NQTNTFFGSPPAATEATHVVSTIPESLQ | 28 | 1 | 28 | 1 |
| VLASVVGTTGVSLNLTQLPPFNVHLQTK | 28 | 14 | 15 | 14 |
| SIFDPPVFPVCMLGNRTLSRDQFDICAK | 28 | 15 | 14 | 14 |
| NVSEVSVEYLCSQPCVVNLEAVVSSEFR | 28 | 1 | 28 | 1 |
| NPNVSDSLLLQNIFQALNVYYNYSGQVK | 28 | 3 | 26 | 3 |
| TIQNSSVSPTSSSSSSSSTGETQTQSSSR | 29 | 4 | 26 | 4 |
| DSGIWINGFDYTGISHITPHIPEINDSIR | 29 | 25 | 5 | 5 |
| LETLSLVSCFANNTPLEHLDLSQNLLQHK | 29 | 12 | 18 | 12 |
| TKPFPWGDGNHTLFHNPHVNPLPTGYEDE | 29 | 10 | 20 | 10 |
| ALMVLTEEPLLYIPPPPCQPLINTTESLR | 29 | 23 | 7 | 7 |
| DINTTAQNIMFYDMGSGSTVCTIVTYQMVK | 30 | 3 | 28 | 3 |
| AEDYGPVEVISHWHPNITINIVDDHTPWVK | 30 | 16 | 15 | 15 |
| SSGLWNNTVFIFSTDNGGQTLAGGNNWPLR | 30 | 6 | 25 | 6 |
| DSLNSSHPISSSVQSSEQIEEMFDSLSYFK | 30 | 4 | 27 | 4 |
| KTVTTVETQNLEGLHHDGQFCHKPCPPGER | 30 | 29 | 2 | 2 |
| EYLCVKAMILLNSSMYPLVTATQDADSSRK | 30 | 12 | 19 | 12 |
| VQSIQLSQMQAETTNTSLLQVQLECSLHNK | 30 | 15 | 16 | 15 |
| KMSNITFLNFDPPIEEFHQYYQHIVTTLVK | 30 | 4 | 27 | 4 |
| GDDLYTNVTISLVESLVGFEMDITHLDGHK | 30 | 7 | 24 | 7 |
| NLTCSVSWACEQGTPPIFSWLSAAPTSLGPR | 31 | 1 | 31 | 1 |
| RAEDYGPVEVISHWHPNITINIVDDHTPWVK | 31 | 17 | 15 | 15 |
| EAKKETGFVESAESDHMAIPGGNQSVLAPSR | 31 | 23 | 9 | 9 |
| EDKPLSLSMPVITEEEENESLSGTEFTASKK | 31 | 18 | 14 | 14 |
| VPASLMVSLGEDAHFQCPHNSSNNANVTWWR | 31 | 20 | 12 | 12 |
| DLQAPLGQSFSCSNSSIILSPAVHLDLLSLR | 31 | 14 | 18 | 14 |
| RGDDLYTNVTISLVESLVGFEMDITHLDGHK | 31 | 8 | 24 | 8 |
| DSLNQSLSQDLTMAPGSTLWLSCGVPPDSVSR | 32 | 4 | 29 | 4 |
| YNQETPMEICLNGTPALAYLASAPPPLCPSGR | 32 | 12 | 21 | 12 |
| HSTINIVDIYPMMCHILGLKPHPNNGTFGHTK | 32 | 25 | 8 | 8 |
| EVHLVQFHEPDIYNYSALLLSEDKDTLYIGAR | 32 | 14 | 19 | 14 |
| PAFPNSSWLPFHERLQVQNGECPWQVSIQMSR | 32 | 5 | 28 | 5 |
| VNATNFQALAAEFGGESFTSTFQTQSPPSFYR | 32 | 2 | 31 | 2 |
| TVLENLGNGTSLPVLEGQSLCLVCVTHSSPPAR | 33 | 8 | 26 | 8 |
| WFGTDMQMINFTTGEFQLTEACPYLGTHSEESR | 33 | 10 | 24 | 10 |
| WGISFCNQTGVFNQGPHSPILSLMAQELGISEK | 33 | 7 | 27 | 7 |
| TPPSSNSCTASSSSFSSSGSSMSSSQNQHGSSK | 33 | 32 | 2 | 2 |
| AFDQIGGYGQLEAAYAQAIPSRTIANTTCHLPR | 33 | 26 | 8 | 8 |
| VCECNPNYTGSACDCSLDTSTCEASNGQICNGR | 33 | 7 | 27 | 7 |
| GWTGAAAGAGGATAAPPSGNQNGAEGDQINASK | 33 | 30 | 4 | 4 |
| CDENGNYLPLQCYGSIGYCWCVFPNGTEVPNTR | 33 | 25 | 9 | 9 |

| | | | | |
|---|---|---|---|---|
| EKPFEDFLPSHYNYLQFAYYNIGNYTQAVECAK | 33 | 24 | 10 | 10 |
| TGFQMWLEENRSNILSDNPDFSDEADIIKEGMIR | 34 | 10 | 25 | 10 |
| SPSAAWLLGAAILLAASLSCSGTIQGTNRSSKGR | 34 | 28 | 7 | 7 |
| PMNKSPMLITGIRCLYFAPPEFLLEVQCMHETQK | 34 | 3 | 32 | 3 |
| GRGWTGAAAGAGGATAAPPSGNQNGAEGDQINASK | 35 | 32 | 4 | 4 |
| IIAENIYEGGLNSDYVHLFVATLHFPHASNITLYK | 35 | 30 | 6 | 6 |
| KPQDQGLNHTCNYYLQPEEDVTIGVWHTVPAVWWK | 35 | 8 | 28 | 8 |
| NGSAVGVSRPAANLVCLSGMNGSTLWSSLLPEEAR | 35 | 1 | 35 | 1 |
| ITDTHTPSGLTVNLTLYYMLSCSPAPLLSPSLSHR | 35 | 13 | 23 | 13 |
| IQNVGWDNTTIACAACNSWCSWASPVALNVQYAPR | 35 | 8 | 28 | 8 |
| GHEVEDVDLELFNTSVQLQPPTTAPGPETAAFIER | 35 | 13 | 23 | 13 |
| GHDLNEDGLVSWEEYKNATYGYVLDDPDPDDGFNYK | 36 | 17 | 20 | 17 |
| NLPLGWDGKPIPYWLYKLHGLNINYNCEICGNYTYR | 36 | 32 | 5 | 5 |
| TQMNSFLLSTASQQEIATLDNKTMTDVVGNPEEERR | 36 | 21 | 16 | 16 |
| SIADFVSGAESRTYNLPFKWAGTNHVVYNGSLYFNK | 36 | 29 | 8 | 8 |
| GTILGLDLQNCSLEDPGPNFHQAHTTVIIDLQANPLK | 37 | 10 | 28 | 10 |
| ILTNNSQTPILSPQEVVSCSQYAQGCEGGFPYLIAGK | 37 | 4 | 34 | 4 |
| LNETQPIDLSGLPLQSSKNSYSFQNPSSFDPSMLLQR | 37 | 2 | 36 | 2 |
| LHHVEANRVNFVAAIDVYEDGEAGLLLCYNYSCIYKK | 37 | 30 | 8 | 8 |
| TILPAAAQDVYYRDEIGNVSTSHLLILDDSVEMEIRPR | 38 | 18 | 21 | 18 |
| NQTLAGEYFHKAAQGGHMEGTLWCSLYYITGNLETFPR | 38 | 1 | 38 | 1 |
| FPIVNAELSFFGHAHLGTGDPYTPGFPSFNHTQFPPSR | 38 | 30 | 9 | 9 |
| VHQESVHMIEVTSNLINETLANHPEWANWLPEAQVVQR | 38 | 17 | 22 | 17 |
| NVTLTLSWNVVPNAGILPLVTGSGHVSVPFPDTYEITK | 38 | 1 | 38 | 1 |
| VDFWHSDQCNMINGTSGQMWPPFMTPESSLEFYSPEACR | 39 | 13 | 27 | 13 |
| QCYQDHNLSQNGSAPTFPLCAMQLFSHMHAVISTATCMR | 39 | 7 | 33 | 7 |
| FDNFSSLSIQWESTRPVLASIEPELPMQLVSQDDESGQK | 39 | 3 | 37 | 3 |
| RFDNFSSLSIQWESTRPVLASIEPELPMQLVSQDDESGQK | 40 | 4 | 37 | 4 |
| HVFPHNHTADIQSEVHCIFSPQIEEPSQCPDCVVSALGAK | 40 | 6 | 35 | 6 |
| FDNFSSLSIQWESTRPVLASIEPELPMQLVSQDDESGQKK | 40 | 3 | 38 | 3 |
| GRGWTGAAAGAGGATAAPPSGNQNGAEGDQINASKNEEDAGK | 42 | 32 | 11 | 11 |
| SFLNGYNCTVLAYGATGAGKTHTMLGSADEPGVMYLTMLHLYK | 43 | 7 | 37 | 7 |
| SGEDAHEALLTLVVPPALLLSSVRPPGACQANETIFCELGNPFK | 44 | 32 | 13 | 13 |
| LDFVAPATGAHNYTLYFMSDAYMGCDQEYKFSVDVKEAETDSDSD | 45 | 12 | 34 | 12 |
| NTTALFSQFVSGNNRTAAPITLPIEEVLQTLQDLIAYFQPPEEEMR | 46 | 1 | 46 | 1 |
| ENGTDTVQEEEESPAEGSKDEPGEQVELKEEAEAPVEDGSQPPPPEPK | 48 | 2 | 47 | 2 |
| HDISESVNFDEETDGISQSACLERPNSASSQNSTFSSFCQKTEDTGYR | 48 | 32 | 17 | 17 |
| ALGFSEELIVELHTLCDCNCSDTQPQAPHCSDGQGHLQCGVCSCAPGR | 48 | 19 | 30 | 19 |
| APGYICHGLSPGGLSNGSREECYWGHVFWDQVSTVHPAPATQQATHGGPR | 50 | 16 | 35 | 16 |
| PGATVTLRCVGNGSVEWDGPPSPHWTLYSDGSSSILSTNNATFQNTGTYR | 50 | 12 | 39 | 12 |
| FSAPLFLSHPHFLNADPVLAEAVTGLHPNQEAHSLFLDIHPVTGIPMNCSVK | 52 | 48 | 5 | 5 |
